# Genomic SEM reveals shared genetic architecture and expands locus discovery for locomotor activity in heterogeneous stock rats

**DOI:** 10.64898/2026.10.01.756043

**Authors:** Apurva S. Chitre, Thiago Missfeldt Sanches, Gavrila Ang, Nana K. Amissah, Rodolfo Avila, Hannah Bimschleger, Paola Campo, Nazzareno Cannella, Denghui Chen, Riyan Cheng, Katarina A. Cohen, Ayteria D. Crow, Eric Dereschewitz, Michelle R. Doyle, Maya Eid, Anthony M. George, Aidan P. Horvath, Keita Ishiwari, Benjamin B. Johnson, Elaine Keung, Christopher King, Connor D. Martin, Angel Garcia Martinez, Daniel Munro, Alesa H. Netzley, Khai-Minh Nguyen, Beverly Peng, Dominika Pullmann, Sara R. M. U. Rahman, Analyse T. Roberts, Mohammad Sadegi, Deborah Sevigny-Resetco, Laura Soverchia, Celine L. St. Pierre, Nina Suzuki, Brady M. Thompson, Shambhavi Tyagi, Massimo Ubaldi, Robert M. Vogel, Tengfei Wang, Yizhi Wang, Hao Chen, Roberto Ciccocioppo, Giordano de Guglielmo, David Dietz, Shelly B. Flagel, Thomas Jhou, Jicai Jiang, Peter W. Kalivas, Brittany N. Kuhn, Paul J. Meyer, Suzanne H. Mitchell, Terry E. Robinson, Oksana Polesskaya, Abraham A. Palmer

## Abstract

Locomotor response to a novel environment is a heritable, addiction-relevant behavioral phenotype in rodents. We characterized its genetic basis in 9,341 male and female heterogeneous stock (HS) rats from eight studies. Locomotor activity was heritable across studies, while pairwise genetic correlations were positive but variable, indicating shared yet nonidentical genetic influences. Given nonuniform genetic architecture, we compared three complementary approaches to locus discovery that analyzed SNP effects 1) on a pooled locomotor phenotype (MEGA), 2) across studies under a fixed-effect model (METAL), or 3) on a shared genetic factor (Genomic SEM). To ensure comparability, all three locus-discovery approaches were applied to a common subset of seven studies. Together, these approaches identified 38 loci, with both convergence and method-specific discovery across analyses. Genomic SEM identified 31 loci, compared to 8 for MEGA and 13 for METAL. To interpret the loci, we integrated brain molecular QTLs, predicted coding and splice consequences, and open-field locomotor phenotypes from knockouts of orthologous mouse genes. Our results highlighted established neurobehavioral candidates spanning locomotor and reward-related biology (*Negr1* and *Lmo4*), serotonergic signaling (*Htr1b*), and GABAergic signaling (*Gabrg2*), as well as relatively novel genes including *Tcerg1l, Pcdh10,* and *Tmem132e*. Our results illustrate how Genomic SEM can integrate genetically correlated yet heterogeneous behavioral measures and enable discovery of loci underlying their shared genetic architecture, highlighting its utility for complex-trait studies in rodents.

## 1 Introduction

Substance use disorders (SUDs) are common, heritable psychiatric disorders that impose a substantial health and socioeconomic burden worldwide (Gerring et al., 2024). Human genetic studies have shown that genetic liability is shared both across SUDs and with broader externalizing traits related to behavioral disinhibition and self-regulation (Karlsson Linnér et al., 2021; Poore et al., 2026). Locomotor response to a novel environment has been studied in rodents for nearly a century and is an addiction-relevant behavioral phenotype (Wise and Bozarth, 1987). Early work in rats by Hall characterized individual differences in ambulatory activity and emotional behavior and demonstrated inherited variation through selective breeding (Hall, 1936, 1938). Later studies showed that rats with higher locomotor responses to novelty were more likely to acquire psychostimulant self-administration (Piazza et al., 1989). Selective breeding for high and low exploratory locomotion has further linked this trait to broader, heritable temperamental dimensions associated with sensation seeking, externalizing and internalizing behavior (Stead et al., 2006; Flagel et al., 2014; Zhou et al., 2019; Chitre et al., 2023), while recent work in outbred Sprague–Dawley rats has continued to examine novelty-induced locomotion among genetically influenced behavioral traits relevant to addiction (Chitre et al., 2023; Chitre et al., 2026).

Genetic studies in rats and mice have established that exploratory locomotor behavior is heritable and genetically complex, however, the specific genes that influence this trait remain largely unknown. A previous genome-wide association study (GWAS) that included a subset (*N* = 1,246) of the current sample identified quantitative trait loci (QTLs) for open-field behaviors including locomotor activity (Gunturkun et al., 2022). Studies in Sprague–Dawley rats have also examined the genetic basis of exploratory locomotion and related addiction-relevant behaviors in both selectively bred and commercially available outbred populations (Chitre et al., 2023; Chitre et al., 2026), while mouse genetic studies have mapped open-field, locomotor, and related behavioral traits in multiple genetically diverse populations, including Diversity Outbred, BXD recombinant inbred, advanced-intercross, and commercially outbred mice (Logan et al., 2013; Parker et al., 2016; Nicod et al., 2016; Delprato et al., 2017; Gonzales et al., 2018; Zou et al., 2022). Heterogeneous stock (HS) rats are particularly well suited for genetic mapping of complex traits because their genetic diversity and accumulated recombination enable association signals to be mapped at relatively high resolution (Solberg Woods and Palmer, 2019). Larger sample sizes substantially increase QTL discovery in HS rats, underscoring the importance of large samples for mapping complex traits (Chitre et al., 2023).

In the present study, we sought to characterize the genetic basis of locomotor activity using data from 9,341 male and female HS rats collected across eight studies that were part of a multisite collaborative project. These studies examined addiction-relevant behavioral phenotypes that included locomotor activity. Because the studies differed in various ways, we first estimated SNP heritability and genetic correlations among the locomotor measures. We then compared three complementary approaches to genome-wide locus discovery: 1) pooled individual-level analysis (MEGA), 2) fixed-effect meta-analysis (METAL), and 3) Genomic SEM, which models genetic variation shared across the locomotor measures (Grotzinger et al., 2019).

This study represents the first application of Genomic SEM in rodents. We also evaluated cross-study SNP-effect patterns and integrated functional genomic and comparative phenotype information to prioritize candidate genes. Together, these analyses provide by far the most powerful genetic characterization of locomotor activity in any rodent population.

## 2 Methods

### 2.1 Subjects and cohorts

Locomotor response to a novel environment was measured in 9,341 male and female HS rats from three HS colonies: NMcwi:HS (RRID:RGD_2314009), NMcwiWfsm:HS (RRID:RGD_13673907), and McwiWfsmAap:HS (HS West; RRID:RGD_155269102), across eight cohorts. These comprised the Dietz and Meyer laboratories at the University at Buffalo (UB-1 and UB-2, respectively), the Jhou and Kalivas laboratories at the Medical University of South Carolina (MUSC-1 and MUSC-2, respectively), the Mitchell laboratory at Oregon Health & Science University (OHSU), the Chen laboratory at the University of Tennessee Health Science Center (UTHSC), the Flagel and Robinson laboratories at the University of Michigan (UMich), and the de Guglielmo laboratory at the University of California San Diego (UCSD). Cohort sample sizes and demographic, breeding, environmental, and procedural characteristics are provided in Supplementary Table 1.

All animal procedures were approved by the relevant Institutional Animal Care and Use Committee.

### 2.2 Locomotor phenotyping

Locomotor activity was quantified as total distance traveled during the initial cohort-specific analysis window. Six cohorts used the first 15 min, whereas UB-2 and MUSC-1 used the first 10 and 20 min, respectively. Locomotor activity was recorded using beam-break or photobeam systems in UB-1, OHSU, MUSC-2, MUSC-1, and UB-2 and by video tracking in UTHSC, UMich, and UCSD. Arena dimensions, tracking systems, testing phase, lighting conditions, and prior behavioral or drug exposure varied among cohorts and are detailed in Supplementary Table 1.

### 2.3 Phenotype processing

Within each cohort, locomotor activity was quantile normalized. Candidate covariates included sex, coat color, phenotyping batch, age, and experimental box. The effects of covariates explaining more than 2% of phenotypic variance were regressed out. The resulting residualized phenotypes were used for SNP-heritability estimation, genetic-correlation analysis, and cohort-specific GWAS.

For MEGA, the residualized locomotor phenotypes from the cohorts included in the primary association analyses were concatenated and quantile normalized again to generate the pooled locomotor phenotype used for association analysis.

### 2.4 DNA sequencing, genotype processing, and variant quality control

Following behavioral testing, spleen tissue was collected for DNA extraction. Low-coverage whole-genome sequencing or genotyping-by-sequencing was performed at the University of Chicago until 2016 and at the University of California San Diego thereafter, as described previously (Chen et al., 2025). Variants were represented on the GRCr8 (rn8) rat reference assembly (Li et al., 2024).

Before variant-level quality control, the genotype dataset contained 7,659,686 biallelic SNPs on autosomes. Variants were excluded for genotype missingness ≥10%, minor-allele frequency ≤0.005, or Hardy–Weinberg equilibrium *P* < 1 × 10^-10^ based on the full sample size. The final genotype set contained 5,523,299 biallelic SNPs.

### 2.5 Genetic relatedness, SNP heritability, and genetic correlations

A genomic relatedness matrix (GRM) was constructed from the filtered genotype data. SNP heritability (*h*^2^) was estimated separately for locomotor activity in each of the eight cohorts using univariate restricted maximum likelihood (REML) in GCTA (Yang et al., 2011). Pairwise genetic correlations *r*_G_ among the cohort-specific locomotor phenotypes were estimated using bivariate GCTA REML.

### 2.6 Genetic-correlation downsampling

To evaluate the variability of genetic-correlation estimates at sample sizes comparable to the smaller primary cohorts, we used UB-1, the largest cohort (*N* = 2,473). In each of 100 repetitions, UB-1 animals were randomly divided into two nonoverlapping subsets of 618 rats (which is similar to the sample size for the OHSU and MUSC-1 cohorts), and *r*_G_ was estimated between the subsets using bivariate GCTA REML.

### 2.7 Genomic SEM covariance estimation and factor-model selection

Genomic SEM was used to model the genetic component shared across the cohort-specific locomotor measures and to perform multivariate association analysis (Grotzinger et al., 2019). The genetic covariance matrix (*S*) and its sampling covariance matrix (*V*) were obtained from individual-level data using a joint covariance-component REML workflow implemented with MPH (Jiang, 2024), rather than the multivariate linkage disequilibrium score regression (LDSC) workflow commonly used to generate Genomic SEM inputs in human genetic studies. Because LDSC-based approximations were developed primarily for human datasets with limited relatedness and rapidly decaying linkage disequilibrium (LD), we used MPH-based REML to accommodate the relatedness and LD characteristics of HS rats (Jiang, 2024).

Covariance matrices were initially constructed using all eight cohorts. The eight-cohort *S* matrix required substantial smoothing, and model fitting was therefore repeated after excluding the UCSD cohort, which had the lowest *h*^2^ estimate. The resulting seven-cohort *S* matrix did not require smoothing, and the seven-cohort covariance matrices were used for subsequent Genomic SEM analyses.

Alternative factor models were fitted using diagonally weighted least squares (DWLS). The model-comparison set comprised a common factor model; common factor models including a residual covariance between UTHSC and UMich, between UB-2 and MUSC-1, or both; and a two-factor model separating the five 15-min cohorts from the two non-15-min cohorts. UTHSC and UMich both used video tracking, motivating the first residual covariance. UB-2 and MUSC-1 were the two cohorts with non-15-min analysis windows and were also the two cohorts with prior drug exposure, motivating the second residual covariance and the two-factor specification.

Model fit was summarized using χ^2^, Akaike information criterion (AIC), comparative fit index (CFI), and standardized root mean square residual (SRMR). Relative model fit and theoretical interpretability were considered when selecting the model for multivariate association analysis (Supplementary Table 4; Supplementary Figure 3).

### 2.8 Genome-wide association analyses

To allow direct comparison across approaches, the same seven cohorts were used for the primary MEGA, METAL, and Genomic SEM analyses; thus the UCSD cohort was not included in the MEGA, METAL or Genomic SEM analyses.

#### 2.8.1 Cohort-specific GWAS and MEGA

Cohort-specific GWAS were performed using the mixed-linear-model association with leave-one-chromosome-out approach implemented in GCTA (MLMA-LOCO; Listgarten et al., 2012; Cheng et al., 2013). Summary statistics from the seven primary cohort GWAS were used as inputs for METAL and Genomic SEM. Individual-cohort GWAS were also evaluated separately for secondary locus discovery.

For MEGA, individual-level genotype data from the same seven cohorts (*N* = 8,595) were analyzed jointly using MLMA-LOCO and the pooled locomotor phenotype described above.

To identify conditionally independent association signals in MEGA, conditional analyses were restricted to variants within ±10 Mb of each initial QTL; when multiple initial QTLs occurred on the same chromosome, the union of these regions was analyzed. The strongest lead SNP was included as a covariate and GCTA MLMA was rerun iteratively within this regional candidate set, adding additional genome-wide-significant lead SNPs as covariates until no further significant signals remained. This conditional analysis was specific to MEGA.

#### 2.8.2 METAL

Summary statistics from the seven primary cohort GWAS were combined using fixed-effect inverse-variance meta-analysis in METAL (Willer et al., 2010). SNP-effect estimates and standard errors were aligned to a common effect allele before meta-analysis. No additional genomic-control correction was applied in METAL.

#### 2.8.3 Genomic SEM multivariate GWAS

The cohort-specific GWAS summary statistics, together with the MPH-derived *S* and *V* matrices, were used for the Genomic SEM multivariate GWAS. The selected factor model was then used to estimate genome-wide SNP effects on the common genetic factor, with the expected pattern of SNP effects across cohorts defined by the factor loadings (Grotzinger et al., 2019).

### 2.9 Genome-wide significance and QTL definition

Genome-wide discovery used simulation-based empirical 5% and 10% thresholds of−log_10_(*P*) = 5.449 and 5.171, respectively. Thresholds were estimated from 1,000 simulated standard-normal null phenotypes by retaining the genome-wide maximum −log_10_(*P*) from each replicate and taking the corresponding empirical percentiles. The same thresholds were applied to MEGA, METAL, Genomic SEM, and individual-cohort GWAS. The primary analyses were restricted to the autosomes because an X-chromosome-specific MPH–Genomic SEM workflow, including appropriate sex and dosage-compensation handling, was not implemented and validated in this study.

Method-specific QTL intervals were defined using variants in LD (*r*^2^ ≥ 0.8) with the corresponding method-specific lead SNP, adapting the LD-based QTL framework used previously in HS rats (Chitre et al., 2020). Signals identified by different methods were assigned to the same QTL when their method-specific lead SNPs were in LD (*r*^2^ ≥ 0.6); physical overlap between QTL intervals alone was not used to group signals. QTLs supported by MEGA, METAL, and/or Genomic SEM comprised the primary discovery set, whereas loci detected only in individual-cohort GWAS were reported separately as secondary findings.

### 2.10 Cross-cohort SNP-effect diagnostics

Cross-cohort SNP-effect patterns were evaluated at the exact METAL and Genomic SEM lead SNPs. For METAL, Cochran’s *Q* tested whether cohort-specific SNP-effect estimates varied more than expected from sampling error under the fixed-effect meta-analysis, and *I*^2^ summarized the proportion of observed variability attributable to between-cohort heterogeneity. For Genomic SEM, *Q*_SNP_ tested whether the pattern of cohort-specific SNP effects departed from that expected under the selected factor model (Grotzinger et al., 2019). Cochran’s *Q* and *Q*_SNP_ therefore test different null hypotheses.

Bonferroni and Benjamini–Hochberg false-discovery-rate (FDR) corrections were applied separately to Cochran’s *Q* heterogeneity *P* values across the METAL lead SNPs and to *Q*_SNP_ *P* values across the Genomic SEM lead SNPs.

### 2.11 Functional annotation

#### 2.11.1 Brain molecular QTL and predicted variant annotation

Genes within finalized QTL intervals were annotated using brain molecular QTLs (xQTLs) and predicted coding or splice consequences. Brain xQTL data were obtained from RatGTEx (Munro et al., 2022; Munro et al., 2024) and included gene expression, splicing, RNA stability, isoform ratio, alternative-transcription-start-site, and alternative-polyadenylation QTLs from brain-hemisphere and brain-region datasets. Molecular-QTL associations were retained if nominal *P* < 1 × 10^-4^ and the molecular-QTL variant was in LD with the relevant method-specific GWAS lead SNP (*r*^2^ > 0.6).

Predicted coding and splice consequences were annotated using SnpEff (Cingolani et al., 2012) and classified by predicted impact. LD between annotated variants and the relevant method-specific lead SNP was used for biological interpretation.

#### 2.11.2 Mouse ortholog and IMPC phenotype annotation

Genes within finalized QTL intervals were mapped to mouse orthologs using Ensembl Compara release 116 (Herrero et al., 2016). Mouse orthologs were subsequently queried against International Mouse Phenotyping Consortium (IMPC) Data Release 24 (Wilson et al., 2026). Open Field findings were classified as direct locomotor, tested-null, or complex; other motor or coordination, body or mechanical, sensory, and viability or developmental phenotypes were retained as additional functional context. Detailed orthology and phenotype-classification criteria are provided in Supplementary Table 7.

#### 2.11.3 Human genetic context

For additional human genetic context, genes within finalized QTL intervals were also cross-referenced with two recent large-scale studies of personality (Schwaba et al., 2026) and psychiatric disorders (Grotzinger et al., 2026); overlapping evidence is reported in Supplementary Table 7.

## 3 Results

### 3.1 Locomotor activity is heritable across cohorts and shows positive but variable genetic correlations

SNP heritability was estimated separately for locomotor activity in each of the eight cohorts.

Estimates were significantly greater than zero in all cohorts, ranging from 0.168 ± 0.058 in UCSD to 0.397 ± 0.068 in OHSU (Supplementary Table 2; Supplementary Figure 1). Across all cohort pairs, genetic-correlation point estimates were positive but varied substantially in magnitude and precision, ranging from *r*_G_ = 0.062 ± 0.139 for UTHSC–UB-2 to *r*_G_ = 0.795 ± 0.180 for UTHSC–UCSD (Supplementary Table 3; Supplementary Figure 1).

It was not clear if these levels of genetic correlation indicated true differences between cohorts, or reflected expected levels of genetic correlations given the modest sample sizes used (e.g. *N =* 538–673; Supplementary Table 1A). To address this question, we performed repeated downsampling of UB-1, the largest cohort (*N* = 2,473). In each of 100 repetitions, UB-1 animals were divided into two non-overlapping random subsets of 618 rats (25% of 2,473) and *r*_G_ was estimated for the same locomotor phenotype between the two subsets. The mean *r*_G_ was 0.93 and the median was 1.00, with estimates ranging from 0.51 to 1.00 (Supplementary Figure 2). Thus, when data were drawn from a single cohort (UB-1), observed genetic correlations were higher than between-cohort genetic correlations, suggesting that there were true cohort-specific differences.

### 3.2 A common factor model with two residual covariances was selected for Genomic SEM

We next used Genomic SEM to model the shared genetic component across the cohort-specific locomotor measures and select a factor model for multivariate GWAS. With all eight cohorts included, the estimated genetic covariance matrix *S* was non-positive definite and required substantial smoothing, with a maximum absolute change in an *S*-matrix element of 0.058, exceeding the Genomic SEM warning threshold of 0.025. Given the substantial smoothing required for the eight-cohort matrix, we repeated model fitting after excluding UCSD, which had the lowest SNP-heritability estimate (*h*^2^ = 0.168 ± 0.058) among the eight cohorts.

The resulting seven-cohort *S* matrix did not require smoothing, and subsequent Genomic SEM analyses used these seven cohorts only.

We first evaluated a common factor model in which all seven cohorts loaded on a single common genetic factor. Alternative specifications were motivated by biologically and experimentally interpretable features of the contributing protocols. UTHSC and UMich both used video tracking, motivating a residual covariance between these measures. UB-2 and MUSC-1 were the two cohorts with non-15-min analysis windows and were also the two cohorts with prior drug exposure, motivating a second residual covariance. We additionally evaluated a two-factor model that separated the 15-min and non-15-min cohorts (Supplementary Table 4; Supplementary Figure 3).

Among the candidate models that yielded proper solutions, the common factor model with residual covariances between UTHSC and UMich and between UB-2 and MUSC-1 provided the best fit (χ^2^ = 23.56, df = 12, *P* = 0.023, AIC = 55.56, CFI = 0.945, SRMR = 0.113) while maintaining a theoretically interpretable one-factor structure. We therefore selected this specification for the Genomic SEM multivariate GWAS. All seven cohort measures loaded positively on the common factor, with standardized loadings ranging from 0.29 to 0.92; UTHSC showed the lowest loading (0.29; Figure 1).

**Figure 1.**
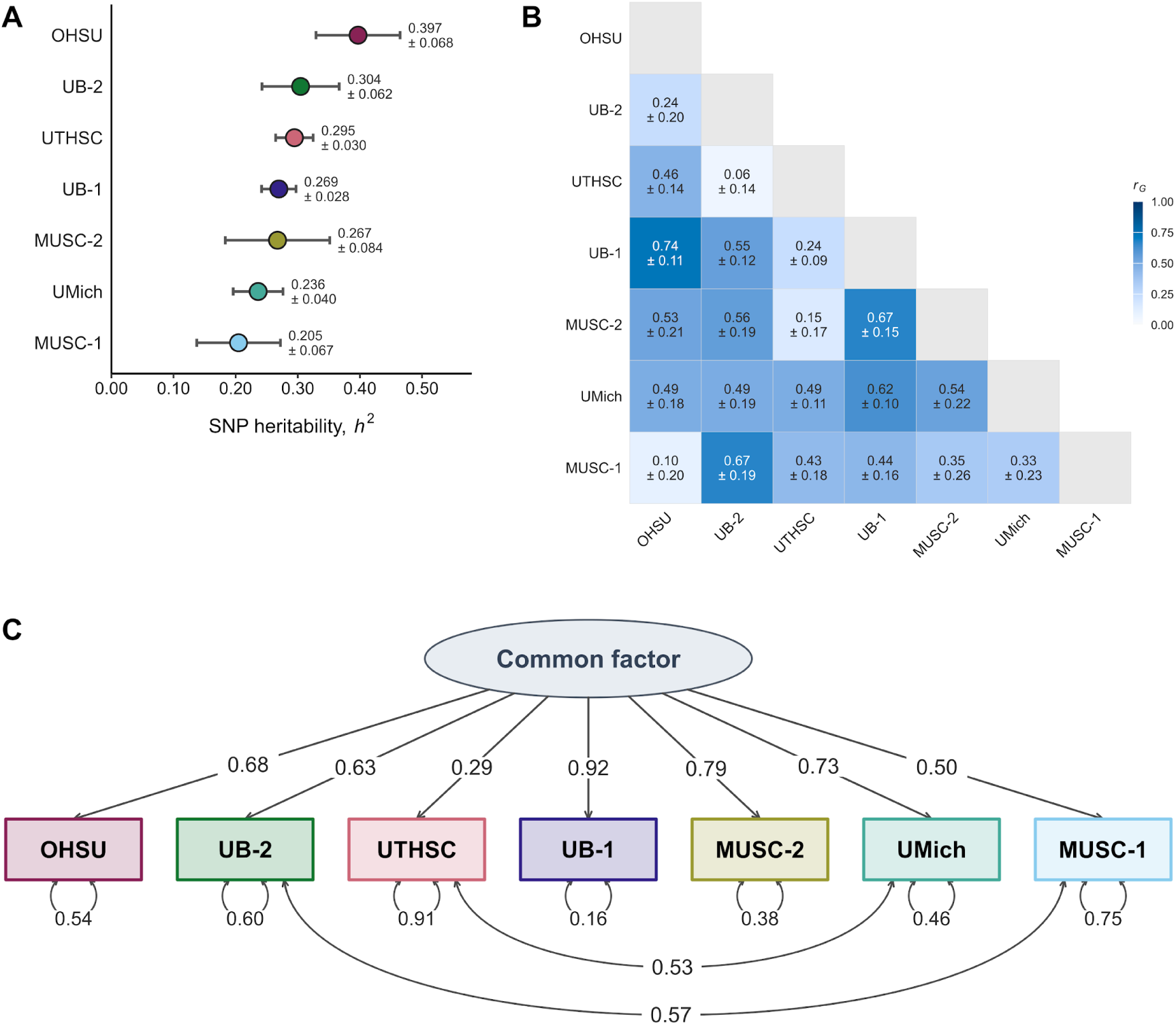
Locomotor activity is heritable across cohorts and shows positive but variable genetic correlations. **(A)** SNP-heritability (*h*^2^) point estimates across the seven primary cohorts ranged from 0.205 to 0.397. Cohort-specific estimates were obtained using univariate GCTA restricted maximum likelihood (REML); points show *h*^2^ estimates, horizontal bars indicate standard errors, and numerical values are estimate ± SE. **(B)** Pairwise genetic correlations (*r*_G_) were positive, with point estimates ranging from 0.06 to 0.74. Correlations were estimated using bivariate GCTA REML; cells show *r*_G_ ± SE, with color intensity indicating the magnitude of the genetic correlation. **(C)** In the selected seven-trait Genomic SEM model, all cohorts loaded positively on a common genetic factor, with standardized loadings ranging from 0.29 to 0.92. The model additionally included residual genetic covariances between UTHSC and UMich and between UB-2 and MUSC-1. Values on single-headed paths are standardized factor loadings, values associated with self-loops are standardized residual variances, and values on curved double-headed paths are standardized residual covariances. Cohort ordering is consistent across panels.

### 3.3 QTL discovery across complementary analyses

We applied three discovery approaches (MEGA, METAL, and Genomic SEM) using the same seven cohorts. MEGA pooled quantile-normalized individual-level data across cohorts and estimated SNP effects on a pooled locomotor phenotype. METAL retained cohort-specific association estimates and combined them using fixed-effect inverse-variance meta-analysis.

Genomic SEM multivariate GWAS instead tested SNP associations with the common genetic factor defined by the selected factor model, as described above.

Genome-wide discovery used simulation-based empirical 5% and 10% thresholds of −log_10_ (*P*) = 5.449 and 5.171, respectively. Across autosomes, the three analyses identified 38 primary QTLs (Figure 2; Supplementary Table 5A). Eight QTLs were supported by MEGA, 13 by METAL, and 31 by Genomic SEM. Thirty-five QTLs reached the empirical 5% threshold and three reached the empirical 10% threshold. Eight additional loci detected only in individual-cohort analyses were retained as secondary findings (Supplementary Table 5B; Supplementary Figure 4).

**Figure 2.**
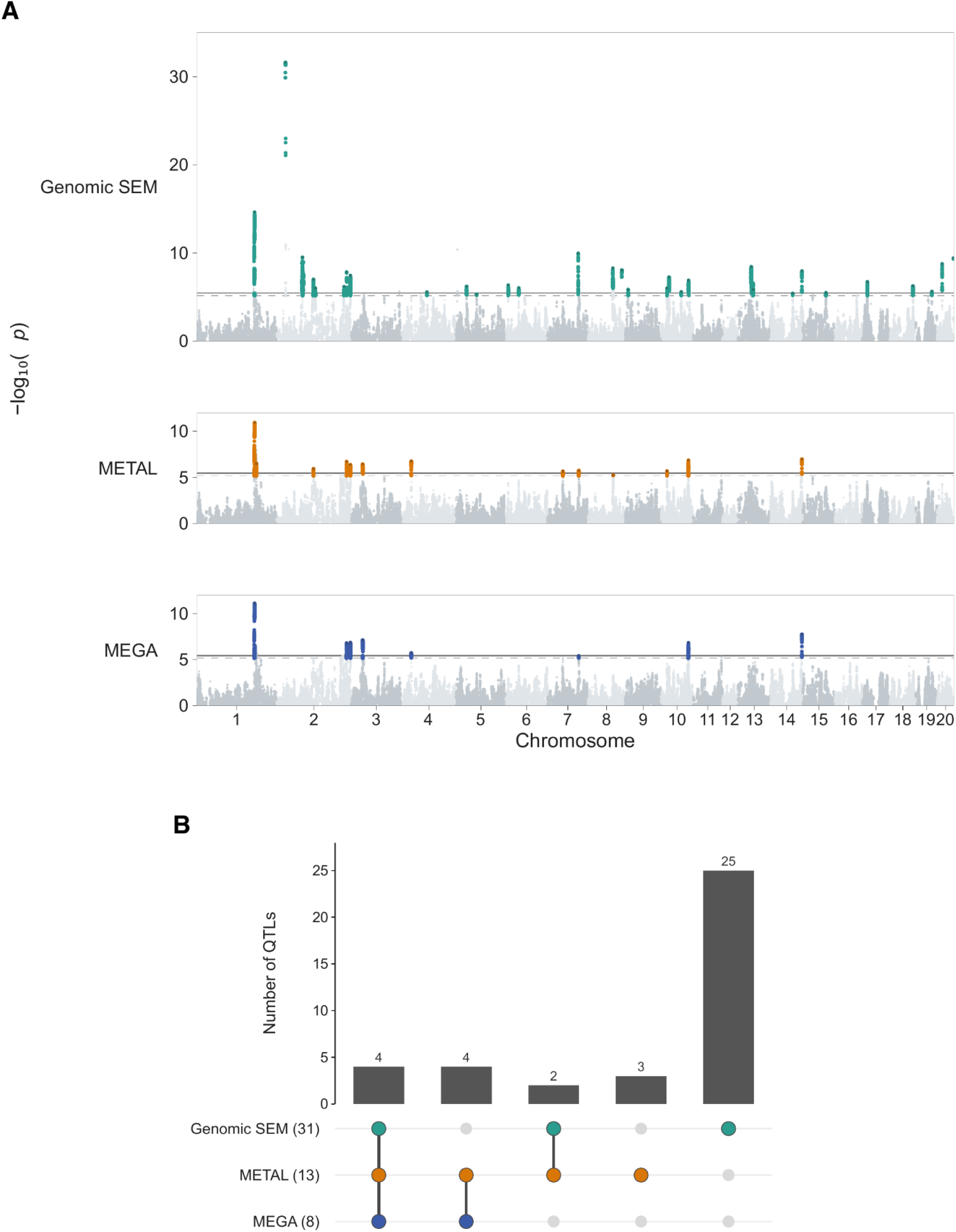
Complementary genome-wide analyses identify shared and method-specific locomotor-activity QTLs. **(A)** Stacked Manhattan plots of the Genomic SEM, METAL, and MEGA association analyses across autosomes. SNPs within method-specific QTL intervals meeting the 10% discovery threshold are colored according to discovery method; other SNPs are shown in alternating neutral chromosome tones. Solid and dashed horizontal lines indicate the simulation-based empirical 5% (−log_10_(*P*) = 5.449) and 10% (−log_10_(*P*) = 5.171) discovery thresholds, respectively. Method-specific y-axis ranges are used to preserve within-method resolution. **(B)** Set-overlap plot summarizing mutually exclusive support patterns among the 38 primary QTLs identified by MEGA, METAL, and Genomic SEM. QTL overlap information is provided in Supplementary Table 5A.

The three approaches showed both convergence and method-specific discovery (Figure 2). All eight MEGA-supported QTLs were also detected by METAL, which identified five additional QTLs. Genomic SEM identified 25 unique QTLs and six additional QTLs that were also detected by MEGA, METAL, or both (Figure 2). Signals identified by different methods were assigned to the same QTL based on LD between their method-specific lead SNPs (*r*^2^ ≥ 0.6).

### 3.4 Cross-cohort SNP-effect diagnostics

We next examined cross-cohort SNP-effect estimates at the exact lead SNPs from METAL and Genomic SEM.

#### 3.4.1 METAL heterogeneity

Cochran’s *Q* tests whether SNP-effect estimates vary across cohorts more than expected from sampling error. *I*^2^ indicates how much of the observed variation in SNP effects reflects differences between cohorts rather than sampling error. None of the 13 exact METAL lead SNPs showed nominally significant Cochran heterogeneity (*P*_het_ > 0.05; Supplementary Table 6A). Six loci had *I*^2^ = 0%, six had *I*^2^ between 25% and <50%, and chr3:59.4 Mb had the highest *I*^2^, at 51.7%, although it did not reach nominal Cochran heterogeneity significance (*P*_het_ = 0.053).

Figure 3 illustrates this range: chr4:33.6 Mb showed particularly coherent cohort effects (*Q* = 1.84, *P*_het_ = 0.934, *I*^2^ = 0%; Figure 3A), whereas chr3:59.4 Mb showed the greatest between-cohort variability among the METAL leads (Figure 3B).

**Figure 3.**
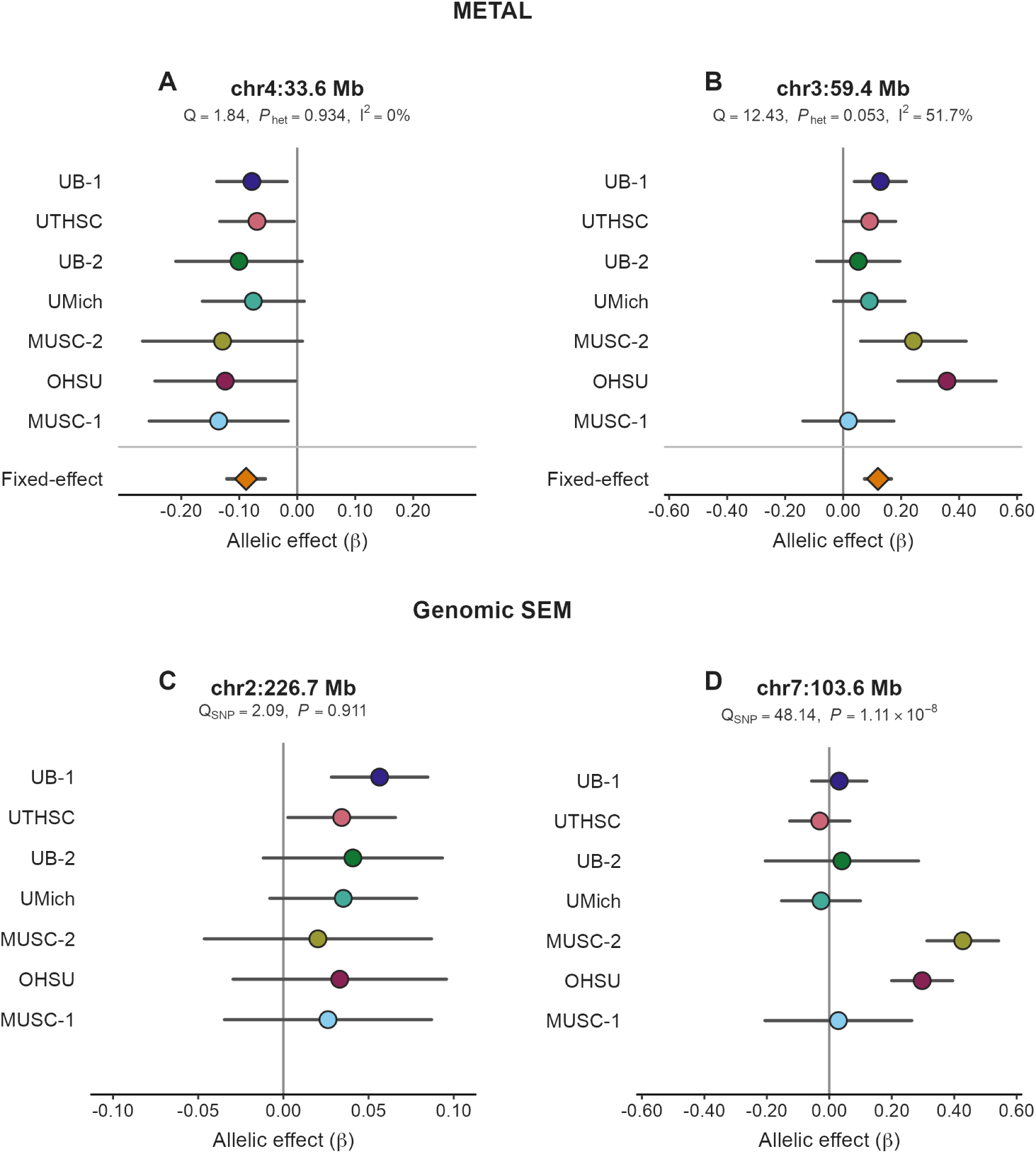
Representative cross-cohort SNP-effect structures. Cohort-specific allelic-effect estimates (*β*; circles) and 95% confidence intervals are shown for the seven primary cohorts, with effects oriented to the effect allele of each method-specific lead SNP. **(A)** METAL-supported chr4:33.6 Mb locus showing highly consistent cohort effects. **(B)** METAL-supported chr3:59.4 Mb locus showing greater cross-cohort variability. **(C)** Genomic SEM-supported chr2:226.7 Mb locus with cohort effects closely matching the pattern predicted by the selected factor model. **(D)** Genomic SEM-supported chr7:103.6 Mb locus showing marked departure from the factor-implied pattern. For METAL panels, orange diamonds and horizontal lines show the fixed-effect summary estimate and its 95% confidence interval; Cochran’s *Q*, heterogeneity *P* value (*P*_het_), and *I*^2^ summarize between-cohort heterogeneity. For Genomic SEM panels, *Q*_SNP_ tests whether the cohort-specific SNP-effect pattern departs from that predicted by the selected factor model; no pooled summary estimate is shown. Vertical gray lines mark *β* = 0.

#### 3.4.2 Genomic SEM *Q*_SNP_

Genomic SEM *Q*_SNP_ tests whether the pattern of SNP effects across the seven cohorts departs from that expected under the selected factor model. By contrast, Cochran’s *Q* in the METAL fixed-effect meta-analysis tests whether cohort-specific SNP effects vary more than expected around a common effect. Among the 31 exact Genomic SEM lead SNPs, 11 showed nominal *Q*_SNP_ significance, six remained significant after Benjamini-Hochberg FDR correction, and five remained significant after Bonferroni correction (Supplementary Table 6B). Therefore, 20 of the 31 lead SNPs were not significant for the *Q*_SNP_ test.

The chr2:226.7 Mb locus provides a representative low-*Q*_SNP_ example (*Q*_SNP_ = 2.09, *P* = 0.911; Figure 3C). In contrast, chr7:103.6 Mb showed strong evidence of additional SNP-specific cohort-effect structure (*Q*_SNP_ = 48.14, *P* = 1.11 × 10^−8^), with substantially larger positive effects in MUSC-2 and OHSU than in the other 5 cohorts (Figure 3D). A SNP may be associated with the common genetic factor while also showing significant *Q*_SNP_, indicating that its cohort-specific effect pattern is not fully captured by the selected factor model. Together, these examples illustrate the locus-specific nature of SNP-effect heterogeneity. Cohort-specific forest plots for all 13 METAL and 31 Genomic SEM lead SNPs are provided in Supplementary Figure 6.

### 3.5 Functional annotation of QTLs

To characterize the biological context of the 38 QTLs, we first examined the size and gene content of each QTL interval. Interval widths ranged from approximately 0.17 to 11.72 Mb, and the number of genes within QTL intervals ranged from 0 to 37. Of these 38 intervals, 24 contained multiple genes, 10 contained only a single gene, and 4 contained no genes. We then integrated three gene-level evidence layers: brain molecular QTLs spanning gene expression, isoform ratio, splicing, alternative transcription start-site usage, alternative polyadenylation usage, and RNA stability; predicted coding or splice consequences; and standardized IMPC phenotype data for mouse orthologs of rat genes within the QTL intervals. Detailed gene-level annotations are provided in Supplementary Table 7A, with a locus-level summary in Supplementary Table 7B.

Brain molecular QTL evidence was identified for at least one gene within 30 of the 38 QTLs.

Eighteen QTLs contained at least one gene with a predicted MODERATE coding or splice consequence, and one QTL additionally contained a predicted HIGH-impact stop-lost variant in the uncharacterized gene *LOC102546663*. Usable IMPC phenotype data were available for at least one gene in 28 QTL intervals. Twenty-five QTLs contained at least one gene tested in the IMPC Open Field assay, and nine unique genes across seven QTLs showed direct or complex Open Field locomotor phenotypes. IMPC Open Field results included both significant and tested-null outcomes, all of which are reported in Supplementary Table 7.

These complementary annotation layers were used to prioritize candidate genes and loci for biological interpretation, including *Negr1*, *Lmo4*, *Tcerg1l*, and *Camkmt*. Based on these evidence layers and their contrasting discovery patterns, we selected *Negr1*, *Tcerg1l*, and *Camkmt* as representative loci for regional visualization in Figure 4. Regional association plots for the remaining primary harmonized loci are provided in Supplementary Figure 5.

**Figure 4.**
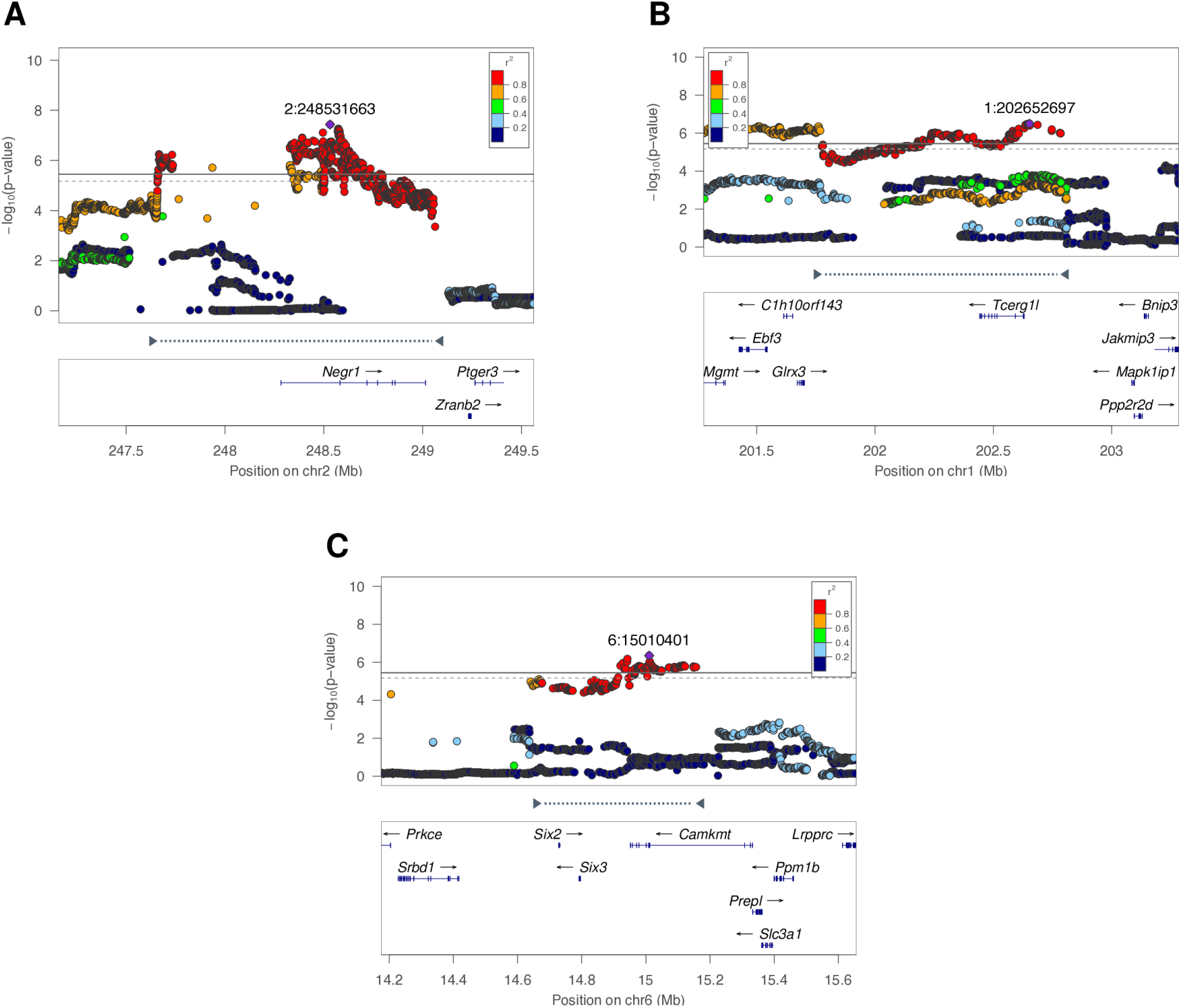
Regional association plots for representative locomotor-activity loci. Regional association plots illustrate complementary discovery patterns at three representative loci. **(A)** The chr2:248.5 Mb *Negr1* locus, supported by MEGA, METAL, and Genomic SEM and displayed using the Genomic SEM association results. **(B)** The chr1:202.7 Mb *Tcerg1l* locus, a METAL-specific discovery. **(C)** The chr6:15.0 Mb *Camkmt* locus, a Genomic SEM-specific discovery. Association significance is plotted as −log_10_(*P*) against rn8 genomic position. Points are colored according to linkage disequilibrium (*r*^2^) with the method-specific lead SNP in each panel; lead SNPs are labeled by chromosome:position. Solid and dashed horizontal lines indicate the simulation-based empirical 5% (−log_10_(*P*) = 5.449) and 10% (−log_10_(*P*) = 5.171) discovery thresholds, respectively. NCBI RefSeq gene models provide local genomic context, and the QTL interval is indicated by inward-facing interval markers connected by a dotted line.

## 4 Discussion

Locomotor response to a novel environment is a widely studied, addiction-relevant behavioral phenotype (e.g. Wise and Bozarth, 1987; Piazza et al., 1989). The present study leveraged a large HS-rat dataset assembled across eight studies within a multisite collaborative project examining addiction-relevant behavioral phenotypes. After filtering the available data, seven cohorts were retained for the primary association analyses (*N* = 8,595). This constitutes the largest rodent GWAS for any complex trait. HS rats are particularly useful for dissecting the genetic architecture of complex traits because their highly recombinant genomes support high-resolution mapping of genetic loci (Baud et al., 2013; Solberg Woods and Palmer, 2019). We first characterized the genetic contribution to locomotor activity within each cohort and estimated the genetic correlations among the cohort-specific measures. Locomotor activity was heritable across cohorts, and showed positive but variable genetic correlations, motivating complementary association analyses. MEGA, METAL, and Genomic SEM identified overlapping and distinct loci, while functional annotation highlighted established neurobehavioral candidates as well as potentially novel candidate genes and loci for locomotor activity.

### 4.1 Shared genetic architecture across heterogeneous cohorts

Locomotor activity was heritable across all eight cohorts, with SNP-heritability estimates ranging from 0.168 ± 0.058 to 0.397 ± 0.068 (Supplementary Table 2; Supplementary Figure 1). The cohorts differed in age, diet and housing, study site, locomotor-testing protocols, and prior behavioral or drug exposure (Supplementary Table 1). Pairwise genetic-correlation estimates were all positive but varied substantially in magnitude and precision, ranging from *r*_G_ = 0.062 ± 0.139 to *r*_G_ = 0.795 ± 0.180 (Supplementary Table 3; Supplementary Figure 1). The cohort measures therefore captured related aspects of locomotor activity that shared genetic influences to varying degrees. This pattern motivated the use of Genomic SEM to model the genetic component shared across related but nonidentical locomotor measures (Grotzinger et al., 2019).

The Genomic SEM model comparison provided a complementary view of this shared but nonuniform genetic architecture. Among the candidate models that yielded proper solutions, the common factor model with residual covariances between UTHSC and UMich and between UB-2 and MUSC-1 provided the most favorable relative fit while retaining a theoretically interpretable one-factor structure (Figure 1). We therefore used this model for the Genomic SEM multivariate GWAS.

UTHSC showed the lowest standardized loading on the common factor (0.29). UTHSC was distinctive in several respects, including its younger age distribution, video-based tracking, and the largest arena (100 × 100 × 50 cm). Because these characteristics varied together with other cohort differences, the lower loading may be due to one or more of these differences.

Several cohort characteristics motivated the alternative Genomic SEM models and may influence which aspects of locomotor behavior are captured. Video tracking records continuous spatial trajectories, whereas photobeam systems quantify movement from discrete beam interruptions. Session duration may also matter because locomotor activity changes due to within-session habituation (Leussis and Bolivar, 2006); 10-, 15-, and 20-min totals can therefore place different emphasis on the initial novelty response and later activity. Prior drug exposure and age also differed across cohorts. Because these characteristics varied together rather than being manipulated independently, they provide biological context for the model specifications rather than causal explanations for the residual covariances.

### 4.2 Complementary association analyses of locomotor activity

#### 4.2.1 Genomic SEM substantially expanded locus discovery

MEGA, METAL, and Genomic SEM address related but distinct questions about the genetic basis of locomotor activity. MEGA pools individual-level data across cohorts and estimates SNP effects on a pooled locomotor phenotype. METAL combines cohort-specific SNP-effect estimates using fixed-effect inverse-variance meta-analysis. Genomic SEM models the genetic covariance among the observed variables and tests SNP associations with the common genetic factor. Together, these analyses evaluate locus discovery under different assumptions about cross-cohort phenotype and effect structure.

Given the positive but variable genetic correlations among the cohort-specific locomotor measures, Genomic SEM provided a way to model their shared genetic component. Genomic SEM identified 31 QTLs compared with 13 for METAL and eight for MEGA, including 25 not detected by either METAL or MEGA.

Genomic SEM has been widely applied in human behavioral genetics to model shared genetic liability across correlated psychiatric and behavioral phenotypes (Grotzinger et al., 2019, 2026; Karlsson Linnér et al., 2021; Poore et al., 2026). In human studies, the covariance inputs required for Genomic SEM are commonly estimated using multivariate LDSC. LDSC-based approximations were developed primarily for human populations with limited relatedness and rapidly decaying LD. Many non-human experimental populations do not meet these assumptions, and both theoretical and empirical simulations have shown that LDSC should not be used in such settings. To address this limitation, we used a joint covariance-component REML workflow implemented with MPH to account for the relatedness and LD characteristics of HS rats (Jiang, 2024). Our study represents the first application of Genomic SEM in rodents.

The MPH-based covariance-estimation approach will allow application of Genomic SEM to other model-organism datasets with similar relatedness and LD patterns.

#### 4.2.2 Locus-specific cross-cohort SNP effects

When combining multiple datasets to perform a GWAS, some loci may have similar effects in each cohort whereas others may be cohort specific. We used cross-cohort effect diagnostics to evaluate whether SNP effects were consistent with the METAL fixed-effect model or the Genomic SEM factor model. In METAL, Cochran’s *Q* tests whether SNP-effect estimates vary across cohorts more than expected from sampling error. None of the 13 METAL lead SNPs showed nominally significant Cochran heterogeneity (Supplementary Table 6A). The chr4:33.6 Mb locus showed particularly coherent cohort effects (Figure 3A). In contrast, chr3:59.4 Mb showed the greatest between-cohort variability among the METAL leads (*I*^2^ = 51.7%), although Cochran’s *Q* did not reach nominal significance (*P*_het_ = 0.053; Figure 3B). In Genomic SEM, *Q*_SNP_ tests whether cohort-specific SNP effects depart from the pattern expected under the selected factor model. Most Genomic SEM lead SNPs showed no evidence of departure, whereas a subset showed additional cohort-specific effects (Supplementary Table 6B). The chr2:226.7 Mb locus closely followed the factor-implied pattern (Figure 3C). In contrast, chr7:103.6 Mb showed a strong departure, with substantially larger positive effects in MUSC-2 and OHSU than in most other cohorts (Figure 3D). Thus, association with the common genetic factor can coexist with locus-specific cohort effects; because cohort characteristics varied together, these analyses cannot determine which features contributed to the observed departures.

### 4.3 Biological interpretation of locomotor QTLs

Candidate genes were prioritized by integrating mapping resolution, cross-method support, brain molecular QTLs, predicted coding and splice consequences, IMPC phenotype data for mouse orthologs, and prior experimental literature. LD with the relevant method-specific lead SNP informed interpretation of regulatory and coding annotations; exact values and provenance are reported in Supplementary Table 7. To place the prioritized loci in a broader biological context, we considered, for each gene, evidence from mouse and rat studies of locomotor activity, psychostimulant response, reward-related behavior, and motor phenotypes, together with human genetic studies of related psychiatric and behavioral traits. Of particular relevance were two recent human genetic studies: Schwaba et al. (2026), which examined the genetic architecture of major personality traits, and Grotzinger et al. (2026), which mapped shared genetic liability across 14 psychiatric disorders. Across the rat QTL gene set, 12 genes were represented in the Schwaba et al. results and 25 showed overlap with the Grotzinger et al. results, with five represented in both studies; the strength and specificity of the human evidence varied across genes (Supplementary Table 7).

#### 4.3.1 Cross-method convergent loci

We first focused on loci supported by MEGA, METAL, and Genomic SEM. At the chr2:248.5 Mb QTL, *Negr1* is the only gene within the 1.41-Mb interval (Figures 2 and 4A). An RNA stability QTL for *Negr1* in brain hemisphere and a predicted MODERATE missense/splice-region variant were both in strong LD with the relevant lead SNP (*r*^2^ = 0.954 and 0.848, respectively).

Increasing or decreasing *Negr1* expression in rat hypothalamic regions has been shown to reduce locomotor activity and alter energy-balance phenotypes (Boender et al., 2014). A recent human GWAS identified a genome-wide-significant association for openness-to-experience at a locus containing *NEGR1* (Schwaba et al., 2026). This association parallels the broader relationship between exploratory locomotion and temperament in rats (Zhou et al., 2019; Chitre et al., 2023). Human GWAS of depression have also prioritized *NEGR1* through brain-expression and integrative gene-prioritization analyses, and recent functional work identified a depression-associated regulatory variant that alters *NEGR1* expression (Meng et al., 2024; Zhang et al., 2025). Together, these findings support *Negr1* as a strong putative candidate while indicating that both neural and metabolic pathways may contribute to variation in locomotor activity.

A second convergent chr2 QTL contained only one gene: *Lmo4*. Brain molecular QTLs for *Lmo4* spanned gene expression, RNA stability, and isoform ratio across brain hemisphere and several regions, including nucleus accumbens; an RNA stability QTL in brain hemisphere was in high LD with the Genomic SEM lead SNP (*r*^2^ = 0.807). Earlier work in *Drosophila* showed that reduced function of the LMO-family gene *Beadex* (*dLmo*) increased acute cocaine sensitivity and disrupted circadian locomotor rhythms (Tsai et al., 2004). Reduced *Lmo4* expression in mice increases sensitivity to the locomotor-stimulant effects of repeated cocaine, and targeted downregulation in nucleus accumbens reproduces this phenotype (Lasek et al., 2010).

Subsequent work identified an LMO4-regulated pathway that influences cocaine sensitization and reward (Lasek et al., 2011). Together, these findings connect the locus to brain regulation and psychostimulant-related locomotor and reward phenotypes.

#### 4.3.2 METAL-specific loci

Among the three METAL-specific QTLs, we highlight a potentially novel locus at chr1:202.7 Mb. The 1.01-Mb interval contains only one gene: *Tcerg1l* (Figures 2 and 4B). A nucleus-accumbens *Tcerg1l* expression QTL (eQTL) is in strong LD with this SNP (*r*^2^ = 0.839). Prior literature provides little direct evidence linking *Tcerg1l* to locomotor or closely related neurobehavioral phenotypes, suggesting that this may represent a novel finding from this study.

#### 4.3.3 Genomic SEM-specific loci

Among the 25 Genomic SEM-specific loci, we investigated whether these additional loci were supported by functional and neurobehavioral evidence. At chr6:15.0 Mb, the 0.48-Mb interval contains *Camkmt*, *Six2*, and *Six3* (Figures 2 and 4C). *Camkmt* is the leading candidate because a nucleus-accumbens splicing QTL is in near-complete LD with the lead SNP (*r*^2^ = 0.9995). Experimental disruption of *Camkmt* in mice causes abnormalities in muscle strength, motor learning, coordination, somatosensory development, and brain function (Haziza et al., 2015). Human GWAS have also implicated the *CAMKMT*-containing 2p21 region in anxiety-related phenotypes (Otowa et al., 2016; Hettema et al., 2020). Together, these findings prioritize *Camkmt* while *Six2* and *Six3* remain alternative interval candidates.

At chr2:226.7 Mb, a nucleus accumbens *Cisd2* splicing QTL and *Ube2d3* eQTL were in high LD with the lead SNP (*r*^2^ = 0.846 and 0.991, respectively). IMPC data showed reduced whole-arena locomotor activity after *Ube2d3* disruption. For *Cisd2*, whole-arena average speed was significantly reduced in both sexes, with a larger estimated effect in males. A separate *Cisd2* mutant mouse study likewise reported reduced open-field activity, particularly in males (Gaines et al., 2021). Together, these findings nominate both *Cisd2* and *Ube2d3* as plausible candidate genes at this locus.

At the chr8 locus, *Htr1b*, which encodes the serotonin 1B receptor (5-HT1B), lies within a two-gene interval and has a prelimbic cortex eQTL in strong LD with the lead SNP (*r*^2^ = 0.943). Mice lacking the serotonin 1B receptor show enhanced locomotor responses to cocaine, providing direct experimental support for a role in psychostimulant-related behavior (Rocha et al., 1998). A subsequent study replicated this effect but showed that it was modified by environmental context (Crabbe et al., 1999). The original study also found greater motivation to self-administer cocaine in the knockout mice (Rocha et al., 1998). In rats, 5-HT1B agonism altered locomotor activity in an age-dependent manner without psychostimulant challenge (McDougall et al., 2022), while *Htr1b* overexpression in the nucleus accumbens shell enhanced cocaine-induced locomotion without altering saline locomotion (Neumaier et al., 2002). In humans, *HTR1B* was significant in a genome-wide gene-based analysis of a broad externalizing factor (Poore et al., 2026). Together, these findings support *Htr1b* as a biologically plausible candidate at this locus and connect the rat locomotor association with serotonergic regulation of locomotor activity and broader externalizing liability.

At the four-gene chr20:54.7 Mb locus, *Ascc3* has a prelimbic cortex eQTL in strong LD with the lead SNP (*r*^2^ = 0.976), while *Cd99* and *Sim1* remain alternative interval candidates based on an alternative-polyadenylation QTL in brain hemisphere and a missense variant, respectively (Supplementary Table 7). Neuronal *Ascc3* deletion increased spontaneous activity and rearing in mice (Meziane et al., 2022). Human GWAS also identified intragenic *ASCC3* associations with neuroticism (Schwaba et al., 2026) and internalizing liability (Grotzinger et al., 2026).

Several other loci that were identified by METAL and Genomic SEM were also of interest. A QTL at chr2:130.3 Mb contained only a single gene: *Pcdh10*, which had an eQTL in brain hemisphere in near-complete LD with the Genomic SEM lead SNP (*r*^2^ = 0.998). The single-gene chr10:67.7 Mb QTL similarly prioritized *Tmem132e* through an RNA stability QTL in basolateral amygdala (*r*^2^ = 0.933) and a missense variant (*r*^2^ = 0.797) linked to the Genomic SEM lead SNP. At chr10:26.7 Mb, multiple brain molecular QTLs supported *Gabrg2*, including an RNA stability QTL in nucleus accumbens in near-complete LD with the Genomic SEM lead SNP (*r*^2^ = 0.9998). At chr3:59.4 Mb, an RNA stability QTL for *Galnt13* in nucleus accumbens was in near-complete LD with the METAL lead SNP (*r*^2^ = 0.998), whereas IMPC Open Field phenotypes supported *Kcnj3*. Prior experimental studies provide broader neurobehavioral and functional evidence for several of these genes: *Pcdh10*, *Gabrg2*, *Galnt13*, and *Kcnj3* (encoding GIRK1) (Hoshina et al., 2022; Zhang et al., 2019; Xu et al., 2016; Mett et al., 2021). The loci highlighted above are selected examples rather than an exhaustive account of the biological evidence. Supplementary Table 7 provides systematic gene-level annotation across all 38 QTLs and identifies additional candidates for future mechanistic studies.

### 4.4 Strengths and limitations

The combination of a large HS-rat dataset with complementary association analyses across heterogeneous cohorts was a strength of this study but also introduced several limitations.

Cohorts differed in age, apparatus, acquisition method, testing duration, and prior experimental history. Because these cohort characteristics varied together rather than being manipulated independently, differences in genetic correlation, factor loading, SNP effects, or *Q*_SNP_ cannot be attributed to any single protocol feature. As all three primary analyses used the same cohorts, cross-method agreement represents analytical convergence rather than independent replication. The cross-method comparison was restricted to autosomes because an X-chromosome-specific MPH–Genomic SEM workflow was not implemented and validated in this study. Although both male and female rats were included, sex-specific genetic effects were not evaluated. Functional annotation provided complementary support for candidate-gene prioritization but does not establish causal genes or variants. Mechanistic studies, including evaluation of additional regulatory and structural variation, will be needed to determine which genes and variants underlie individual QTLs. Overall, the scale and heterogeneity of this HS-rat dataset revealed shared genetic influences on locomotor activity alongside locus-specific variation in SNP effects. These findings highlight the value of modeling shared genetic variation while separately evaluating cross-cohort SNP-effect differences in heterogeneous behavioral datasets.

## 5 Conclusions

This study represents the largest rodent GWAS for any complex trait, with 8,595 HS rats included in the primary association analyses. Across cohorts, locomotor activity was heritable and showed shared but nonidentical genetic influences. Complementary association analyses identified 38 QTLs; this high discovery yield despite cohort heterogeneity was greatly facilitated by our application of Genomic SEM, using genetic covariance and sampling covariance matrices estimated with MPH rather than LDSC. Genomic SEM identified 31 QTLs, including 25 not detected by MEGA or METAL. Functional annotation connected these loci to established locomotor, reward- and psychostimulant-related, serotonergic, and other neurobehavioral biology while also nominating potentially novel candidate genes. Together, these findings provide a genetic framework for locomotor activity and demonstrate the value of modeling shared genetic variation across heterogeneous behavioral measures in model organisms.

## Data Availability

**Genotype data:** Palmer, Abraham A. (2026). Heterogeneous Stock (HS) Rat Genotypes, Version 8. In Genotype Data from: NIDA Center for GWAS in Outbred Rats. UC San Diego Library Digital Collections. https://doi.org/10.6075/J0SN09ZD

**Sequence Read Archive (SRA) reads:** BioProject accession number PRJNA487943 in the NCBI BioProject database: https://www.ncbi.nlm.nih.gov/bioproject/PRJNA487943

**Phenotype data** will be released to UC San Diego Library Digital Collections at the time of publication.

## Code Availability

Code implementing the MPH covariance-estimation and Genomic SEM workflows is available at https://github.com/ApurvaChitre/mph-pipeline and https://github.com/ApurvaChitre/gsem-pipeline.

## Acknowledgements

We thank Wenyan Han, Olivia Harrison, Gwen Johnson, Caroline Jones, Shuangying Leng, and Jun Huang of the University of Tennessee Health Science Center, and Jordan Bromley and Katie Garland of the Translational Neuroeconomics Lab at OHSU, for assistance with data collection.

Funding was provided by the following National Institutes of Health grants: P50DA037844, U01DA045300, U01DA046077, U01DA044468, P30DA060810, and R01AA030048.

