## Supplementary Figures for "Genomic SEM reveals shared genetic architecture and expands locus discovery for locomotor activity in heterogeneous stock rats"

### Supplementary Information

**Contents**

Supplementary Figure 1. SNP heritability and pairwise genetic correlations across all eight cohorts. 2

Supplementary Figure 2. Repeated within-cohort genetic-correlation downsampling analysis in UB-1. 3

Supplementary Figure 3. Alternative seven-trait Genomic SEM models of locomotor activity. 4

Supplementary Figure 4. Individual-cohort genome-wide association results for locomotor activity. 5

Supplementary Figure 5. Regional association plots for primary locomotor-activity QTLs. 7

Supplementary Figure 6. Cross-cohort effect structures at METAL- and Genomic SEM-supported loci. 114

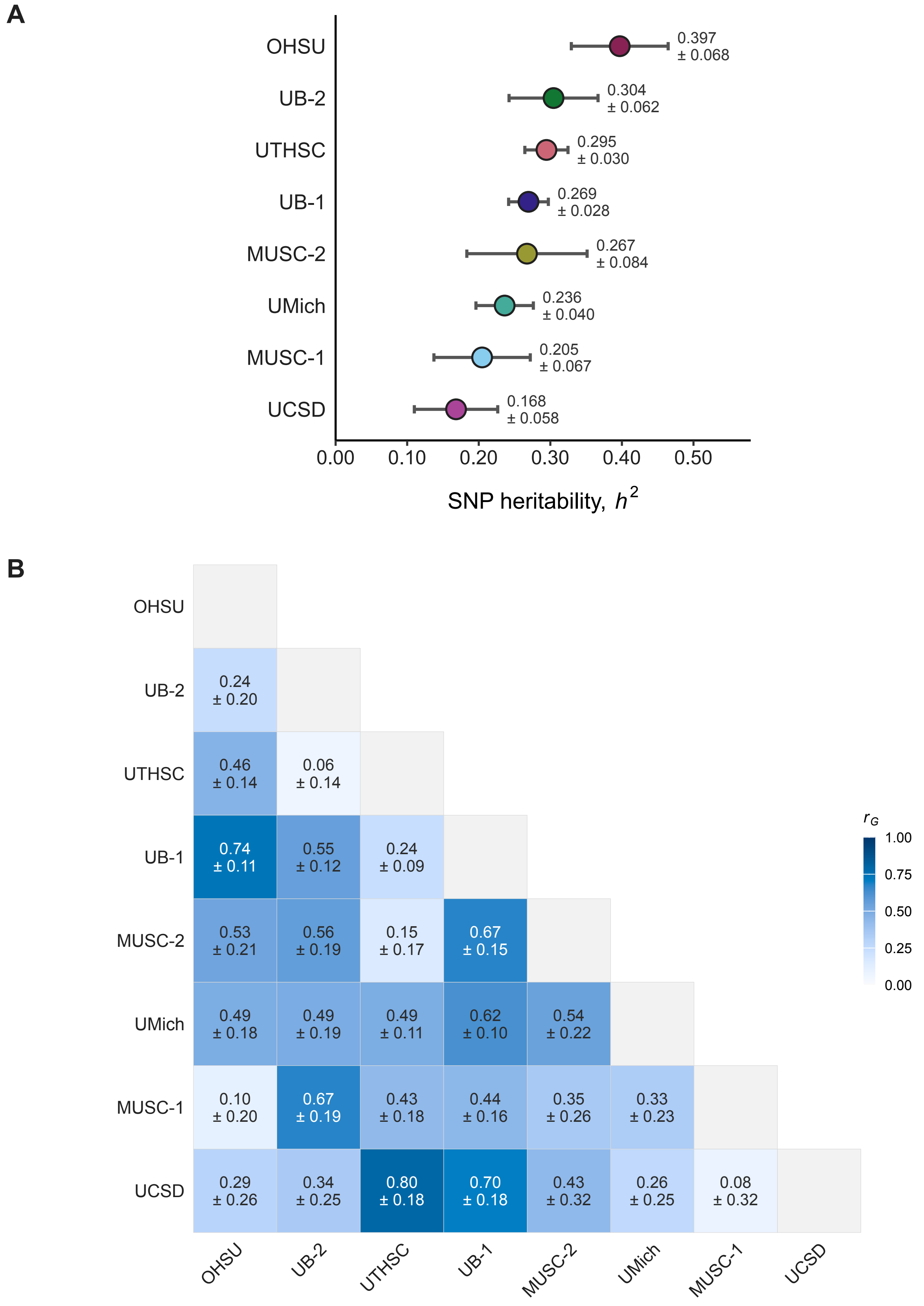


**Supplementary Figure 1. SNP heritability and pairwise genetic correlations across all eight cohorts.**

**(A)** Cohort-specific SNP-heritability estimates; points show $h^2$ estimates and horizontal bars indicate standard errors. **(B)** Pairwise genetic correlations among all eight cohorts; cells show $r_G$ ± SE, with color intensity indicating the magnitude of the genetic correlation. UCSD is included here for descriptive cohort-level analyses but was not included in the primary seven-cohort association analyses.

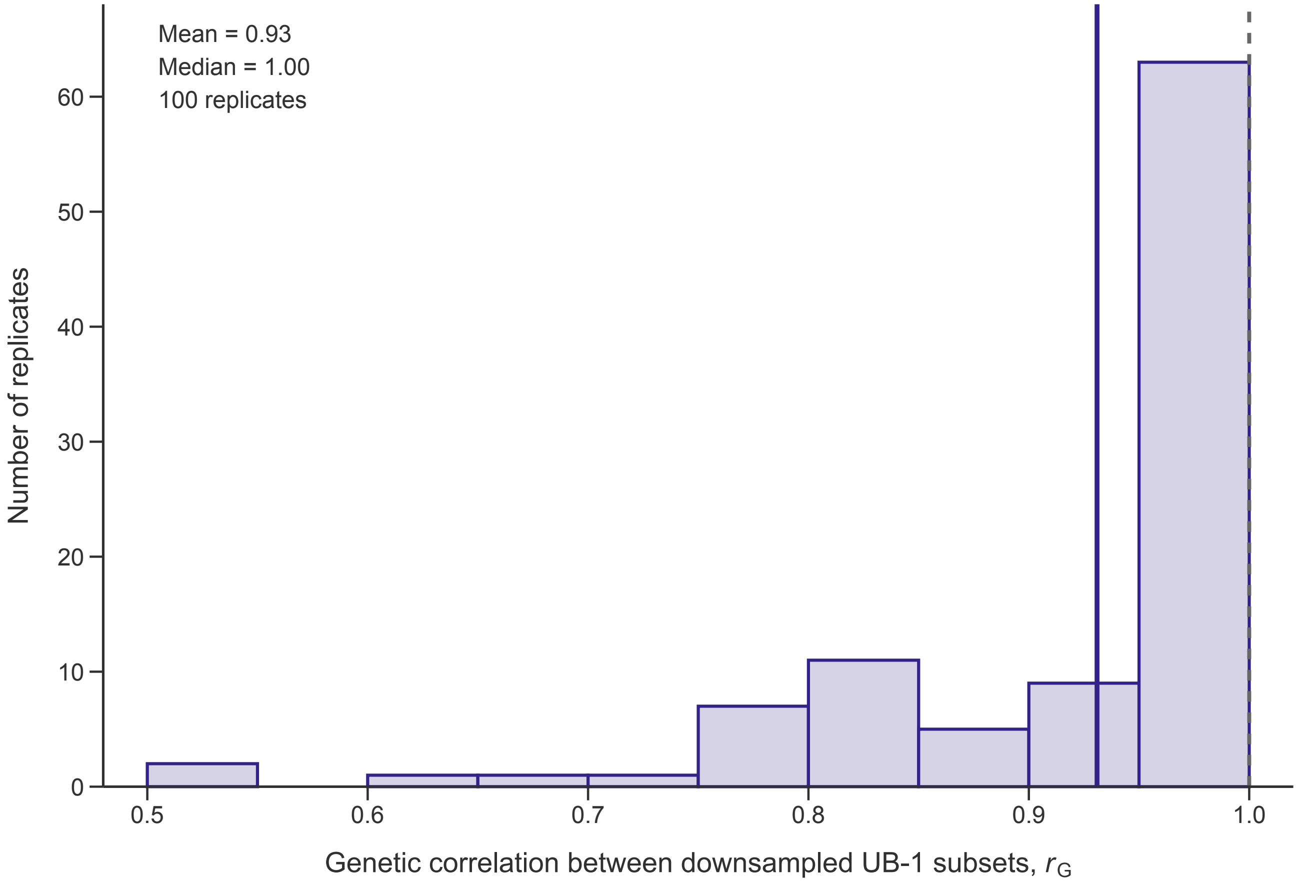


**Supplementary Figure 2. Repeated within-cohort genetic-correlation downsampling analysis in UB-1.**

In each of 100 repetitions, UB-1 was randomly divided into two nonoverlapping subsets of $N$ = 618 rats, and $r_G$ was estimated between the subsets. The histogram shows the resulting distribution; vertical lines mark the mean estimate (0.93) and $r_G$ = 1.00. Estimates at 1.00 reached the upper boundary of the parameter space.

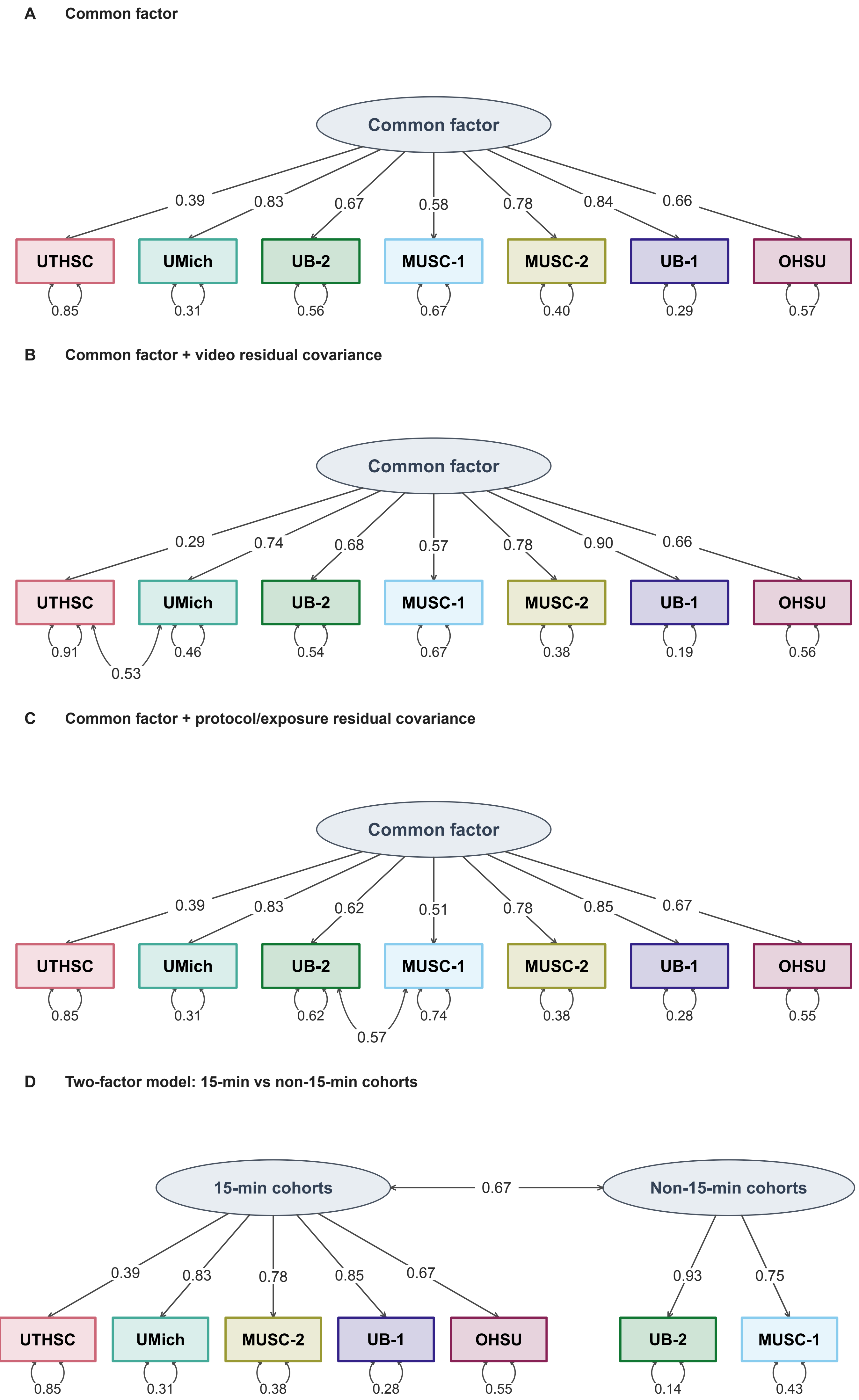


**Supplementary Figure 3. Alternative seven-trait Genomic SEM models of locomotor activity.**

**(A)** Common-factor model. **(B)** Common-factor model with residual genetic covariance between UTHSC and UMich. **(C)** Common-factor model with residual genetic covariance between UB-2 and MUSC-1. **(D)** Two-factor model separating the five 15-min cohorts from the two non-15-min cohorts. Values on single-headed paths are standardized factor loadings, self-loops show standardized residual variances, and double-headed paths show residual covariances or the standardized correlation between latent factors. The selected model containing both residual covariances is shown in Figure 1C.

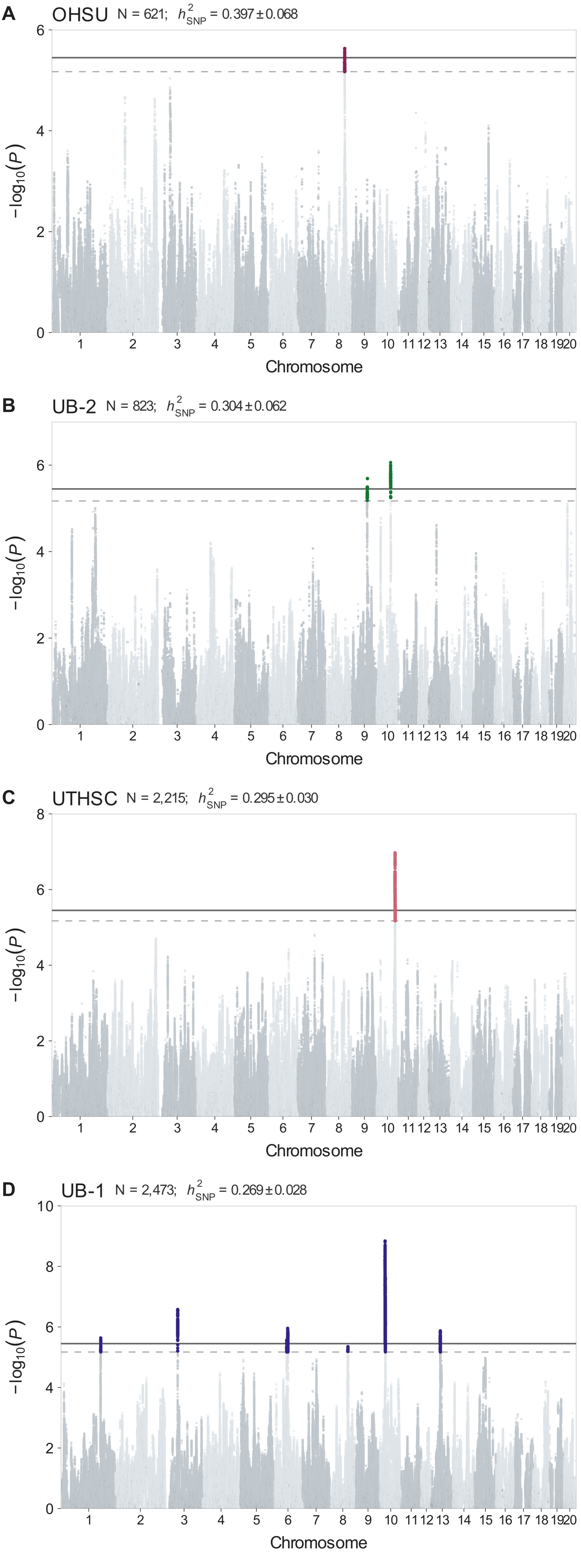


**Supplementary Figure 4. Individual-cohort genome-wide association results for locomotor activity.**

Manhattan plots show cohort-specific association results for **(A)** OHSU, **(B)** UB-2, **(C)** UTHSC, **(D)** UB-1, **(E)** MUSC-2, **(F)** UMich, **(G)** MUSC-1, and **(H)** UCSD. Each panel reports *N* and SNP heritability ($h^2$). SNPs meeting the simulation-based empirical 10% discovery threshold are highlighted. Solid and dashed horizontal lines indicate the simulation-based empirical 5% ($-\log_{10}(P)$ = 5.449) and 10% ($-\log_{10}(P)$ = 5.171) discovery thresholds, respectively. UCSD is shown here as a cohort-specific analysis but was not included in the primary seven-cohort MEGA, METAL, or Genomic SEM analyses.

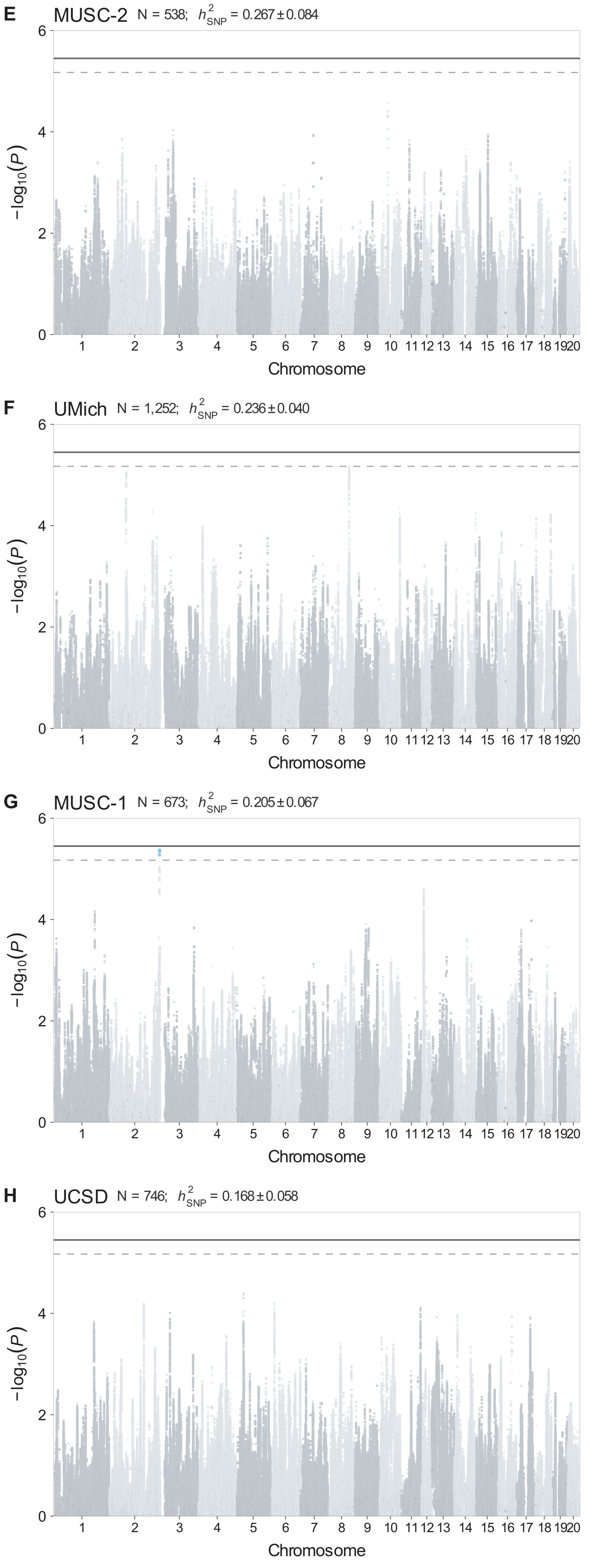

E
MUSC-2 N = 538; $h^2_{SNP}$ = 0.267 ± 0.084
F
UMich N = 1,252; $h^2_{SNP}$ = 0.236 ± 0.040
G
MUSC-1 N = 673; $h^2_{SNP}$ = 0.205 ± 0.067
H
UCSD N = 746; $h^2_{SNP}$ = 0.168 ± 0.058
−log10(P)
0
2
4
6
1 2 3 4 5 6 7 8 9 10 11 12 13 14 15 16 17 18 19 20
Chromosome

**Supplementary Figure 5. Regional association plots for primary locomotor-activity QTLs.**

Narrow and broad regional association plots are shown for primary QTLs not displayed in Figure 4, with separate method-specific views where multiple primary methods support the same QTL. Exact method-specific lead SNPs are listed above each plot. Association significance is plotted as $-\log_{10}(P)$ against rn8 genomic position, with points colored according to linkage disequilibrium ($r^2$) with the corresponding lead SNP. Solid and dashed horizontal lines indicate the simulation-based empirical 5% ($-\log_{10}(P) = 5.449$) and 10% ($-\log_{10}(P) = 5.171$) discovery thresholds, respectively. Headers identify the QTL interval and gene count, and NCBI RefSeq gene models provide local genomic context. For gene-dense regions, broad gene tracks continue on pages labeled “broad (continued).”

**chr1:196.1 Mb - MEGA - narrow**

Lead: 1:196,086,143 | Genes in QTL interval: 23

QTL interval: chr1:194,427,078-196,396,744

Displayed region: chr1:194,089,152-196,596,744

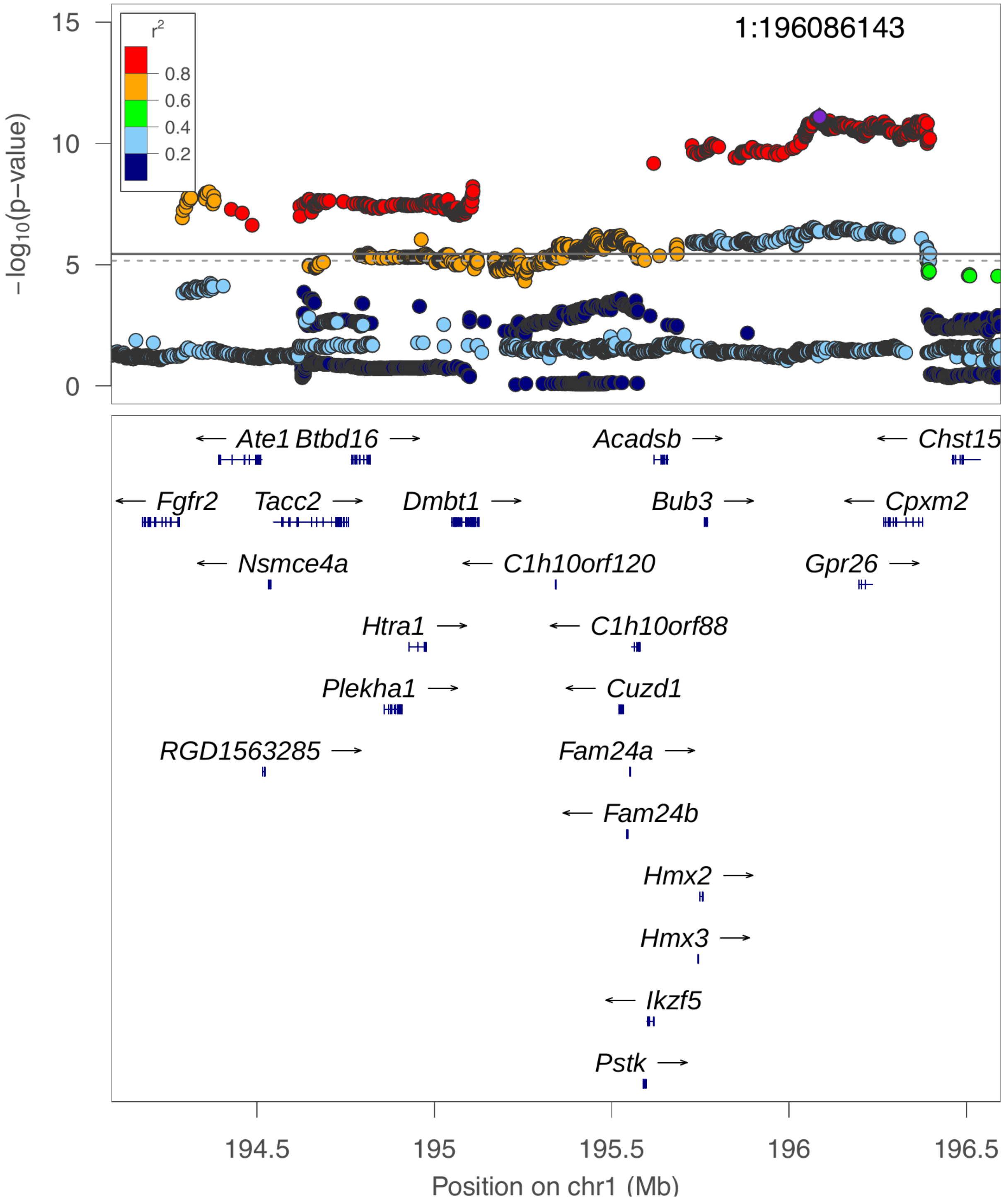

**chr1:196.1 Mb - MEGA - broad**

Lead: 1:196,086,143 | Genes in QTL interval: 23

QTL interval: chr1:194,427,078-196,396,744

Displayed region: chr1:191,289,152-199,396,744

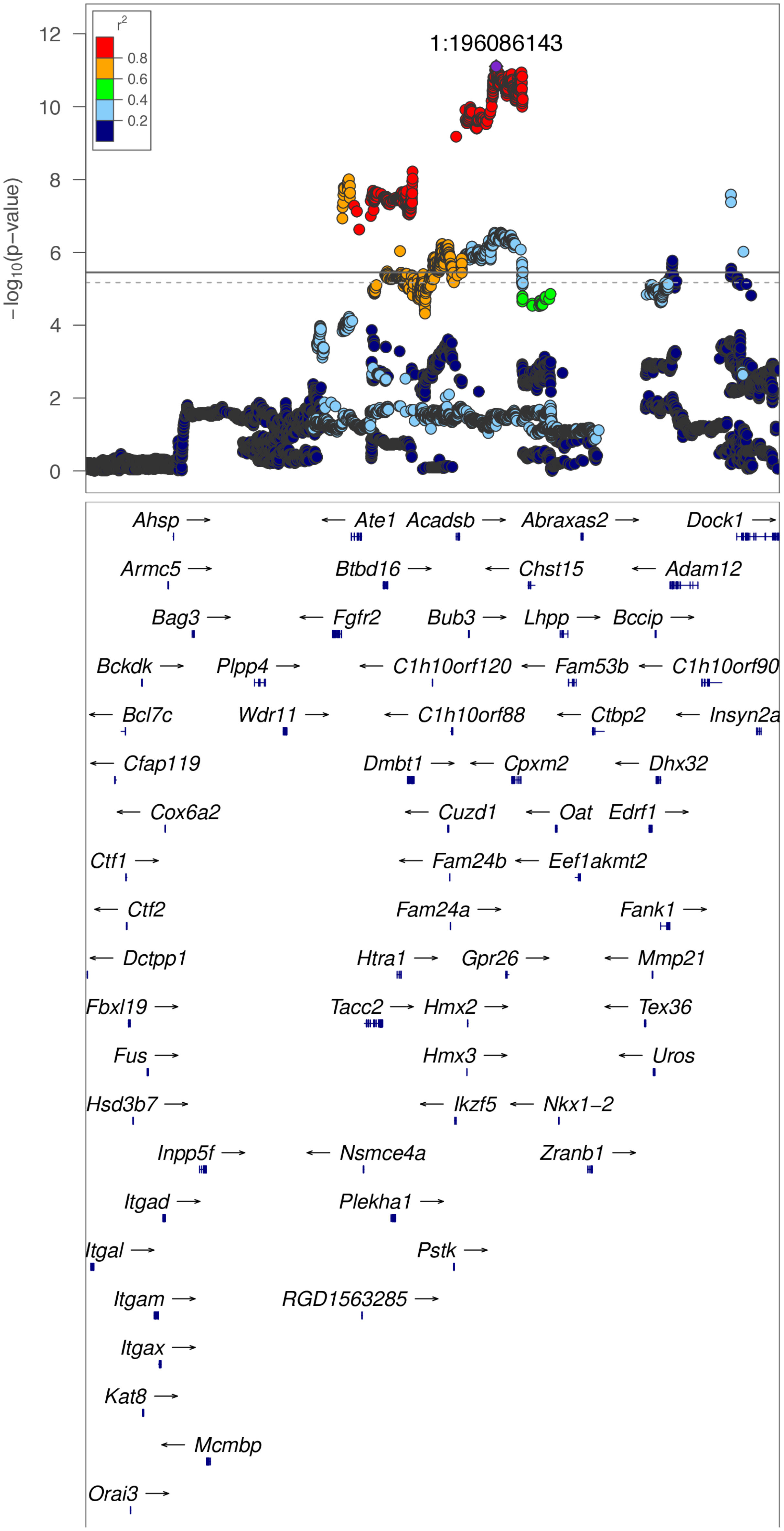

**chr1:196.1 Mb - MEGA - broad (continued)**

Lead: 1:196,086,143 | Genes in QTL interval: 23

QTL interval: chr1:194,427,078-196,396,744

Displayed region: chr1:191,289,152-199,396,744

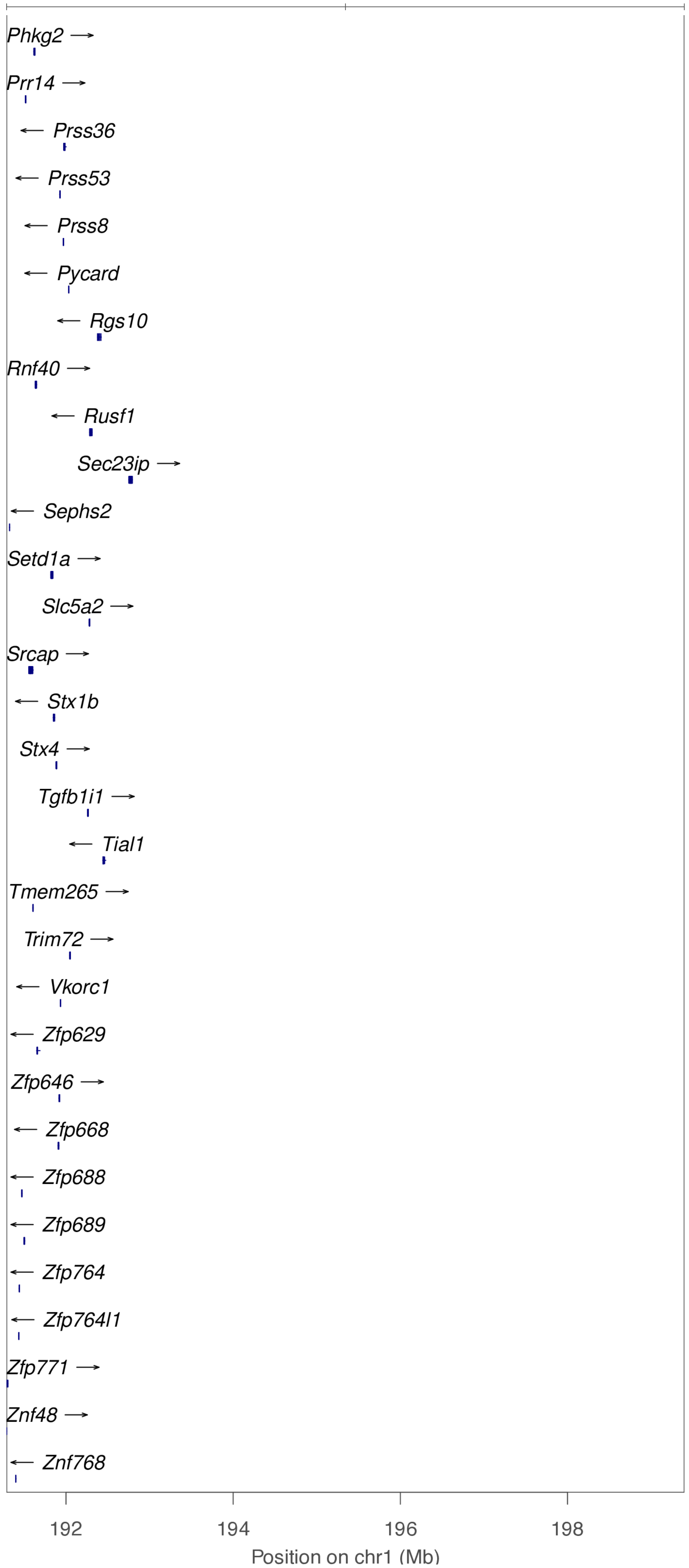

**chr1:196.1 Mb - METAL - narrow**

Lead: 1:196,367,786 | Genes in QTL interval: 23

QTL interval: chr1:194,427,078-196,396,744

Displayed region: chr1:194,086,781-196,596,744

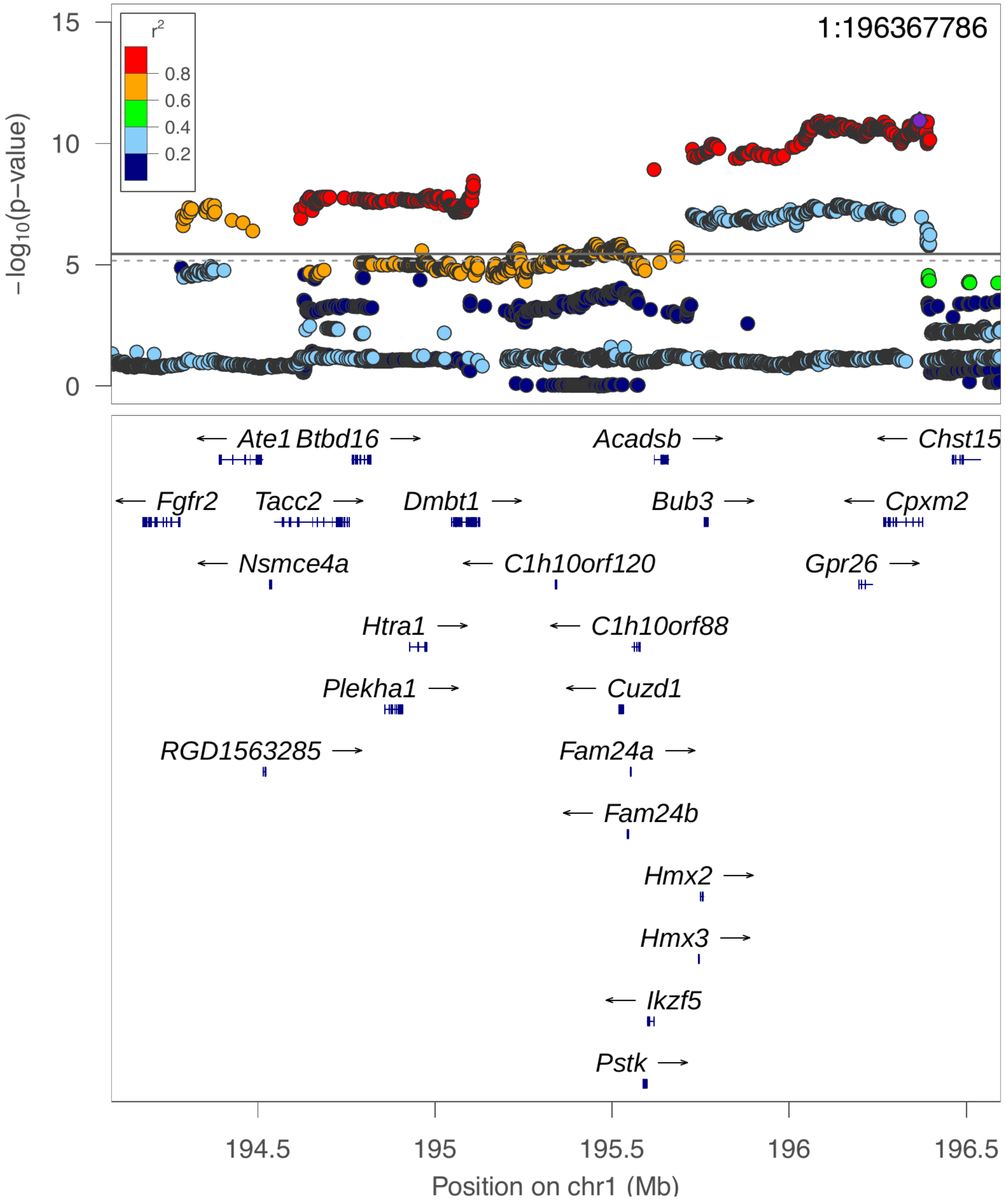

**chr1:196.1 Mb - METAL - broad**

Lead: 1:196,367,786 | Genes in QTL interval: 23

QTL interval: chr1:194,427,078-196,396,744

Displayed region: chr1:191,286,781-199,396,744

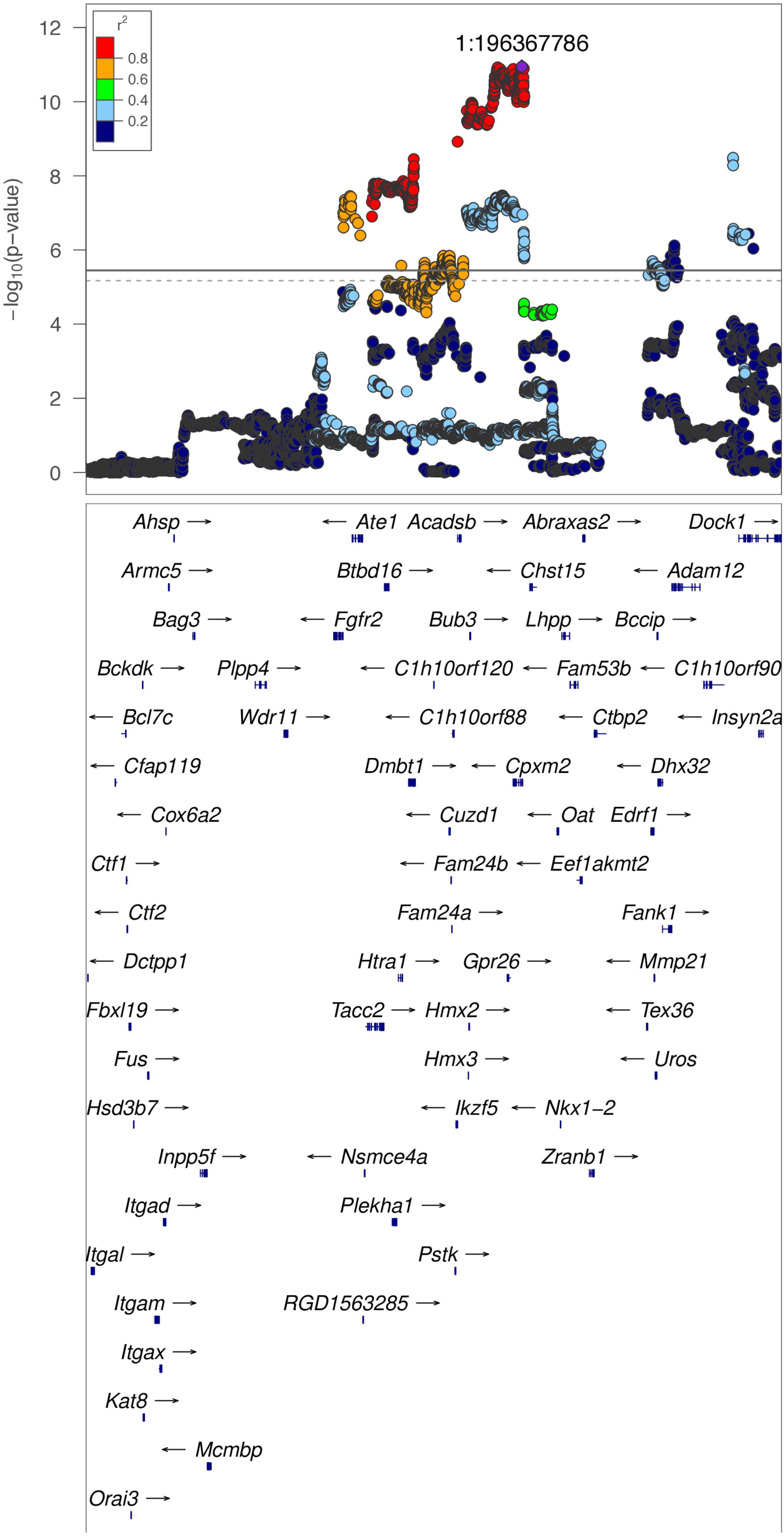

**chr1:196.1 Mb - METAL - broad (continued)**

Lead: 1:196,367,786 | Genes in QTL interval: 23

QTL interval: chr1:194,427,078-196,396,744

Displayed region: chr1:191,286,781-199,396,744

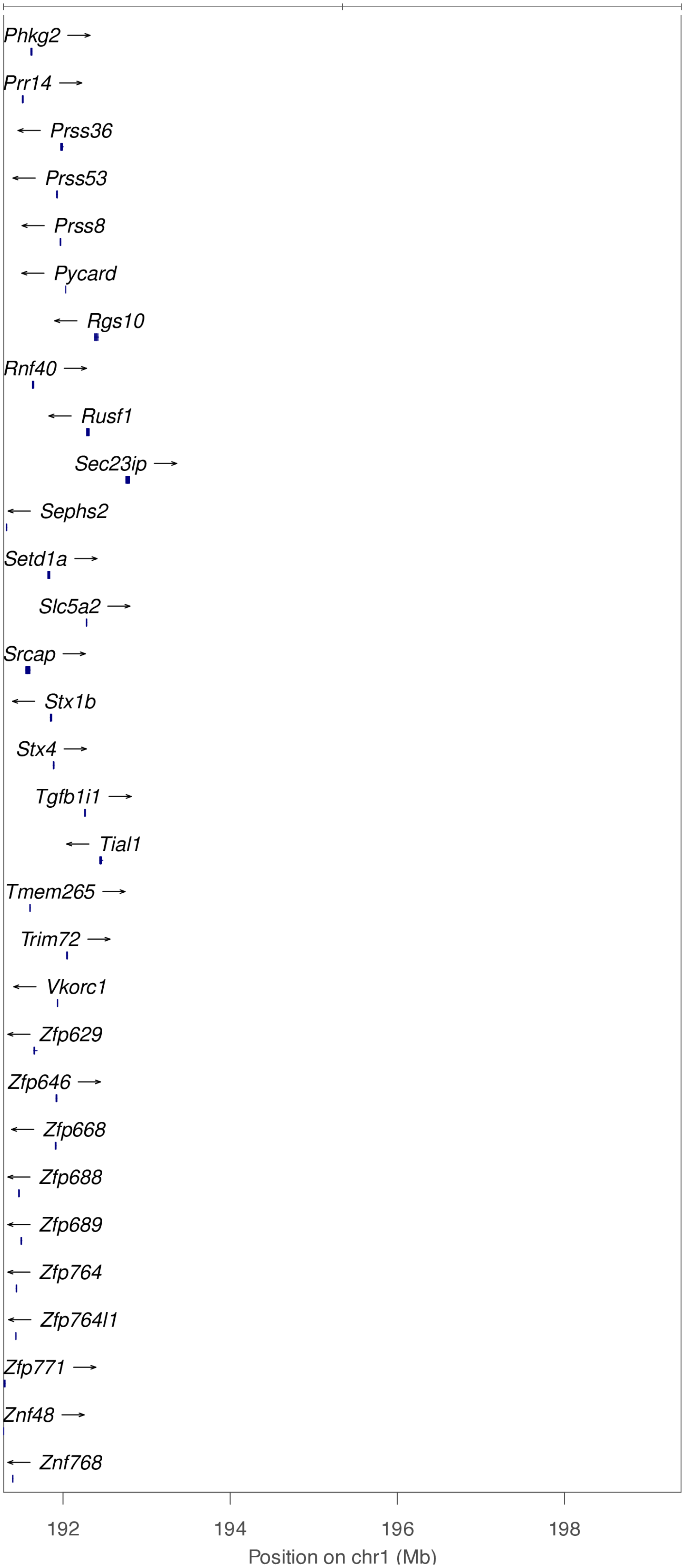

**chr1:196.1 Mb - Genomic SEM - narrow**

Lead: 1:196,113,797 | Genes in QTL interval: 23

QTL interval: chr1:194,427,078-196,396,744

Displayed region: chr1:194,086,781-196,596,744

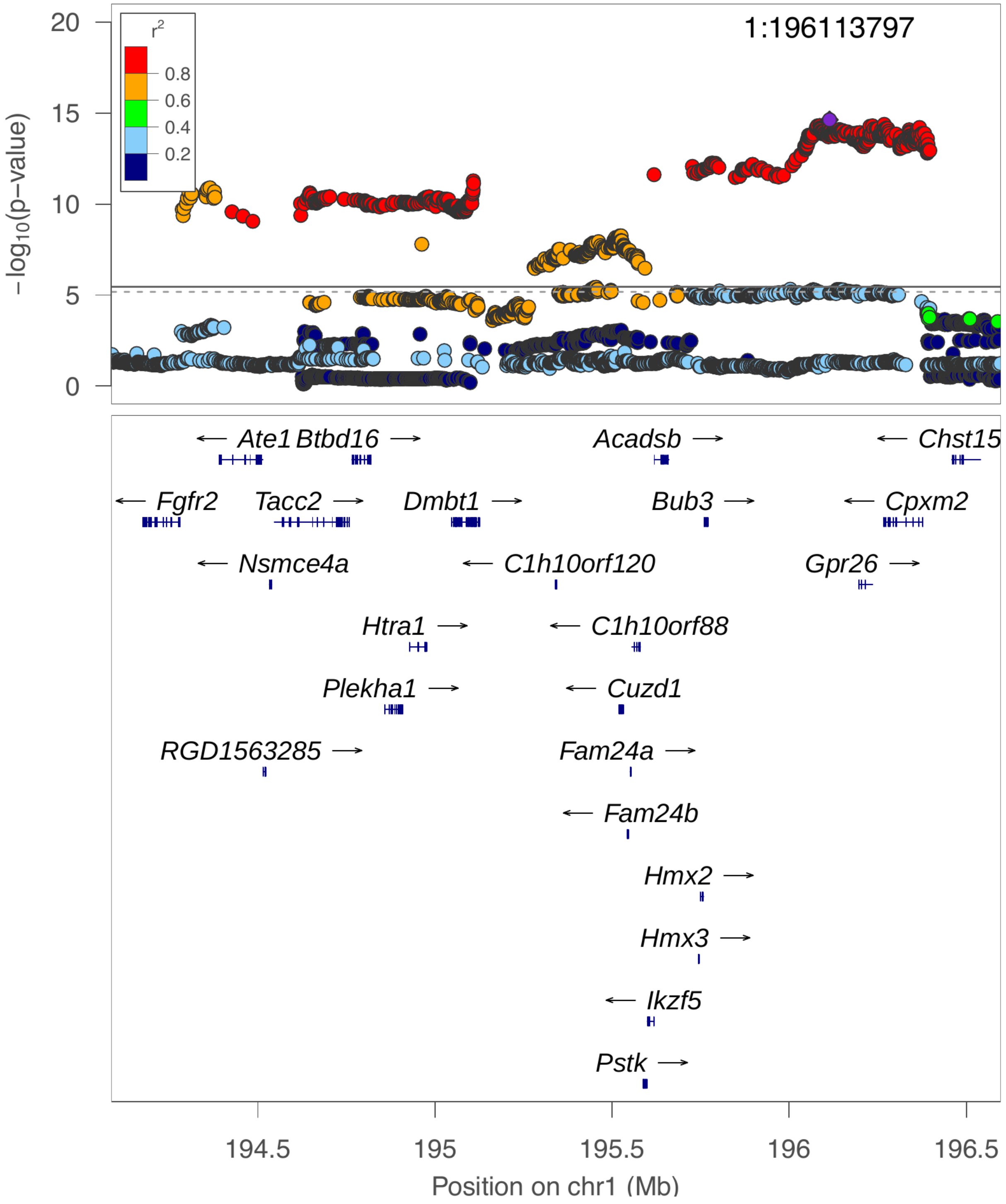

**chr1:196.1 Mb - Genomic SEM - broad**

Lead: 1:196,113,797 | Genes in QTL interval: 23

QTL interval: chr1:194,427,078-196,396,744

Displayed region: chr1:191,286,781-199,396,744

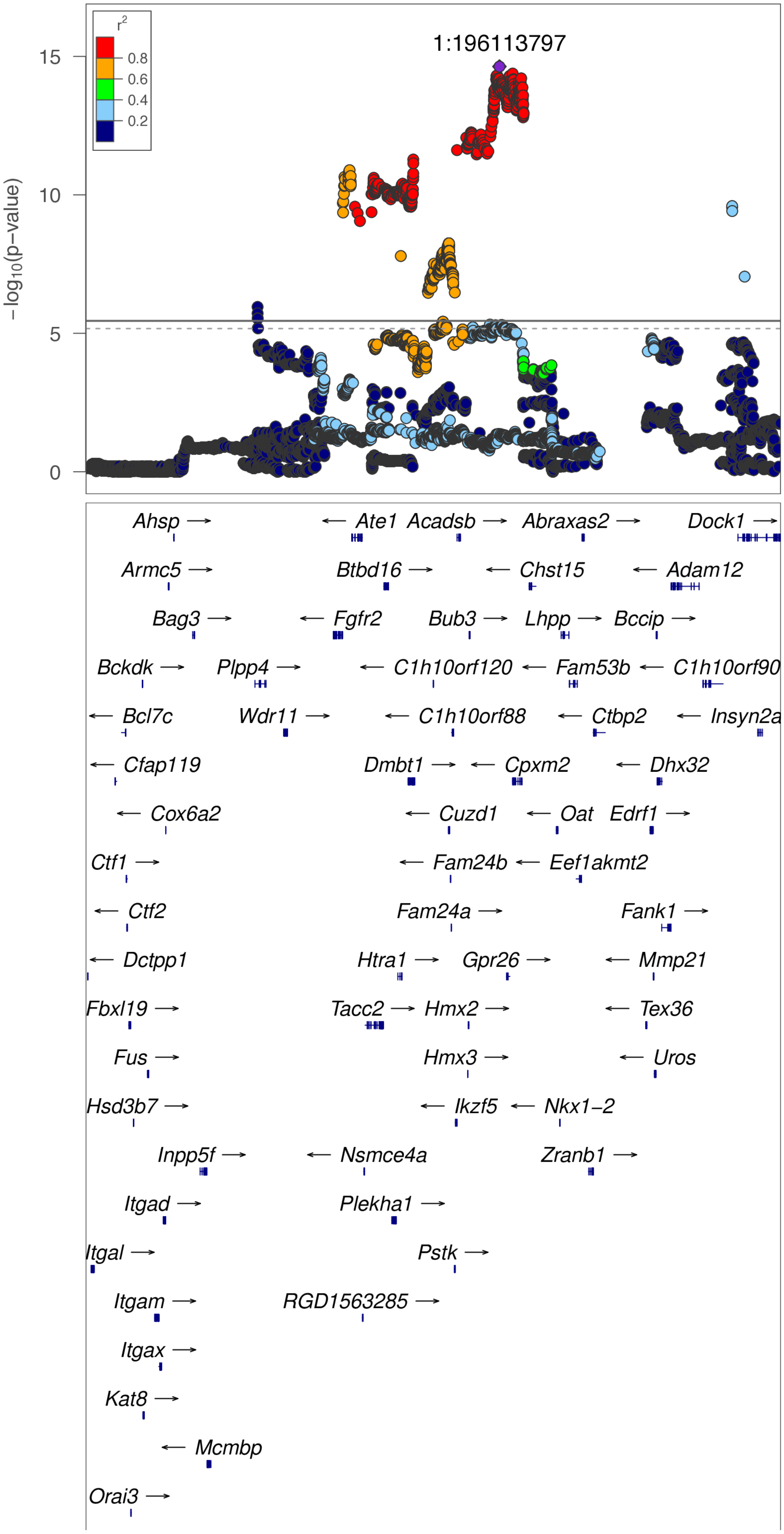

**chr1:196.1 Mb - Genomic SEM - broad (continued)**

Lead: 1:196,113,797 | Genes in QTL interval: 23

QTL interval: chr1:194,427,078-196,396,744

Displayed region: chr1:191,286,781-199,396,744

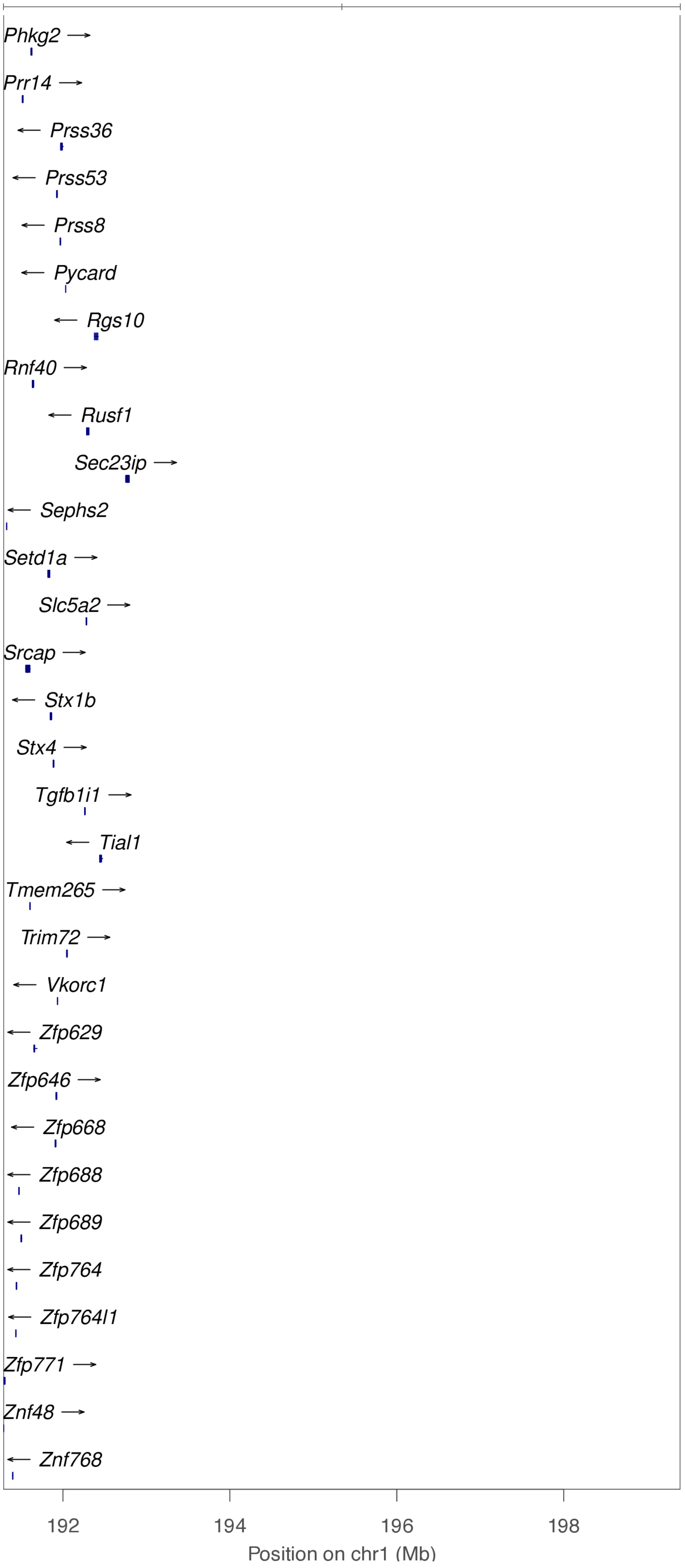

**chr2:30.0 Mb - Genomic SEM - narrow**

Lead: 2:30,004,505 | Genes in QTL interval: 10

QTL interval: chr2:29,601,527-30,444,766

Displayed region: chr2:29,422,877-30,924,904

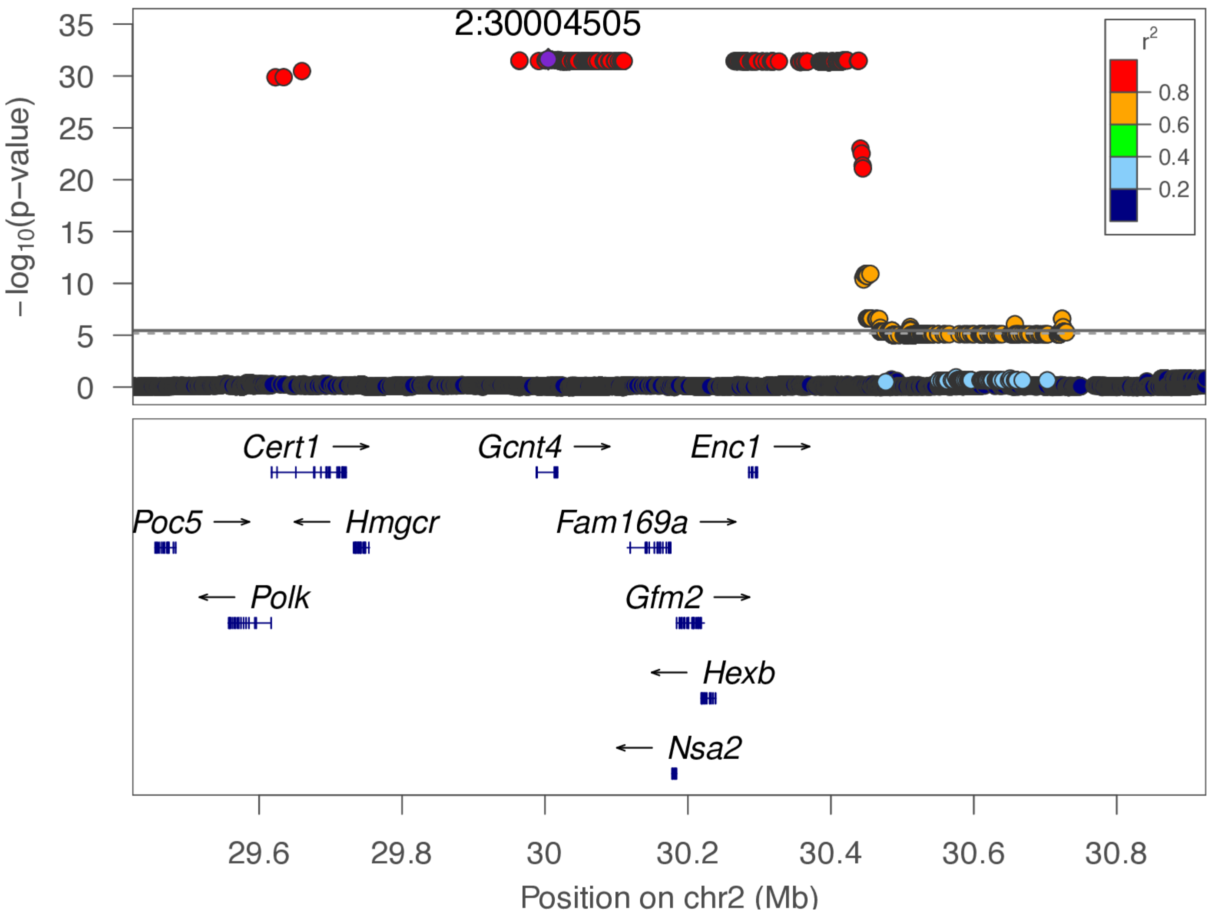

**chr2:30.0 Mb - Genomic SEM - broad**

Lead: 2:30,004,505 | Genes in QTL interval: 10

QTL interval: chr2:29,601,527-30,444,766

Displayed region: chr2:26,622,877-33,724,904

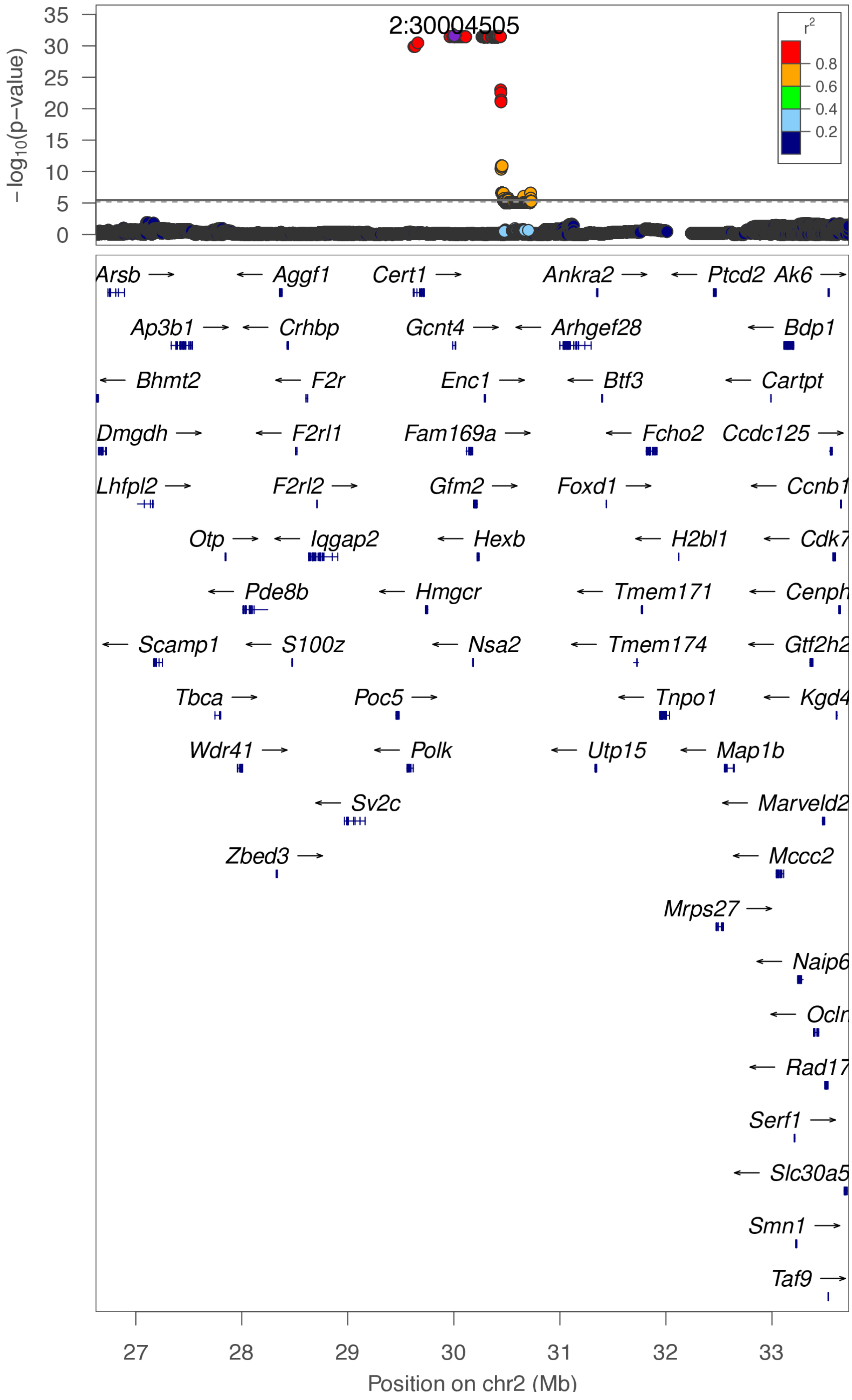

**chr2:87.3 Mb - Genomic SEM - narrow**

Lead: 2:87,278,158 | Genes in QTL interval: 28

QTL interval: chr2:85,729,807-90,636,632

Displayed region: chr2:85,529,807-90,836,632

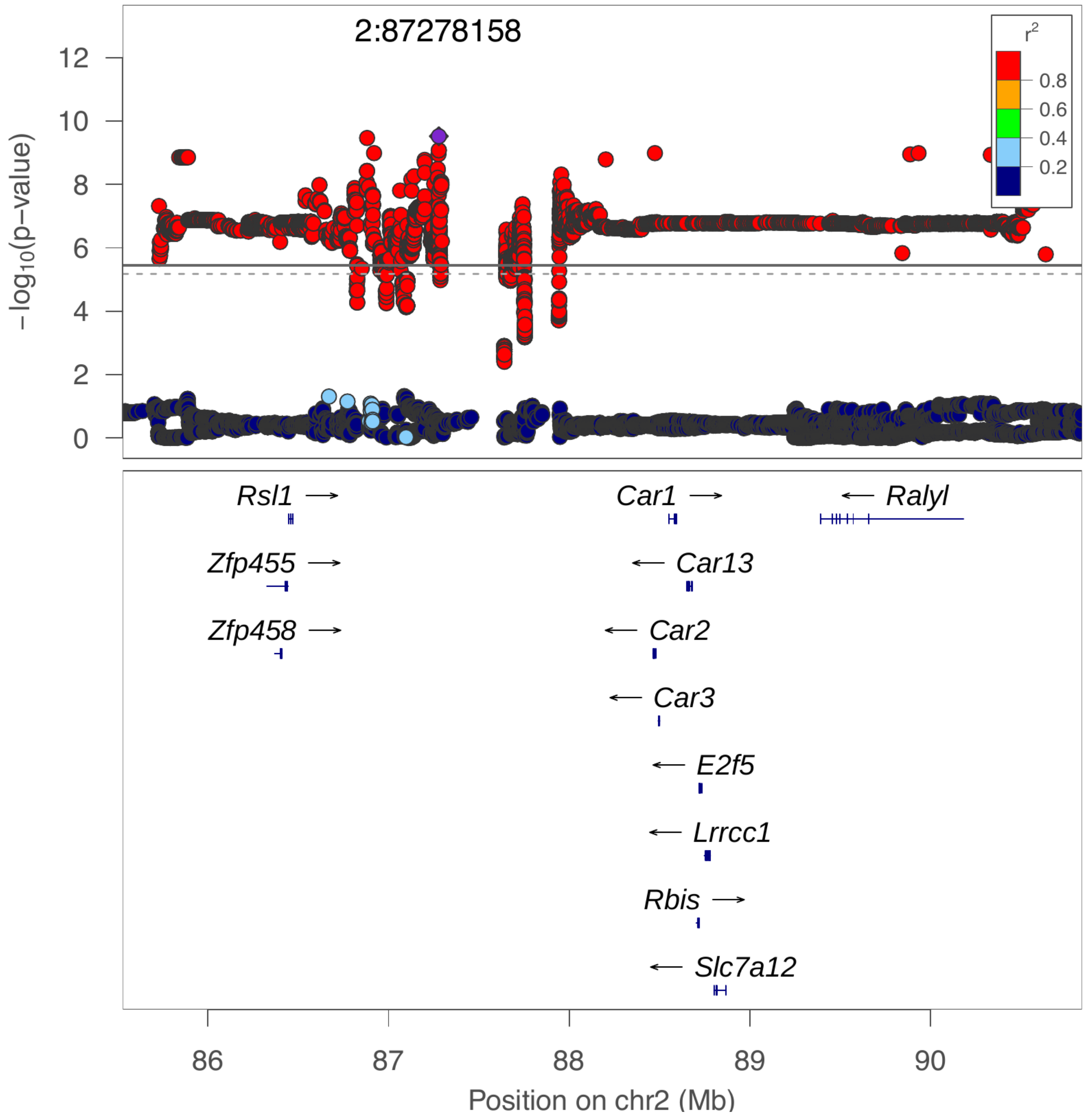

**chr2:87.3 Mb - Genomic SEM - broad**

Lead: 2:87,278,158 | Genes in QTL interval: 28

QTL interval: chr2:85,729,807-90,636,632

Displayed region: chr2:82,729,807-93,636,632

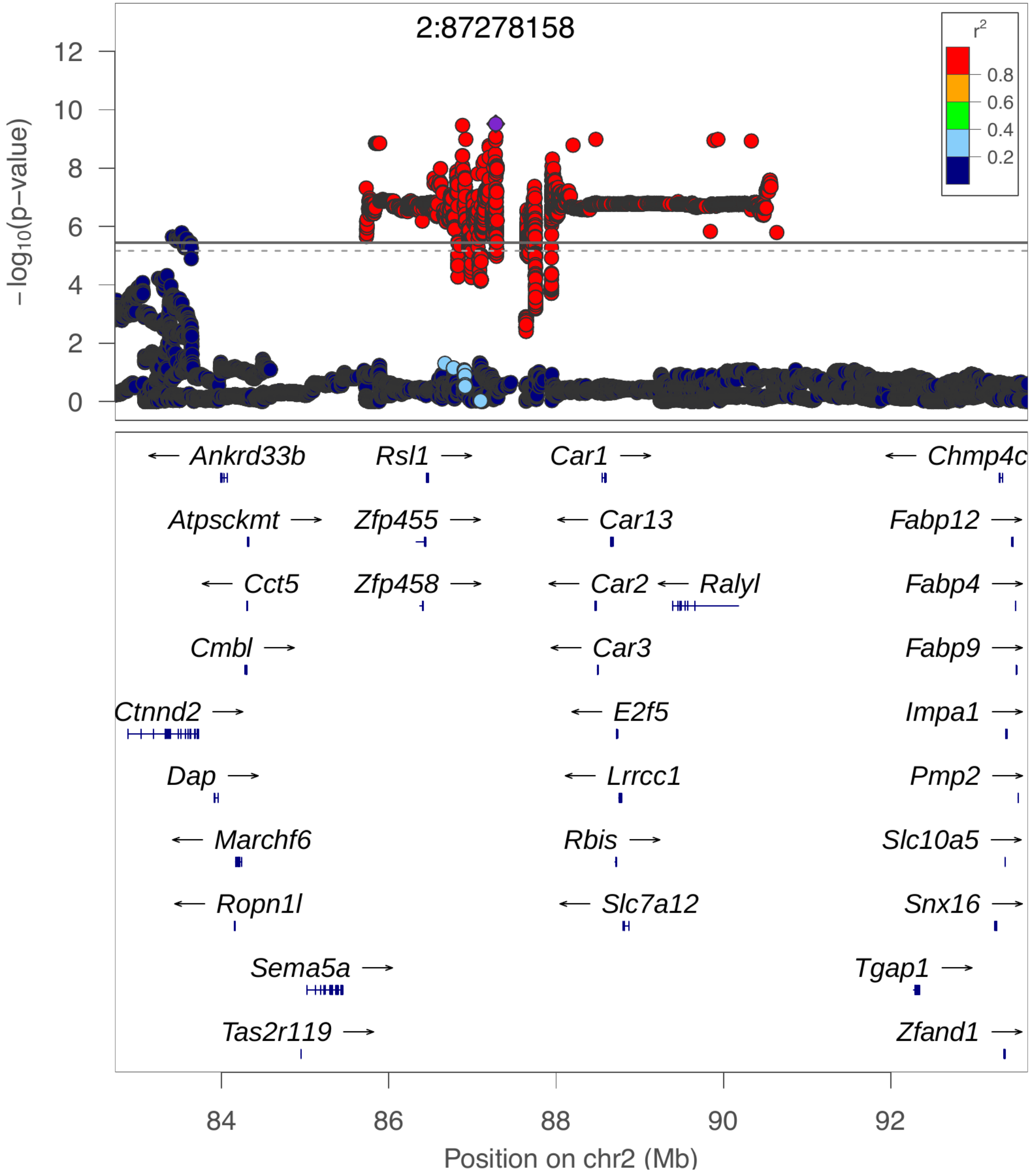

**chr2:124.0 Mb - METAL - narrow**

Lead: 2:124,226,433 | Genes in QTL interval: 5

QTL interval: chr2:122,227,357-124,707,622

Displayed region: chr2:119,910,662-130,098,863

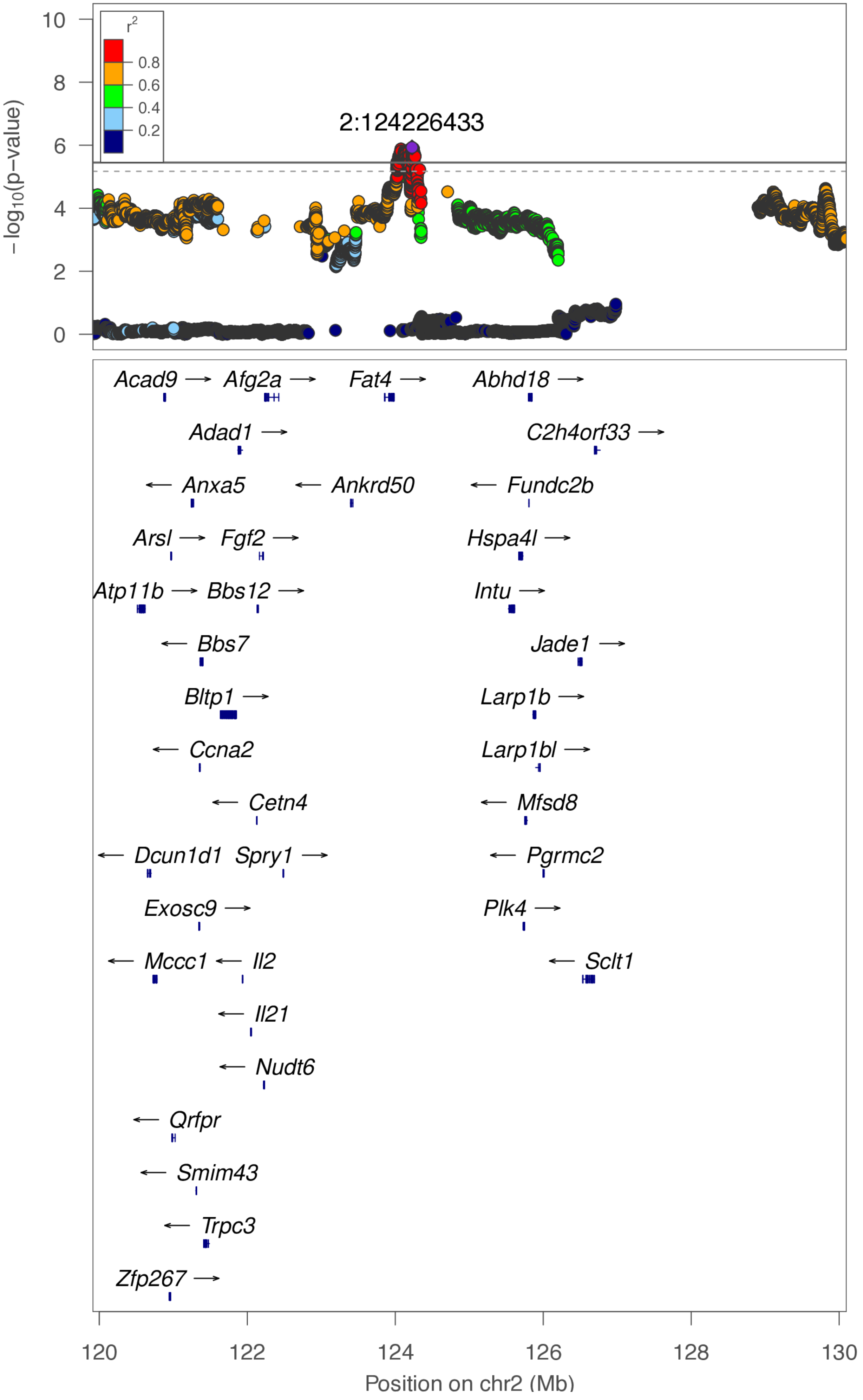

**chr2:124.0 Mb - METAL - broad**

Lead: 2:124,226,433 | Genes in QTL interval: 5

QTL interval: chr2:122,227,357-124,707,622

Displayed region: chr2:117,110,662-132,898,863

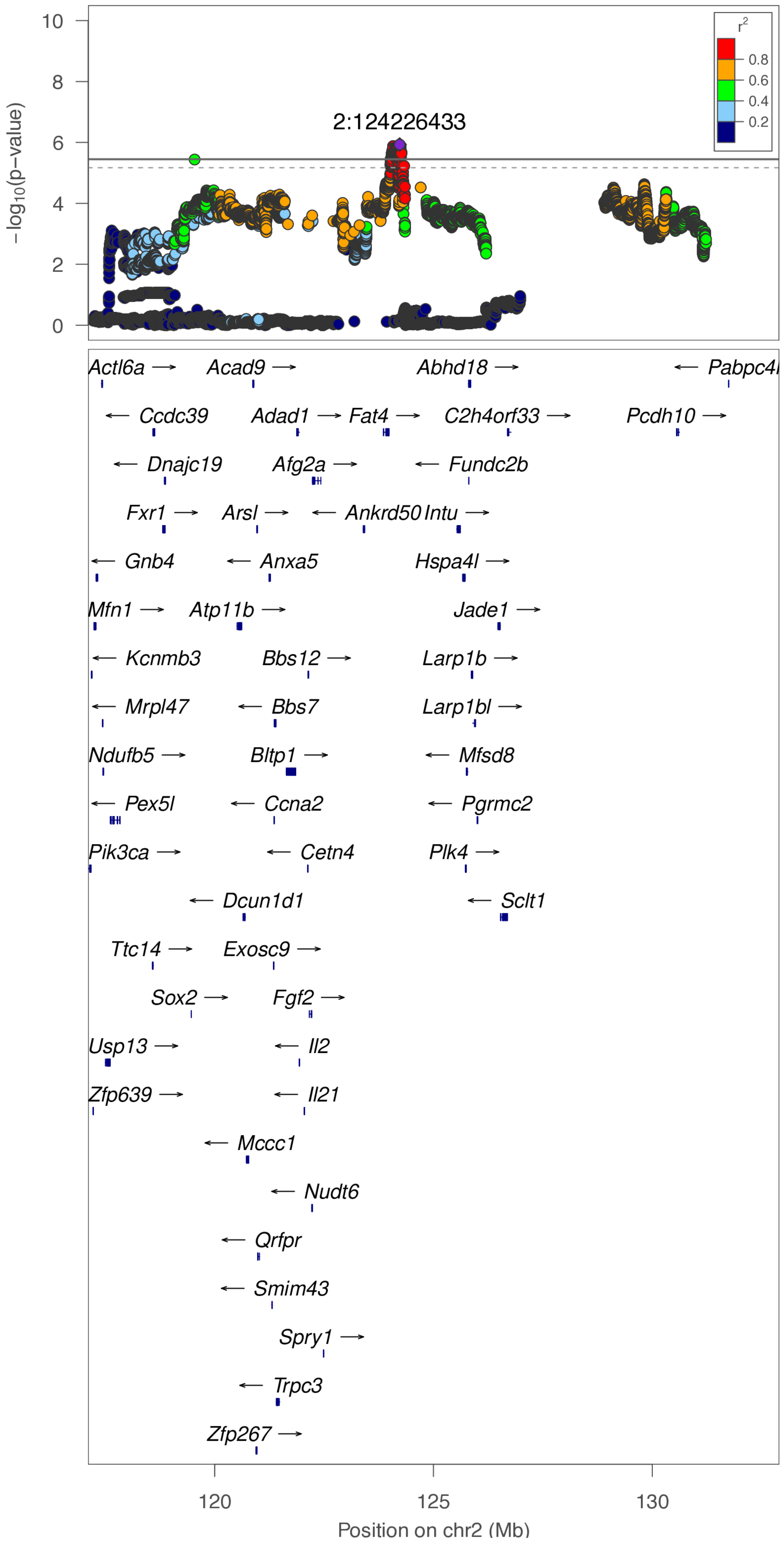

**chr2:124.0 Mb - Genomic SEM - narrow**

Lead: 2:124,046,350 | Genes in QTL interval: 5

QTL interval: chr2:122,227,357-124,707,622

Displayed region: chr2:119,925,681-129,980,432

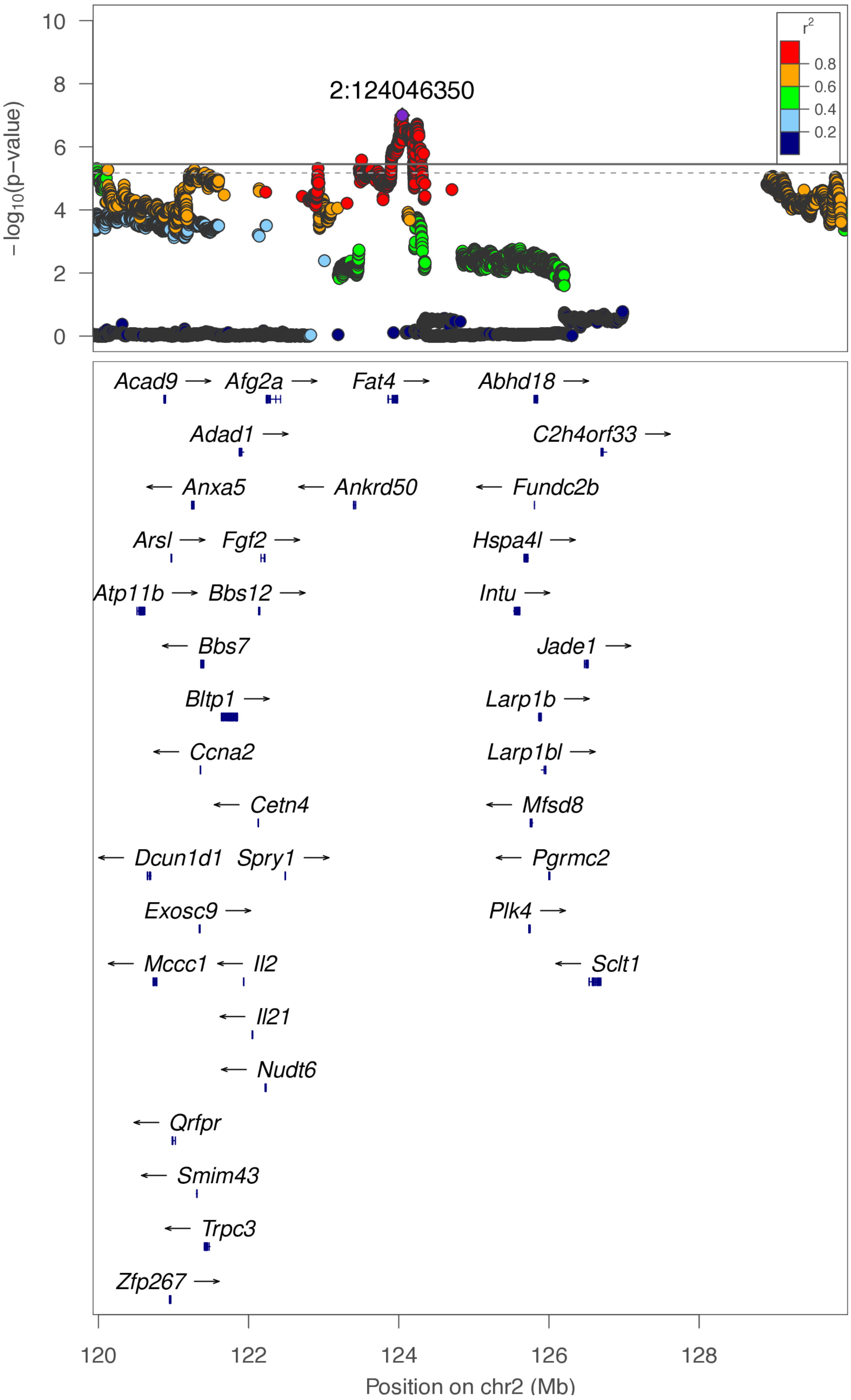

**chr2:124.0 Mb - Genomic SEM - broad**

Lead: 2:124,046,350 | Genes in QTL interval: 5

QTL interval: chr2:122,227,357-124,707,622

Displayed region: chr2:117,125,681-132,780,432

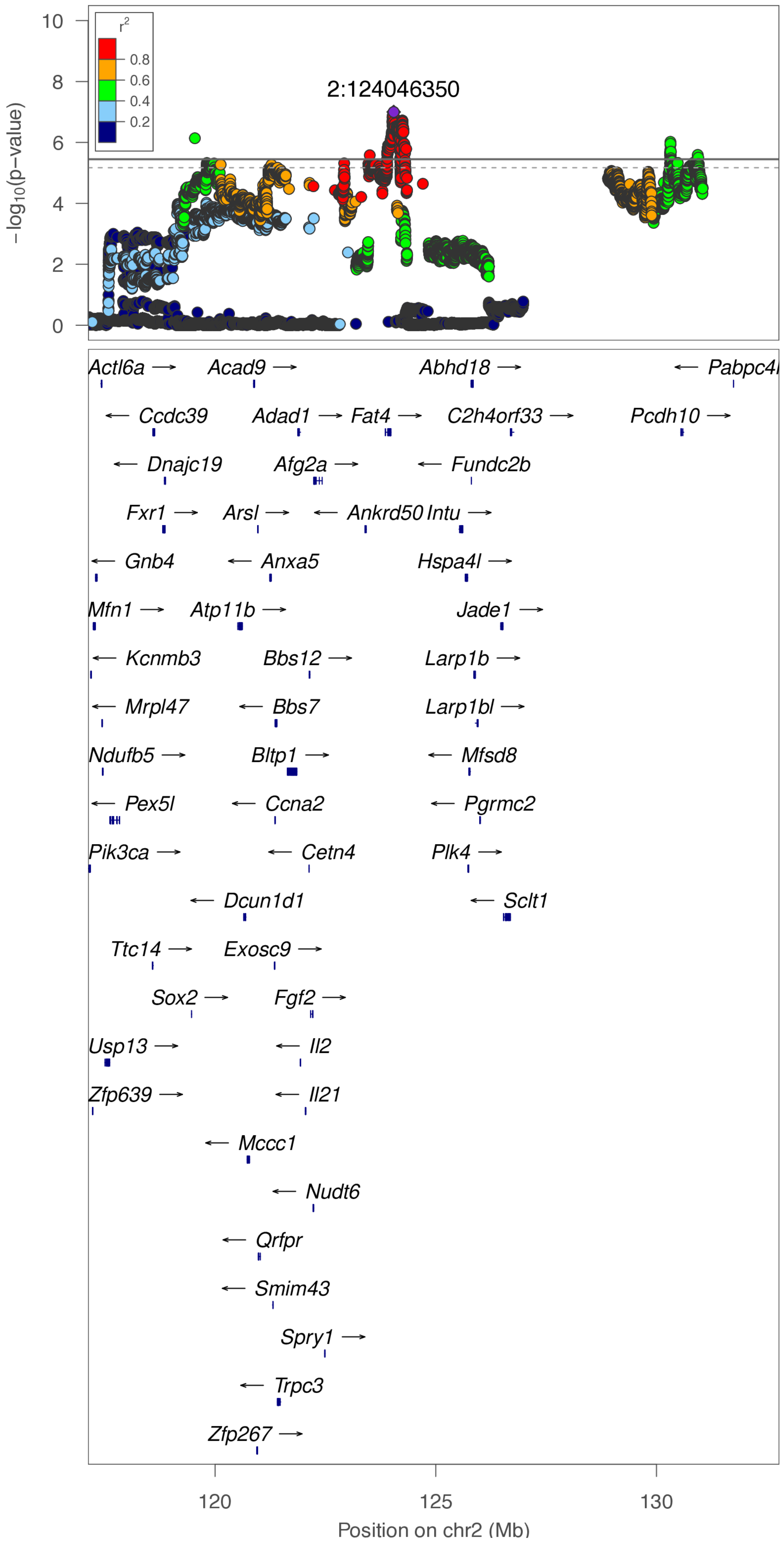

**chr2:130.3 Mb - Genomic SEM - narrow**

Lead: 2:130,316,468 | Genes in QTL interval: 1

QTL interval: chr2:129,777,962-131,183,341

Displayed region: chr2:128,702,959-131,586,667

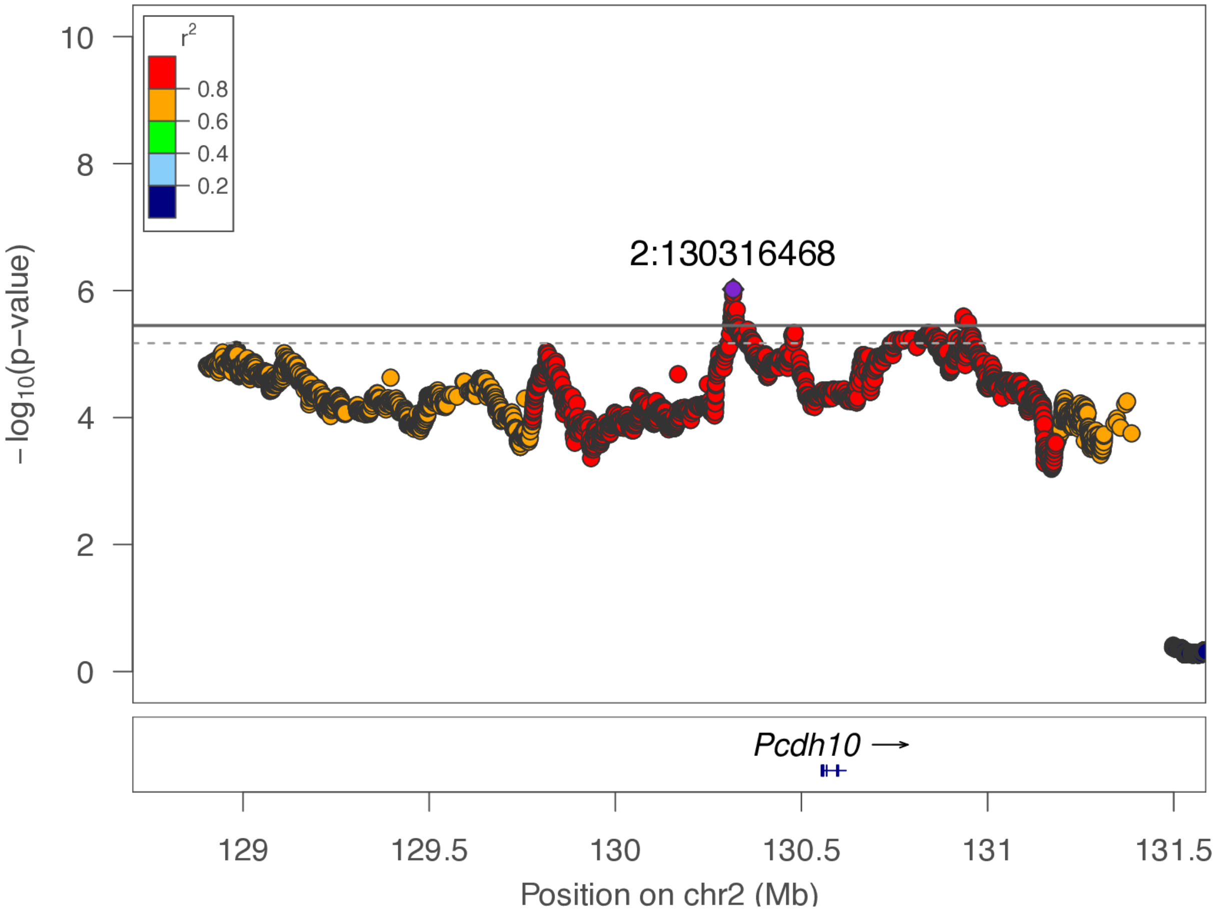

**chr2:130.3 Mb - Genomic SEM - broad**

Lead: 2:130,316,468 | Genes in QTL interval: 1

QTL interval: chr2:129,777,962-131,183,341

Displayed region: chr2:125,902,959-134,386,667

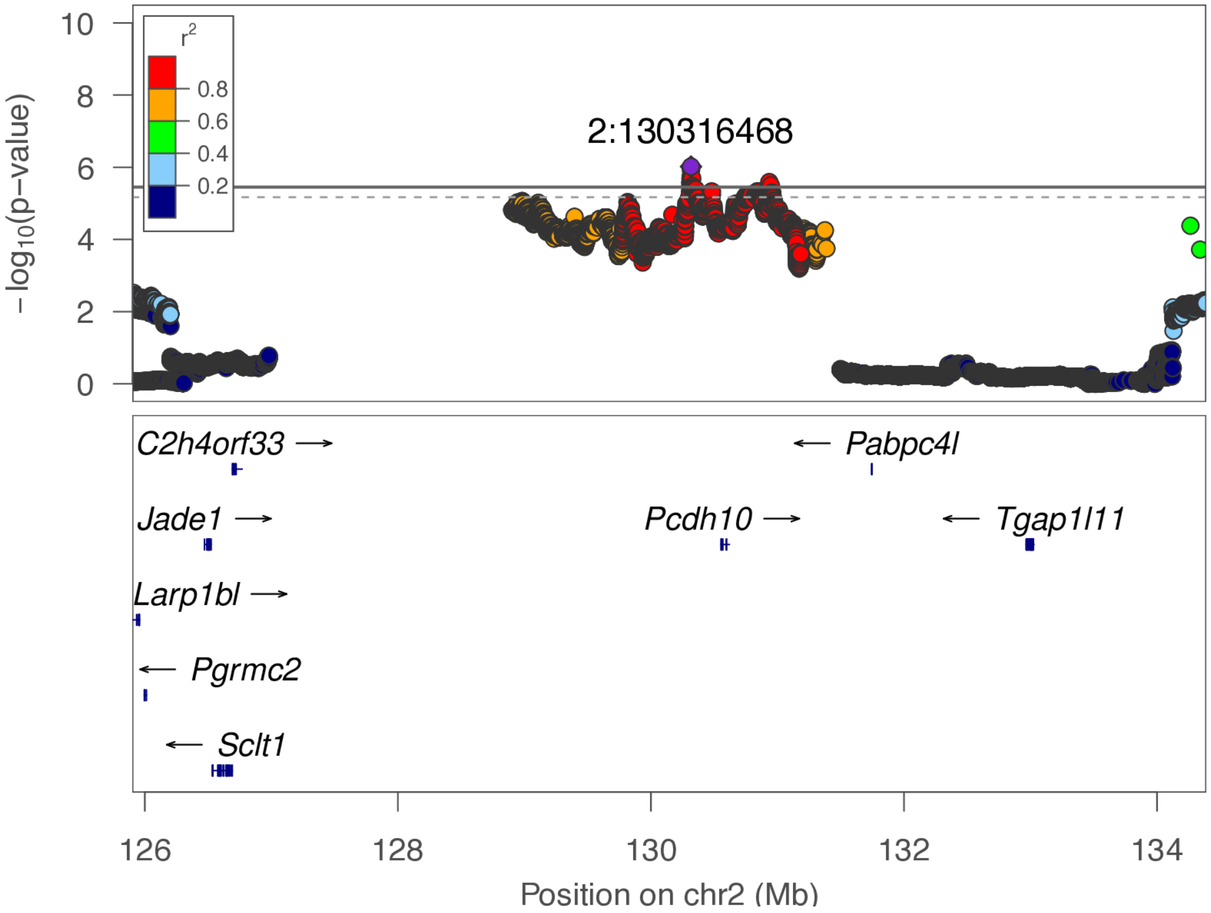

**chr2:226.7 Mb - Genomic SEM - narrow**

Lead: 2:226,722,630 | Genes in QTL interval: 9

QTL interval: chr2:226,372,866-226,894,162

Displayed region: chr2:225,861,681-227,094,162

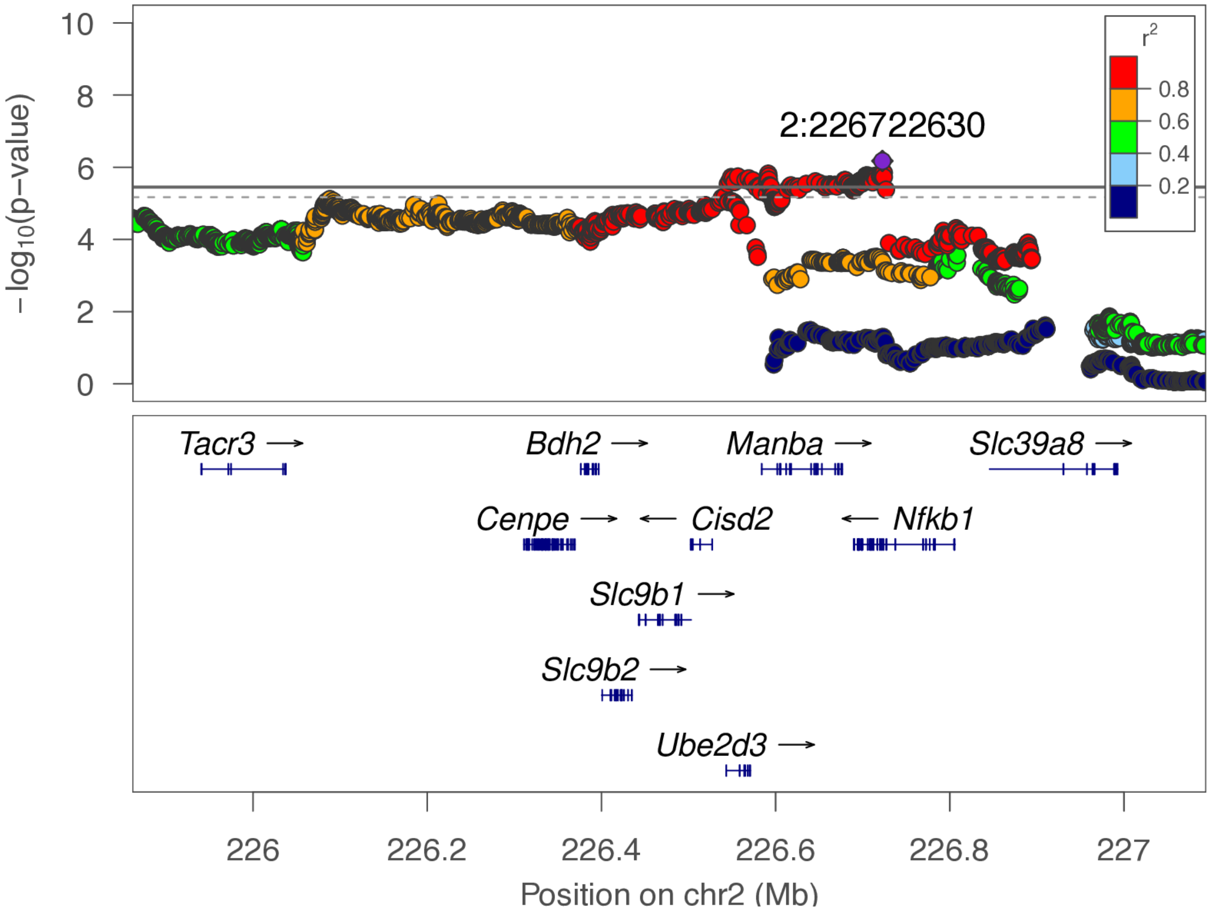

**chr2:226.7 Mb - Genomic SEM - broad**

Lead: 2:226,722,630 | Genes in QTL interval: 9

QTL interval: chr2:226,372,866-226,894,162

Displayed region: chr2:223,061,681-229,894,162

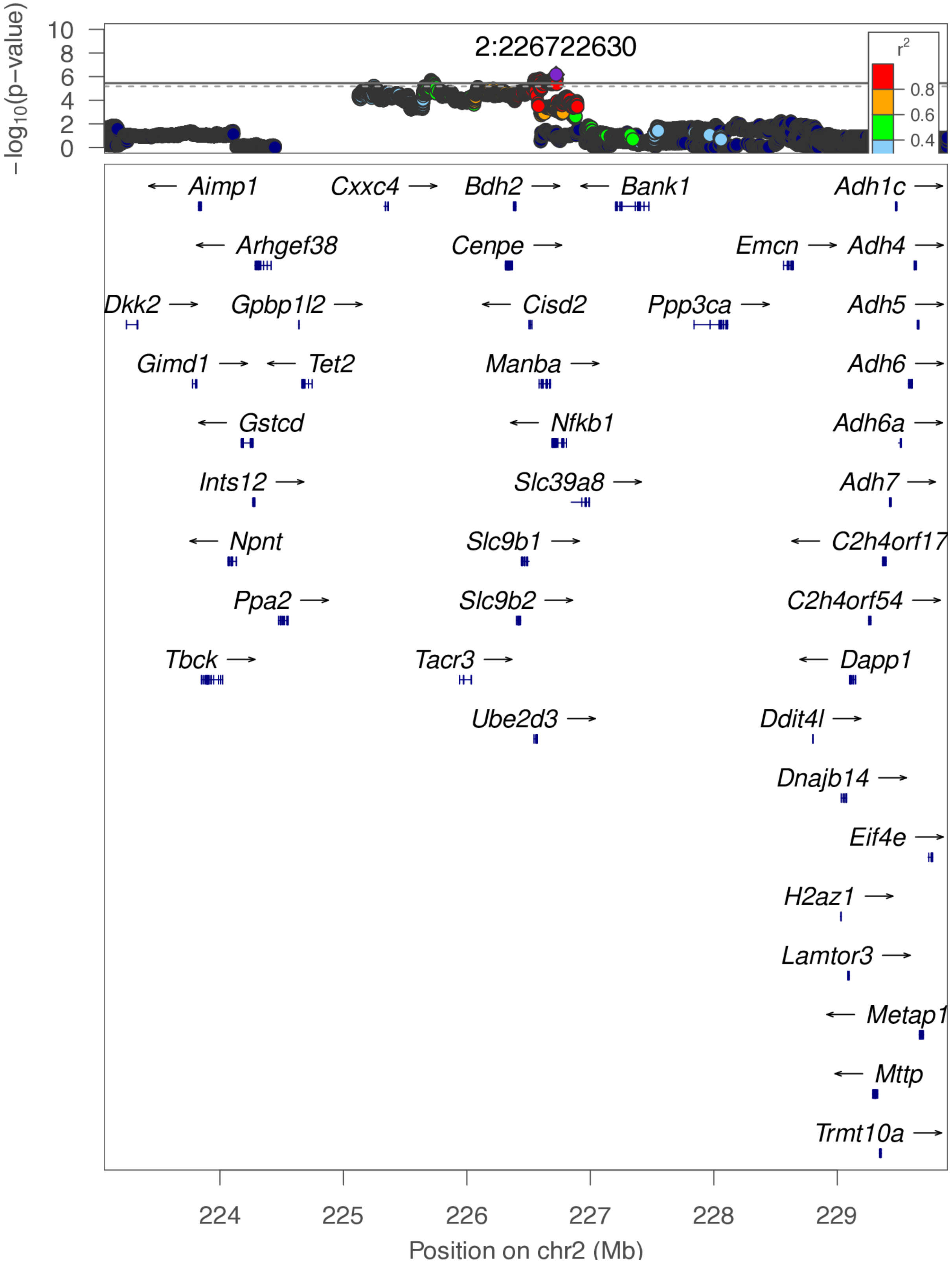

**chr2:235.6 Mb - MEGA - narrow**

Lead: 2:235,932,591 | Genes in QTL interval: 1

QTL interval: chr2:234,722,945-236,162,931

Displayed region: chr2:234,686,261-236,281,993

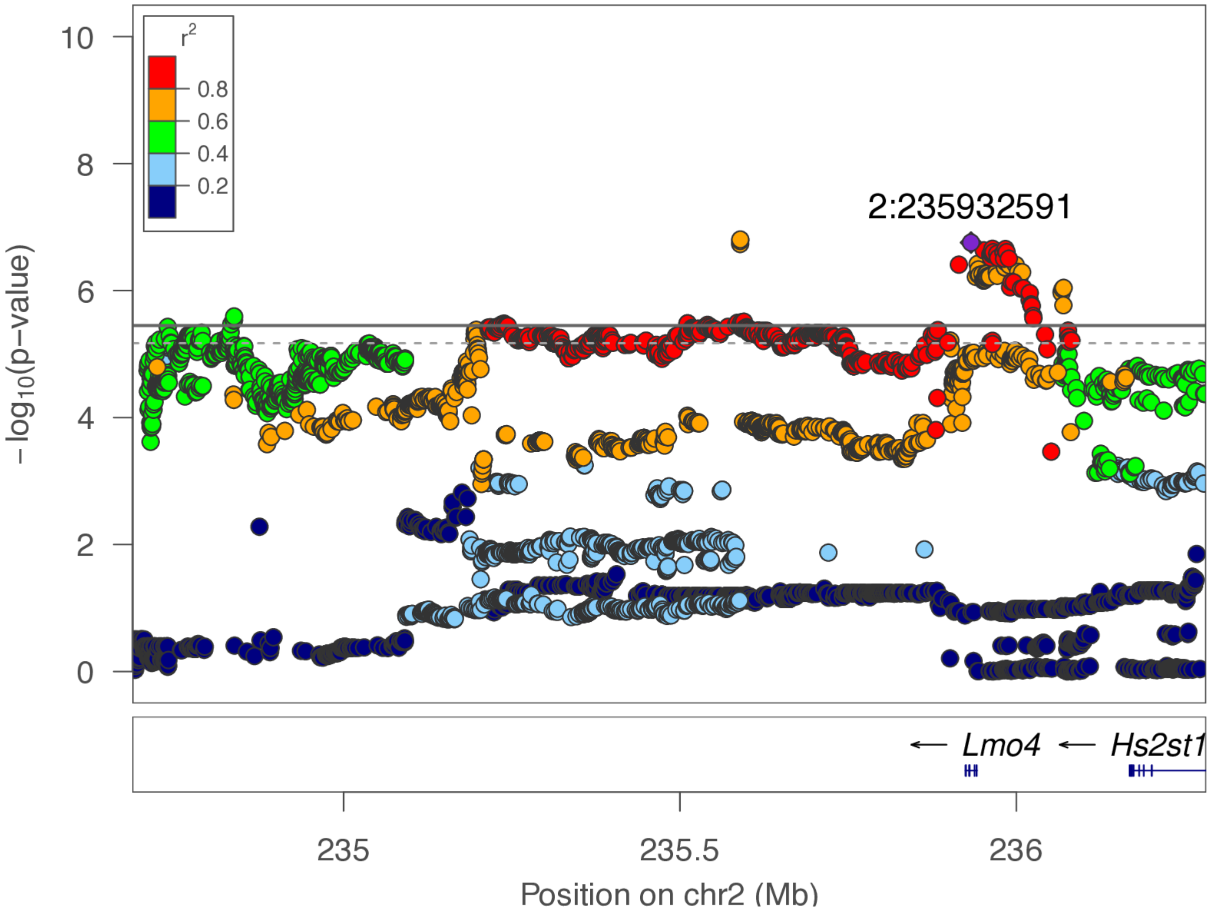

**chr2:235.6 Mb - MEGA - broad**

Lead: 2:235,932,591 | Genes in QTL interval: 1

QTL interval: chr2:234,722,945-236,162,931

Displayed region: chr2:231,886,261-239,081,993

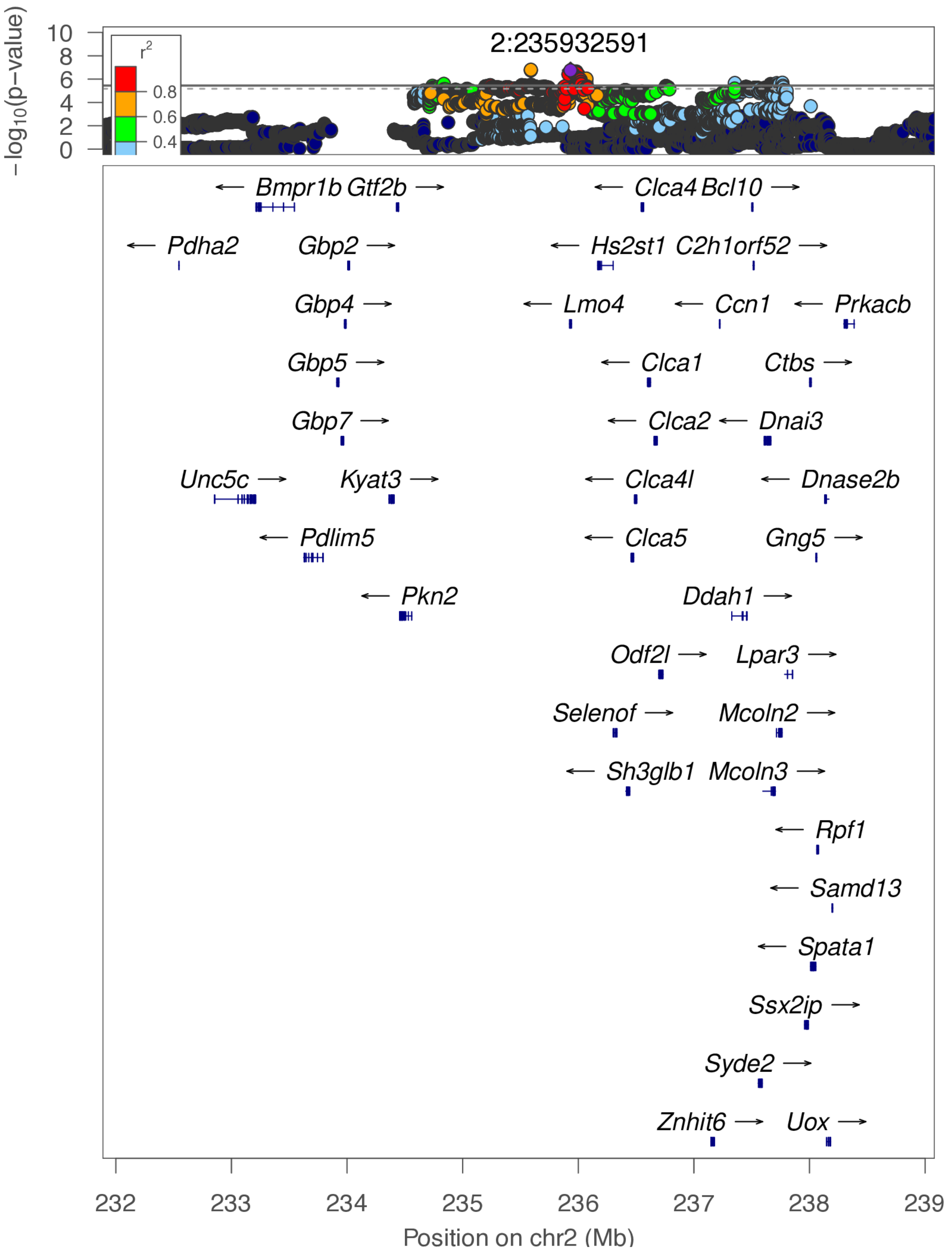

**chr2:235.6 Mb - METAL - narrow**

Lead: 2:235,589,603 | Genes in QTL interval: 1

QTL interval: chr2:234,722,945-236,162,931

Displayed region: chr2:234,508,734-236,665,792

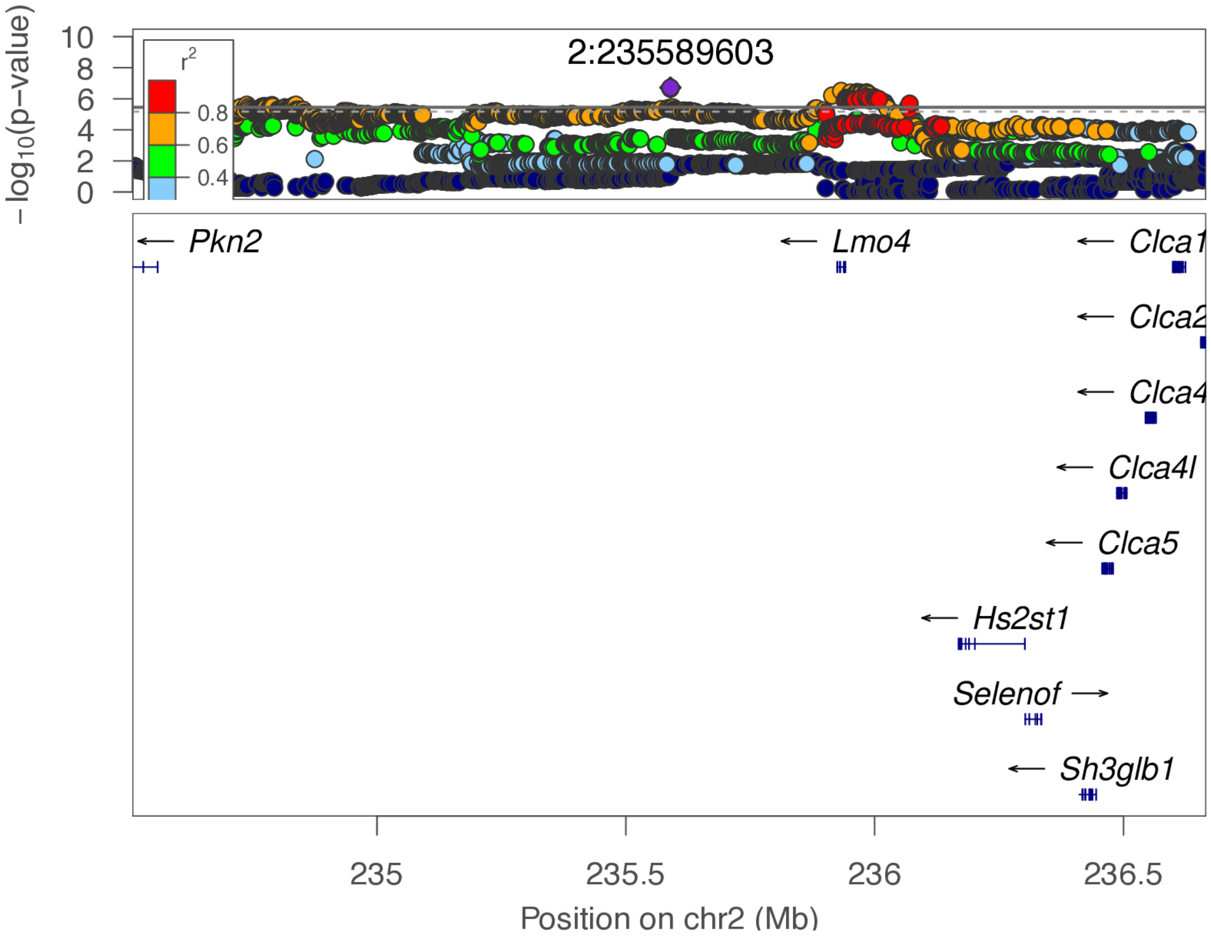

**chr2:235.6 Mb - METAL - broad**

Lead: 2:235,589,603 | Genes in QTL interval: 1

QTL interval: chr2:234,722,945-236,162,931

Displayed region: chr2:231,708,734-239,465,792

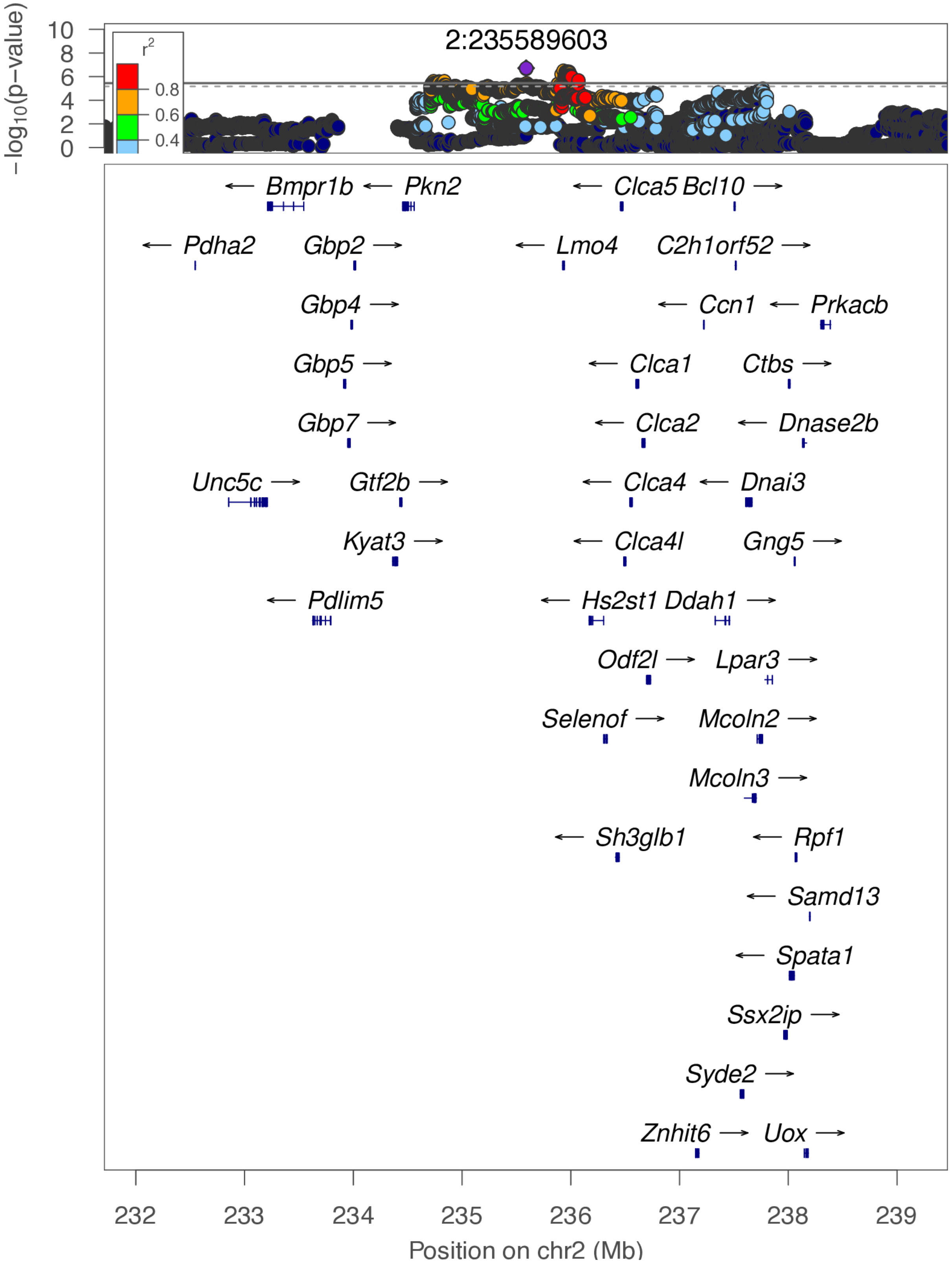

**chr2:235.6 Mb - Genomic SEM - narrow**

Lead: 2:235,589,603 | Genes in QTL interval: 1

QTL interval: chr2:234,722,945-236,162,931

Displayed region: chr2:234,508,734-236,665,792

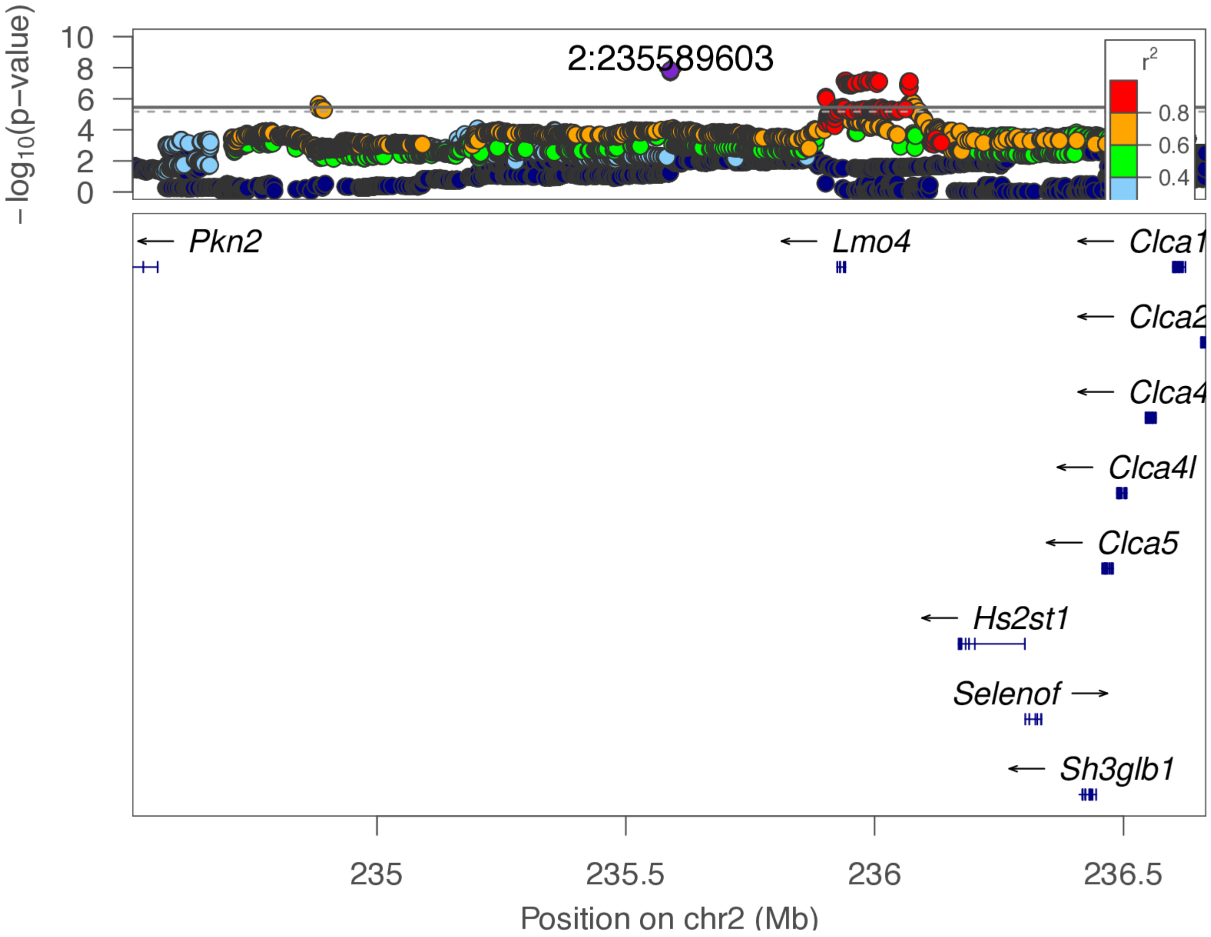

**chr2:235.6 Mb - Genomic SEM - broad**

Lead: 2:235,589,603 | Genes in QTL interval: 1

QTL interval: chr2:234,722,945-236,162,931

Displayed region: chr2:231,708,734-239,465,792

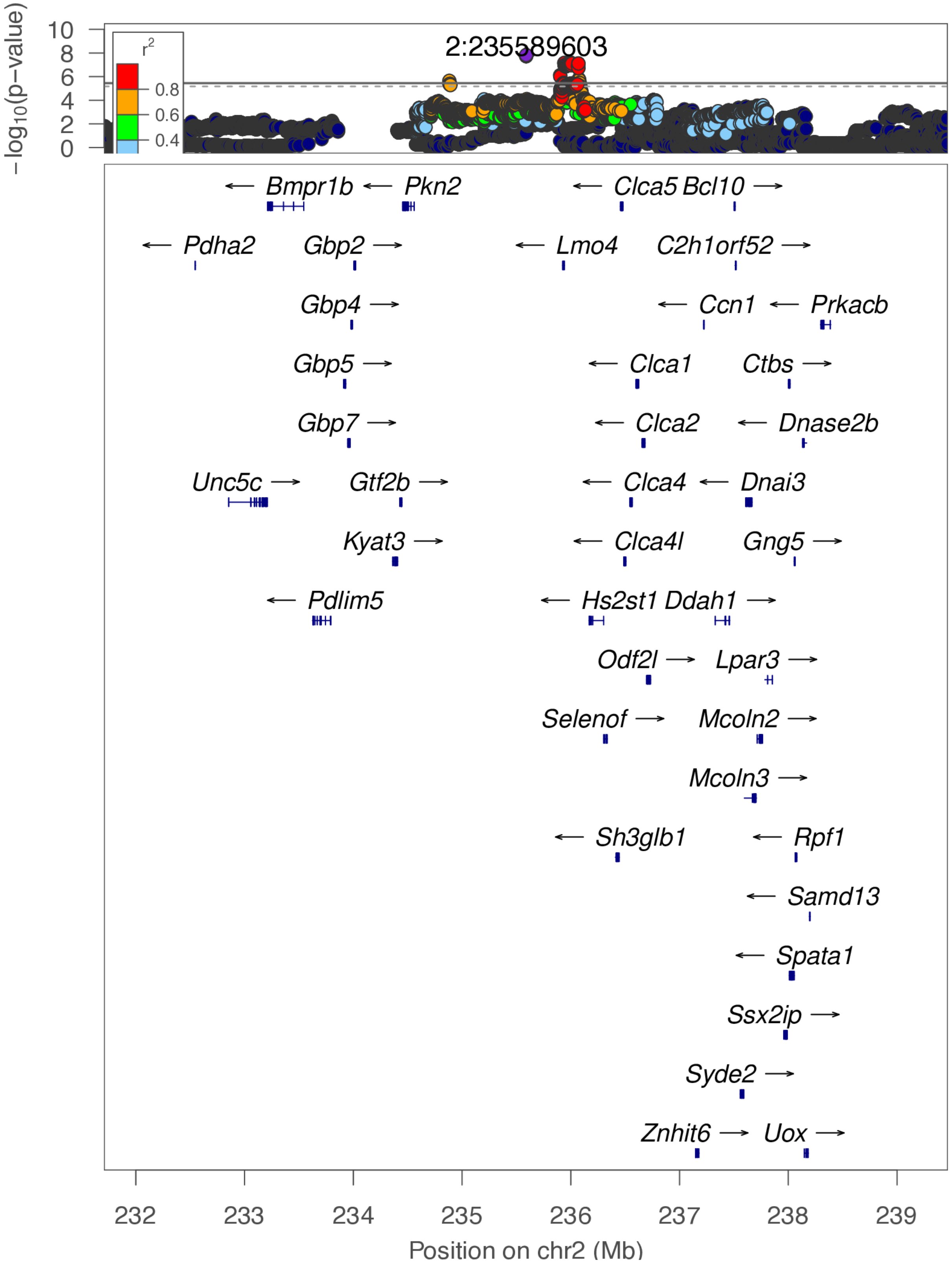

**chr3:59.4 Mb - MEGA - narrow**

Lead: 3:59,395,491 | Genes in QTL interval: 2

QTL interval: chr3:59,313,308-60,365,607

Displayed region: chr3:58,472,382-61,823,014

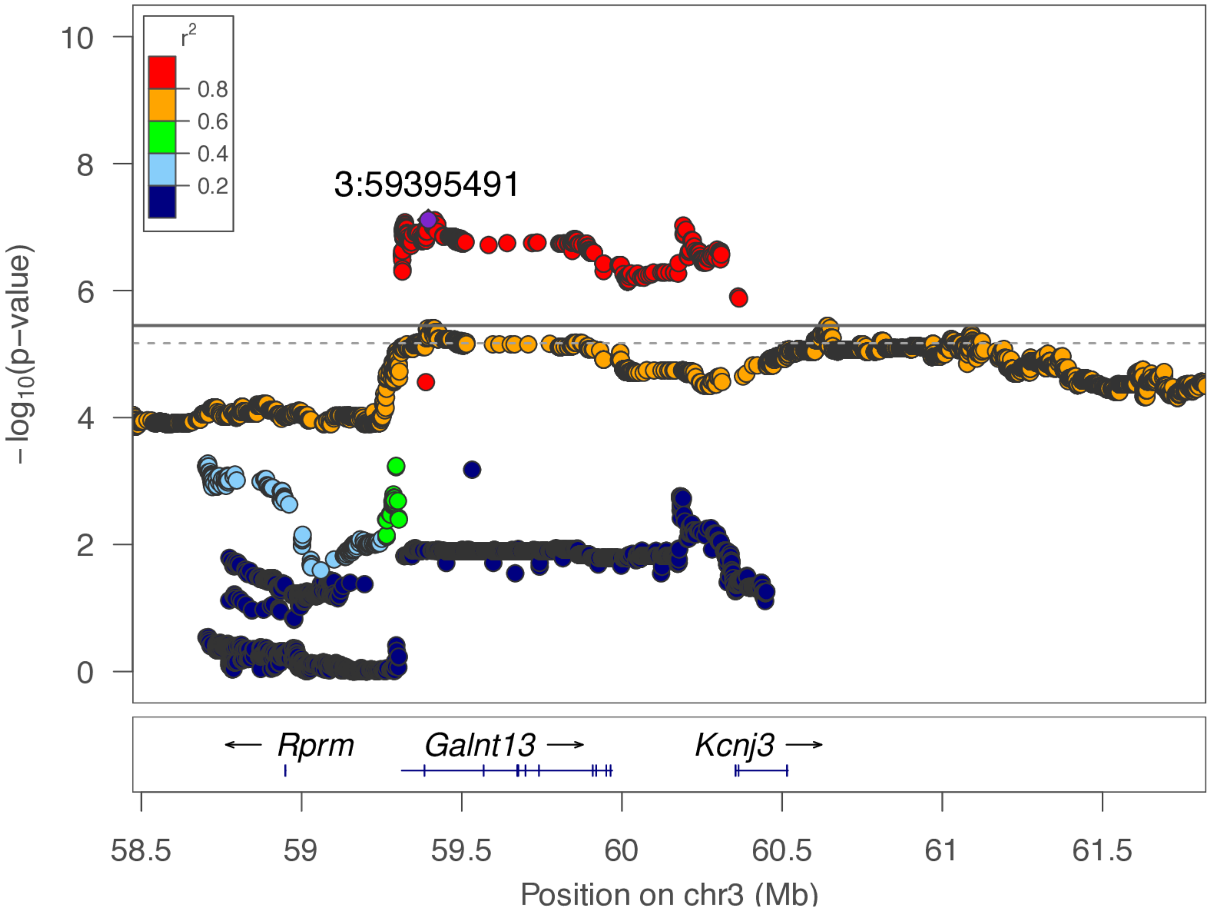

**chr3:59.4 Mb - MEGA - broad**

Lead: 3:59,395,491 | Genes in QTL interval: 2

QTL interval: chr3:59,313,308-60,365,607

Displayed region: chr3:55,672,382-64,623,014

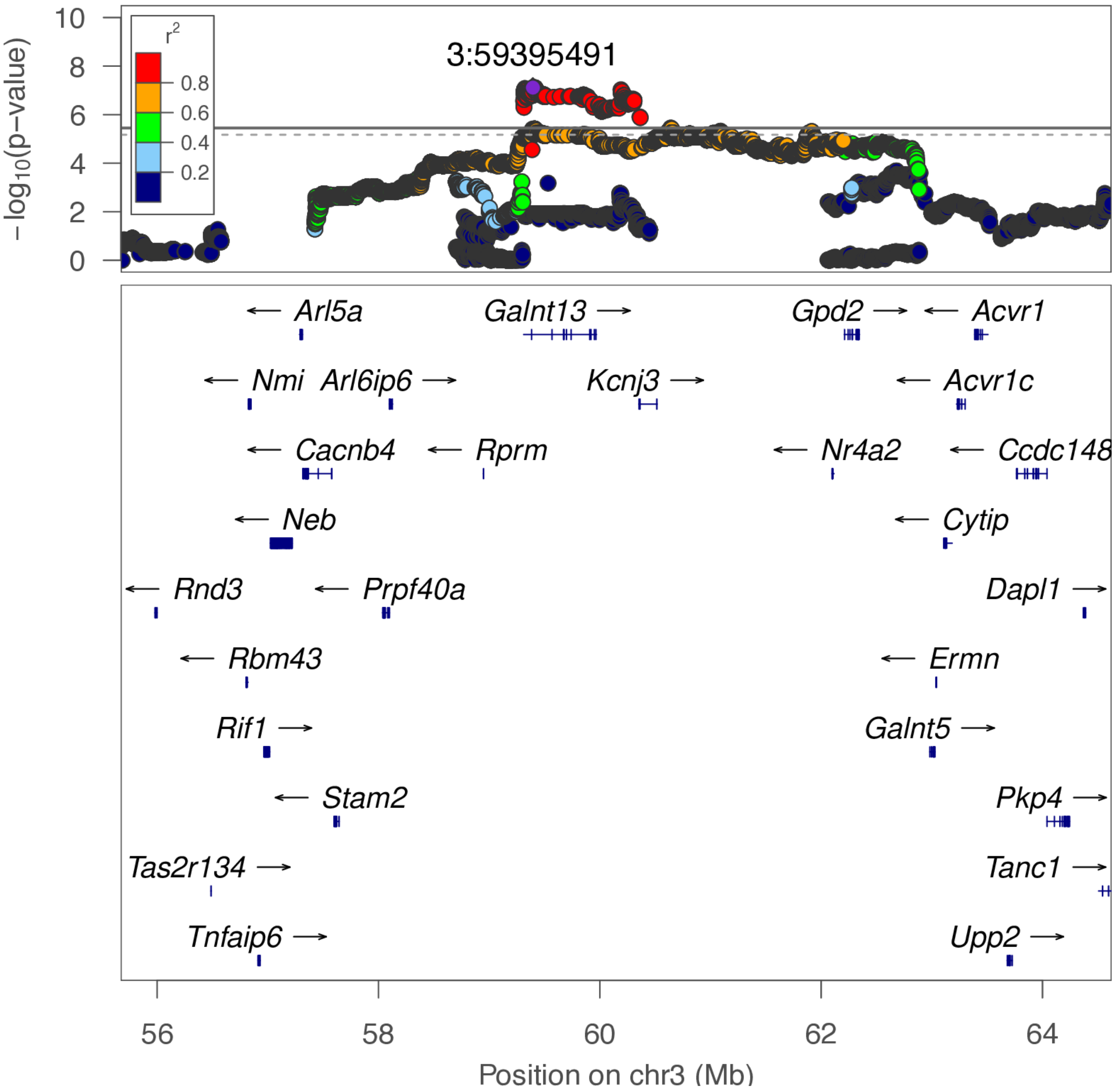

**chr3:59.4 Mb - METAL - narrow**

Lead: 3:59,414,346 | Genes in QTL interval: 2

QTL interval: chr3:59,313,308-60,365,607

Displayed region: chr3:58,632,183-61,931,496

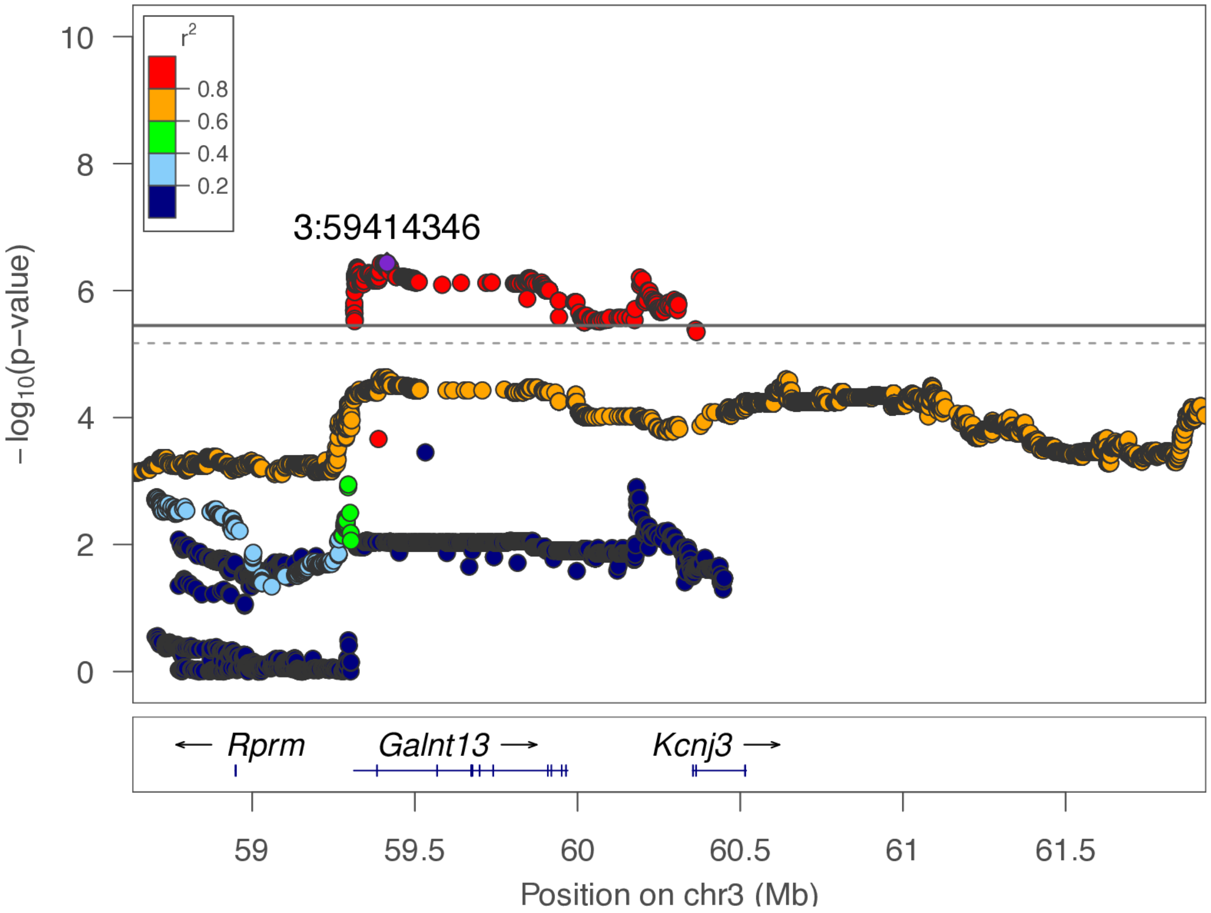

**chr3:59.4 Mb - METAL - broad**

Lead: 3:59,414,346 | Genes in QTL interval: 2

QTL interval: chr3:59,313,308-60,365,607

Displayed region: chr3:55,832,183-64,731,496

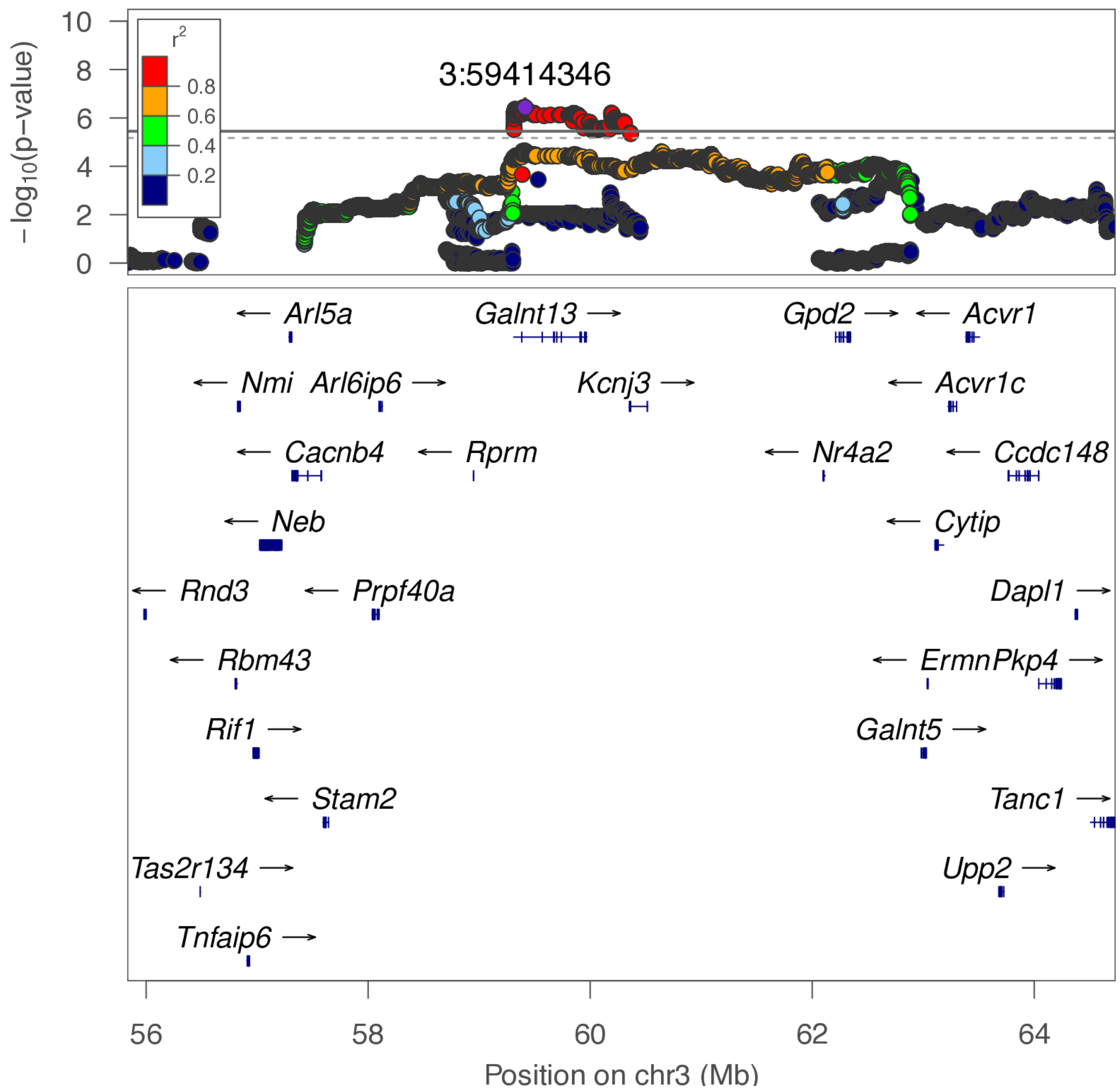

**chr4:33.6 Mb - MEGA - narrow**

Lead: 4:33,666,294 | Genes in QTL interval: 3

QTL interval: chr4:33,427,113-33,822,758

Displayed region: chr4:32,747,715-34,022,758

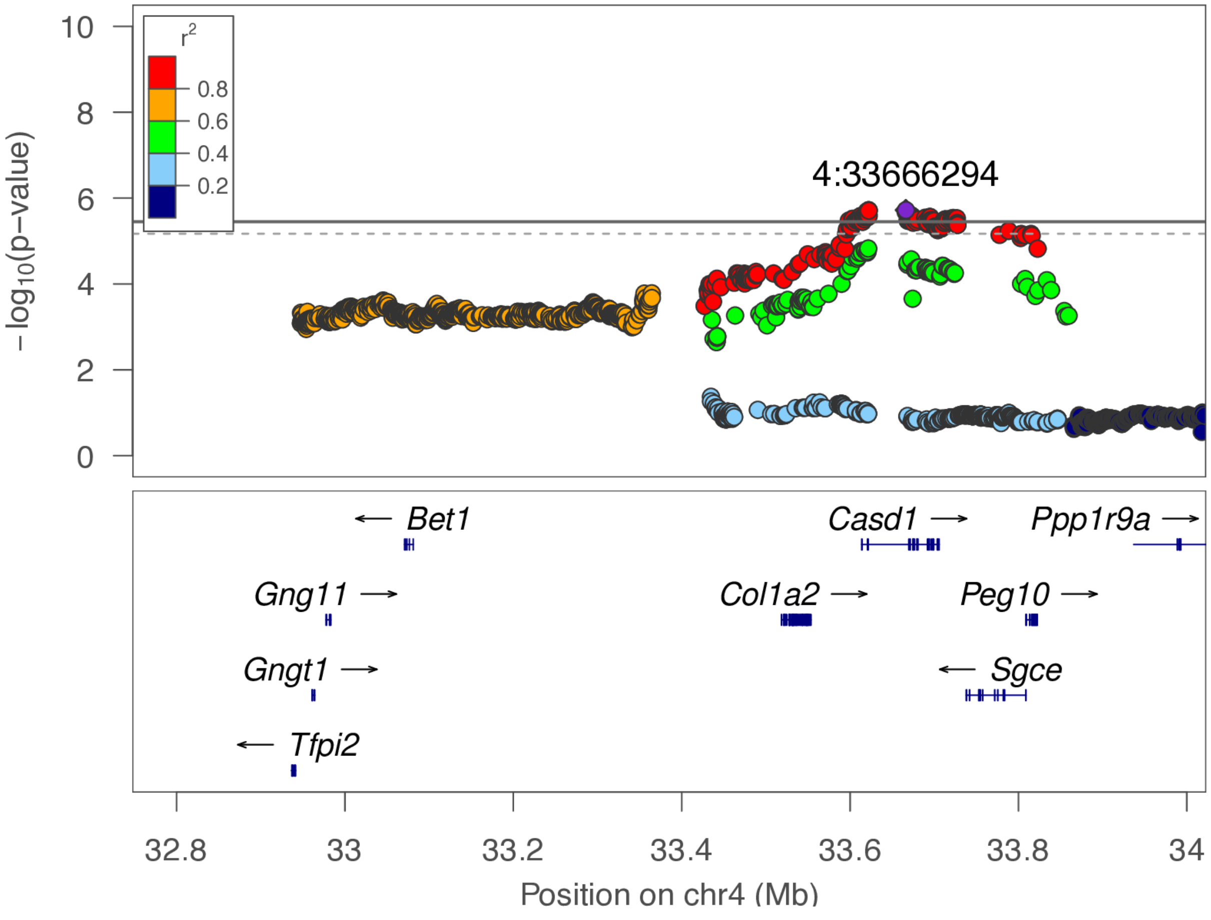

**chr4:33.6 Mb - MEGA - broad**

Lead: 4:33,666,294 | Genes in QTL interval: 3

QTL interval: chr4:33,427,113-33,822,758

Displayed region: chr4:29,947,715-36,822,758

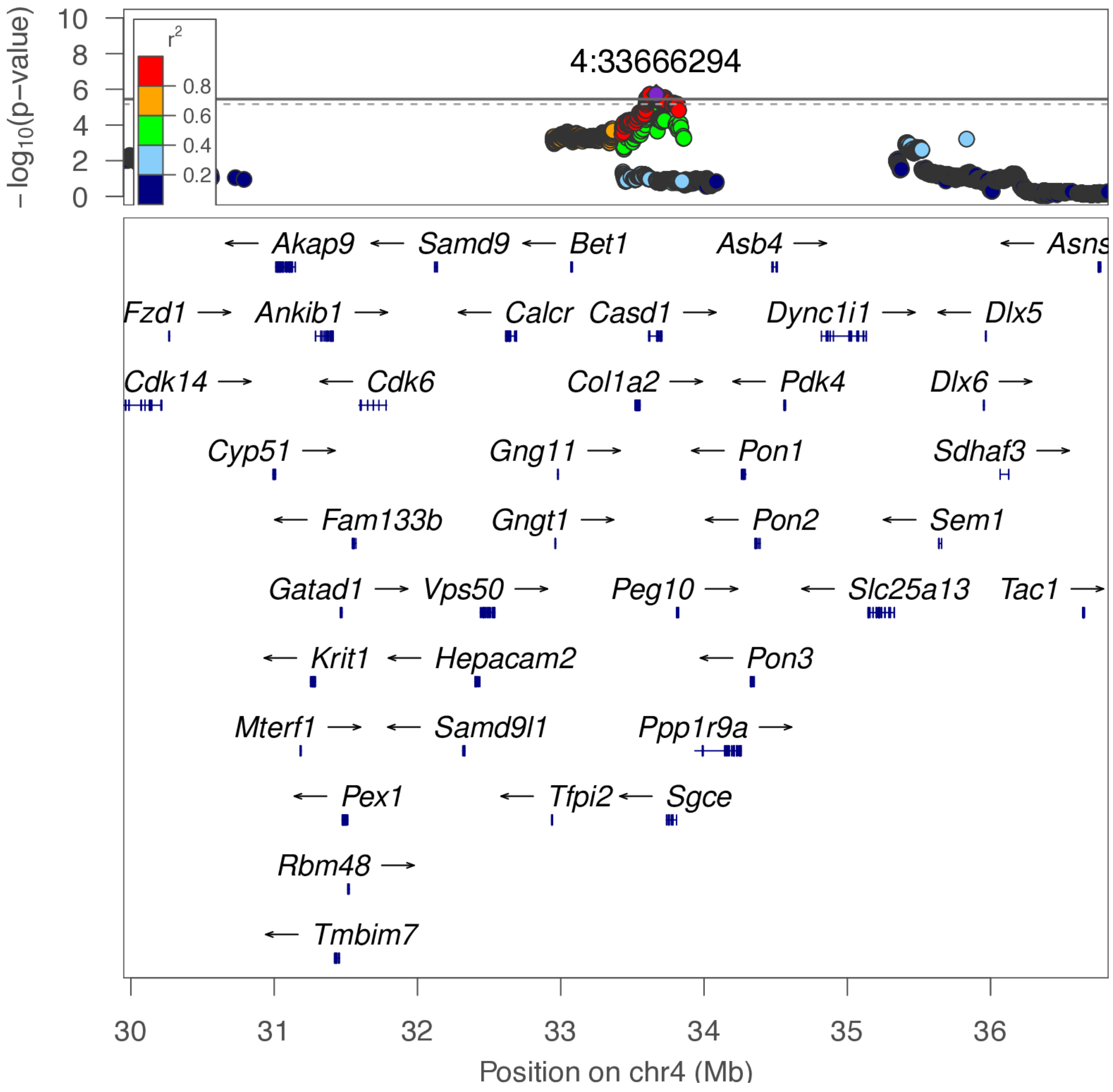

**chr4:33.6 Mb - METAL - narrow**

Lead: 4:33,622,055 | Genes in QTL interval: 3

QTL interval: chr4:33,427,113-33,822,758

Displayed region: chr4:32,747,715-34,022,758

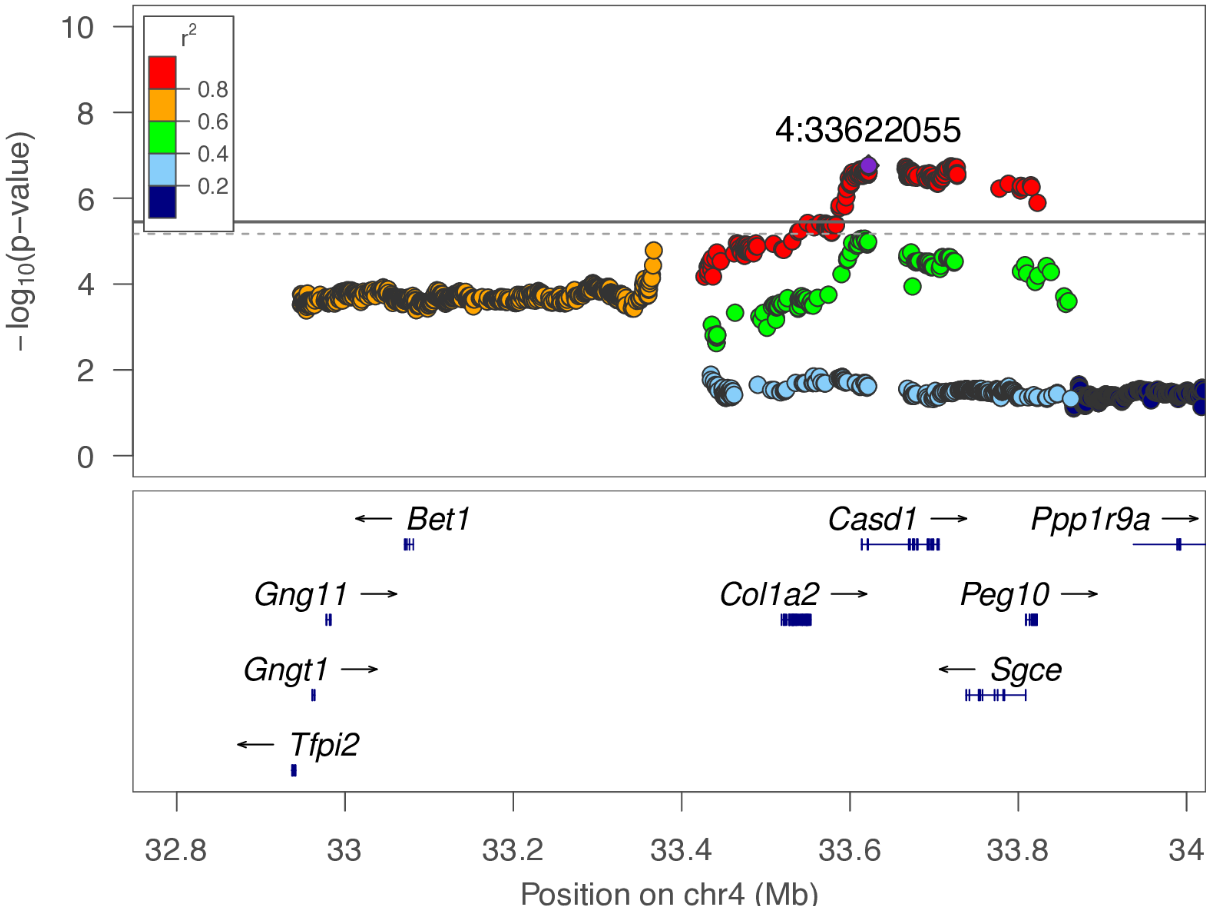

**chr4:33.6 Mb - METAL - broad**

Lead: 4:33,622,055 | Genes in QTL interval: 3

QTL interval: chr4:33,427,113-33,822,758

Displayed region: chr4:29,947,715-36,822,758

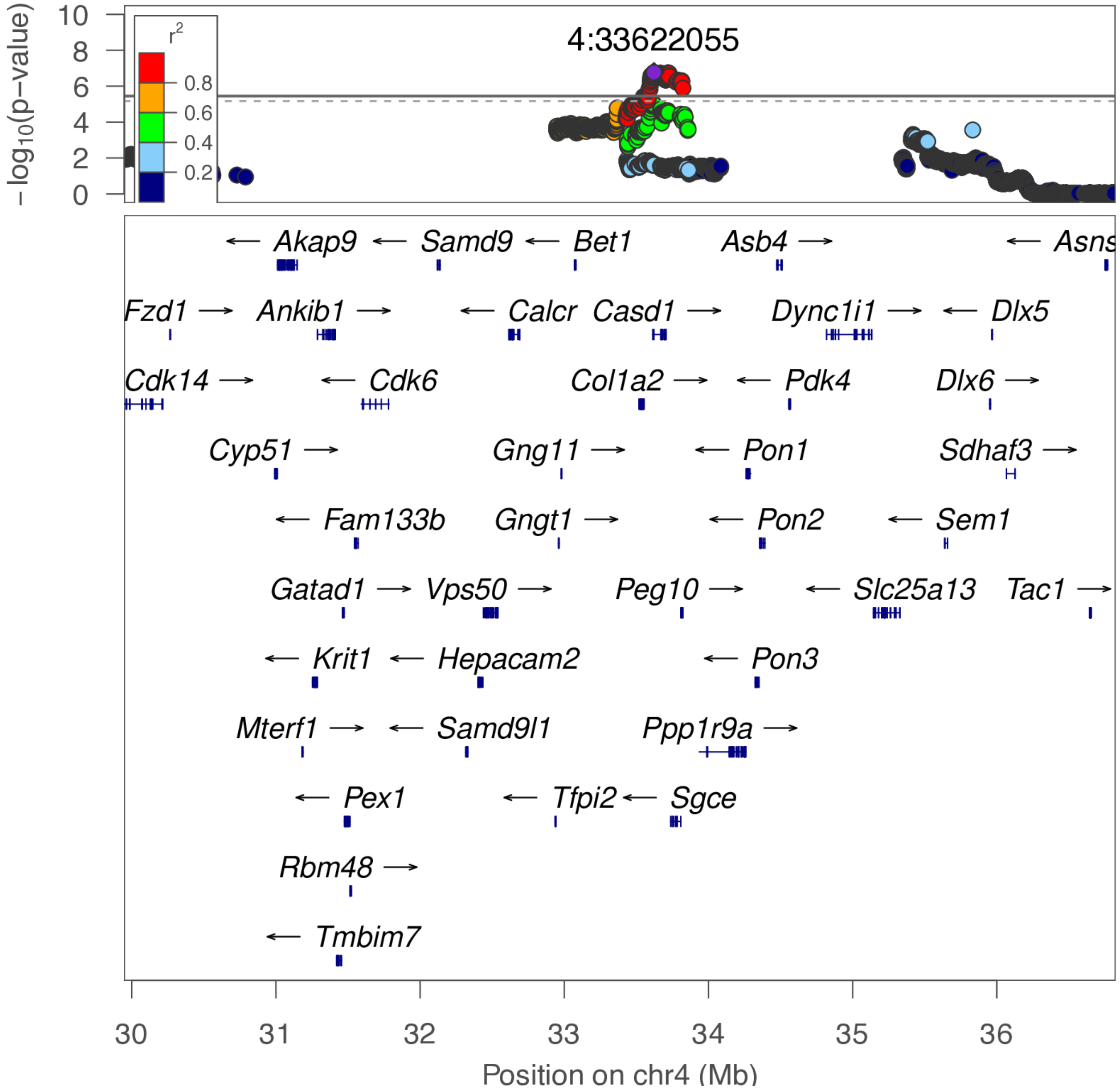

**chr4:86.4 Mb - Genomic SEM - narrow**

Lead: 4:86,372,848 | Genes in QTL interval: 1

QTL interval: chr4:86,328,860-86,497,748

Displayed region: chr4:86,036,535-88,930,367

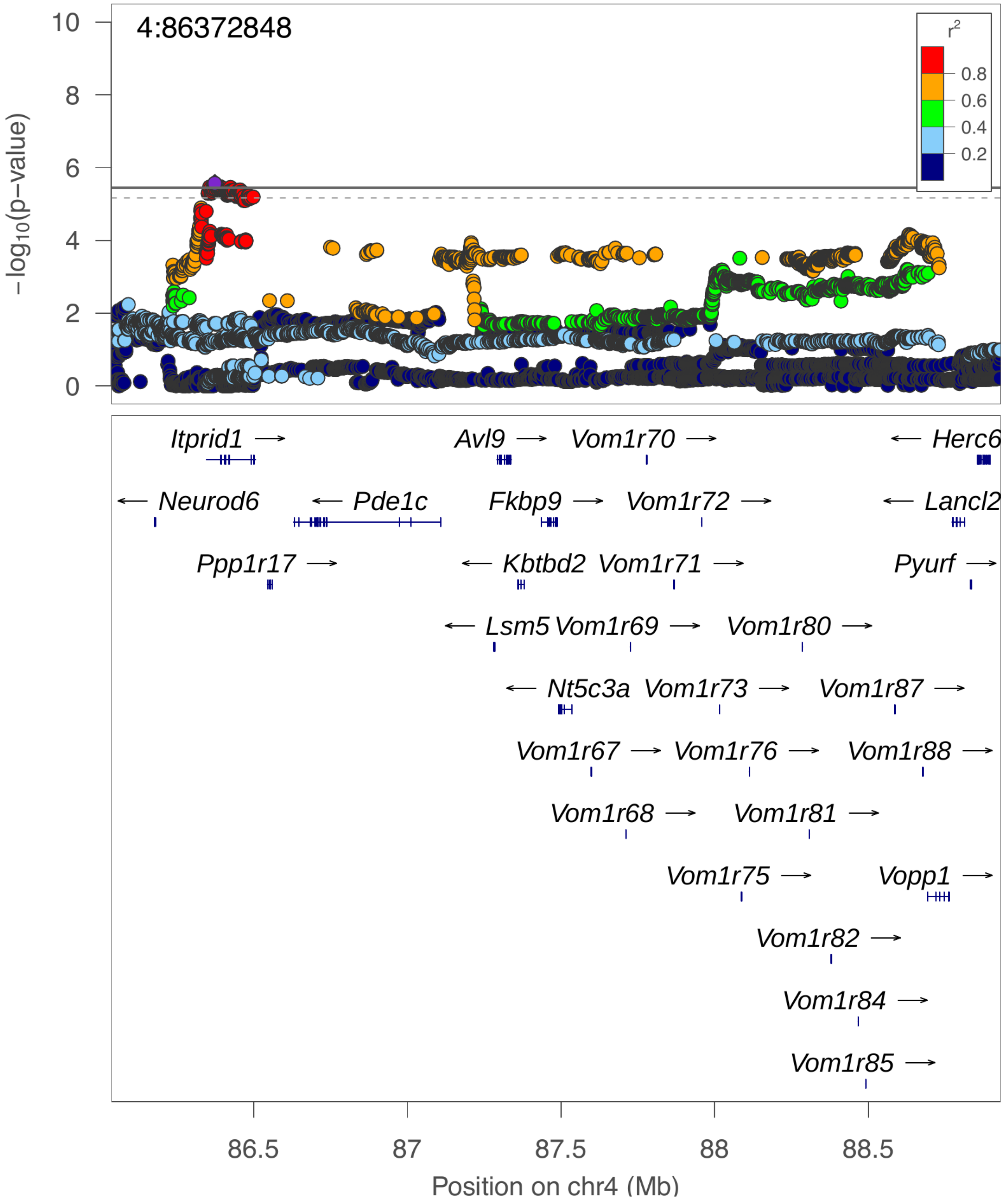

**chr4:86.4 Mb - Genomic SEM - broad**

Lead: 4:86,372,848 | Genes in QTL interval: 1

QTL interval: chr4:86,328,860-86,497,748

Displayed region: chr4:83,236,535-91,730,367

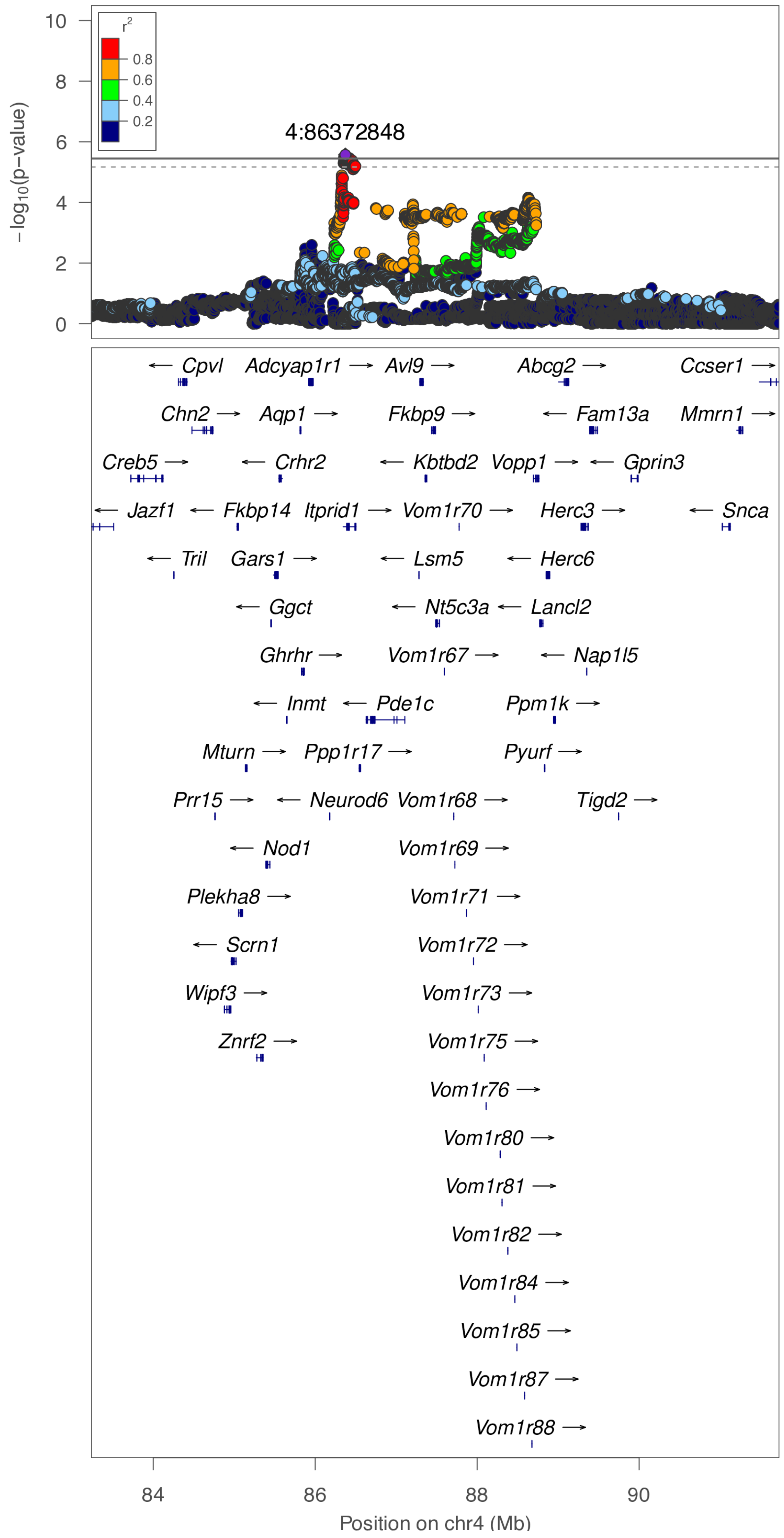

**chr5:40.8 Mb - Genomic SEM - narrow**

Lead: 5:40,848,632 | Genes in QTL interval: 2

QTL interval: chr5:40,753,574-41,614,105

Displayed region: chr5:40,553,574-41,823,683

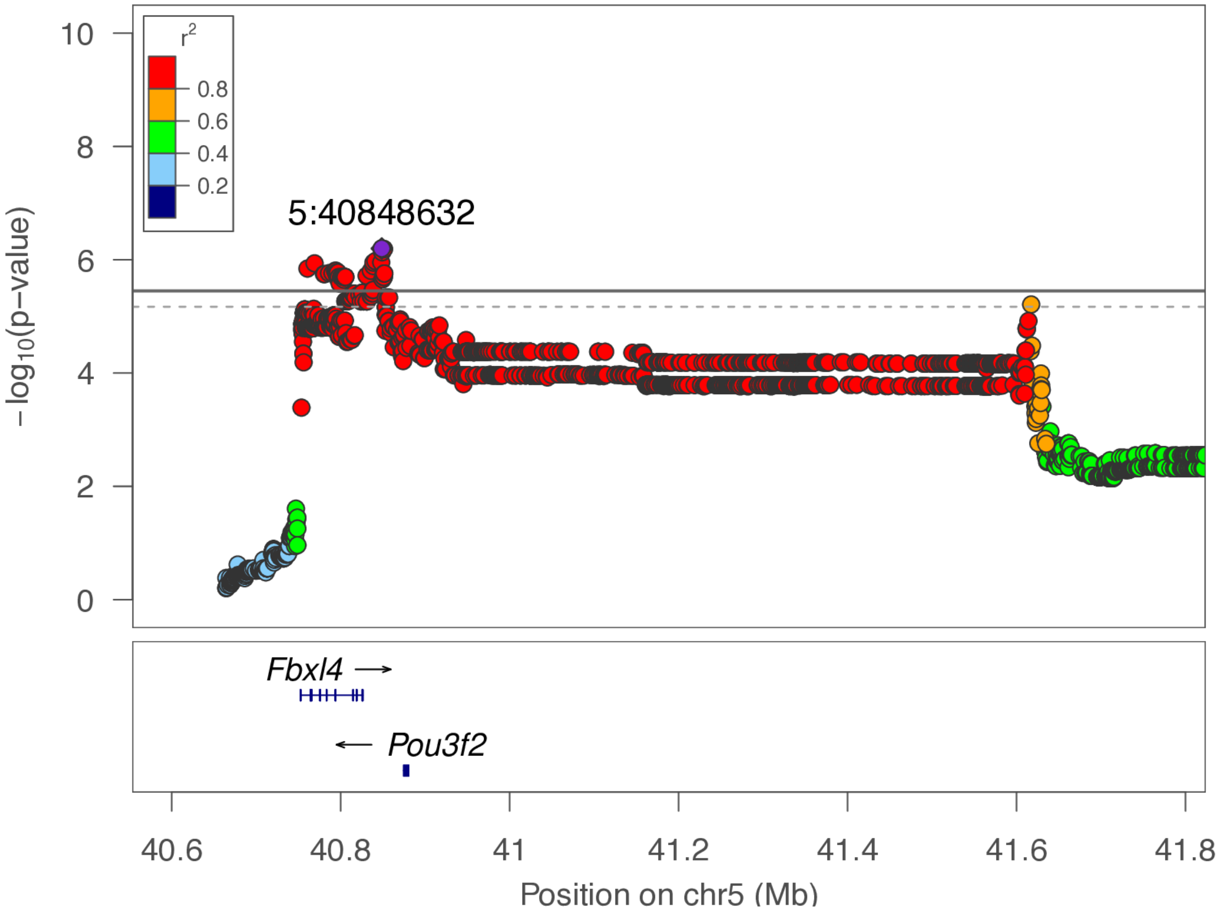

**chr5:40.8 Mb - Genomic SEM - broad**

Lead: 5:40,848,632 | Genes in QTL interval: 2

QTL interval: chr5:40,753,574-41,614,105

Displayed region: chr5:37,753,574-44,623,683

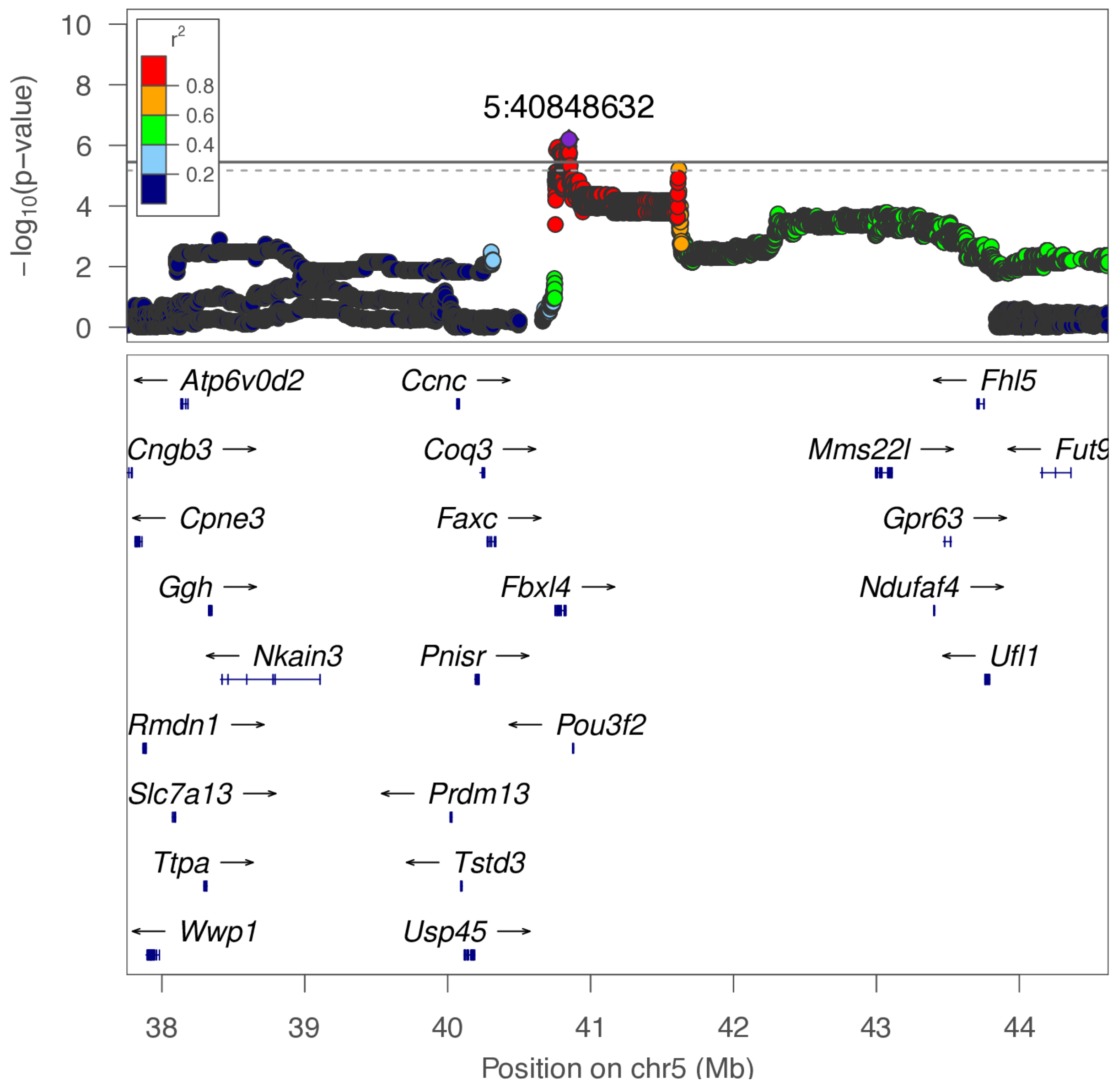

**chr5:74.8 Mb - Genomic SEM - narrow**

Lead: 5:74,835,274 | Genes in QTL interval: 2

QTL interval: chr5:74,820,249-75,275,913

Displayed region: chr5:74,565,509-75,878,337

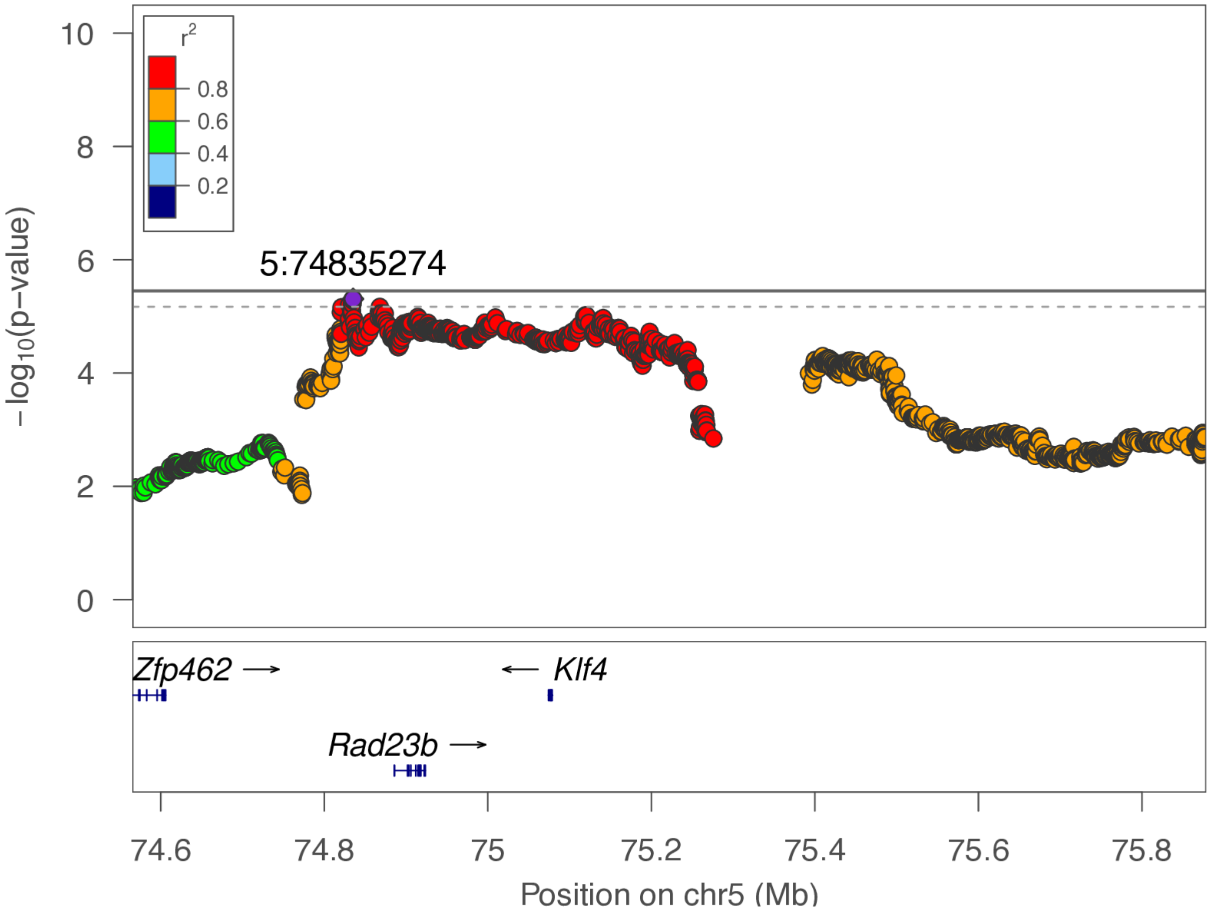

**chr5:74.8 Mb - Genomic SEM - broad**

Lead: 5:74,835,274 | Genes in QTL interval: 2

QTL interval: chr5:74,820,249-75,275,913

Displayed region: chr5:71,765,509-78,678,337

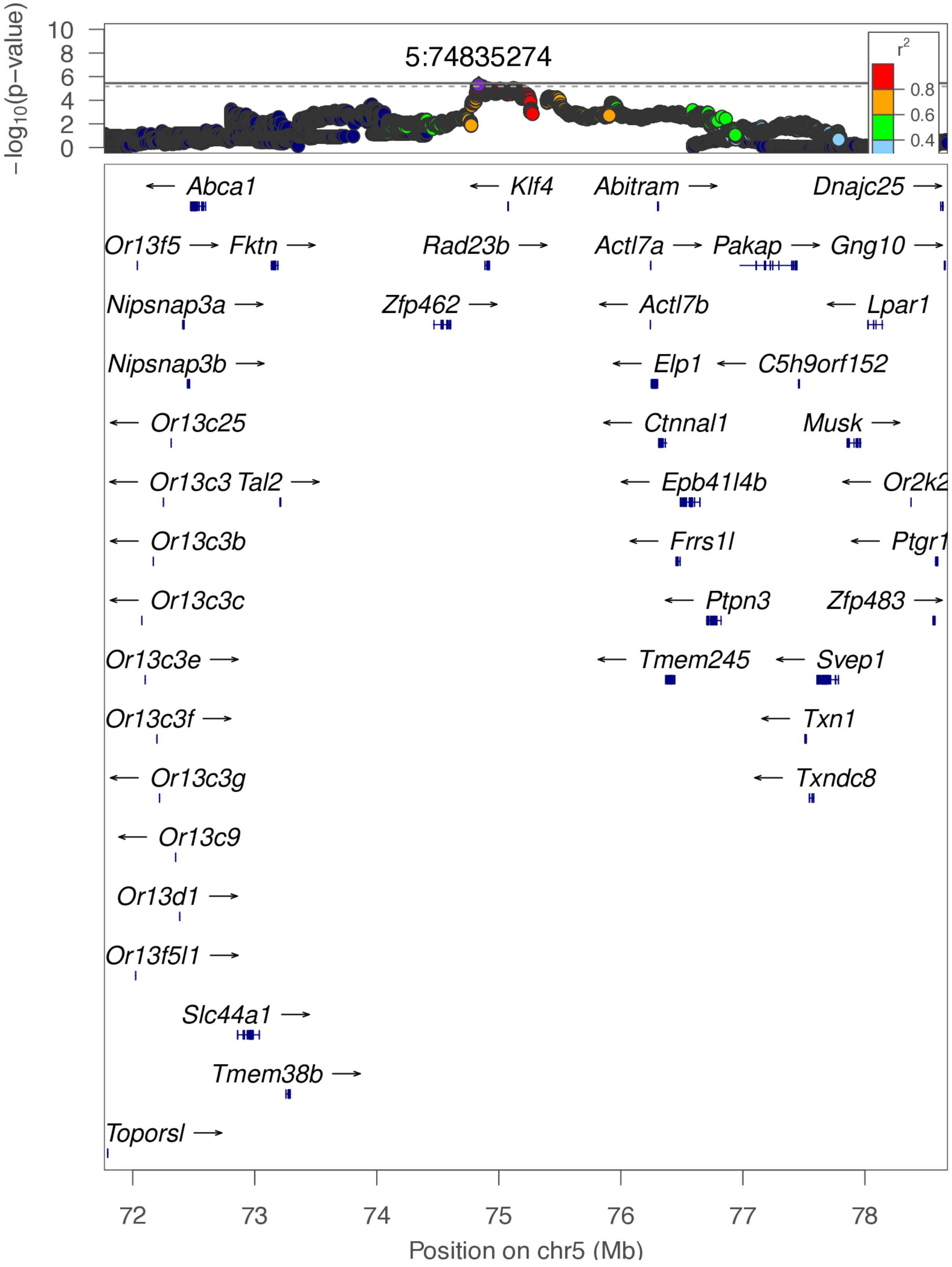

**chr6:50.3 Mb - Genomic SEM - narrow**

Lead: 6:50,315,544 | Genes in QTL interval: 0

QTL interval: chr6:50,209,832-50,416,657

Displayed region: chr6:49,859,630-50,616,657

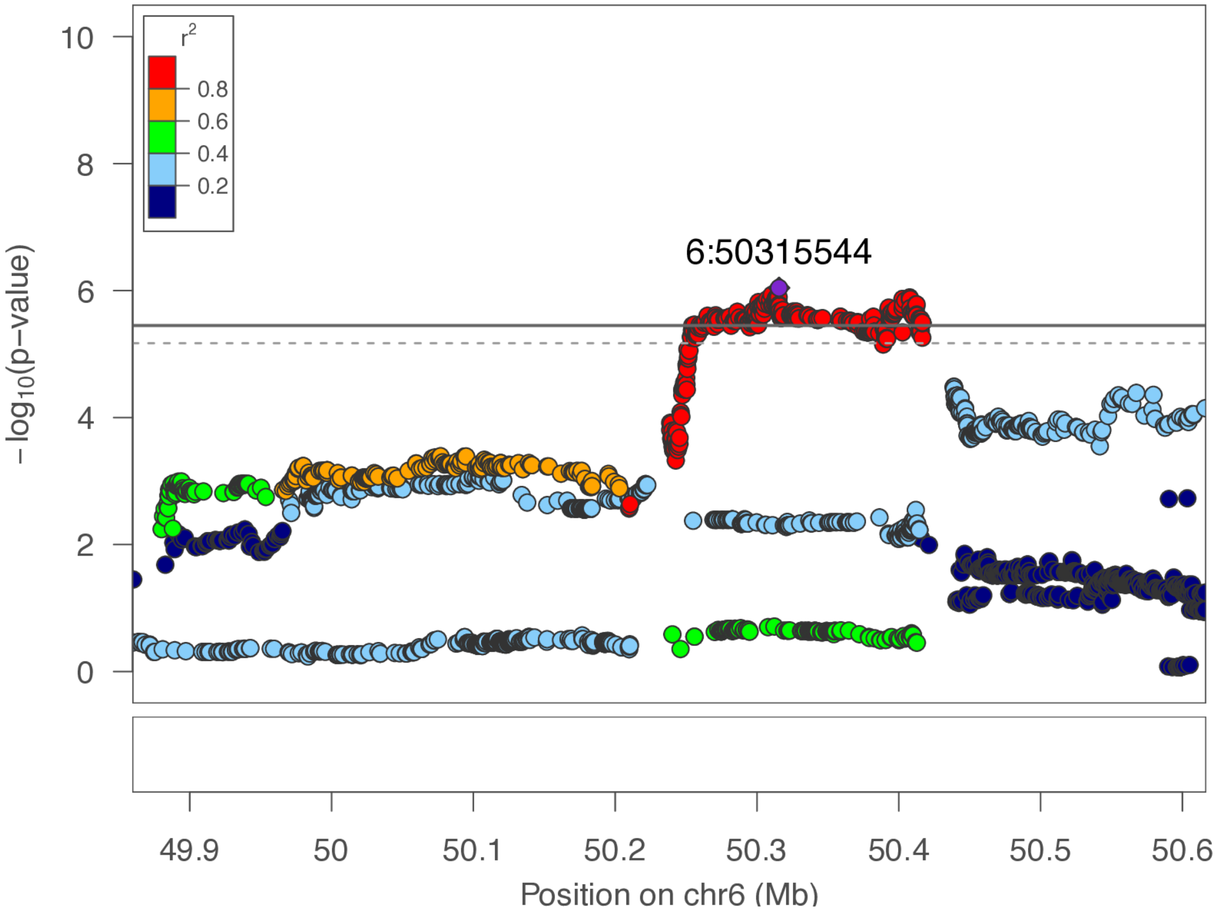

**chr6:50.3 Mb - Genomic SEM - broad**

Lead: 6:50,315,544 | Genes in QTL interval: 0

QTL interval: chr6:50,209,832-50,416,657

Displayed region: chr6:47,059,630-53,416,657

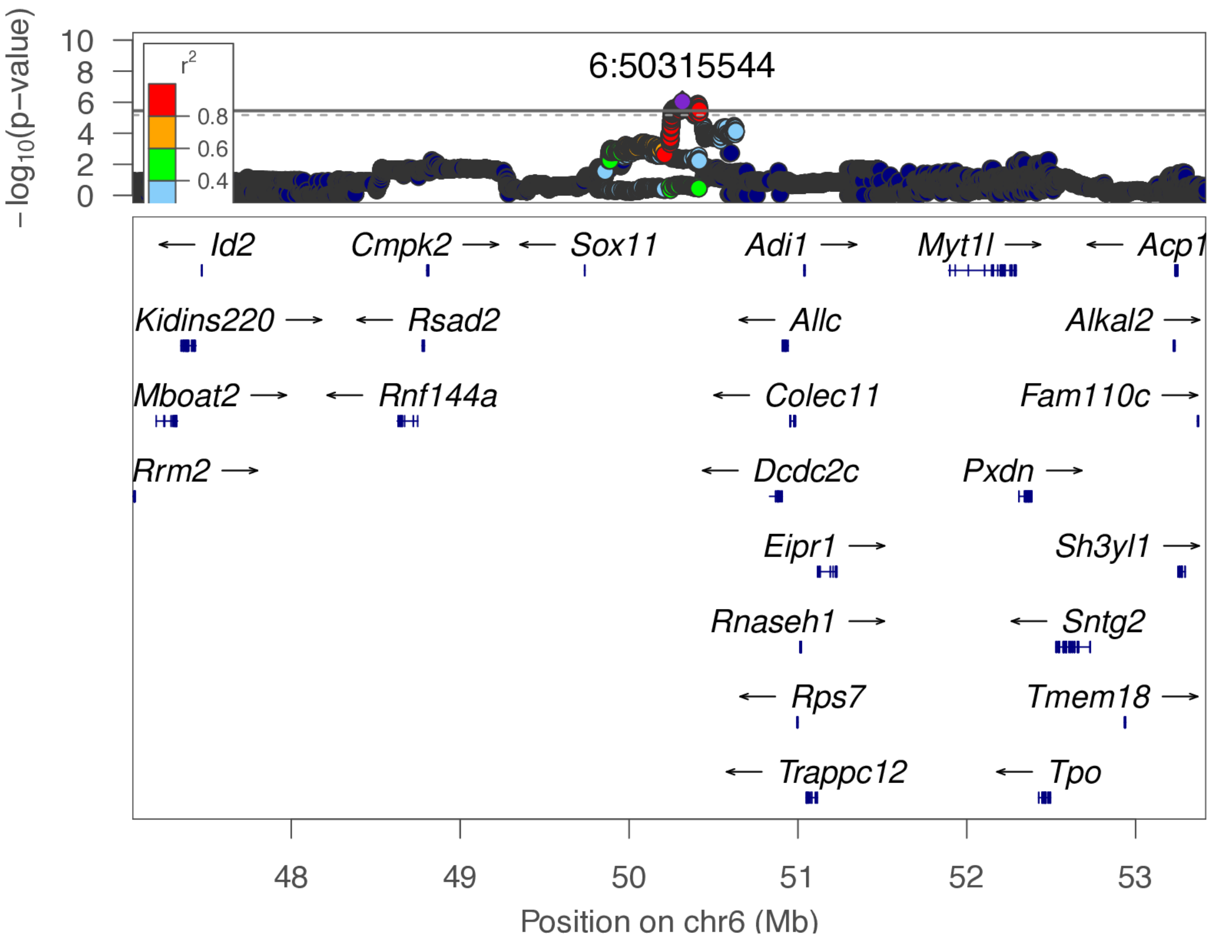

**chr7:52.1 Mb - METAL - narrow**

Lead: 7:52,113,752 | Genes in QTL interval: 12

QTL interval: chr7:51,631,833-53,558,662

Displayed region: chr7:51,428,146-54,128,396

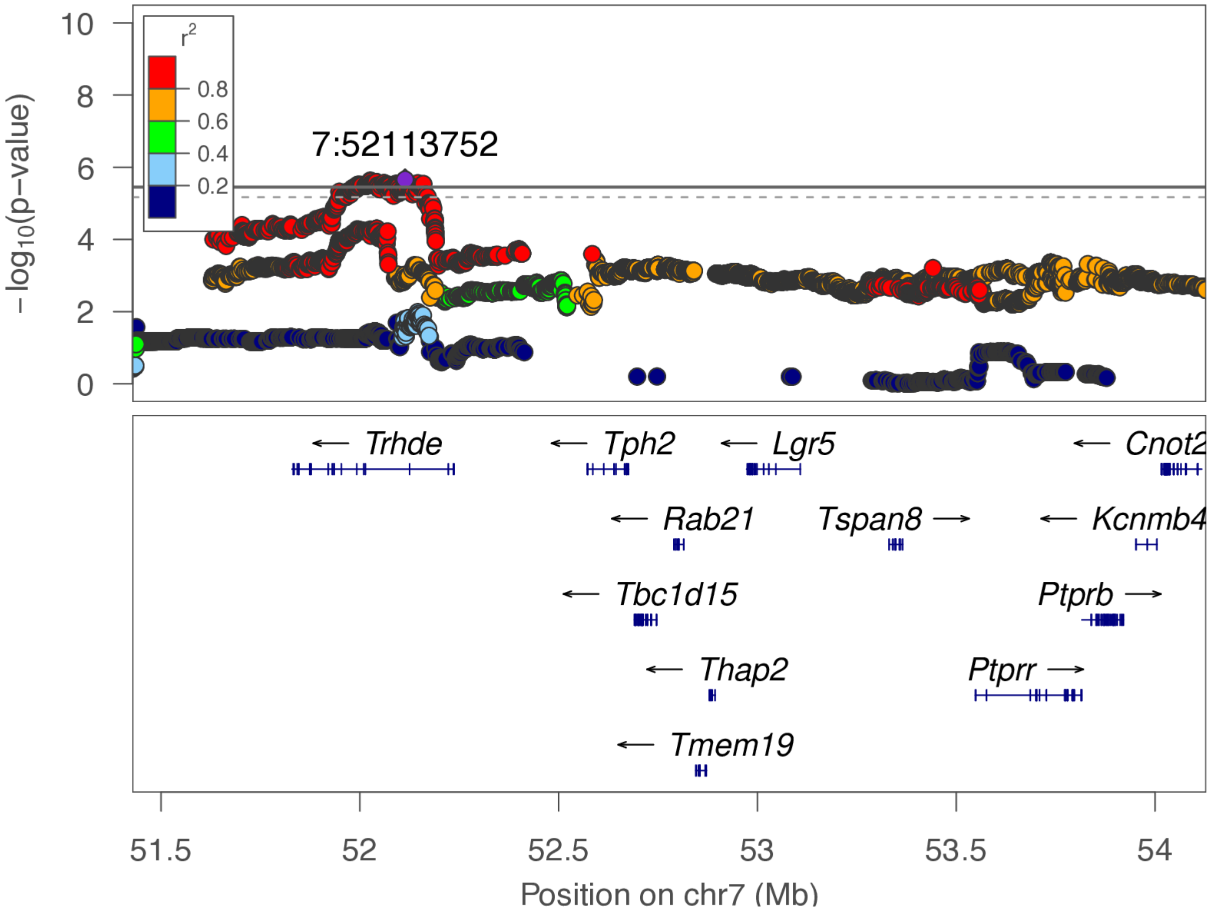

**chr7:52.1 Mb - METAL - broad**

Lead: 7:52,113,752 | Genes in QTL interval: 12

QTL interval: chr7:51,631,833-53,558,662

Displayed region: chr7:48,628,146-56,928,396

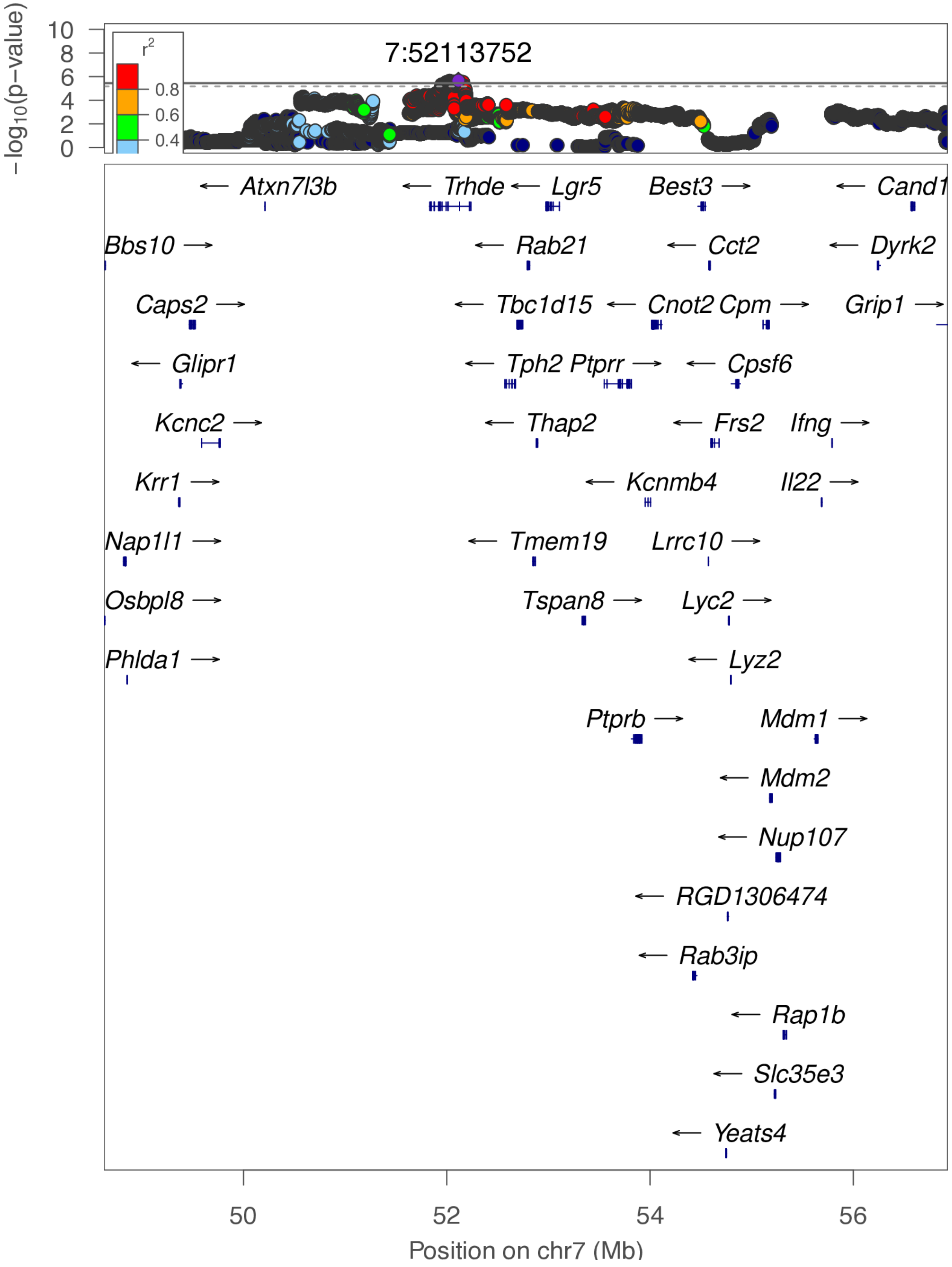

**chr7:103.6 Mb - Genomic SEM - narrow**

Lead: 7:103,612,226 | Genes in QTL interval: 1

QTL interval: chr7:102,209,016-103,668,981

Displayed region: chr7:102,009,016-103,868,981

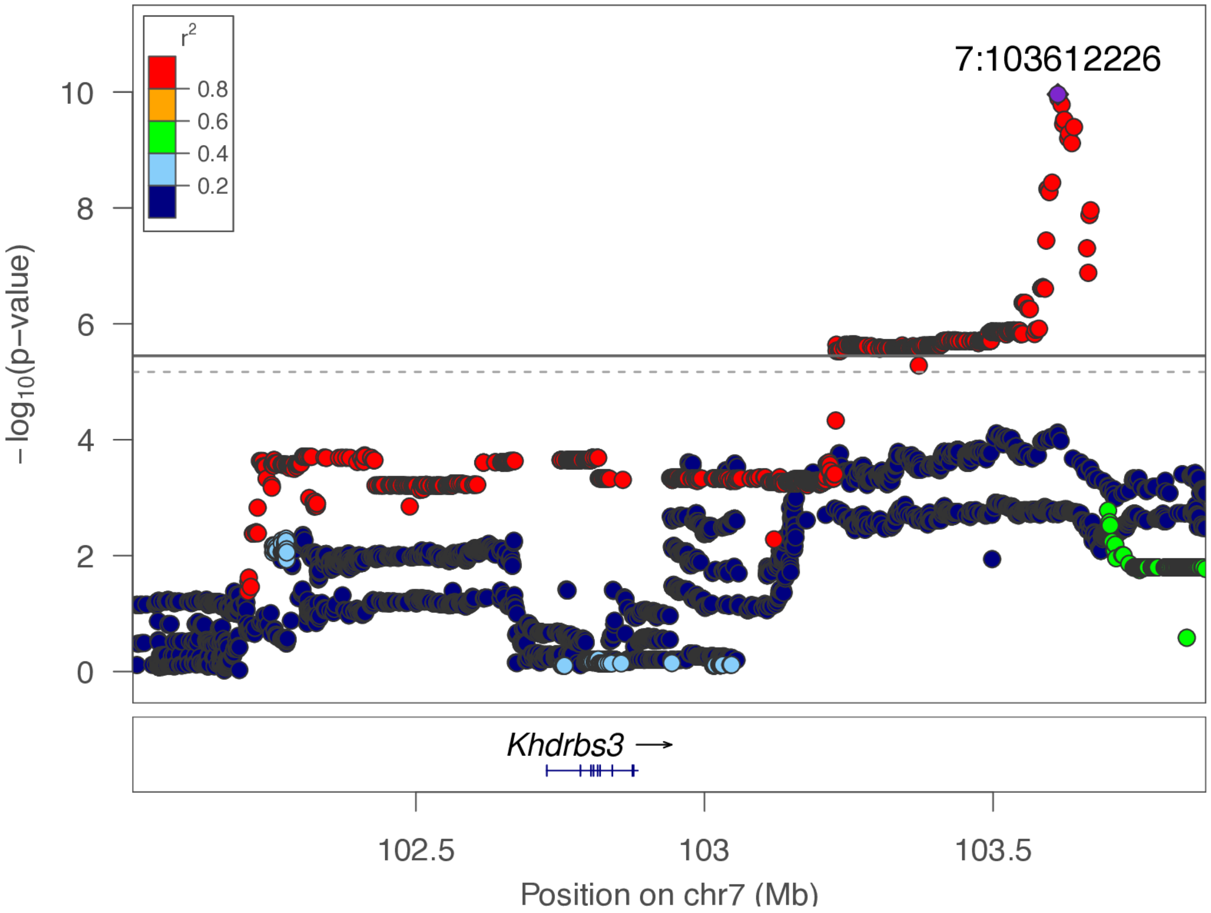

**chr7:103.6 Mb - Genomic SEM - broad**

Lead: 7:103,612,226 | Genes in QTL interval: 1

QTL interval: chr7:102,209,016-103,668,981

Displayed region: chr7:99,209,016-106,668,981

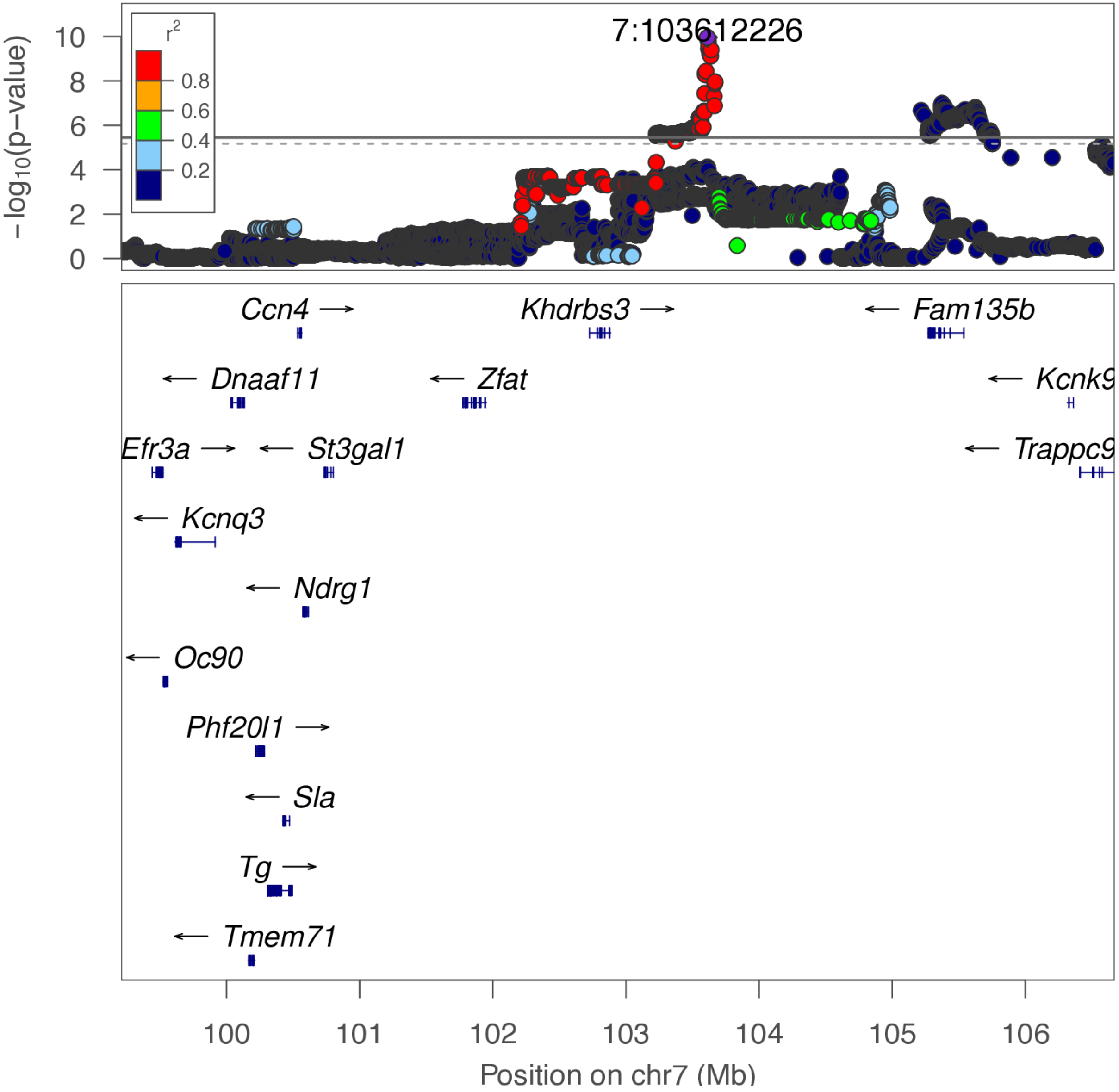

**chr7:105.6 Mb - MEGA - narrow**

Lead: 7:105,217,421 | Genes in QTL interval: 4

QTL interval: chr7:105,217,421-106,595,972

Displayed region: chr7:105,017,421-107,554,972

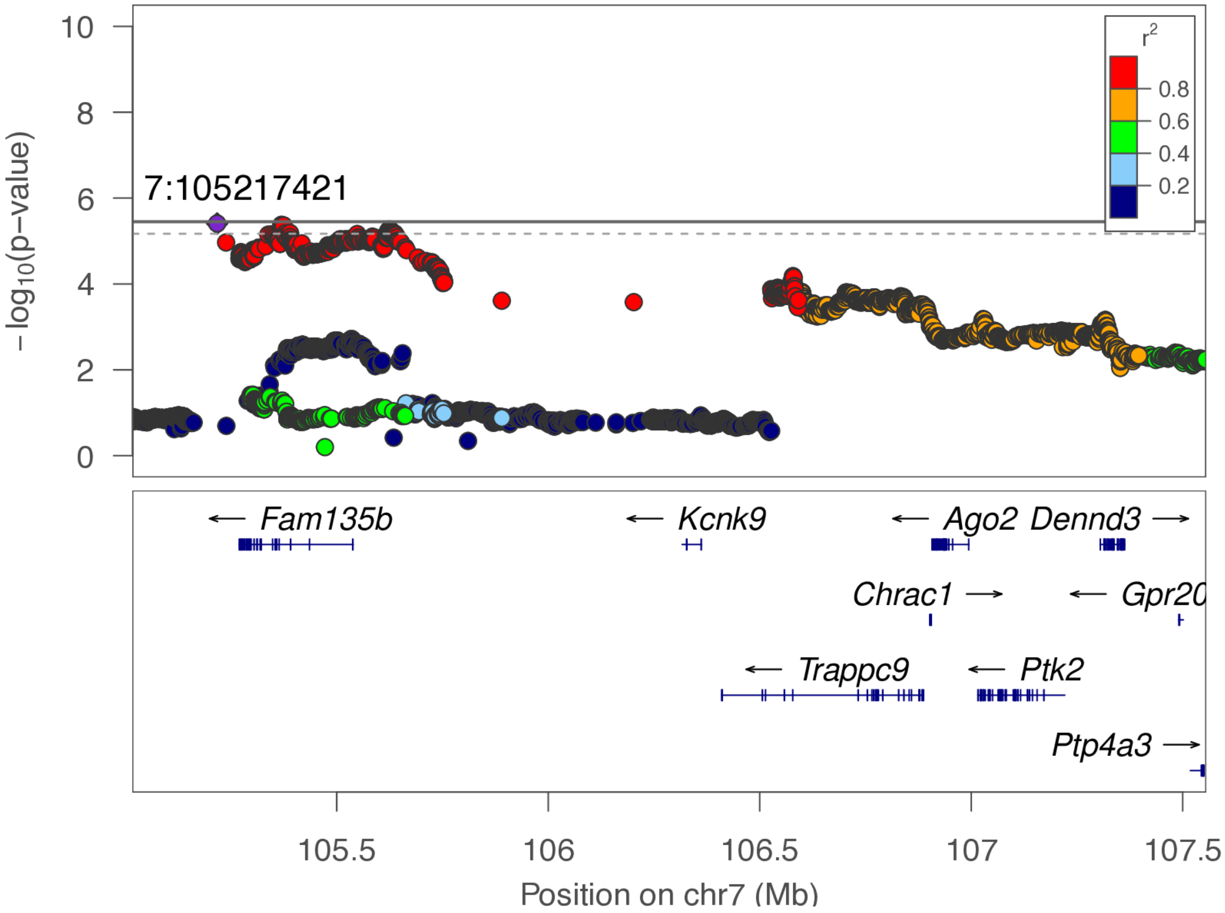

**chr7:105.6 Mb - MEGA - broad**

Lead: 7:105,217,421 | Genes in QTL interval: 4

QTL interval: chr7:105,217,421-106,595,972

Displayed region: chr7:102,217,421-110,354,972

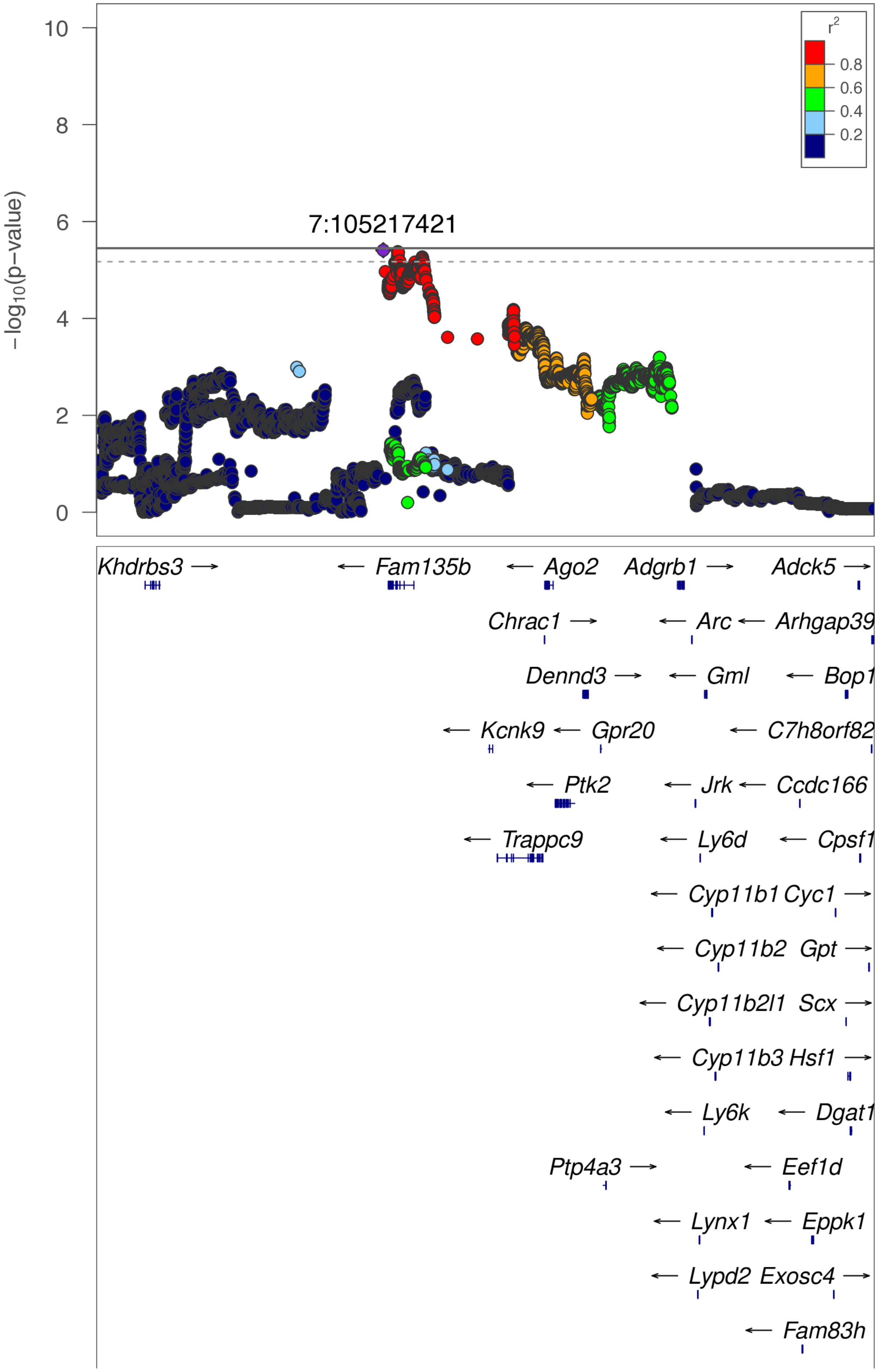

**chr7:105.6 Mb - MEGA - broad (continued)**

Lead: 7:105,217,421 | Genes in QTL interval: 4

QTL interval: chr7:105,217,421-106,595,972

Displayed region: chr7:102,217,421-110,354,972

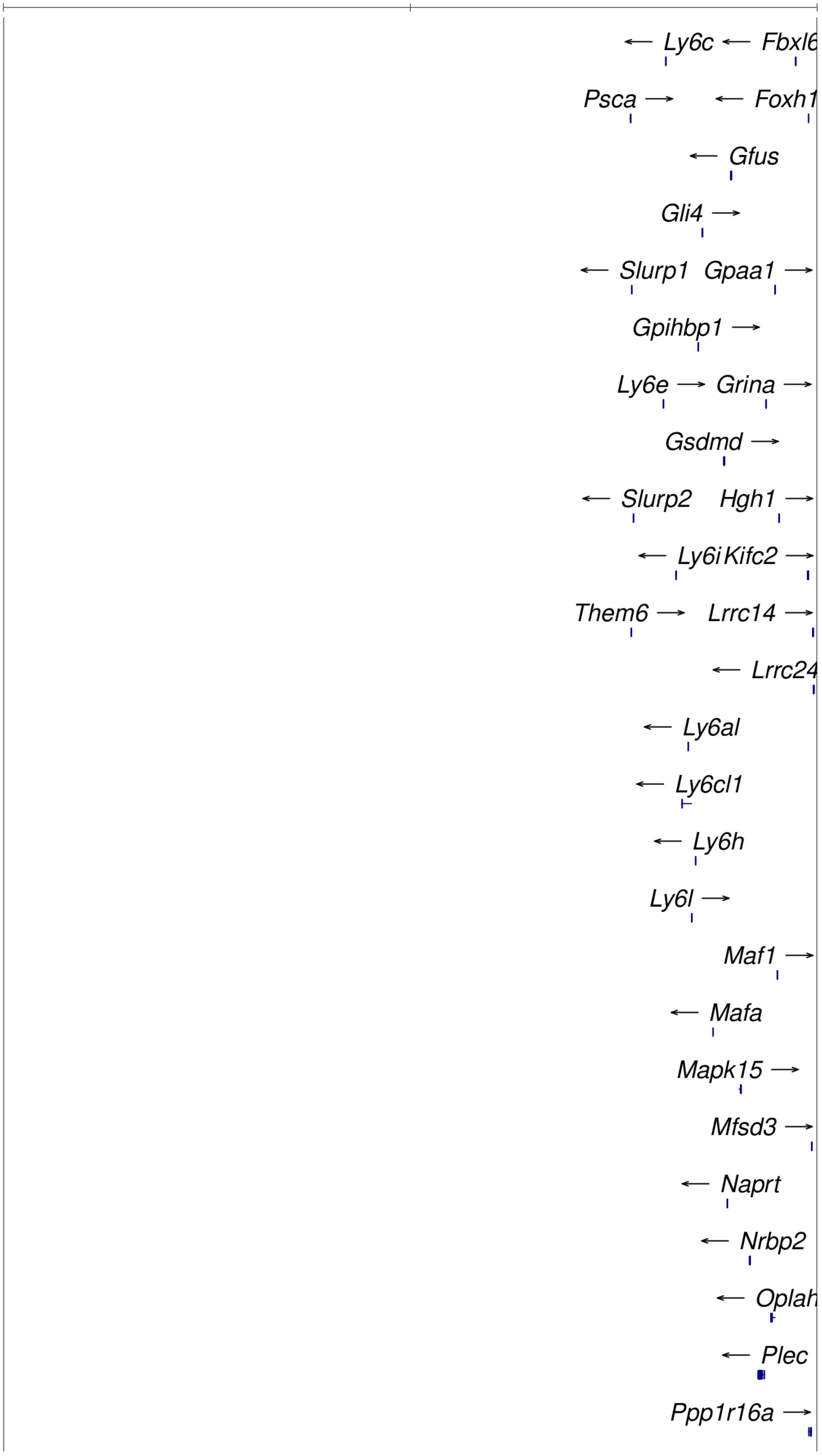

**chr7:105.6 Mb - MEGA - broad (continued)**

Lead: 7:105,217,421 | Genes in QTL interval: 4

QTL interval: chr7:105,217,421-106,595,972

Displayed region: chr7:102,217,421-110,354,972

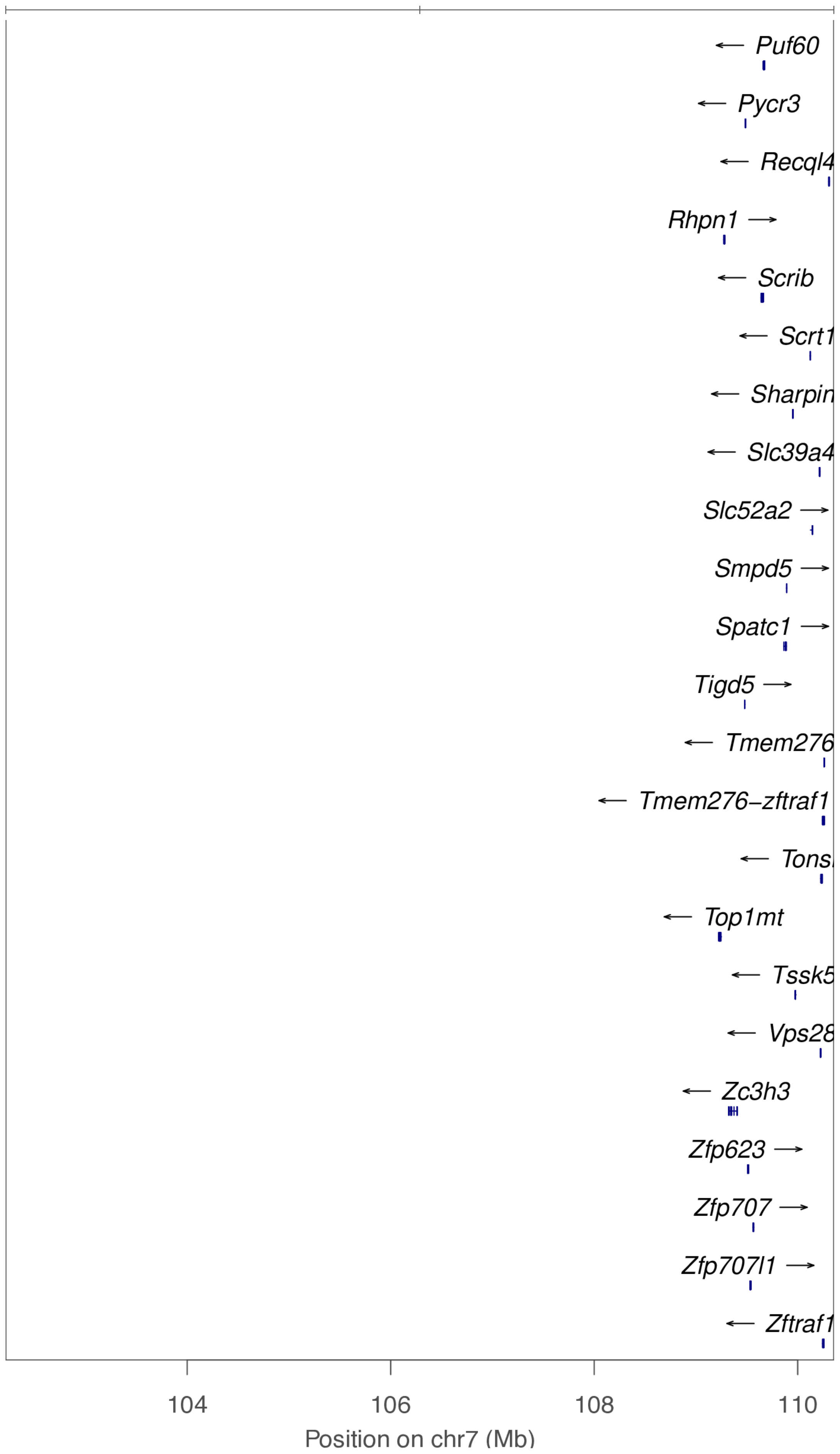

**chr7:105.6 Mb - METAL - narrow**

Lead: 7:105,624,832 | Genes in QTL interval: 4

QTL interval: chr7:105,217,421-106,595,972

Displayed region: chr7:105,017,421-107,545,163

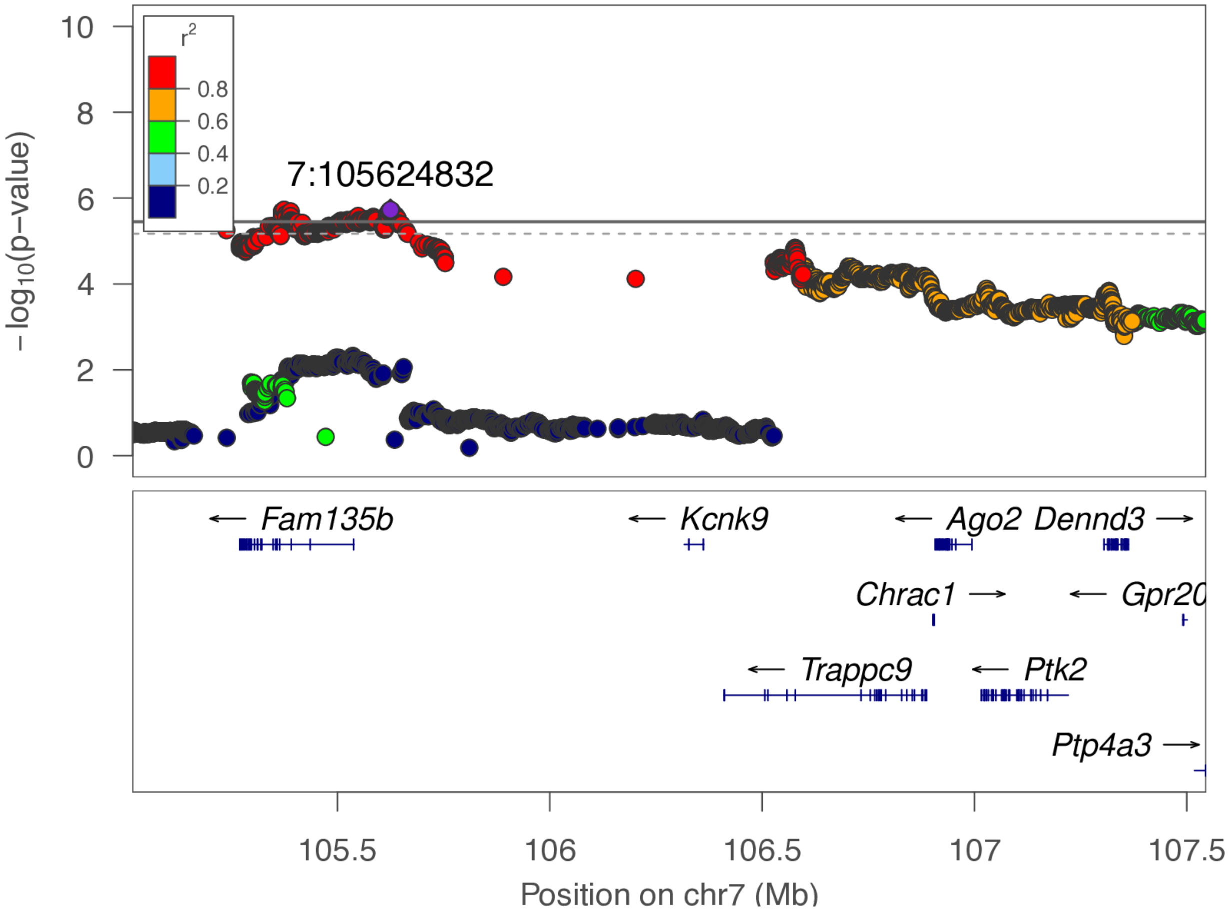

**chr7:105.6 Mb - METAL - broad**

Lead: 7:105,624,832 | Genes in QTL interval: 4

QTL interval: chr7:105,217,421-106,595,972

Displayed region: chr7:102,217,421-110,345,163

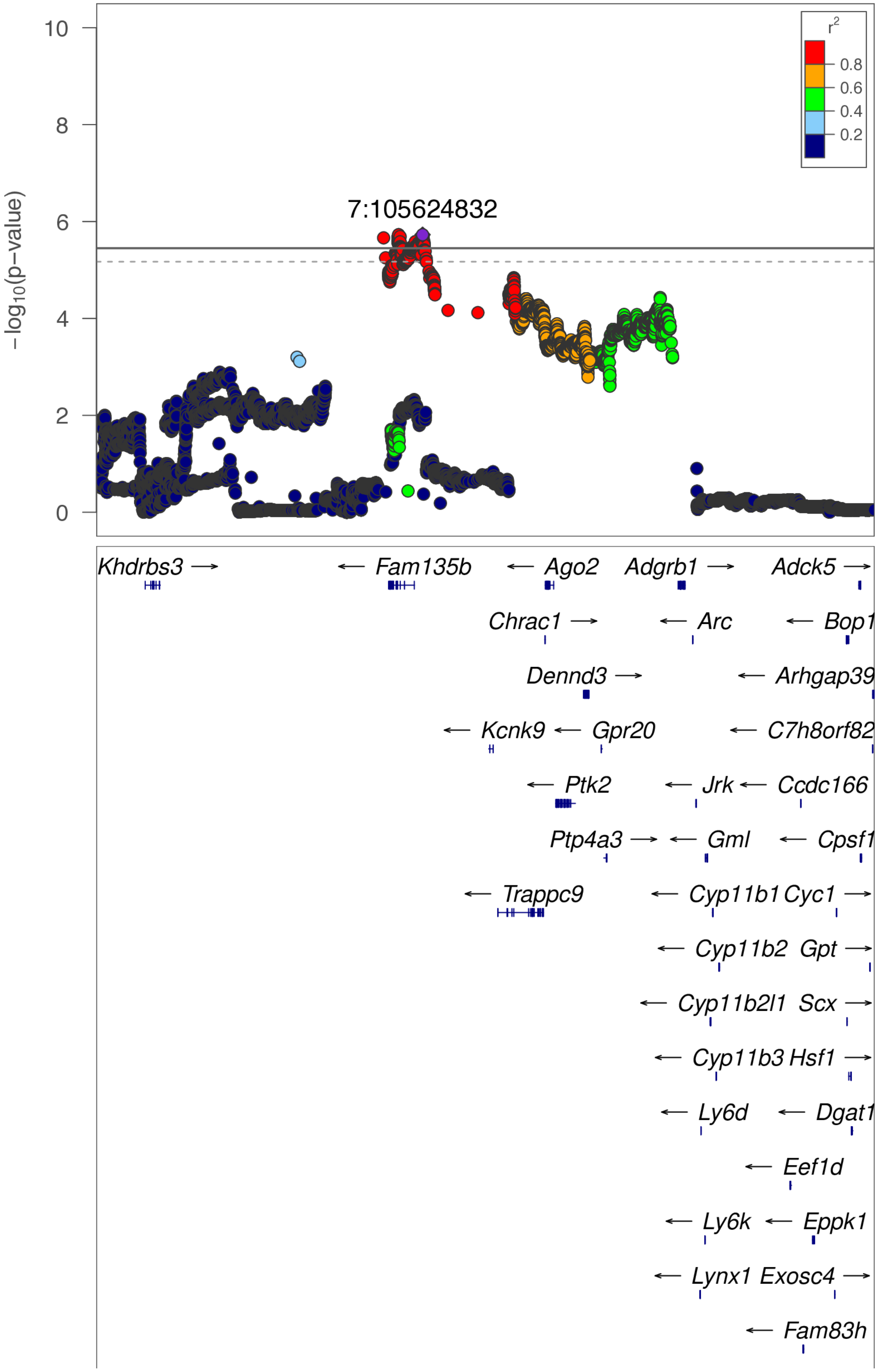

**chr7:105.6 Mb - METAL - broad (continued)**

Lead: 7:105,624,832 | Genes in QTL interval: 4

QTL interval: chr7:105,217,421-106,595,972

Displayed region: chr7:102,217,421-110,345,163

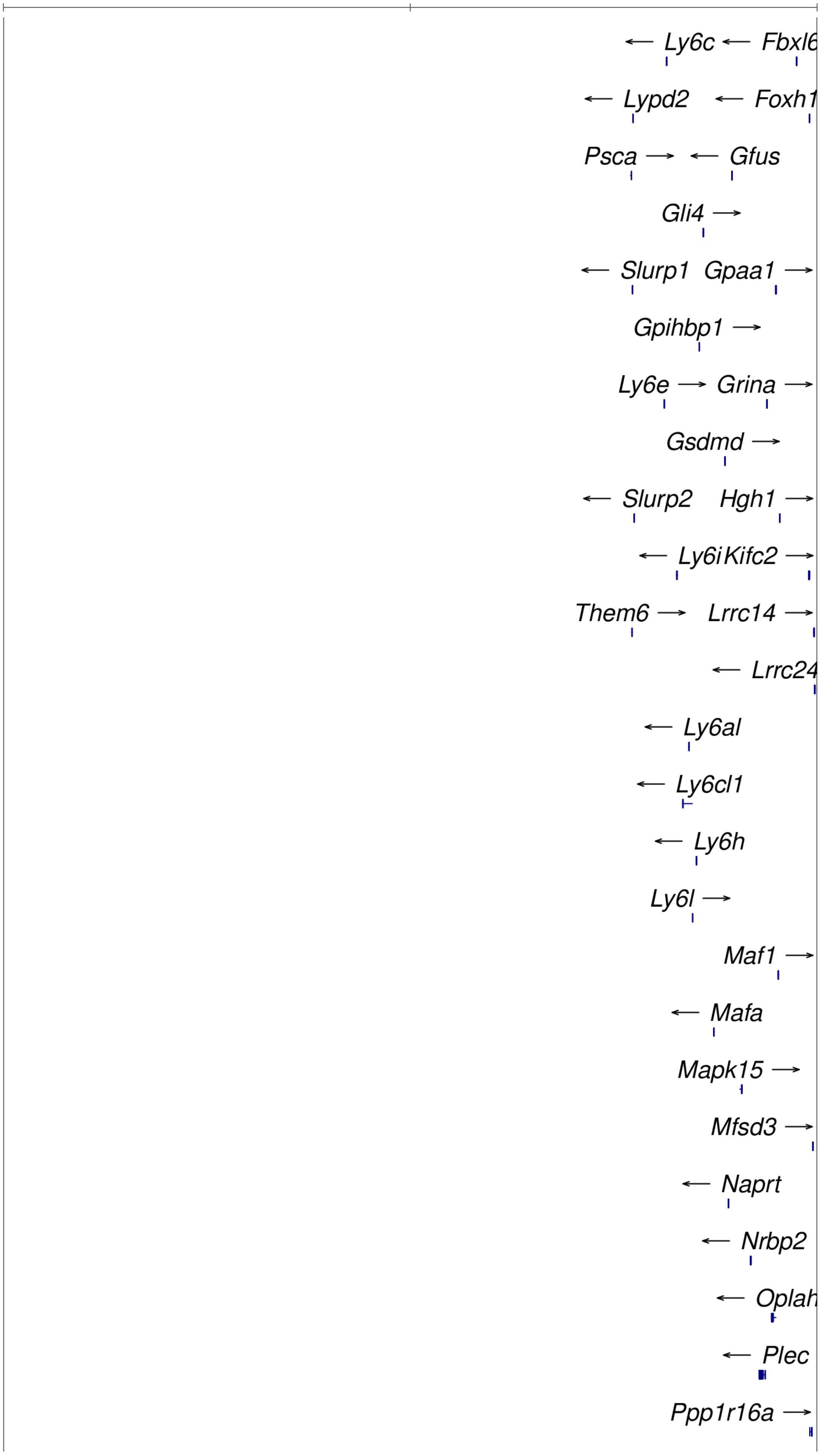

**chr7:105.6 Mb - METAL - broad (continued)**

Lead: 7:105,624,832 | Genes in QTL interval: 4

QTL interval: chr7:105,217,421-106,595,972

Displayed region: chr7:102,217,421-110,345,163

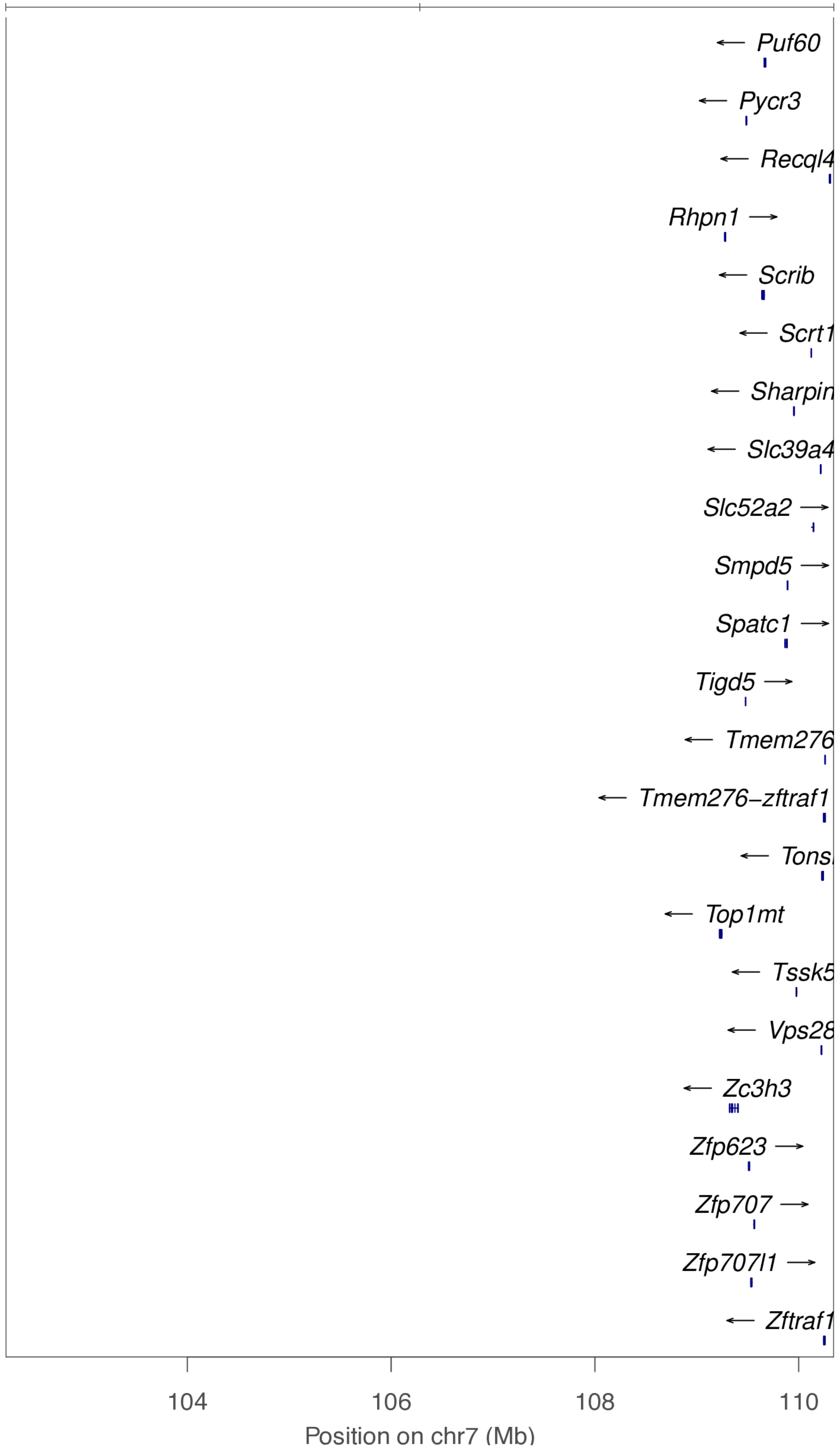

**chr8:91.3 Mb - Genomic SEM - narrow**

Lead: 8:91,271,154 | Genes in QTL interval: 2

QTL interval: chr8:90,878,612-91,893,276

Displayed region: chr8:89,678,105-92,108,103

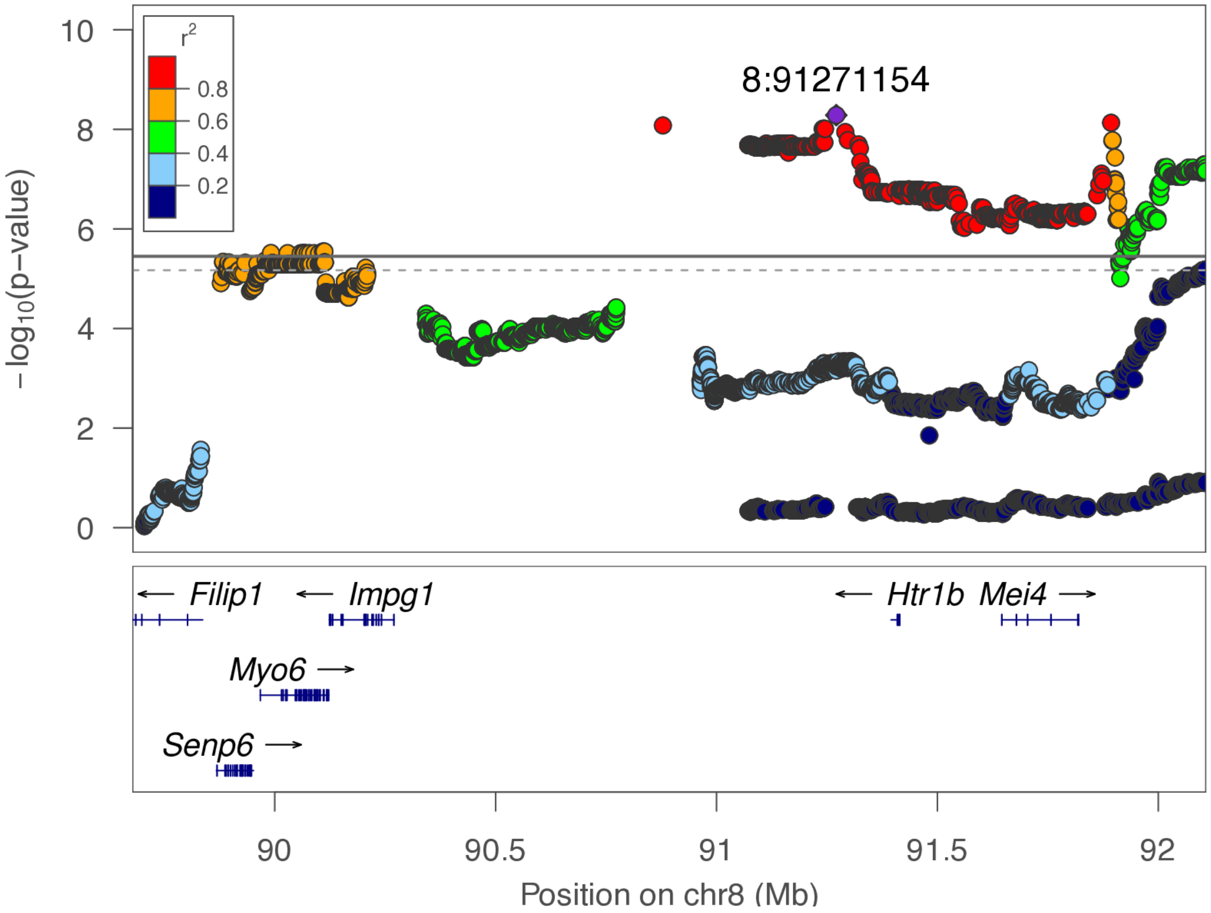

**chr8:91.3 Mb - Genomic SEM - broad**

Lead: 8:91,271,154 | Genes in QTL interval: 2

QTL interval: chr8:90,878,612-91,893,276

Displayed region: chr8:86,878,105-94,908,103

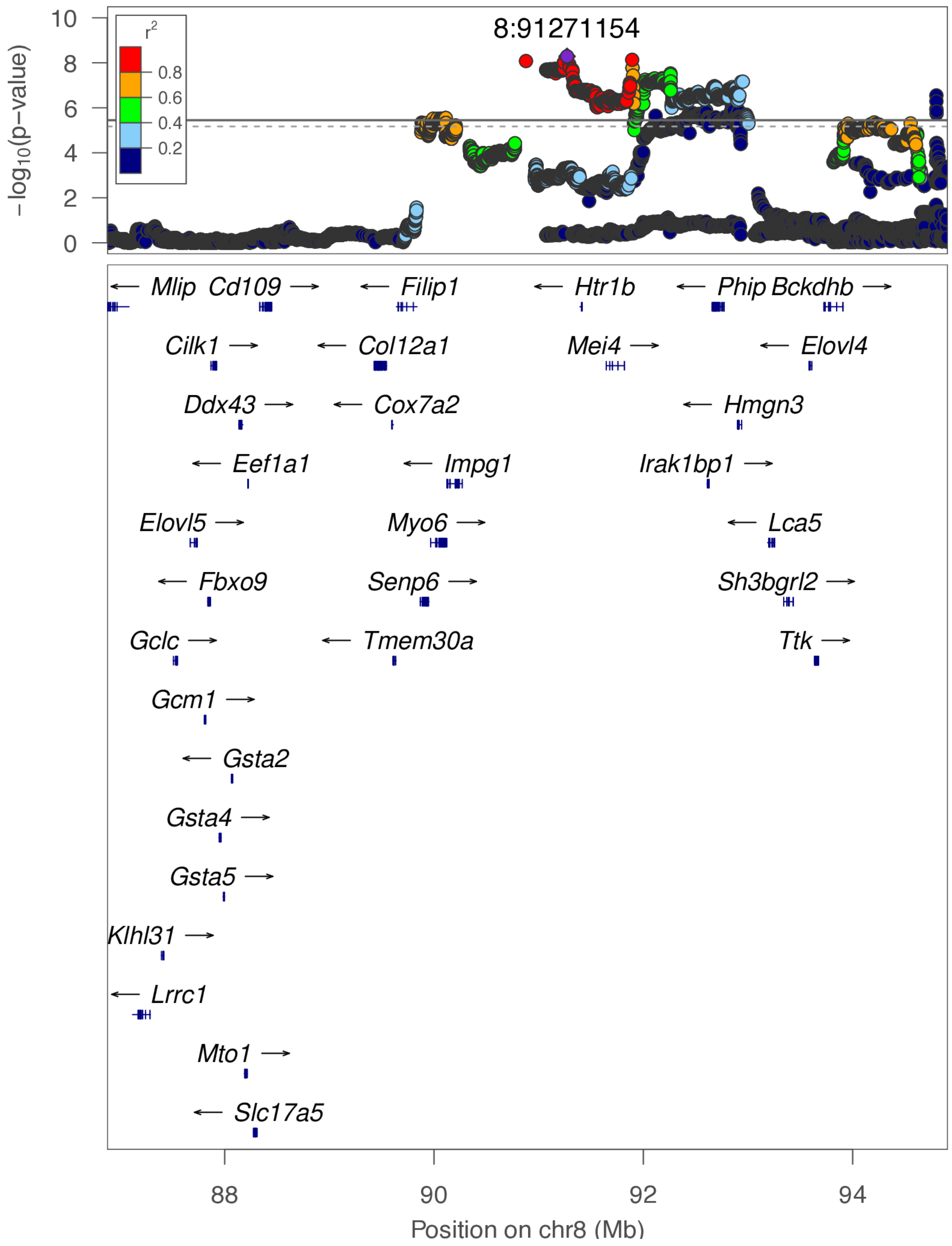

**chr8:92.6 Mb - METAL - narrow**

Lead: 8:92,600,050 | Genes in QTL interval: 2

QTL interval: chr8:91,879,732-92,930,359

Displayed region: chr8:91,448,694-93,130,359

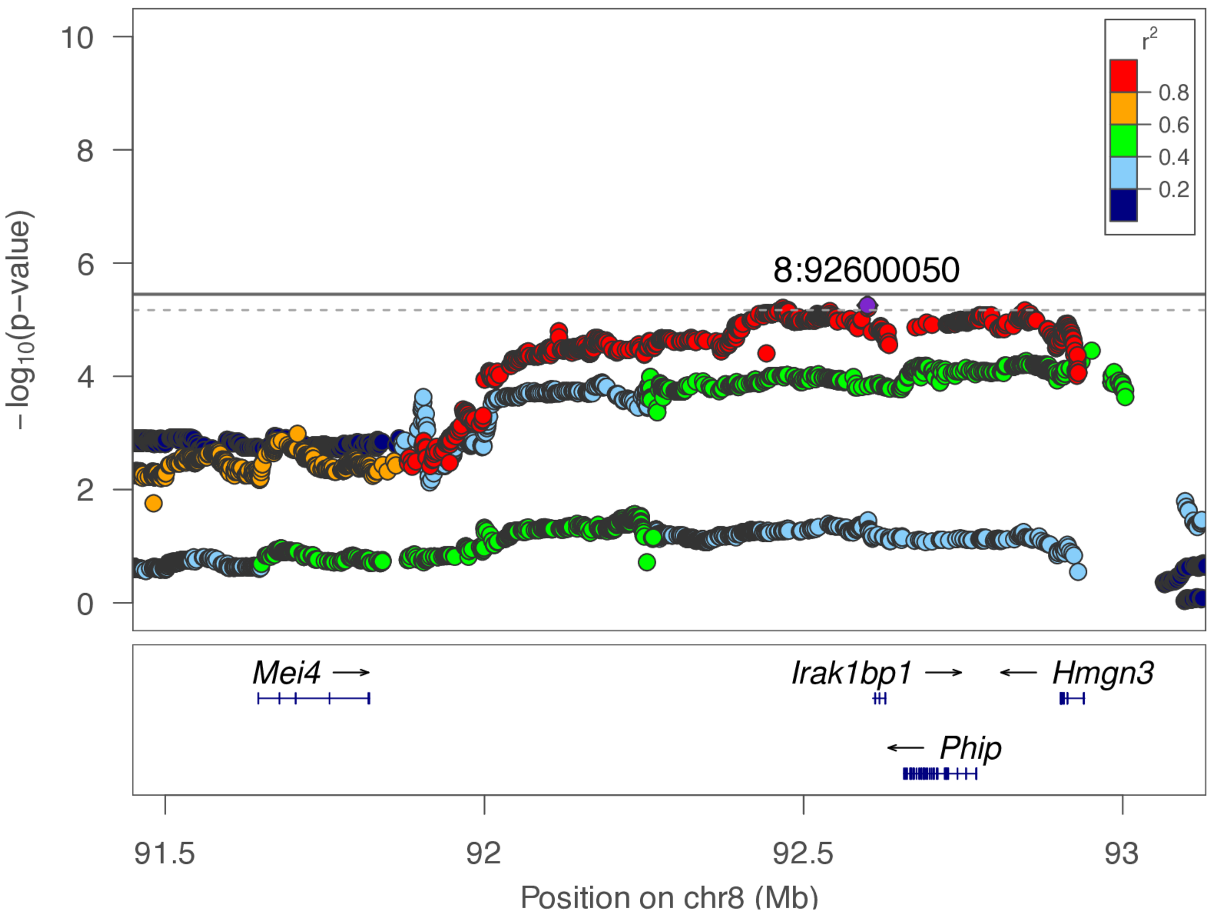

**chr8:92.6 Mb - METAL - broad**

Lead: 8:92,600,050 | Genes in QTL interval: 2

QTL interval: chr8:91,879,732-92,930,359

Displayed region: chr8:88,648,694-95,930,359

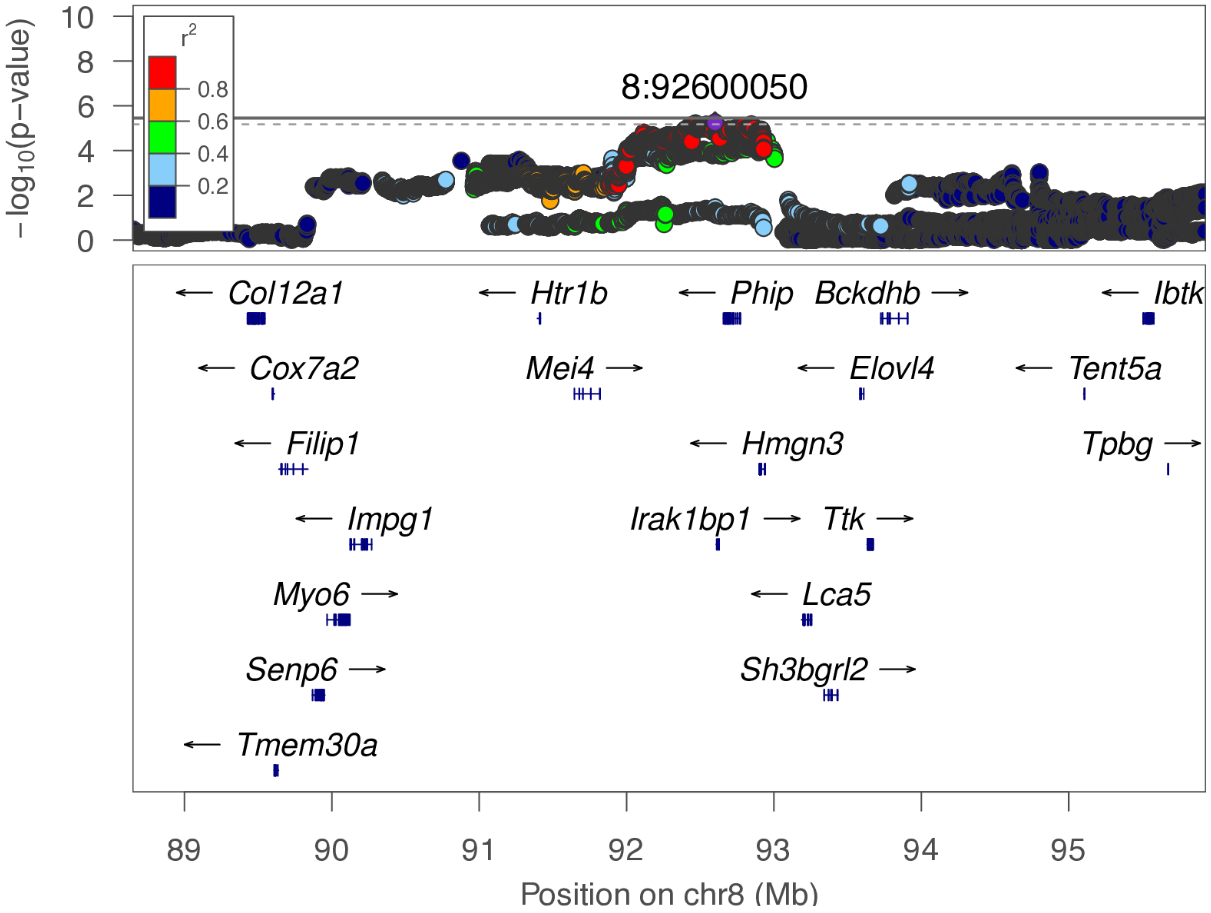

**chr8:121.7 Mb - Genomic SEM - narrow**

Lead: 8:121,711,386 | Genes in QTL interval: 0

QTL interval: chr8:121,506,276-121,807,919

Displayed region: chr8:121,306,276-122,007,919

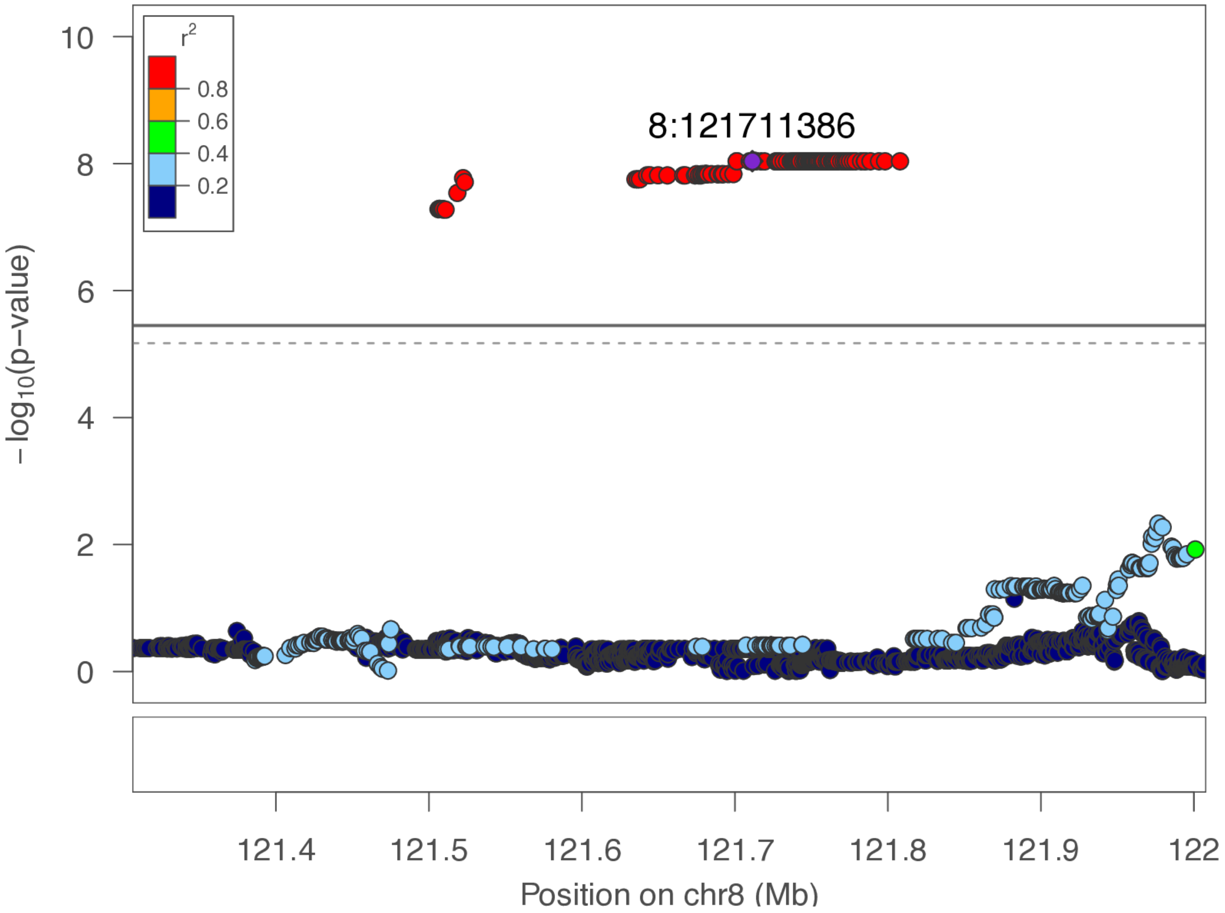

**chr8:121.7 Mb - Genomic SEM - broad**

Lead: 8:121,711,386 | Genes in QTL interval: 0

QTL interval: chr8:121,506,276-121,807,919

Displayed region: chr8:118,506,276-124,807,919

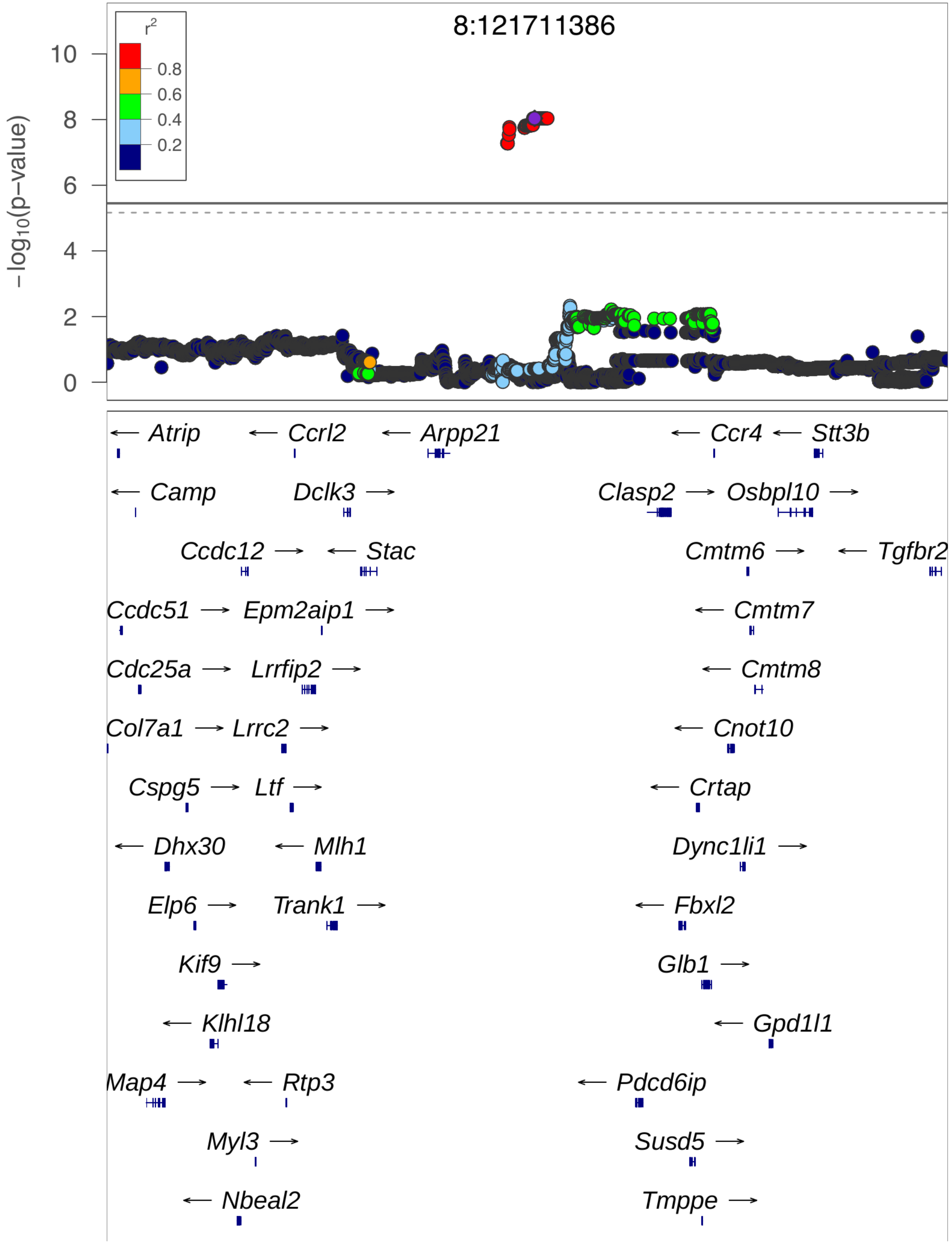

**chr8:121.7 Mb - Genomic SEM - broad (continued)**

Lead: 8:121,711,386 | Genes in QTL interval: 0

QTL interval: chr8:121,506,276-121,807,919

Displayed region: chr8:118,506,276-124,807,919

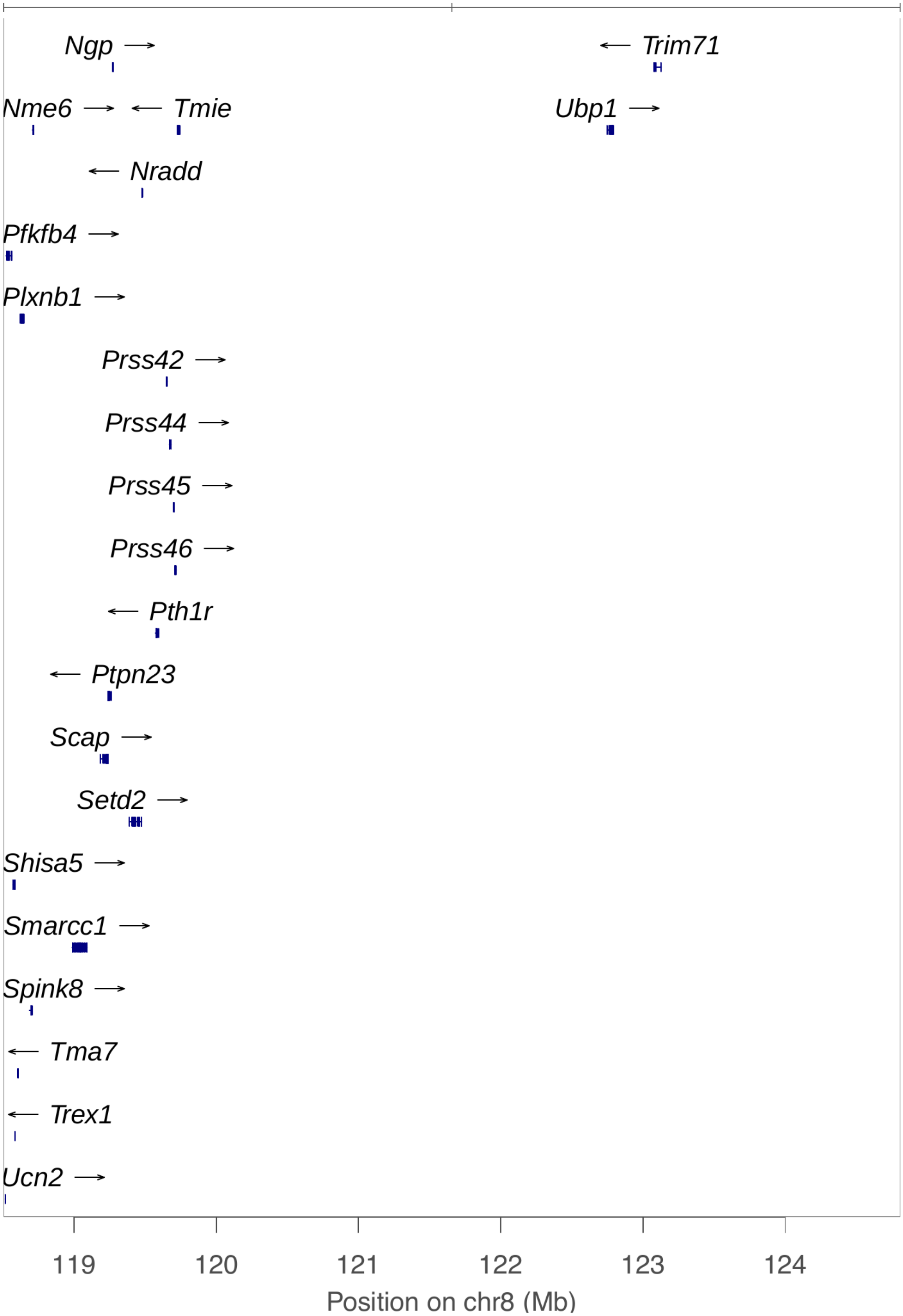

**chr9:10.3 Mb - Genomic SEM - narrow**

Lead: 9:10,273,927 | Genes in QTL interval: 30

QTL interval: chr9:6,595,114-18,312,239

Displayed region: chr9:6,395,114-17,417,344

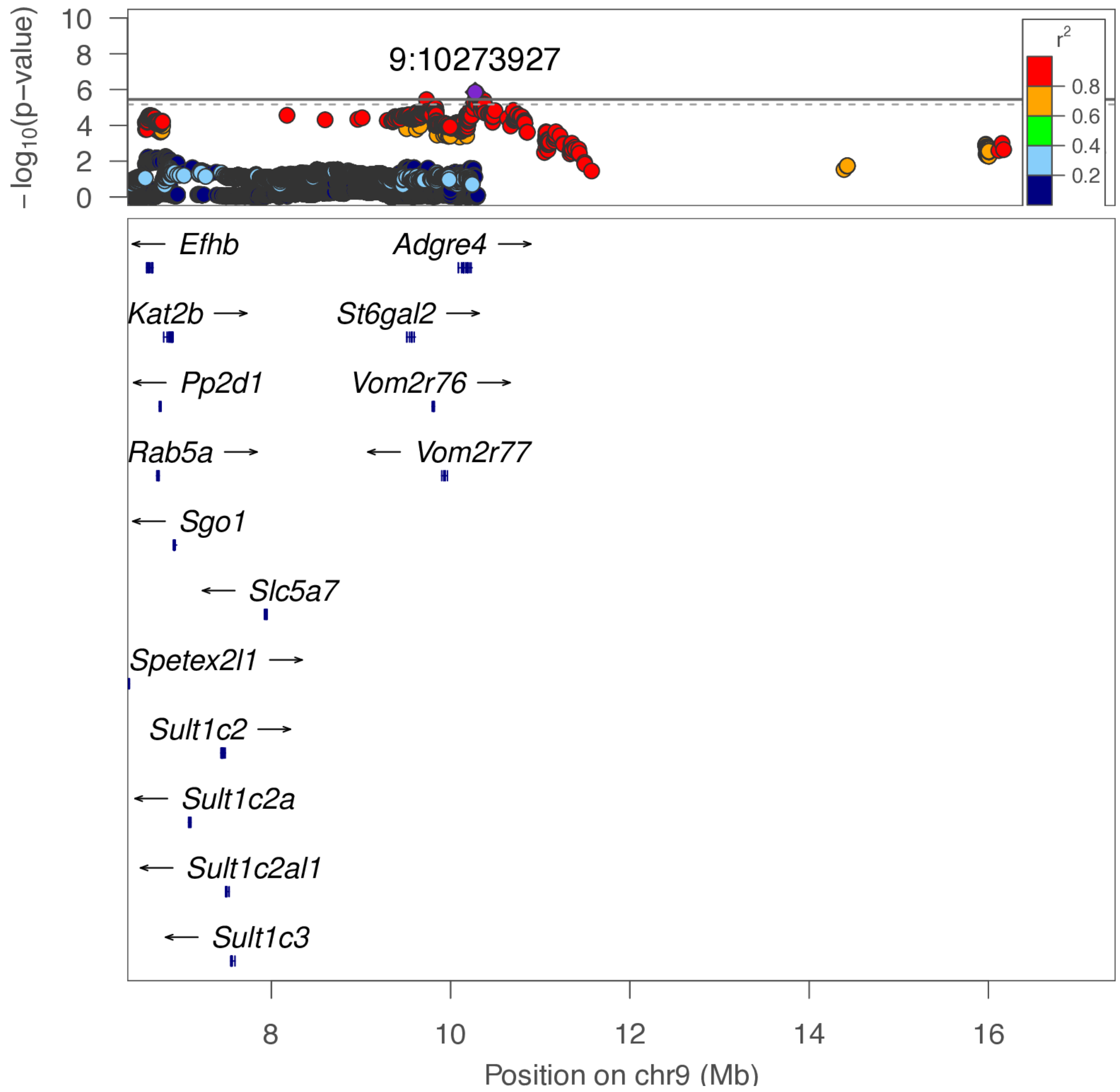

**chr9:10.3 Mb - Genomic SEM - broad**

Lead: 9:10,273,927 | Genes in QTL interval: 30

QTL interval: chr9:6,595,114-18,312,239

Displayed region: chr9:3,595,114-20,217,344

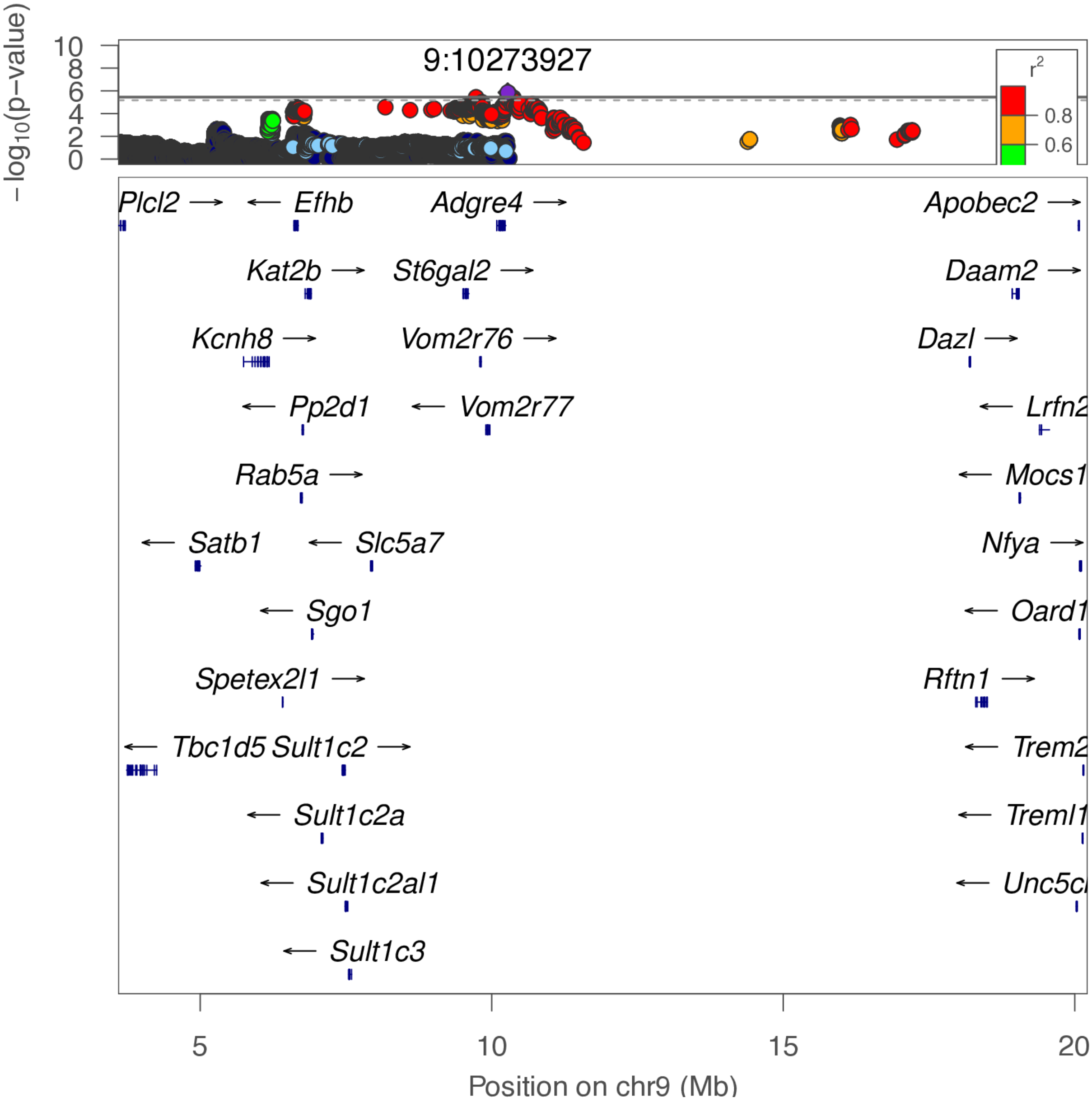

**chr10:19.7 Mb - METAL - narrow**

Lead: 10:19,688,707 | Genes in QTL interval: 2

QTL interval: chr10:19,682,350-20,032,548

Displayed region: chr10:19,482,350-20,775,475

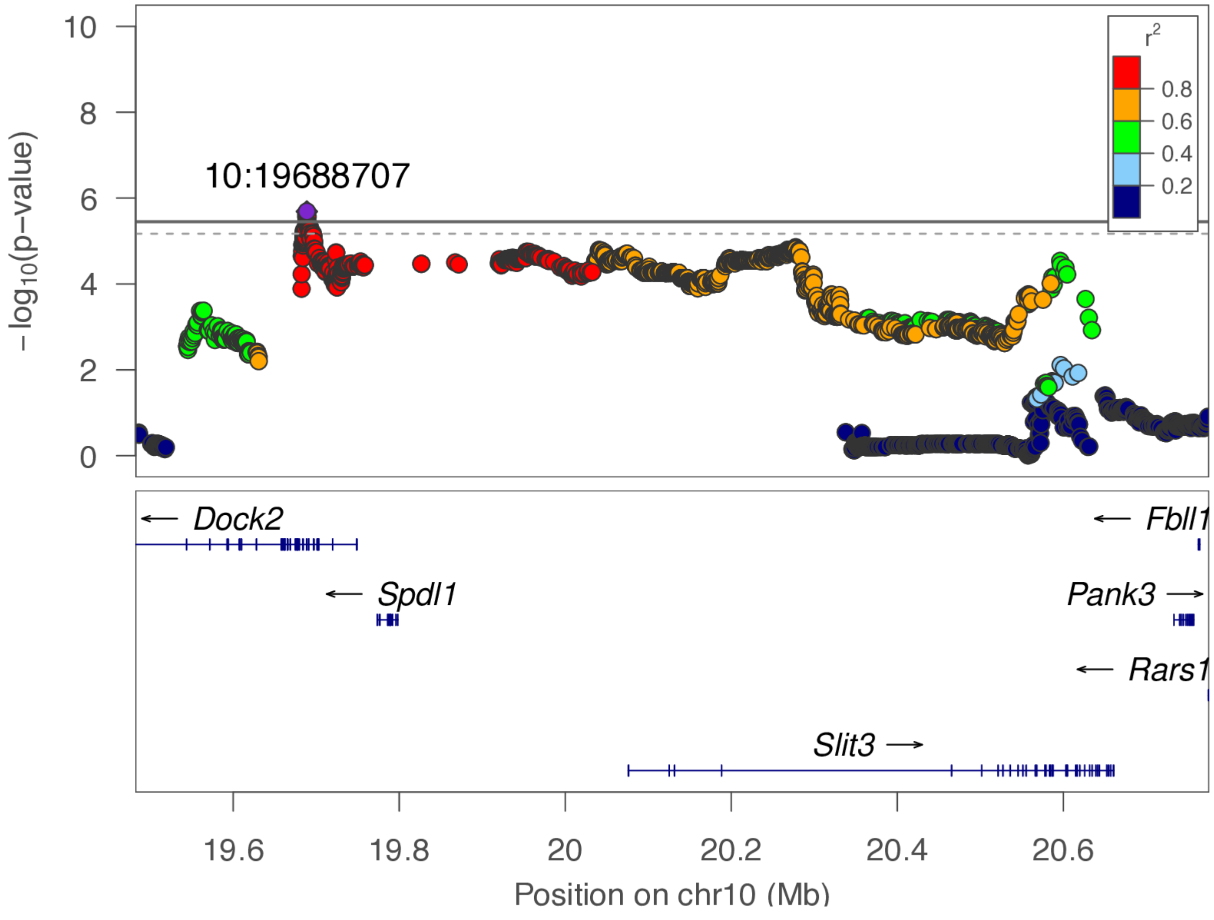

**chr10:19.7 Mb - METAL - broad**

Lead: 10:19,688,707 | Genes in QTL interval: 2

QTL interval: chr10:19,682,350-20,032,548

Displayed region: chr10:16,682,350-23,575,475

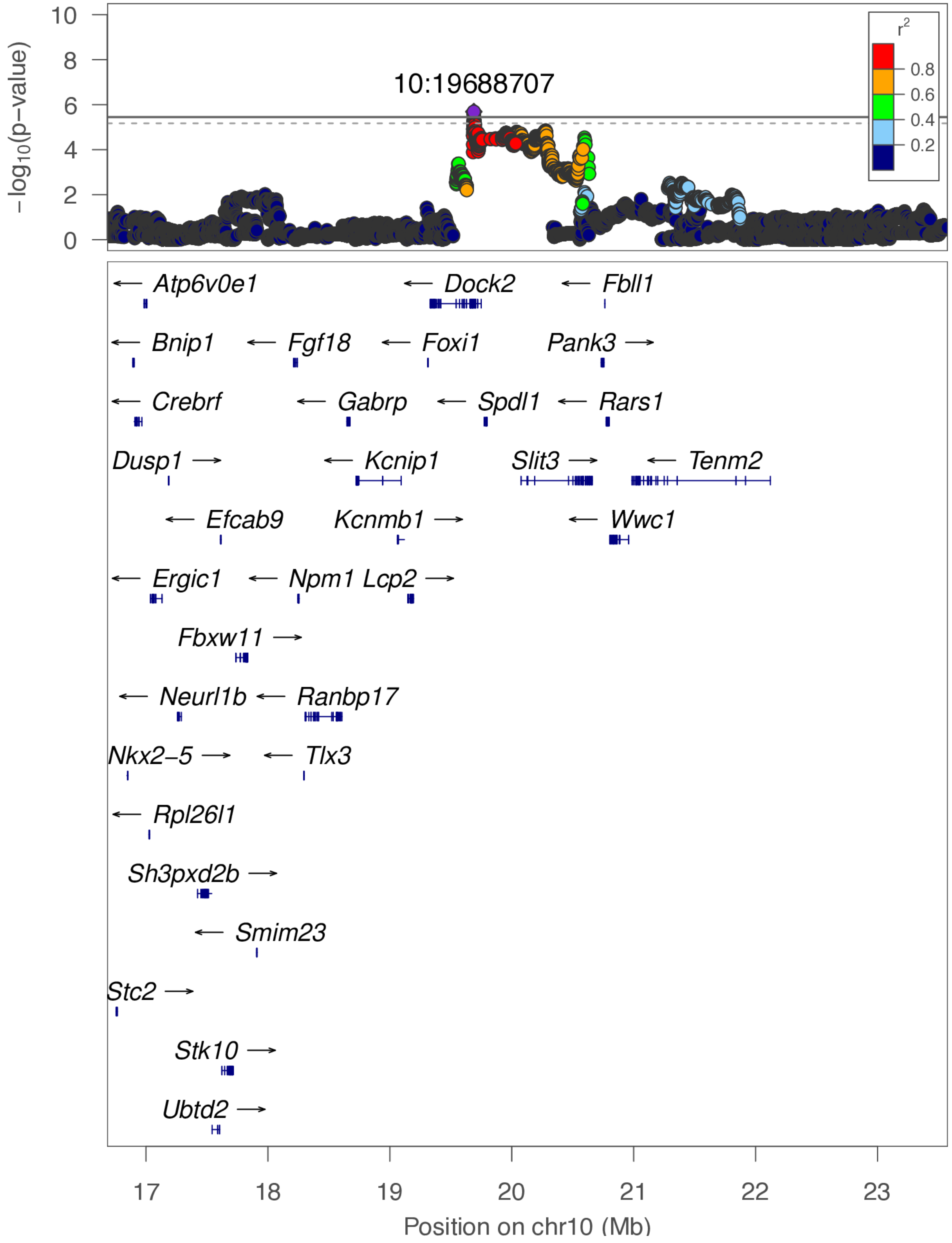

**chr10:19.7 Mb - Genomic SEM - narrow**

Lead: 10:19,688,707 | Genes in QTL interval: 2

QTL interval: chr10:19,682,350-20,032,548

Displayed region: chr10:19,482,350-20,775,475

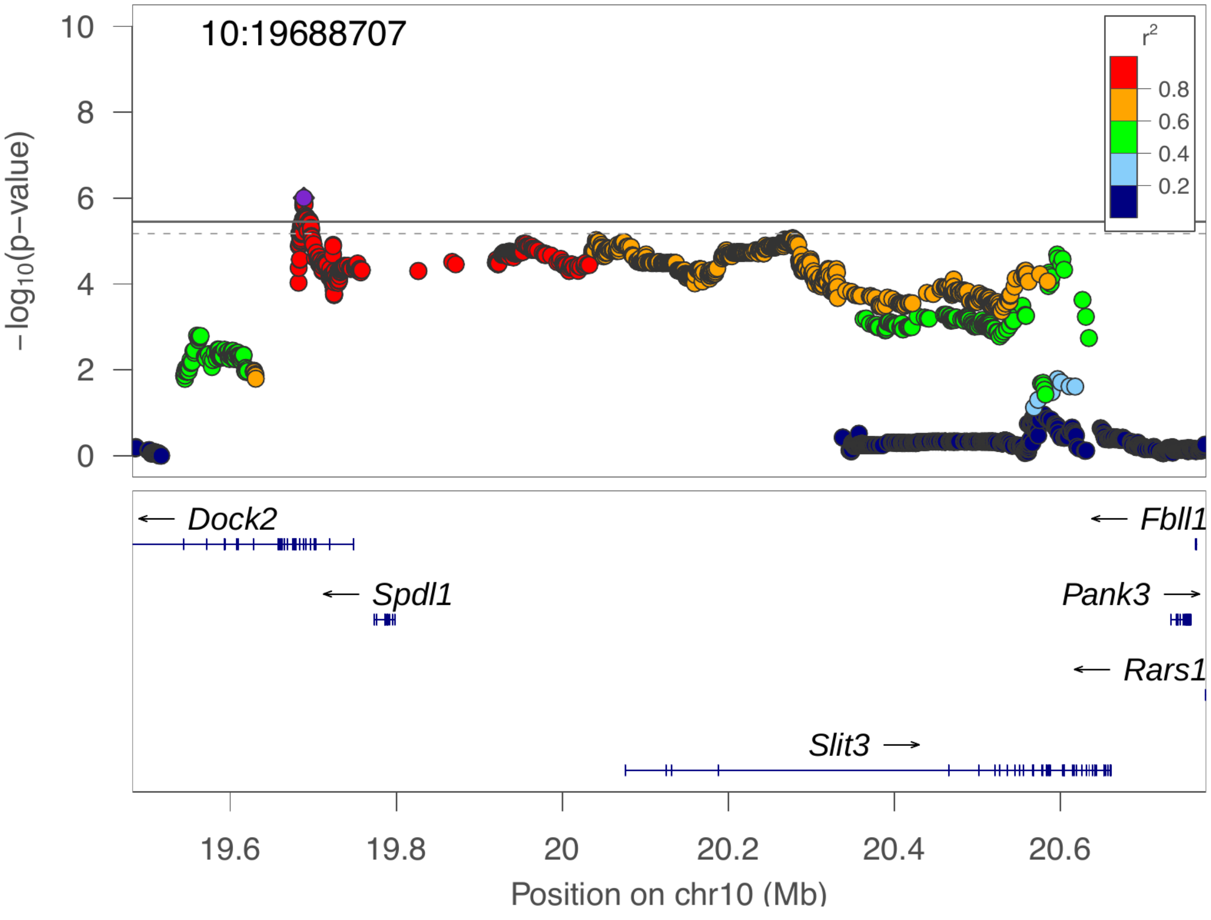

**chr10:19.7 Mb - Genomic SEM - broad**

Lead: 10:19,688,707 | Genes in QTL interval: 2

QTL interval: chr10:19,682,350-20,032,548

Displayed region: chr10:16,682,350-23,575,475

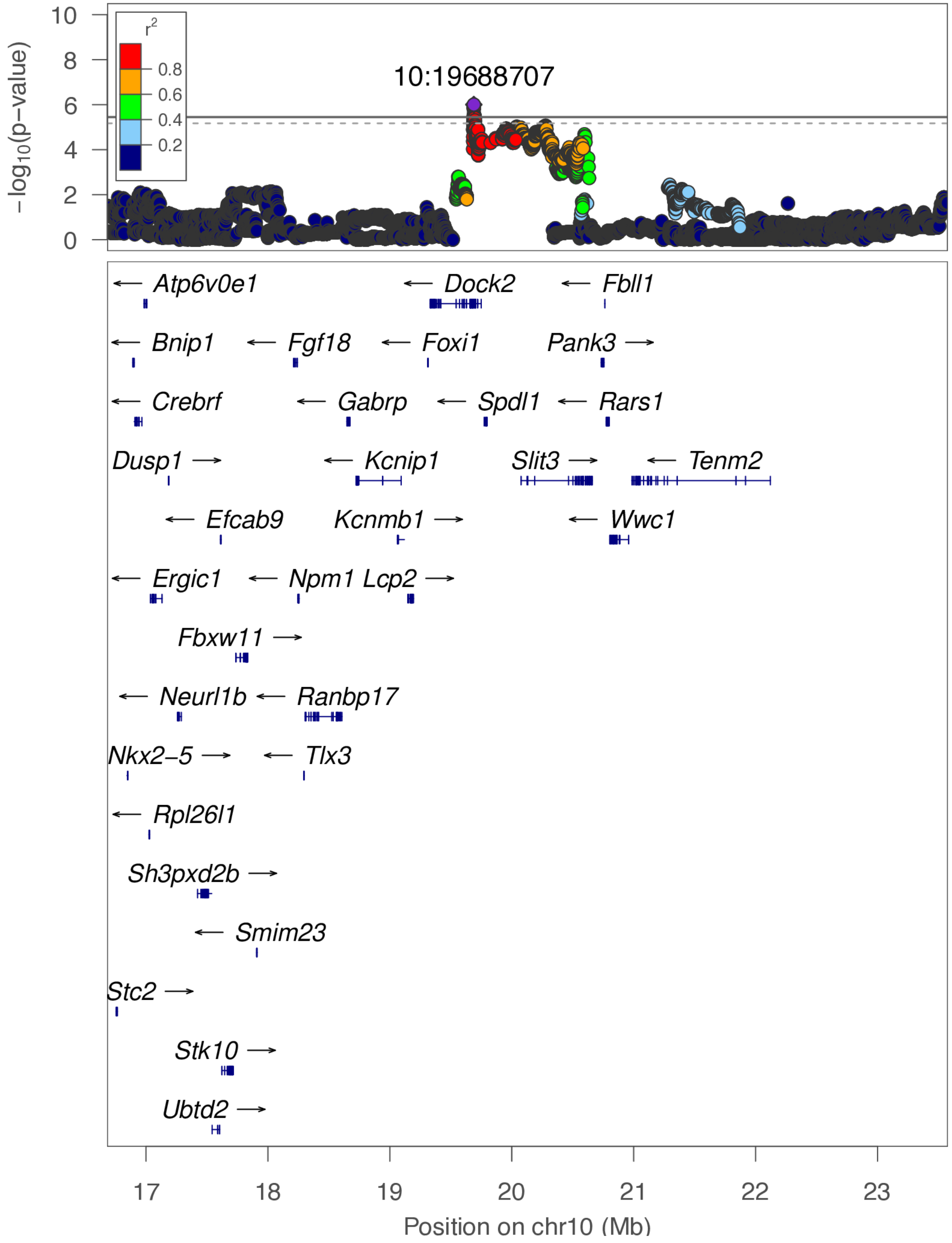

**chr10:26.7 Mb - Genomic SEM - narrow**

Lead: 10:26,668,342 | Genes in QTL interval: 2

QTL interval: chr10:26,483,857-27,074,318

Displayed region: chr10:25,907,459-27,587,816

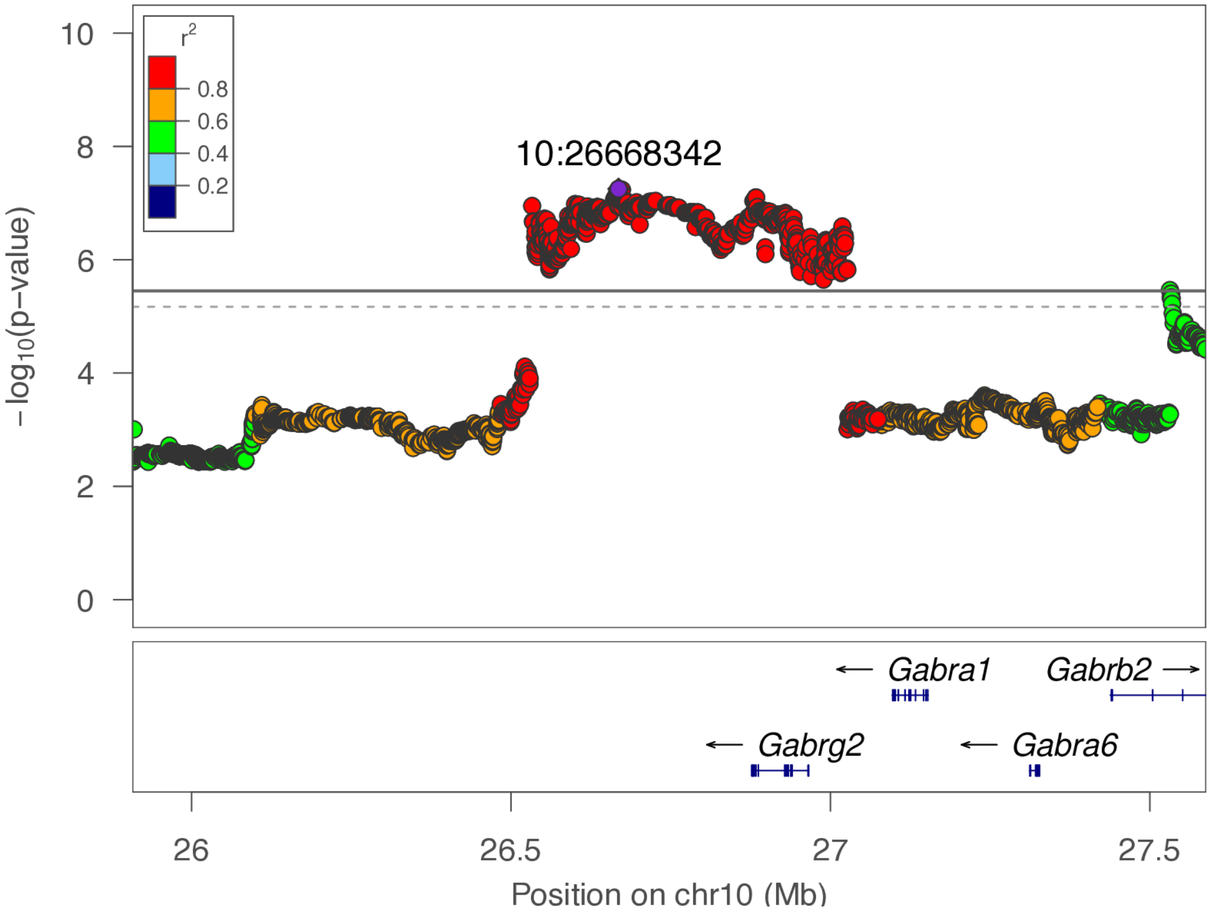

**chr10:26.7 Mb - Genomic SEM - broad**

Lead: 10:26,668,342 | Genes in QTL interval: 2

QTL interval: chr10:26,483,857-27,074,318

Displayed region: chr10:23,107,459-30,387,816

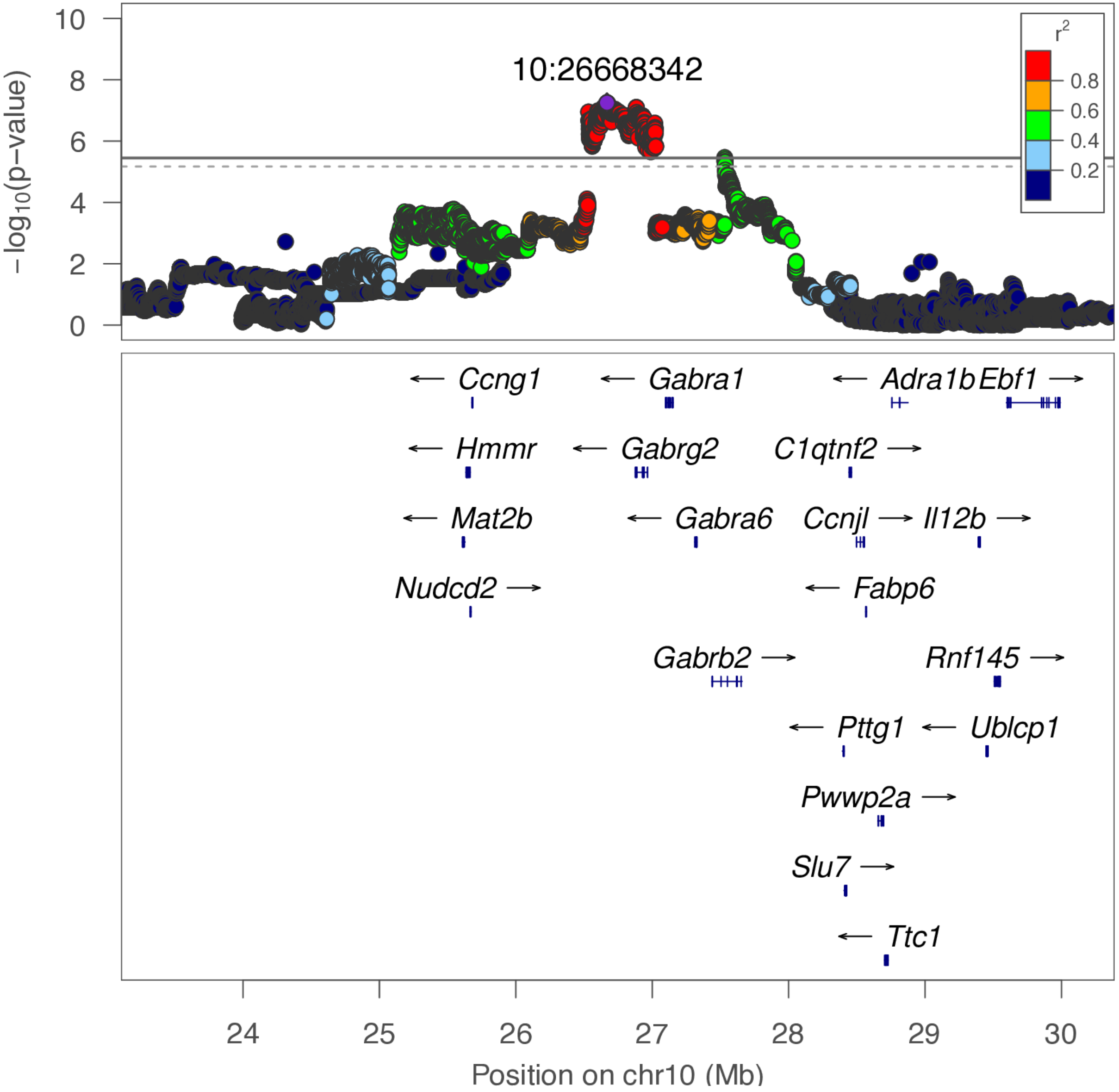

**chr10:67.7 Mb - Genomic SEM - narrow**

Lead: 10:67,678,589 | Genes in QTL interval: 1

QTL interval: chr10:67,659,283-67,843,462

Displayed region: chr10:67,342,044-68,120,122

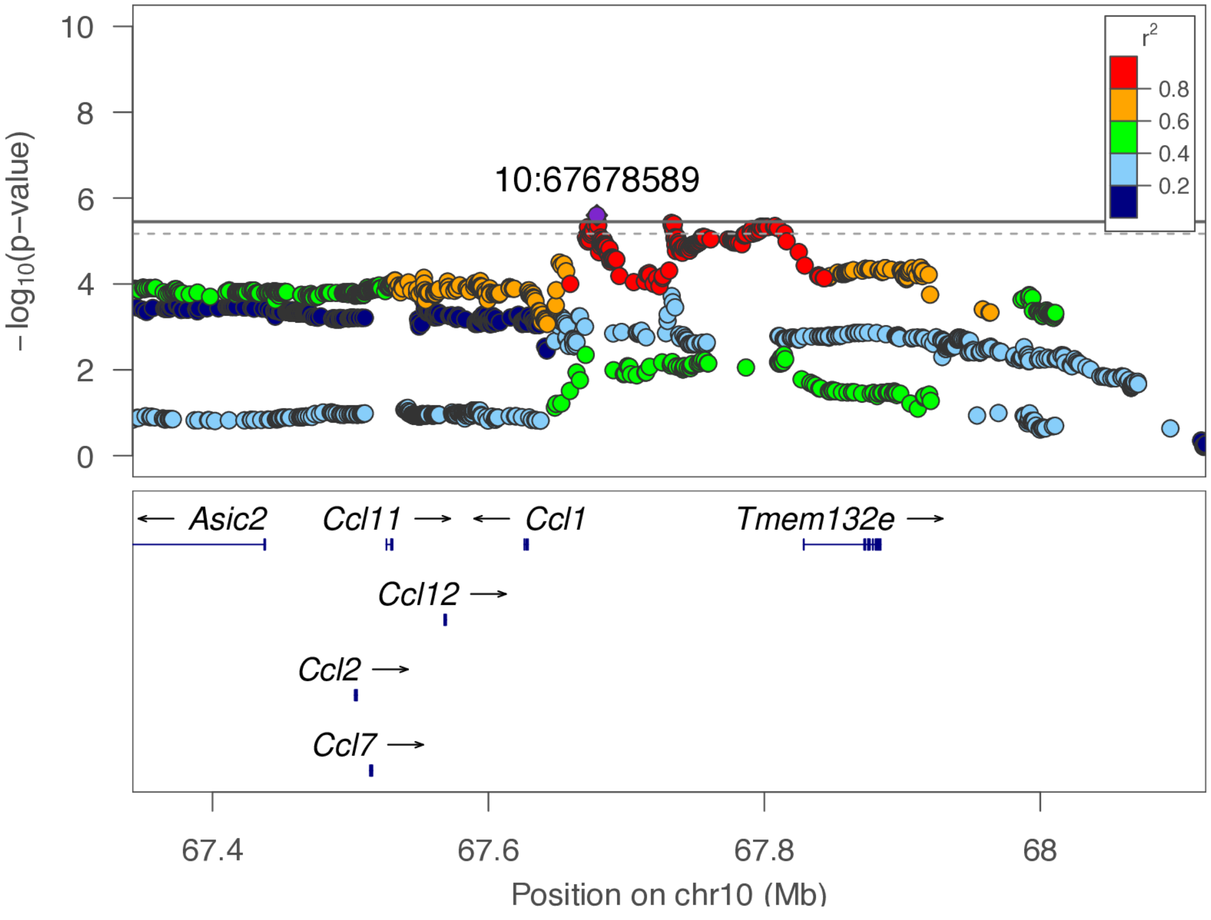

**chr10:67.7 Mb - Genomic SEM - broad**

Lead: 10:67,678,589 | Genes in QTL interval: 1

QTL interval: chr10:67,659,283-67,843,462

Displayed region: chr10:64,542,044-70,920,122

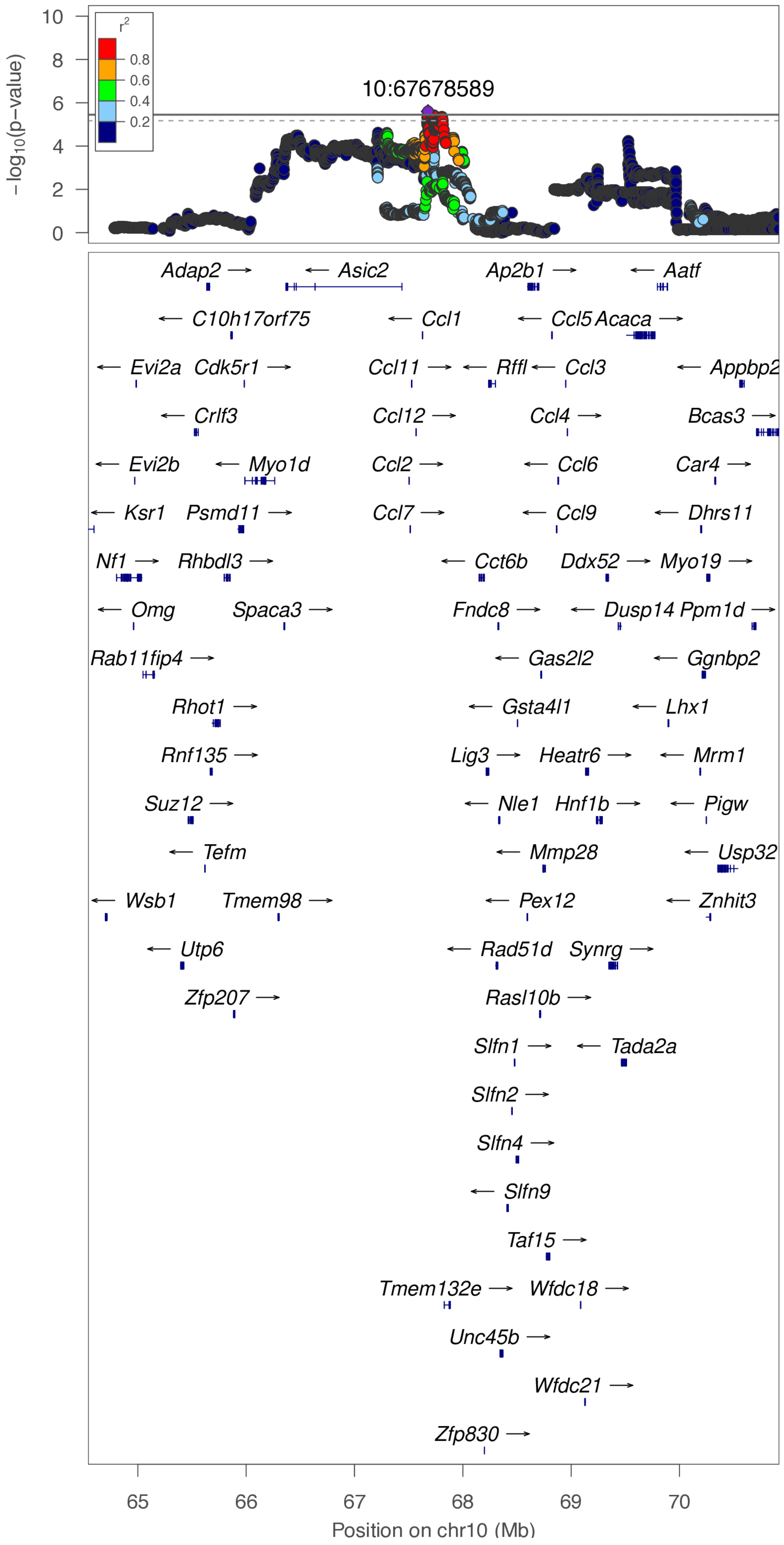

**chr10:91.5 Mb - MEGA - narrow**

Lead: 10:91,726,136 | Genes in QTL interval: 37

QTL interval: chr10:90,586,669-92,719,932

Displayed region: chr10:89,888,192-93,360,153

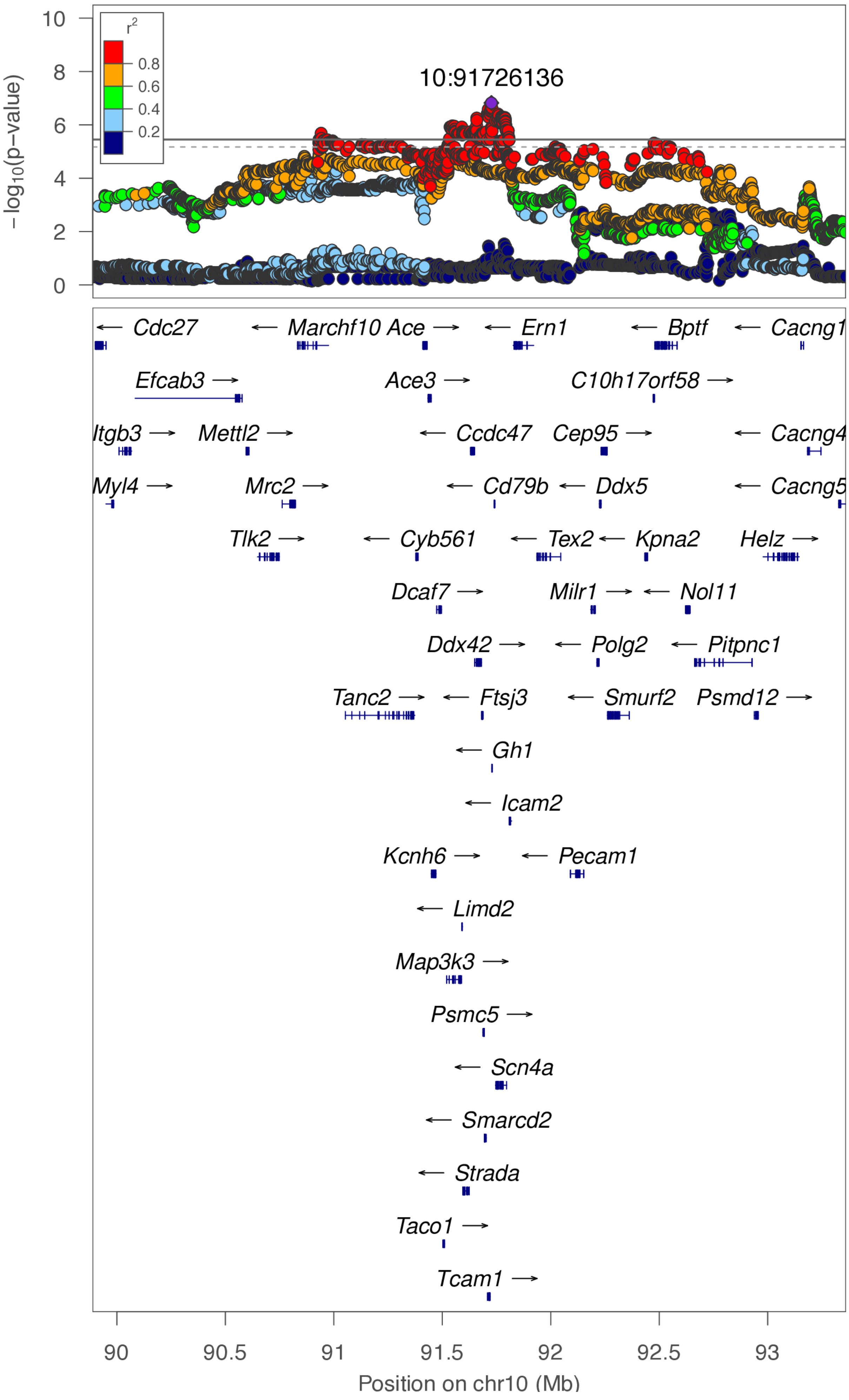

**chr10:91.5 Mb - MEGA - broad**

Lead: 10:91,726,136 | Genes in QTL interval: 37

QTL interval: chr10:90,586,669-92,719,932

Displayed region: chr10:87,088,192-96,160,153

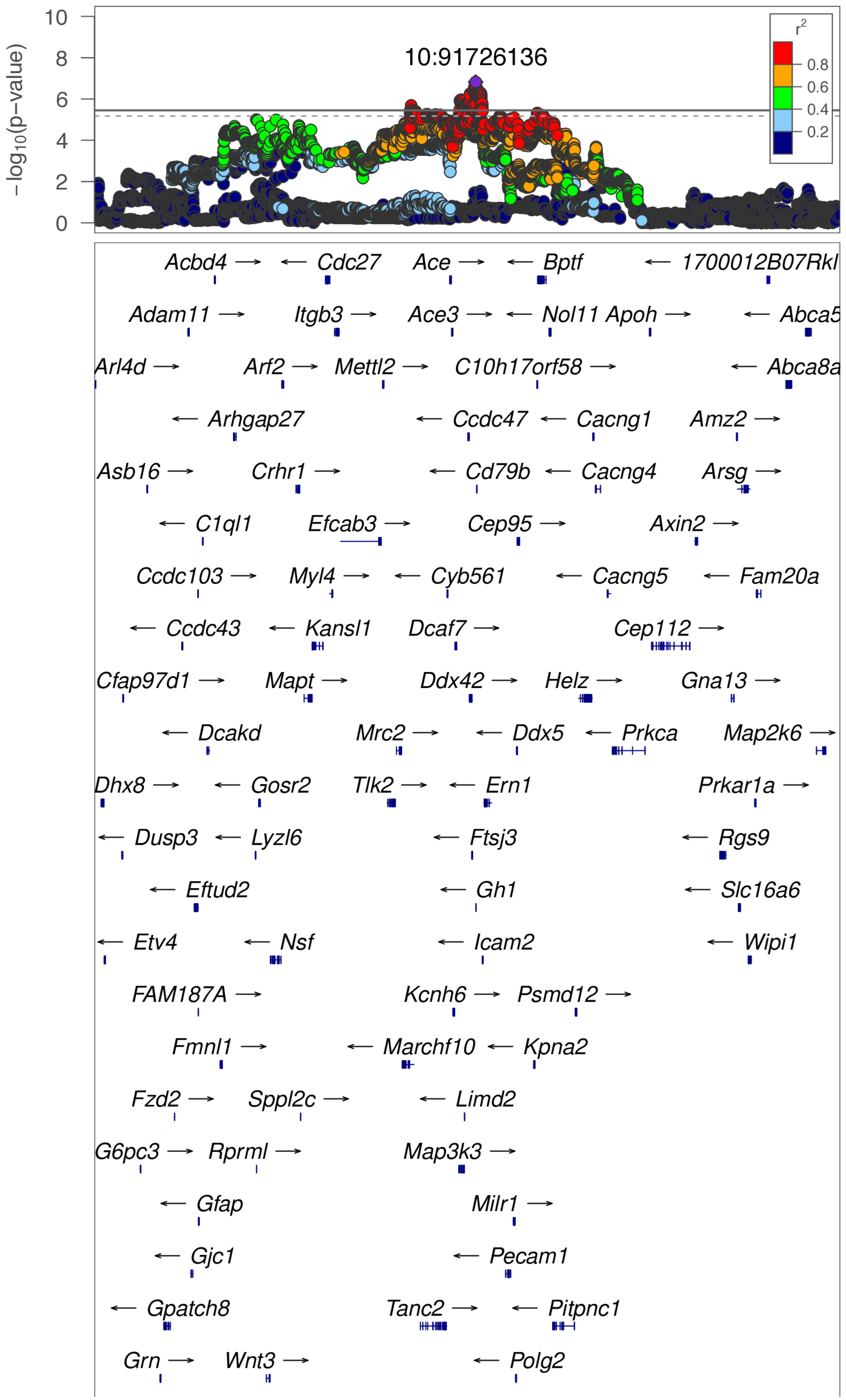

**chr10:91.5 Mb - MEGA - broad (continued)**

Lead: 10:91,726,136 | Genes in QTL interval: 37

QTL interval: chr10:90,586,669-92,719,932

Displayed region: chr10:87,088,192-96,160,153

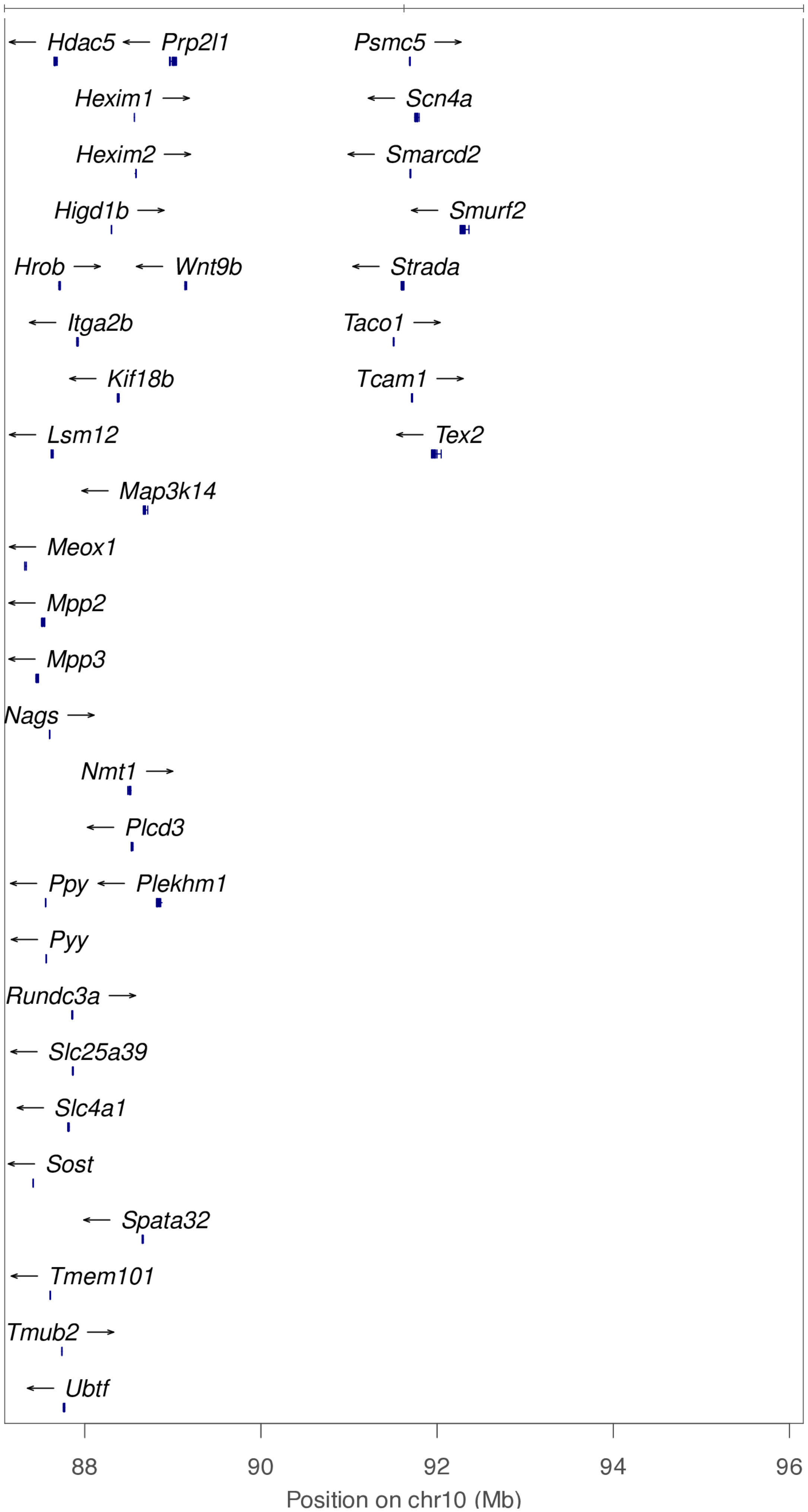

**chr10:91.5 Mb - METAL - narrow**

Lead: 10:91,726,136 | Genes in QTL interval: 37

QTL interval: chr10:90,586,669-92,719,932

Displayed region: chr10:89,888,192-93,357,889

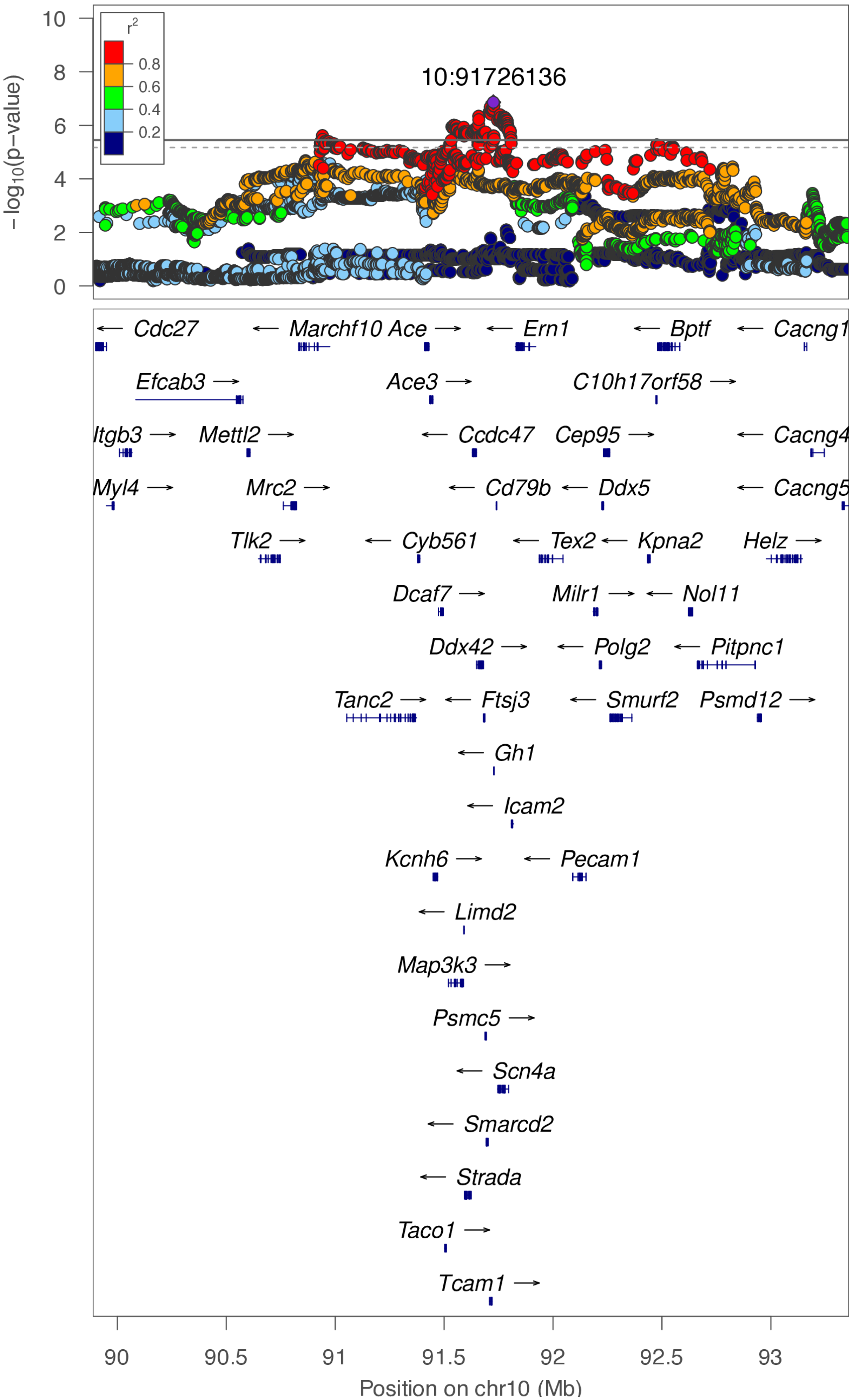

**chr10:91.5 Mb - METAL - broad**

Lead: 10:91,726,136 | Genes in QTL interval: 37

QTL interval: chr10:90,586,669-92,719,932

Displayed region: chr10:87,088,192-96,157,889

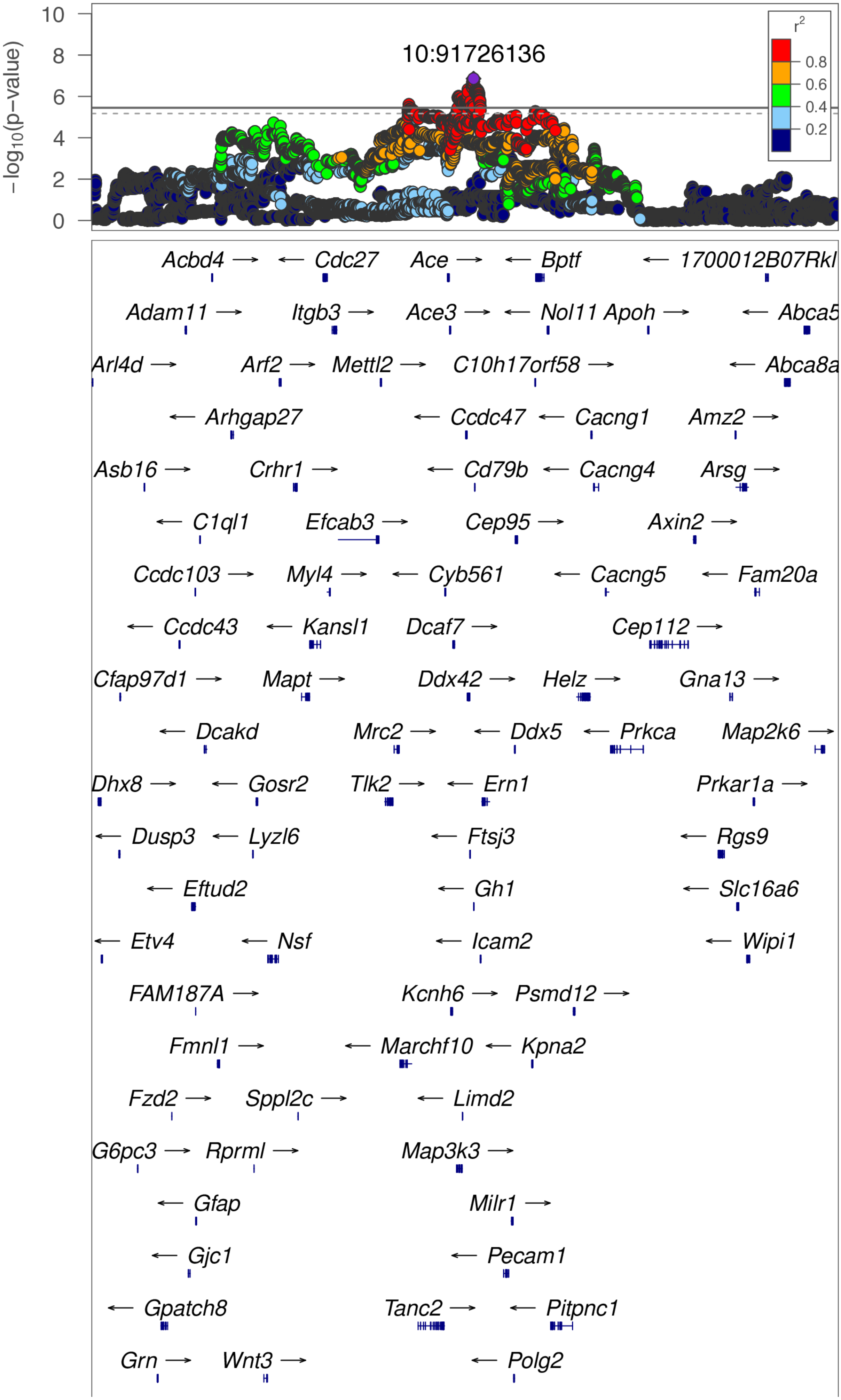

**chr10:91.5 Mb - METAL - broad (continued)**

Lead: 10:91,726,136 | Genes in QTL interval: 37

QTL interval: chr10:90,586,669-92,719,932

Displayed region: chr10:87,088,192-96,157,889

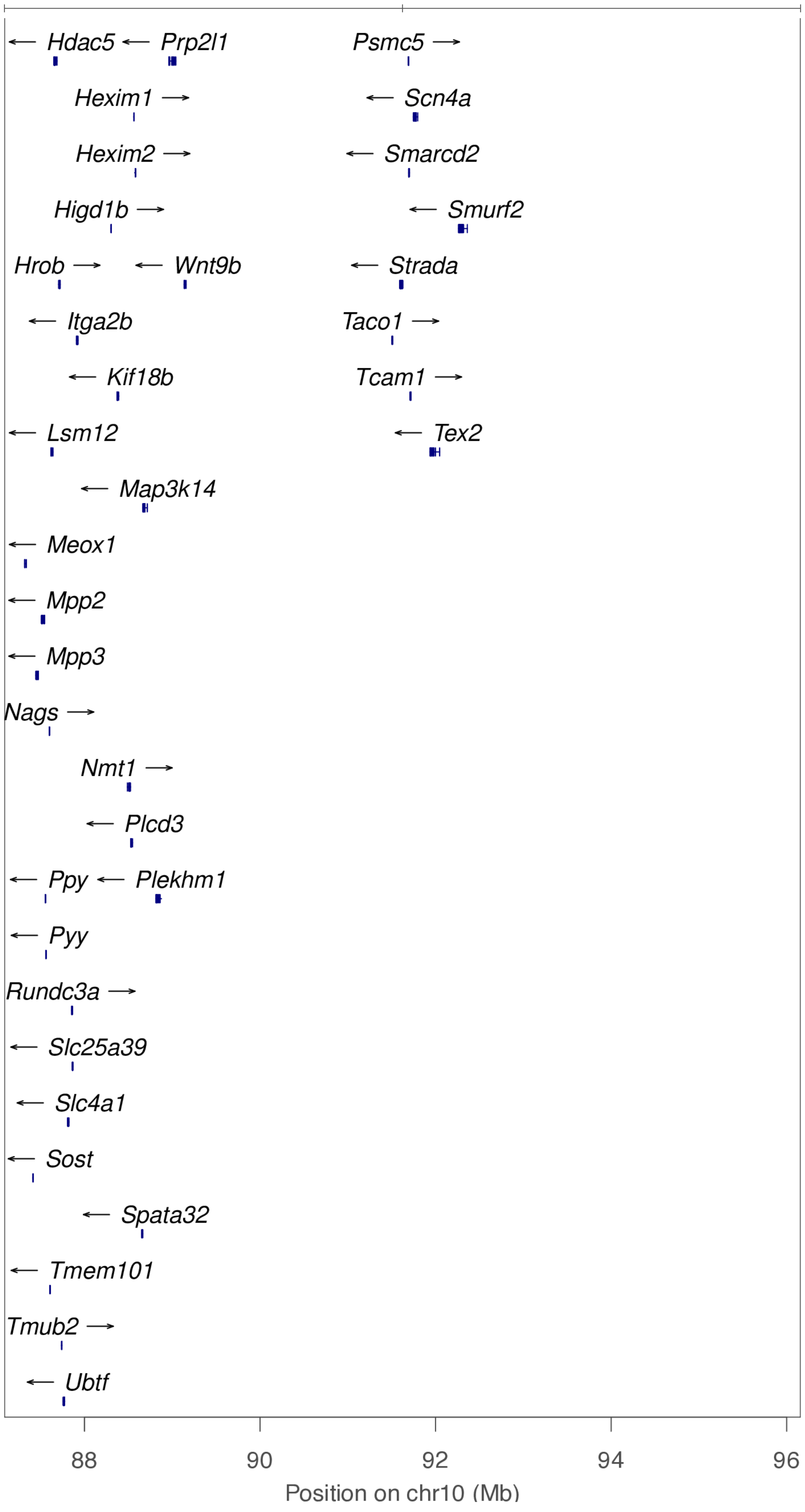

**chr10:92.1 Mb - Genomic SEM - narrow**

Lead: 10:92,130,822 | Genes in QTL interval: 14

QTL interval: chr10:92,120,303-92,987,782

Displayed region: chr10:91,920,303-94,593,220

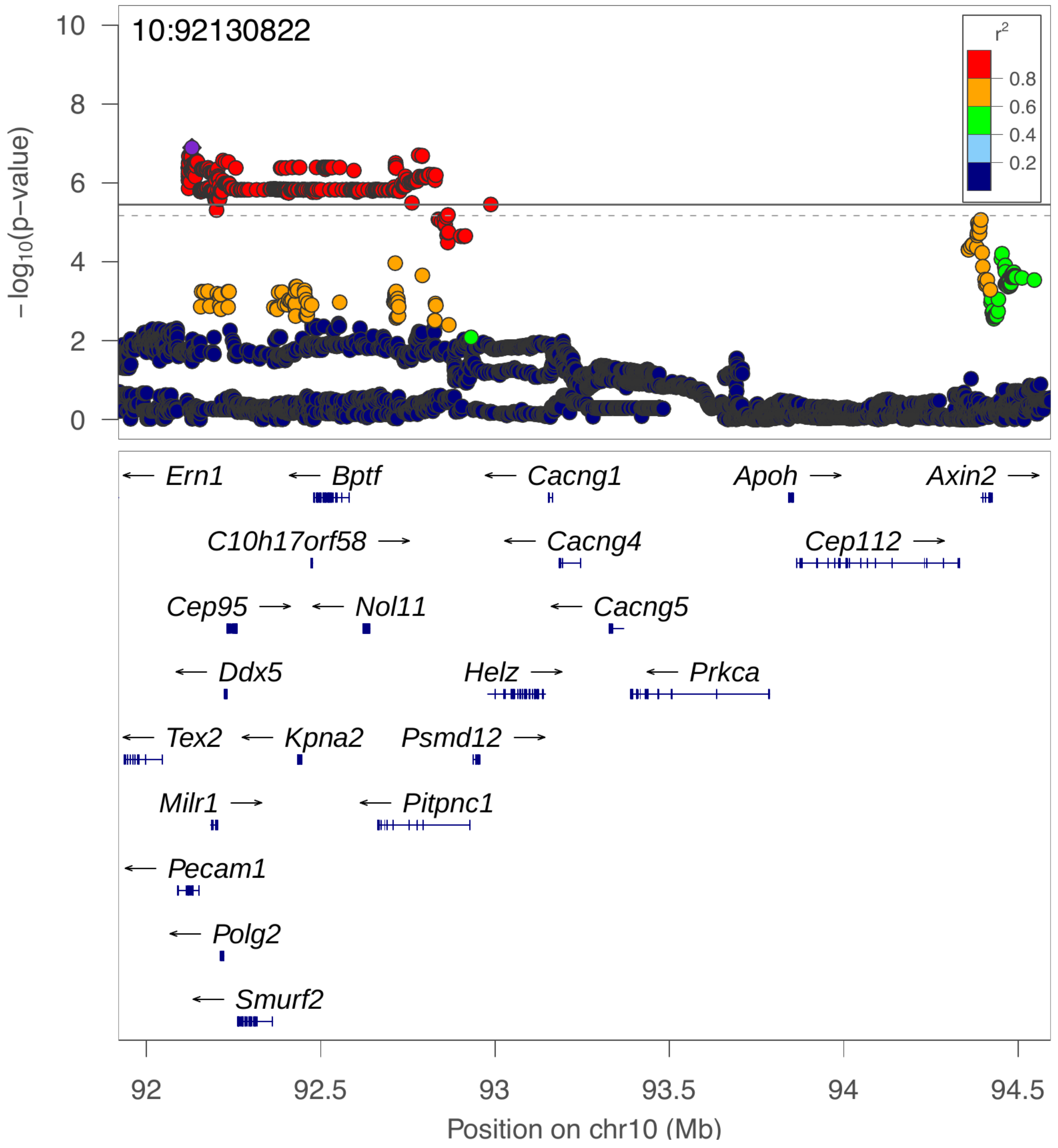

**chr10:92.1 Mb - Genomic SEM - broad**

Lead: 10:92,130,822 | Genes in QTL interval: 14

QTL interval: chr10:92,120,303-92,987,782

Displayed region: chr10:89,120,303-97,393,220

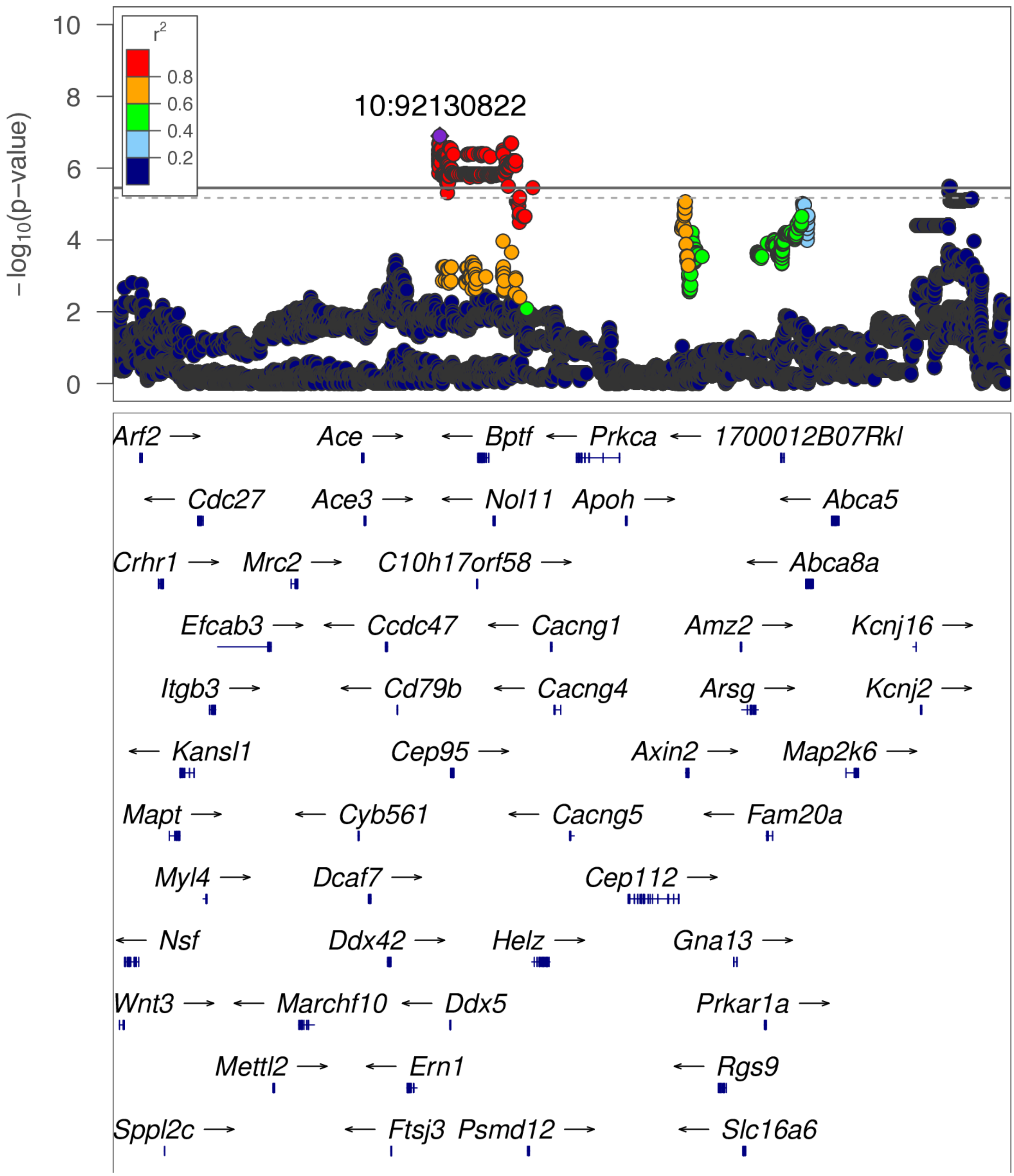

**chr10:92.1 Mb - Genomic SEM - broad (continued)**

Lead: 10:92,130,822 | Genes in QTL interval: 14

QTL interval: chr10:92,120,303-92,987,782

Displayed region: chr10:89,120,303-97,393,220

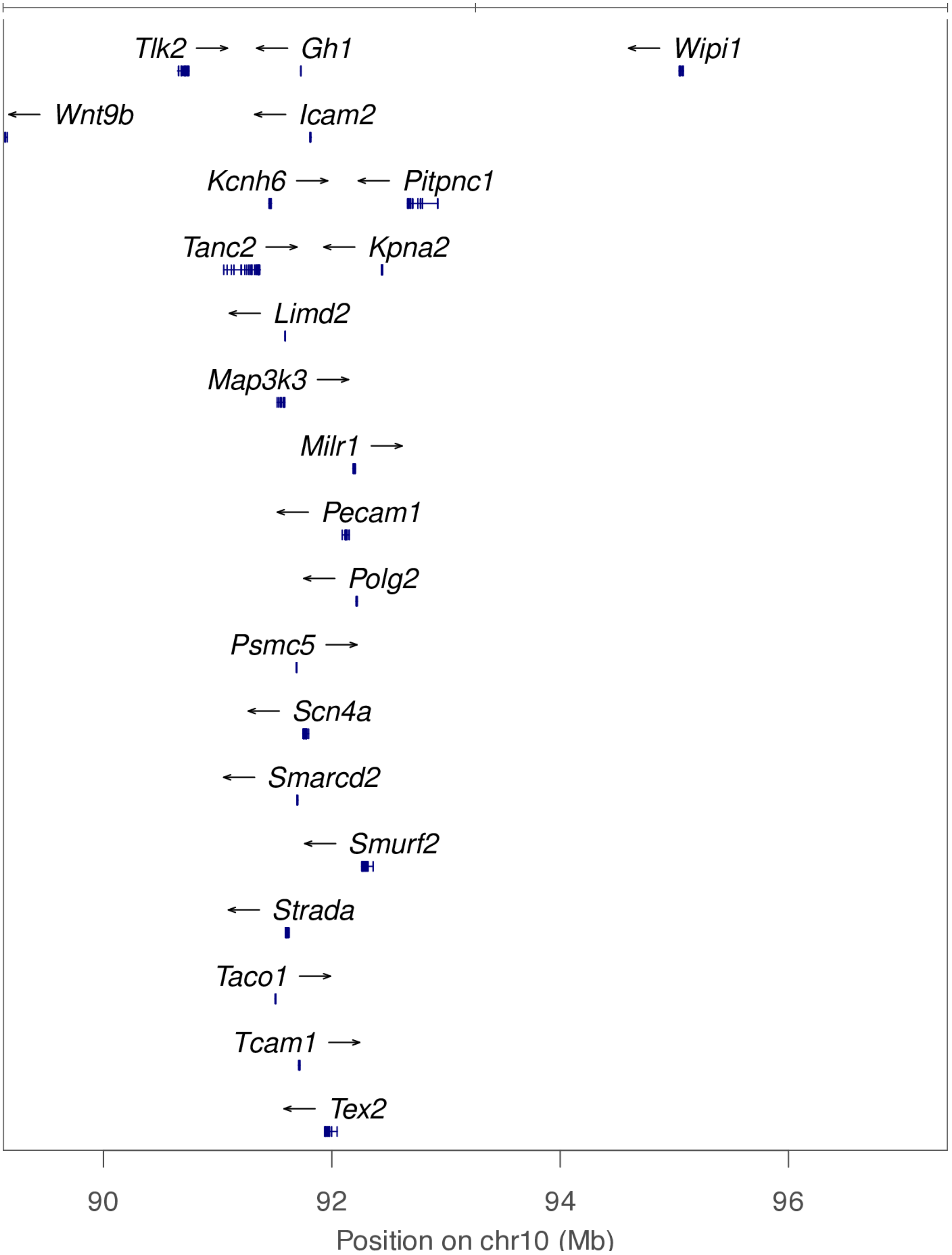

**chr13:44.7 Mb - Genomic SEM - narrow**

Lead: 13:44,662,145 | Genes in QTL interval: 21

QTL interval: chr13:44,496,272-45,668,431

Displayed region: chr13:44,138,510-45,868,431

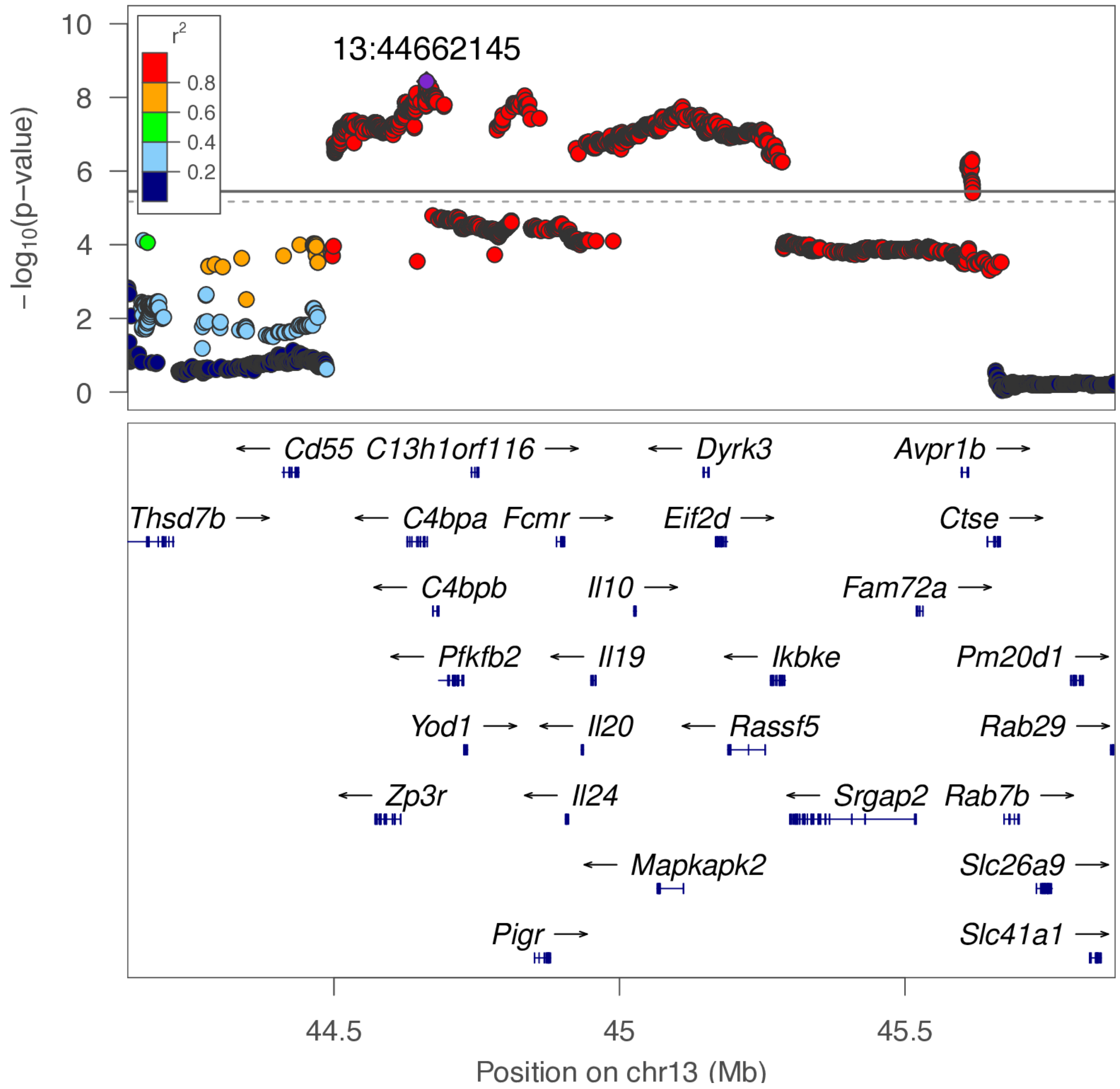

**chr13:44.7 Mb - Genomic SEM - broad**

Lead: 13:44,662,145 | Genes in QTL interval: 21

QTL interval: chr13:44,496,272-45,668,431

Displayed region: chr13:41,338,510-48,668,431

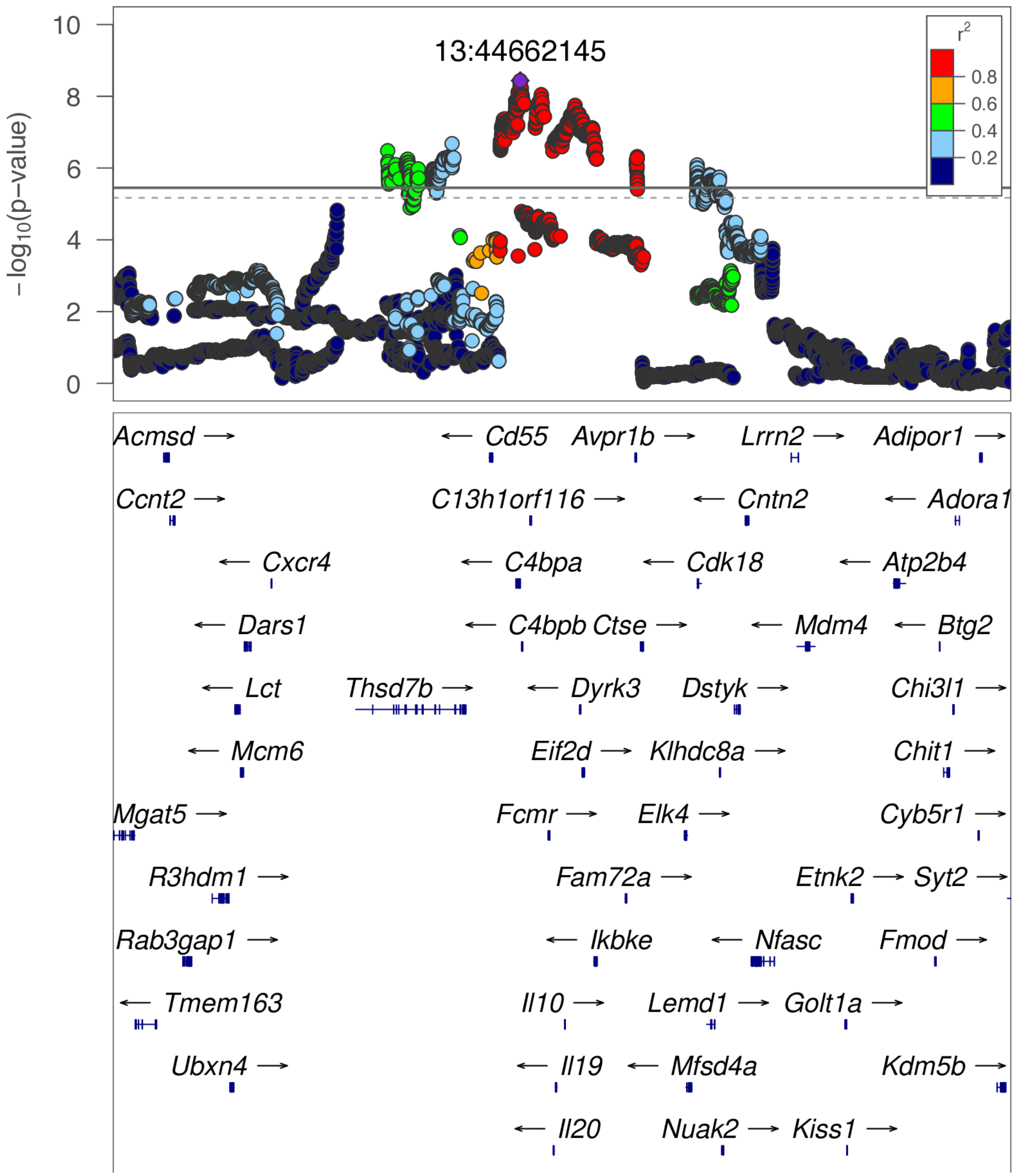

**chr13:44.7 Mb - Genomic SEM - broad (continued)**

Lead: 13:44,662,145 | Genes in QTL interval: 21

QTL interval: chr13:44,496,272-45,668,431

Displayed region: chr13:41,338,510-48,668,431

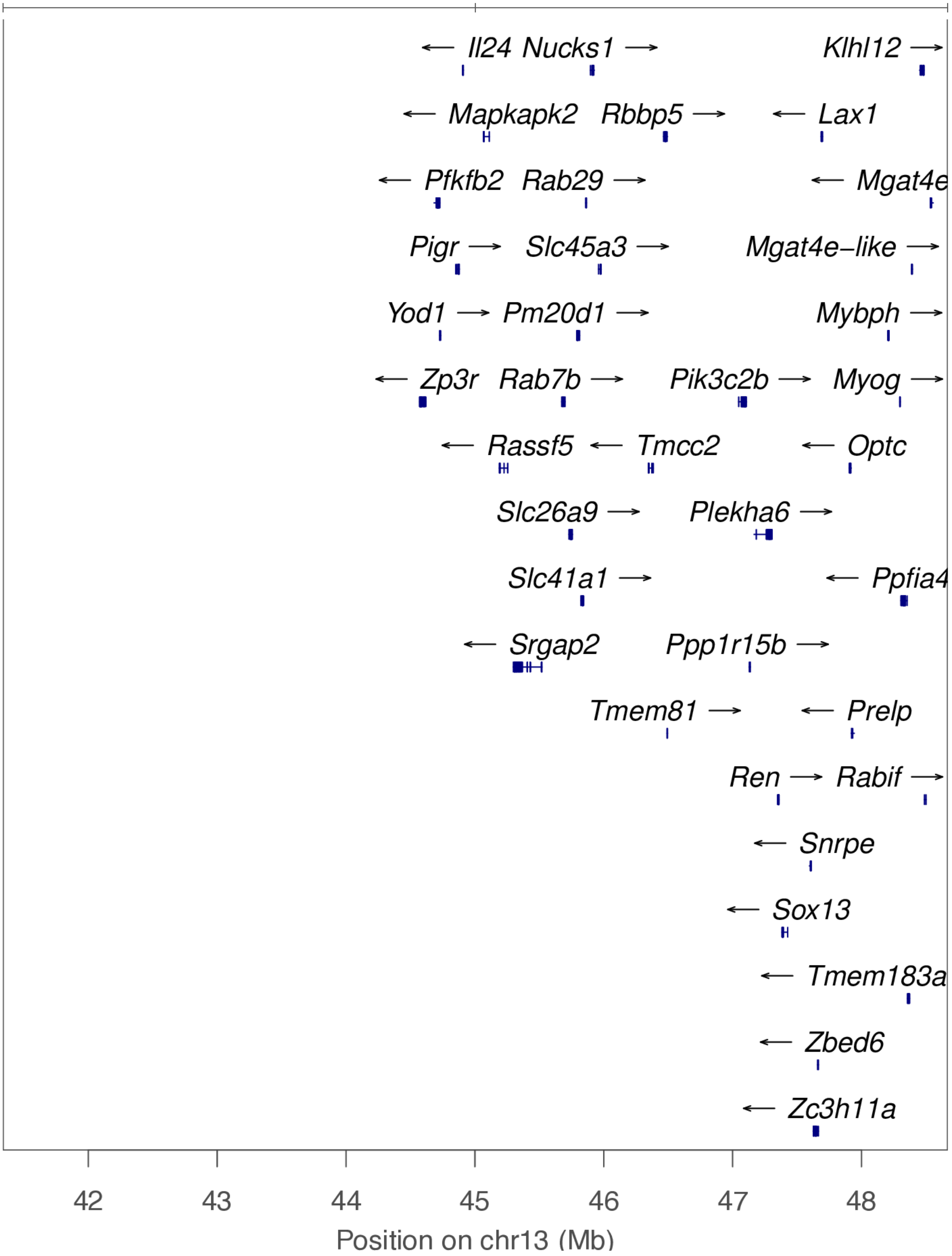

**chr13:50.8 Mb - Genomic SEM - narrow**

Lead: 13:50,762,238 | Genes in QTL interval: 3

QTL interval: chr13:50,576,435-50,921,211

Displayed region: chr13:50,376,435-51,158,892

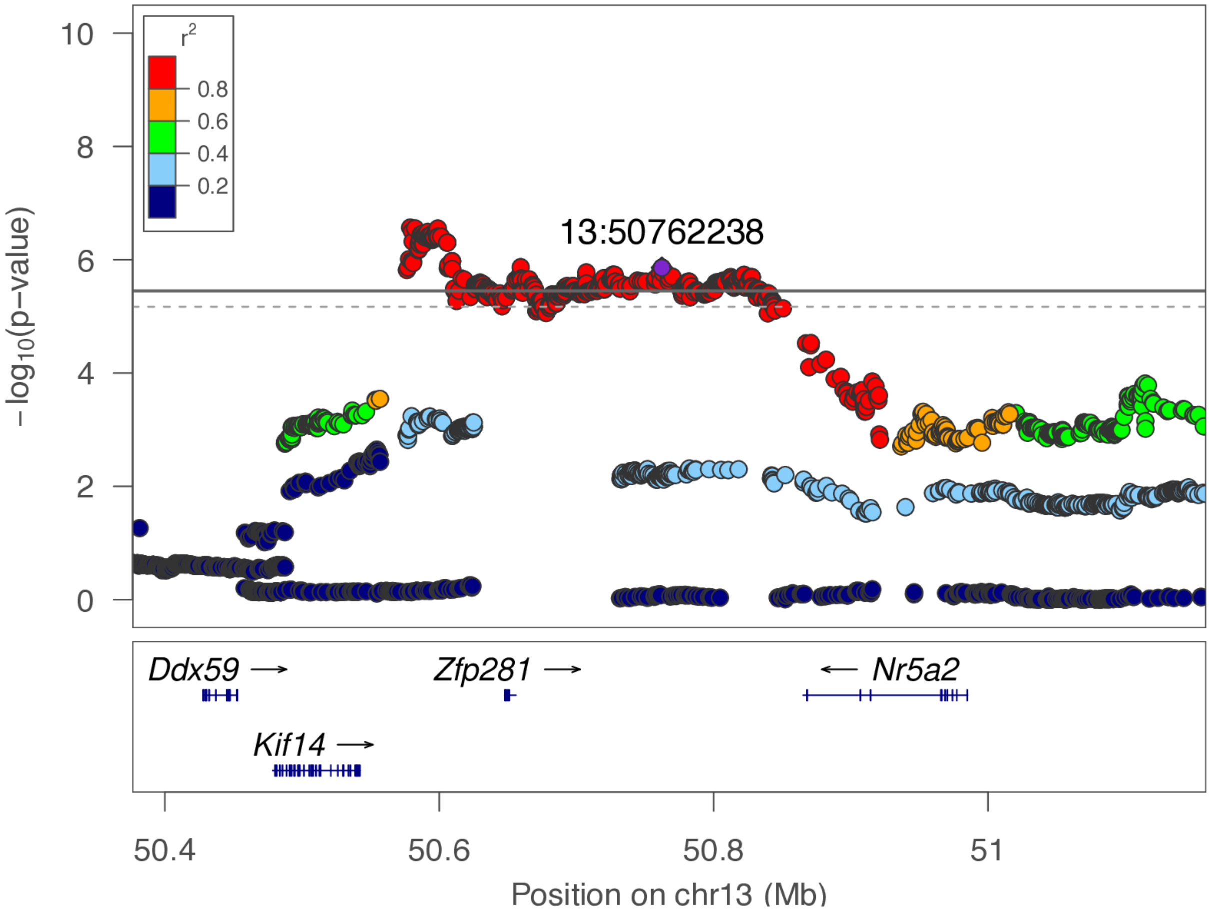

**chr13:50.8 Mb - Genomic SEM - broad**

Lead: 13:50,762,238 | Genes in QTL interval: 3

QTL interval: chr13:50,576,435-50,921,211

Displayed region: chr13:47,576,435-53,958,892

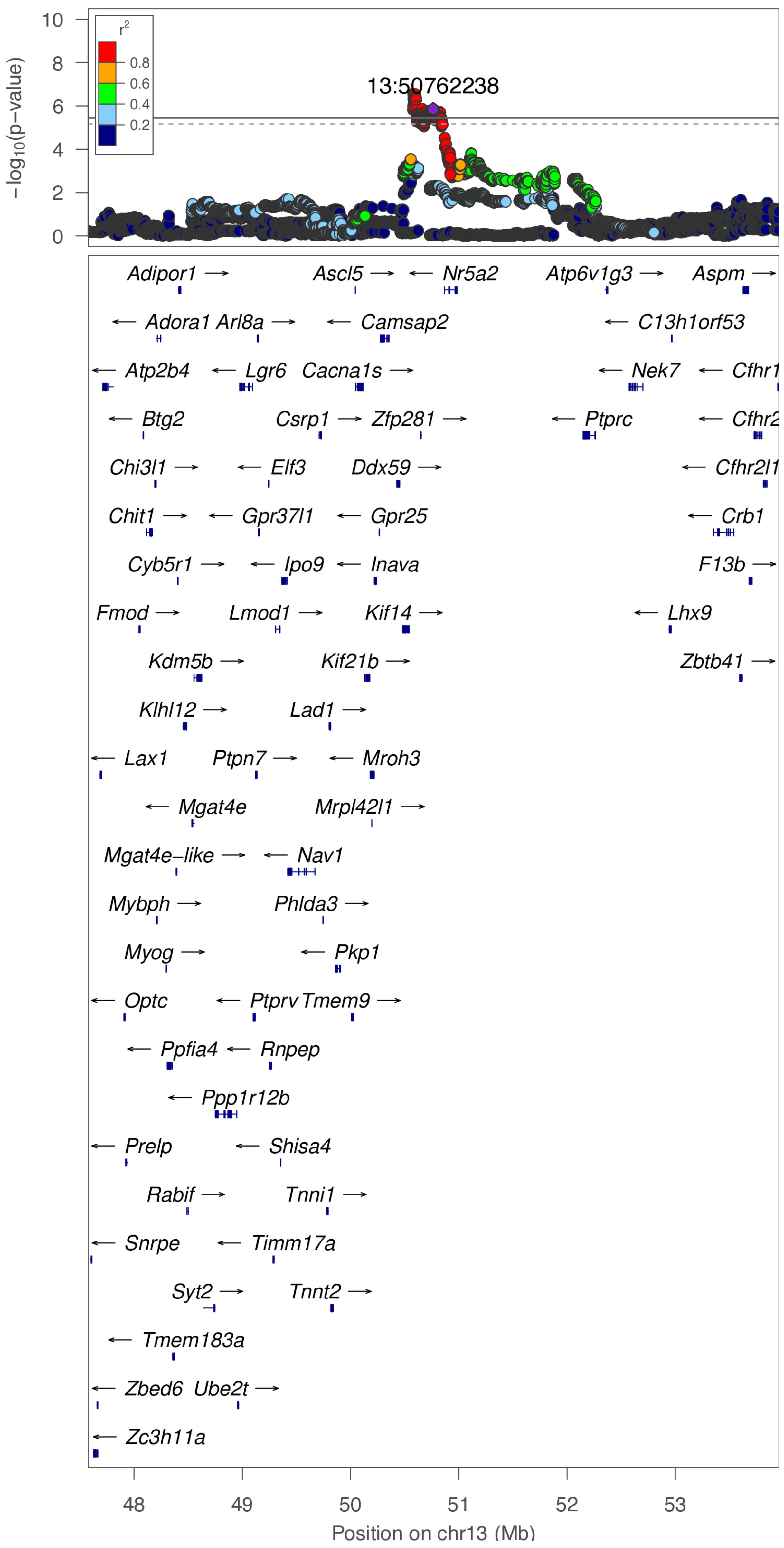

**chr14:74.7 Mb - Genomic SEM - narrow**

Lead: 14:74,727,310 | Genes in QTL interval: 1

QTL interval: chr14:74,415,936-75,722,776

Displayed region: chr14:74,215,936-76,784,922

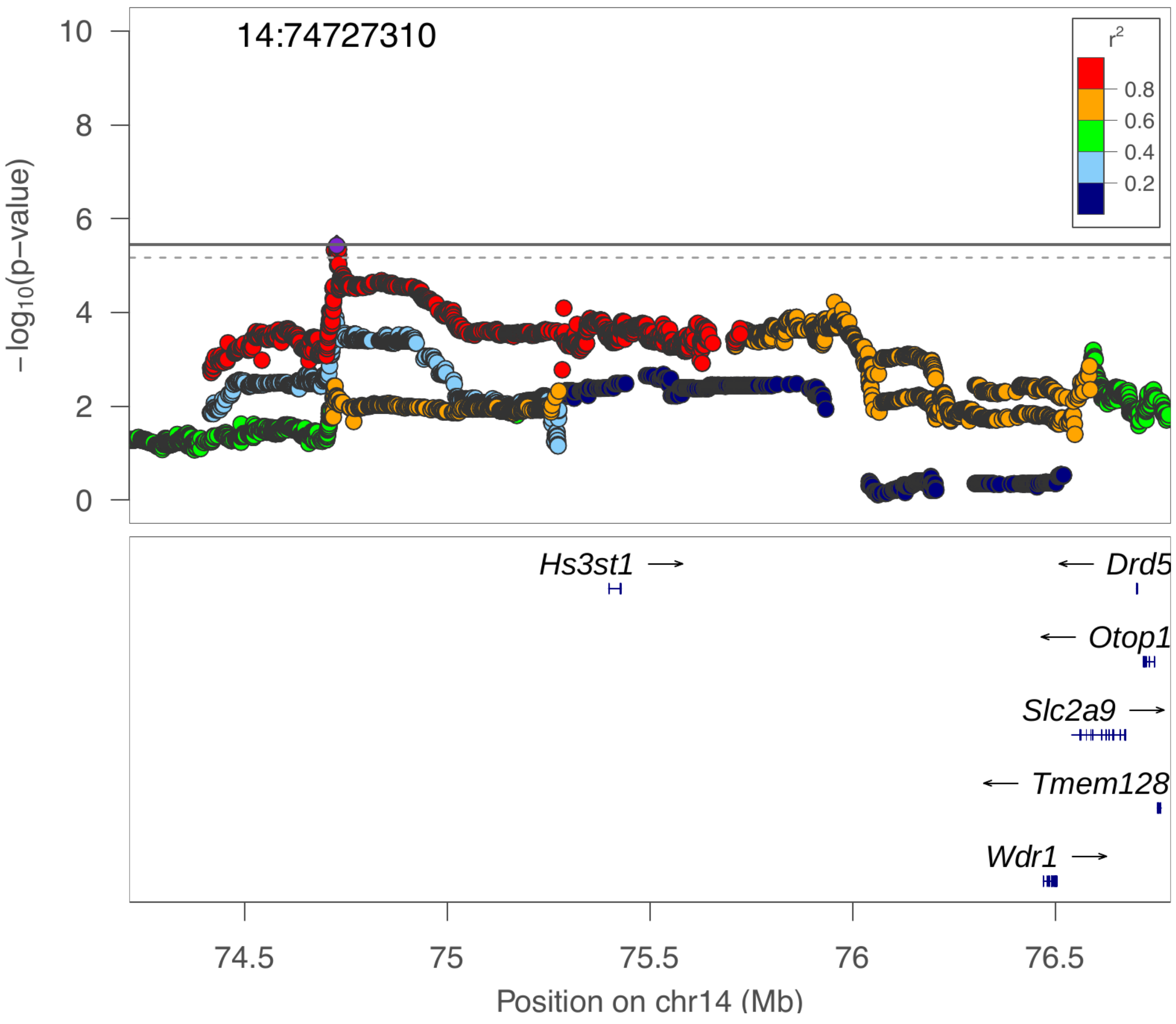

**chr14:74.7 Mb - Genomic SEM - broad**

Lead: 14:74,727,310 | Genes in QTL interval: 1

QTL interval: chr14:74,415,936-75,722,776

Displayed region: chr14:71,415,936-79,584,922

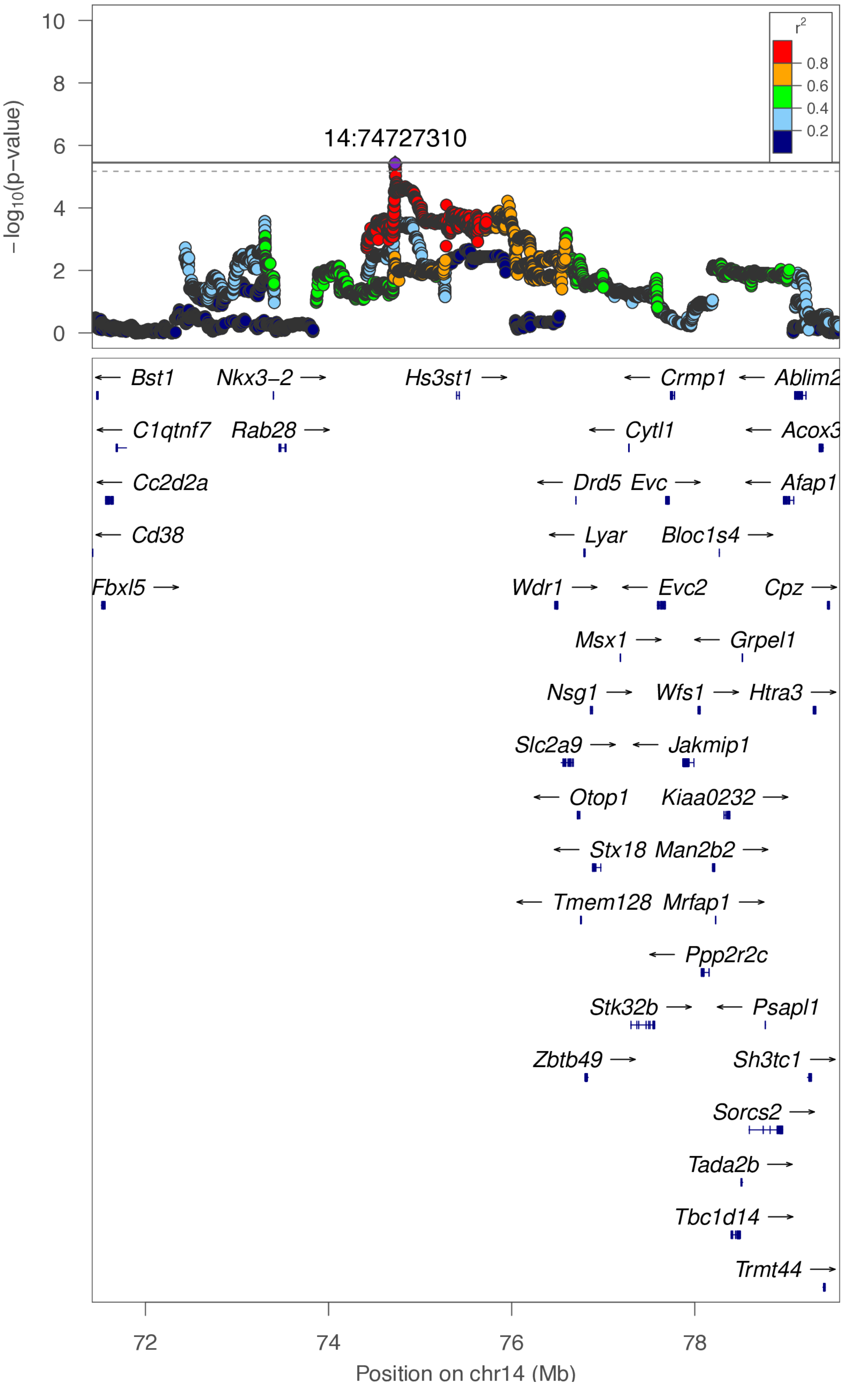

**chr14:105.7 Mb - MEGA - narrow**

Lead: 14:105,642,007 | Genes in QTL interval: 0

QTL interval: chr14:105,537,323-105,732,122

Displayed region: chr14:105,337,323-105,945,228

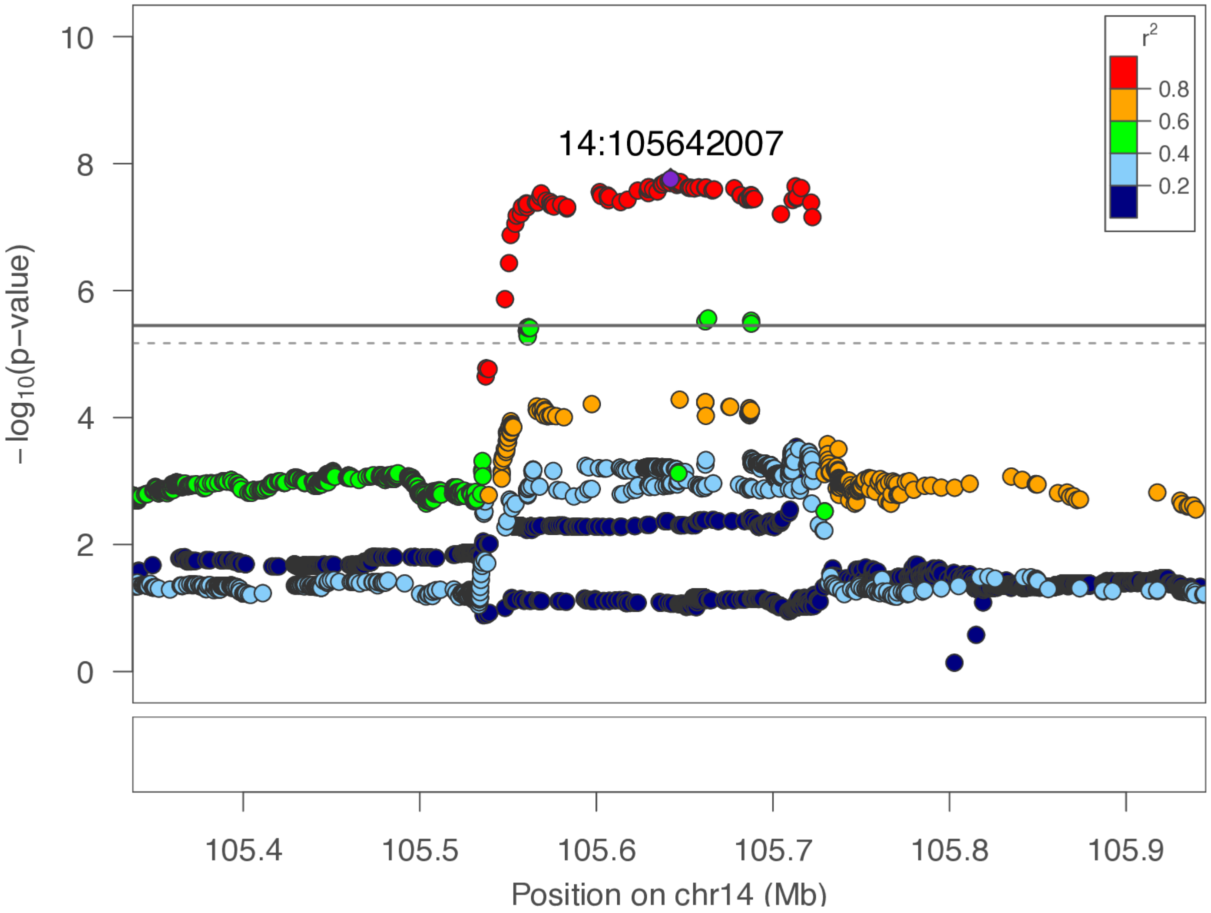

**chr14:105.7 Mb - MEGA - broad**

Lead: 14:105,642,007 | Genes in QTL interval: 0

QTL interval: chr14:105,537,323-105,732,122

Displayed region: chr14:102,537,323-108,745,228

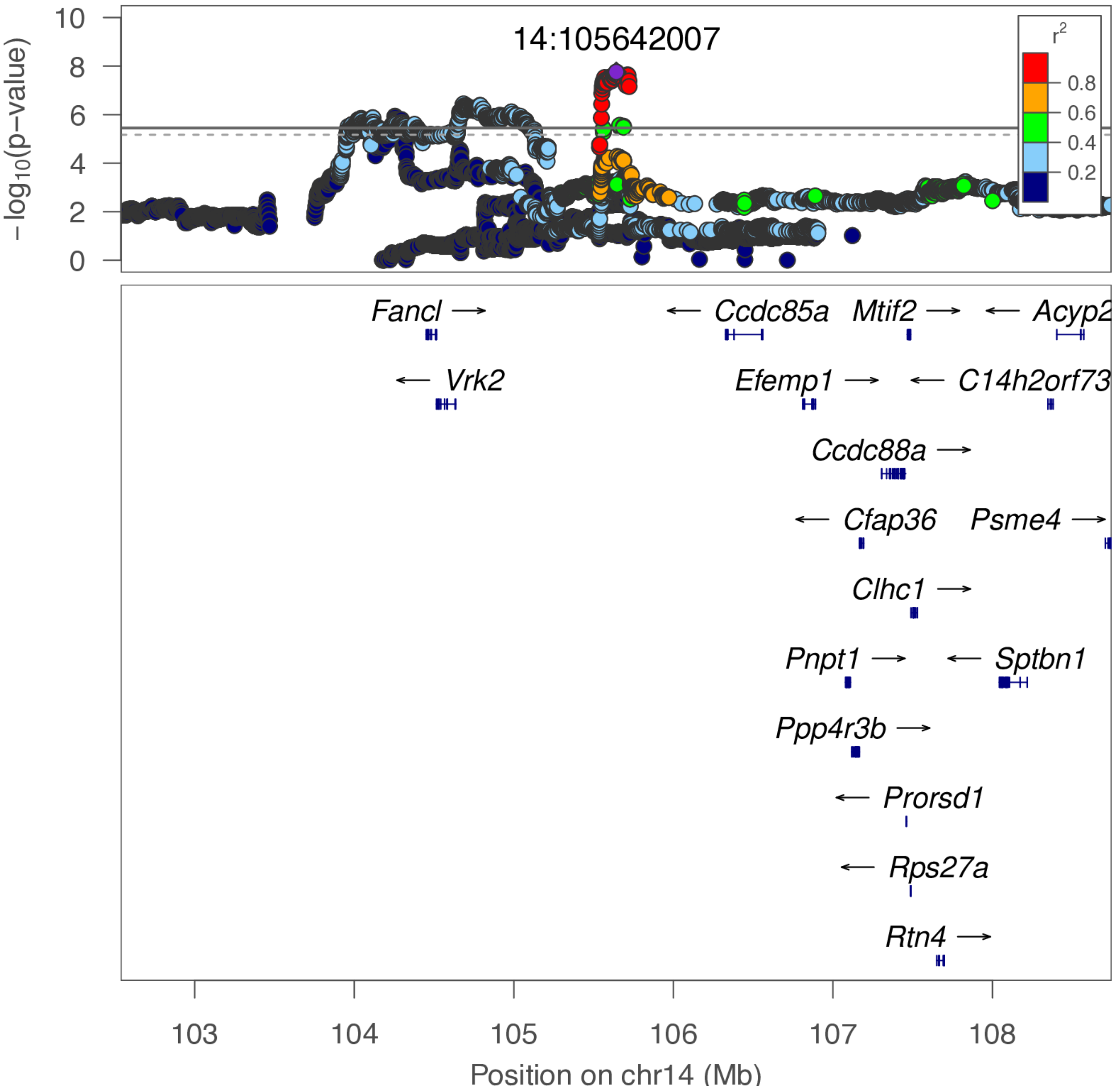

**chr14:105.7 Mb - METAL - narrow**

Lead: 14:105,642,007 | Genes in QTL interval: 0

QTL interval: chr14:105,537,323-105,732,122

Displayed region: chr14:105,337,323-105,977,198

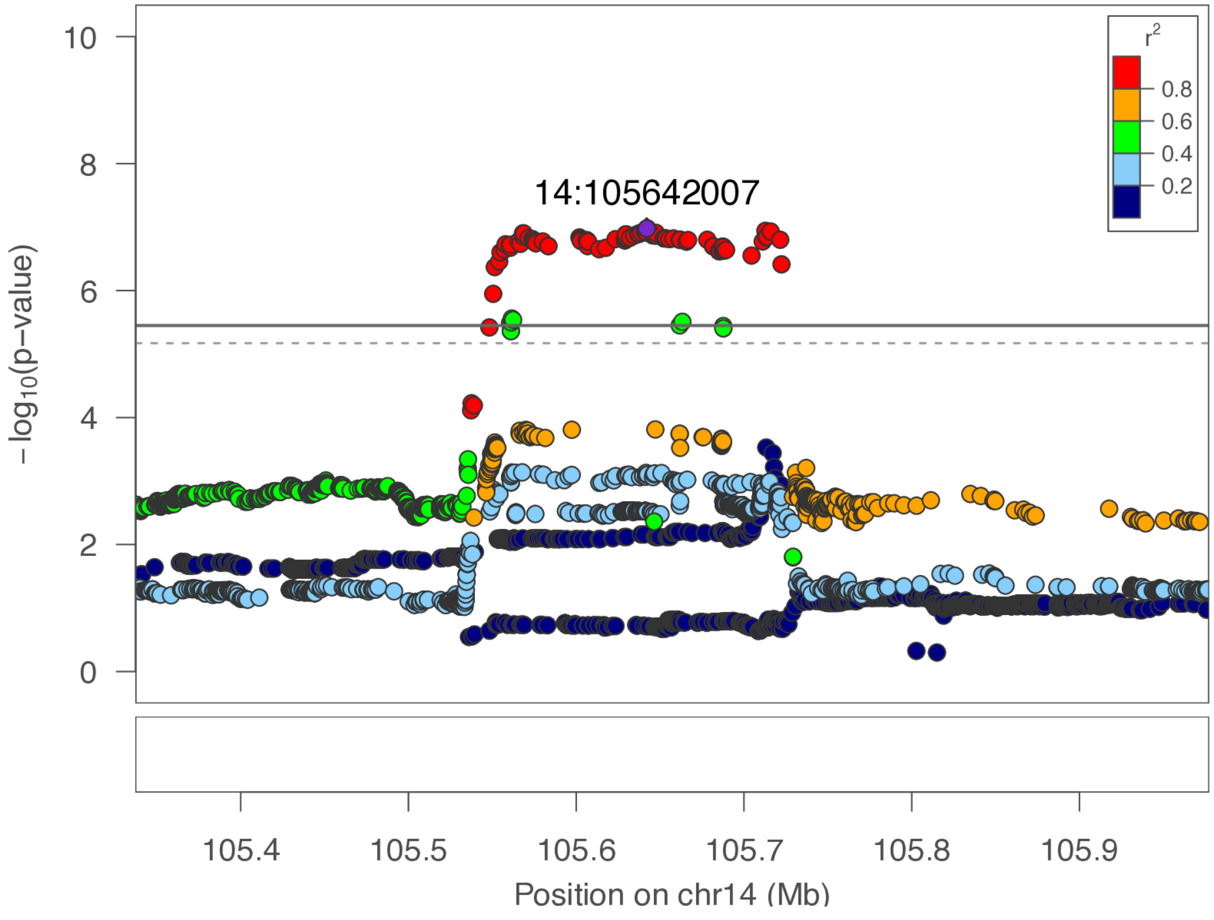

**chr14:105.7 Mb - METAL - broad**

Lead: 14:105,642,007 | Genes in QTL interval: 0

QTL interval: chr14:105,537,323-105,732,122

Displayed region: chr14:102,537,323-108,777,198

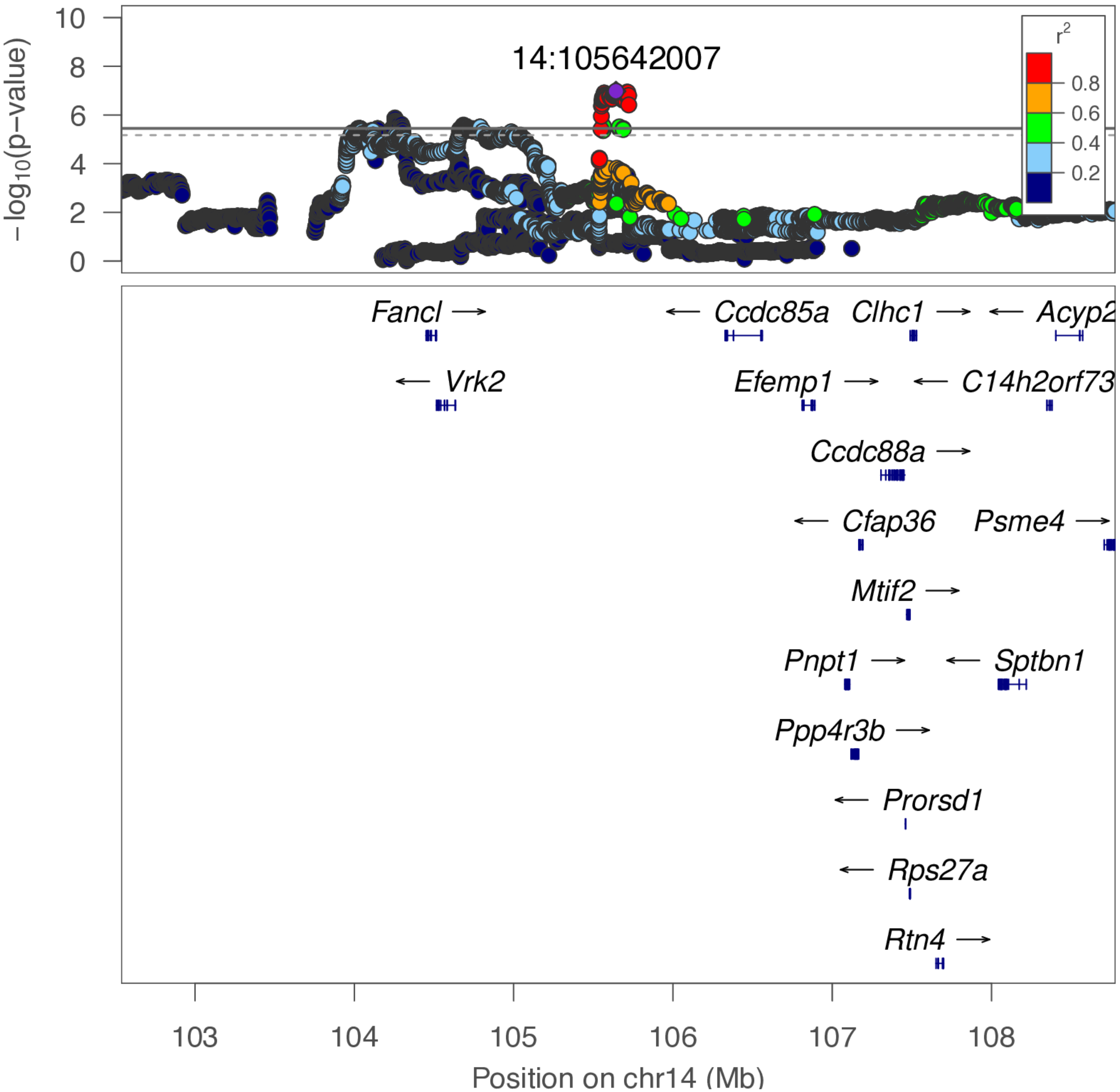

**chr14:105.7 Mb - Genomic SEM - narrow**

Lead: 14:105,721,596 | Genes in QTL interval: 0

QTL interval: chr14:105,537,323-105,732,122

Displayed region: chr14:105,337,323-106,171,559

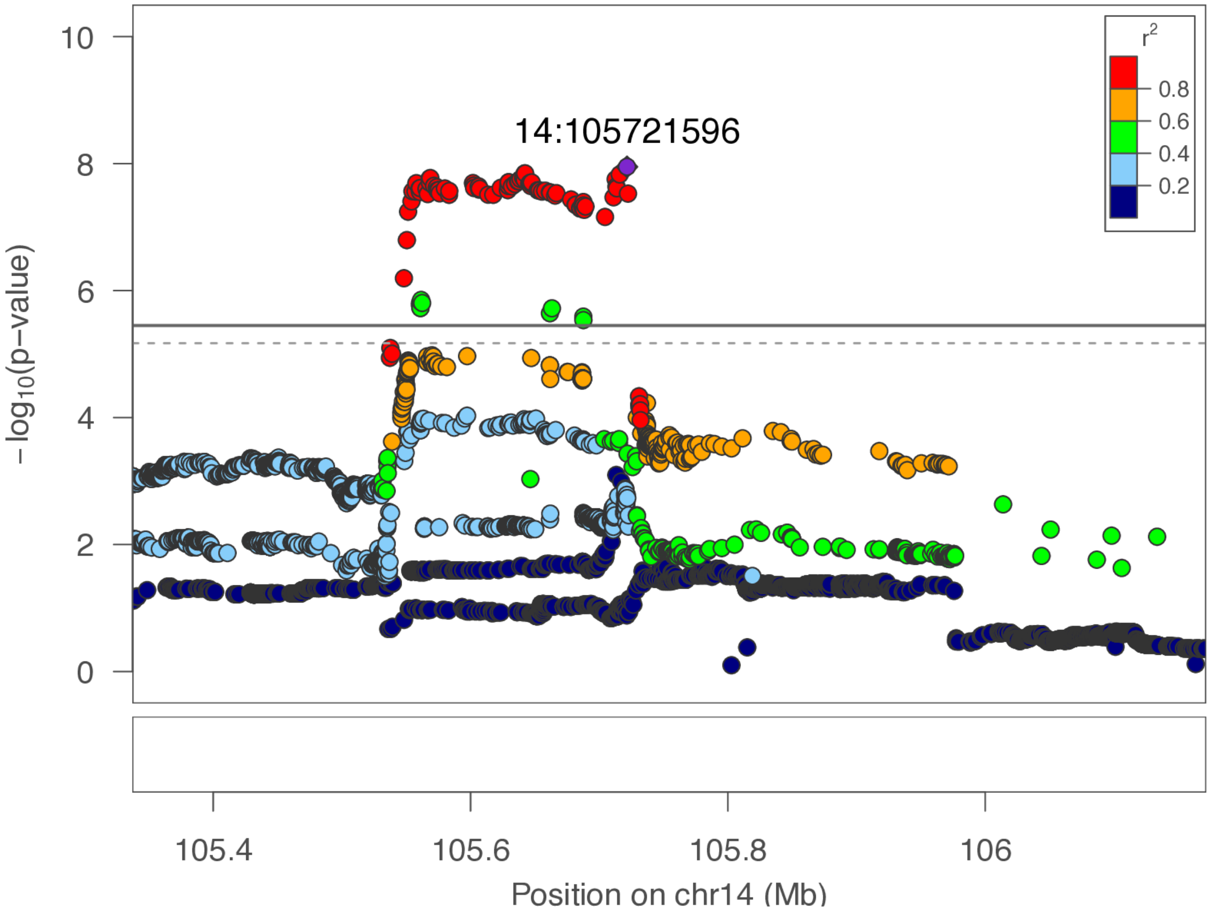

**chr14:105.7 Mb - Genomic SEM - broad**

Lead: 14:105,721,596 | Genes in QTL interval: 0

QTL interval: chr14:105,537,323-105,732,122

Displayed region: chr14:102,537,323-108,971,559

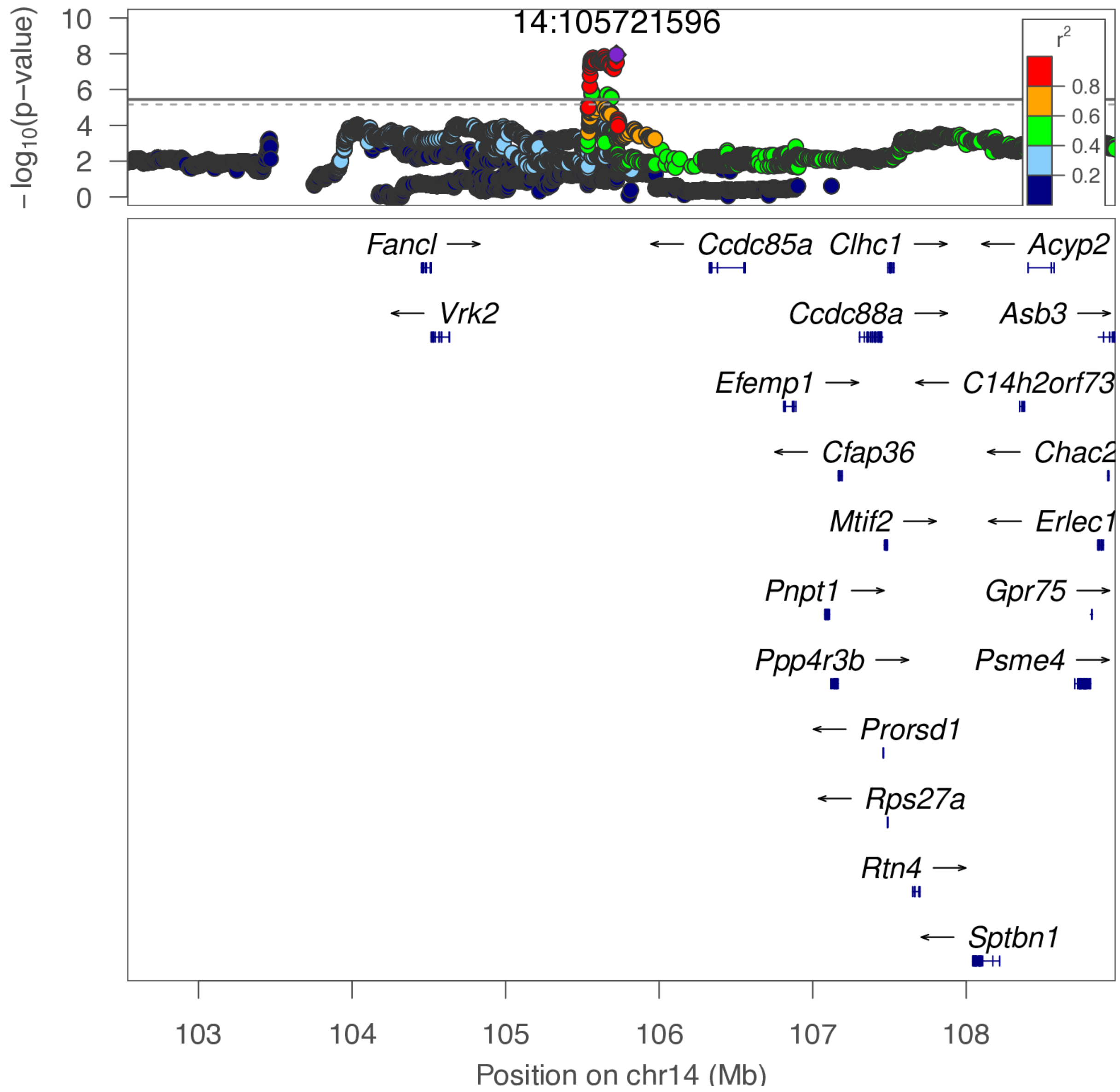

**chr15:77.5 Mb - Genomic SEM - narrow**

Lead: 15:77,491,635 | Genes in QTL interval: 0

QTL interval: chr15:76,700,151-78,264,979

Displayed region: chr15:76,463,265-78,464,979

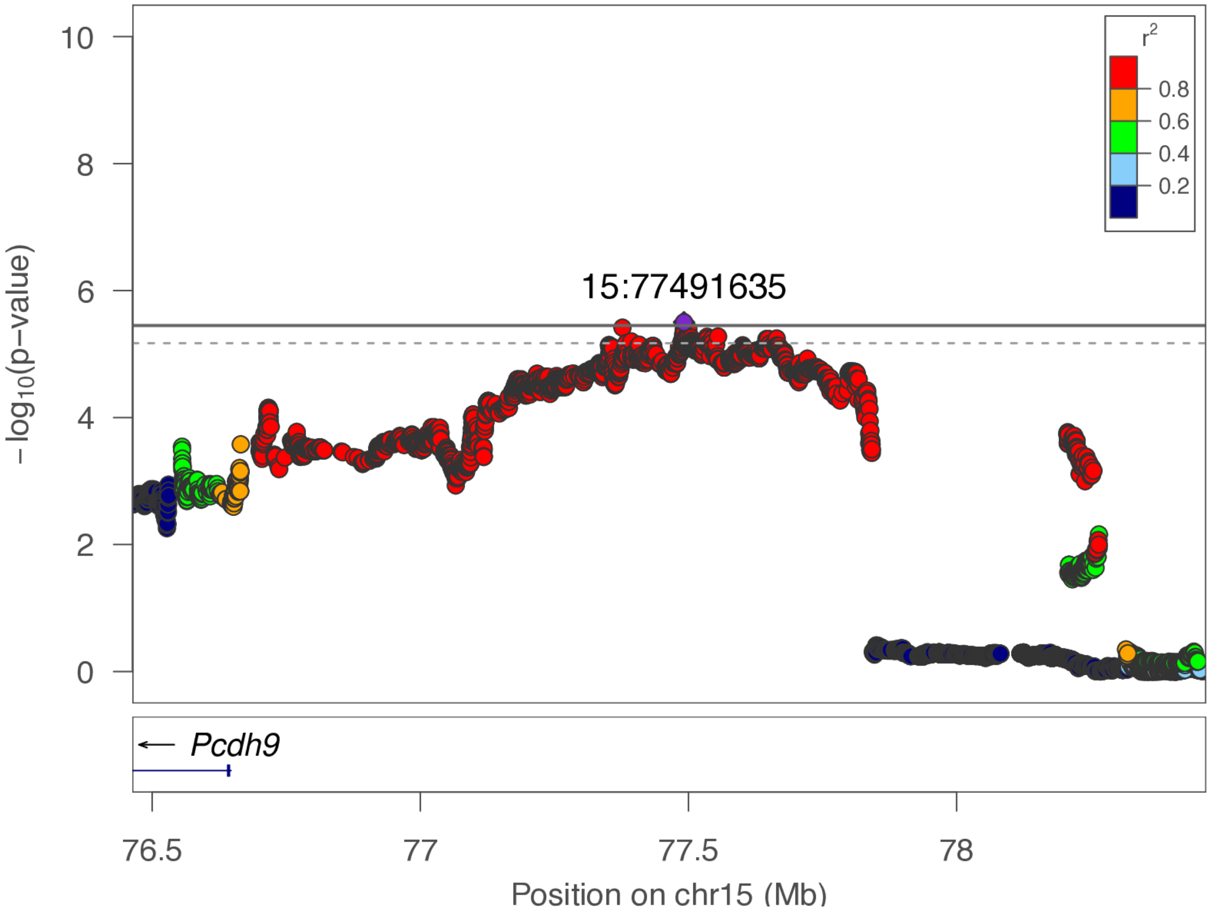

**chr15:77.5 Mb - Genomic SEM - broad**

Lead: 15:77,491,635 | Genes in QTL interval: 0

QTL interval: chr15:76,700,151-78,264,979

Displayed region: chr15:73,663,265-81,264,979

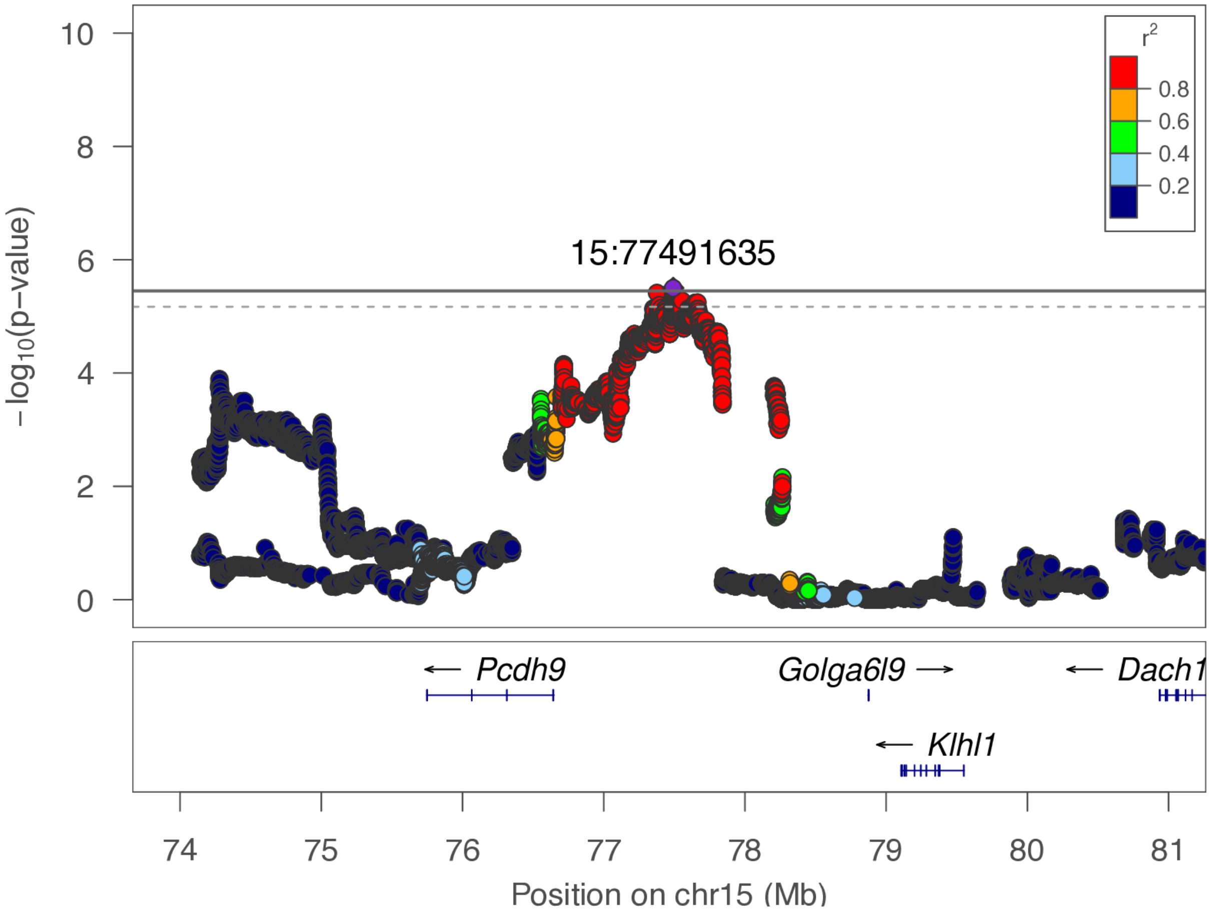

**chr17:17.1 Mb - Genomic SEM - narrow**

Lead: 17:17,106,624 | Genes in QTL interval: 1

QTL interval: chr17:16,554,912-17,272,514

Displayed region: chr17:15,960,825-17,645,717

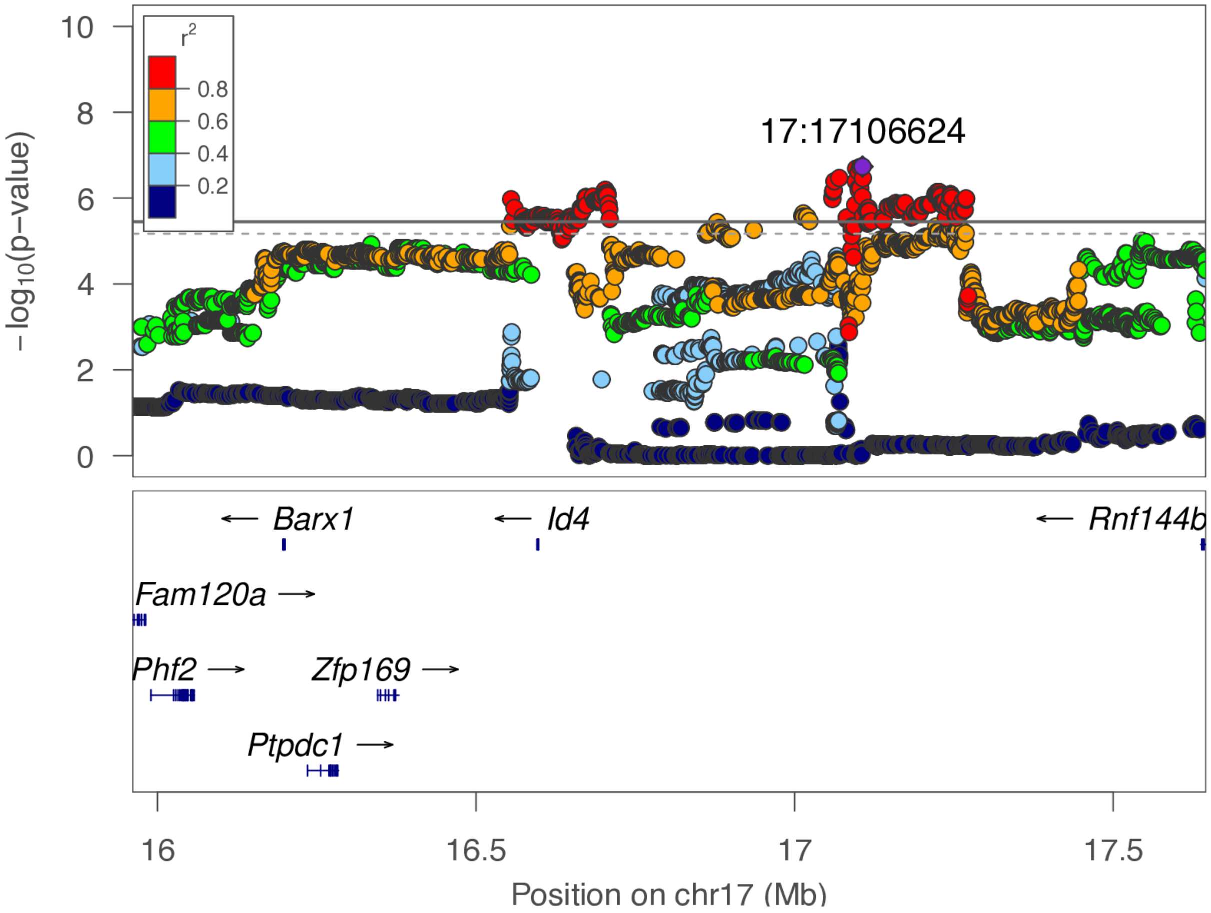

**chr17:17.1 Mb - Genomic SEM - broad**

Lead: 17:17,106,624 | Genes in QTL interval: 1

QTL interval: chr17:16,554,912-17,272,514

Displayed region: chr17:13,160,825-20,445,717

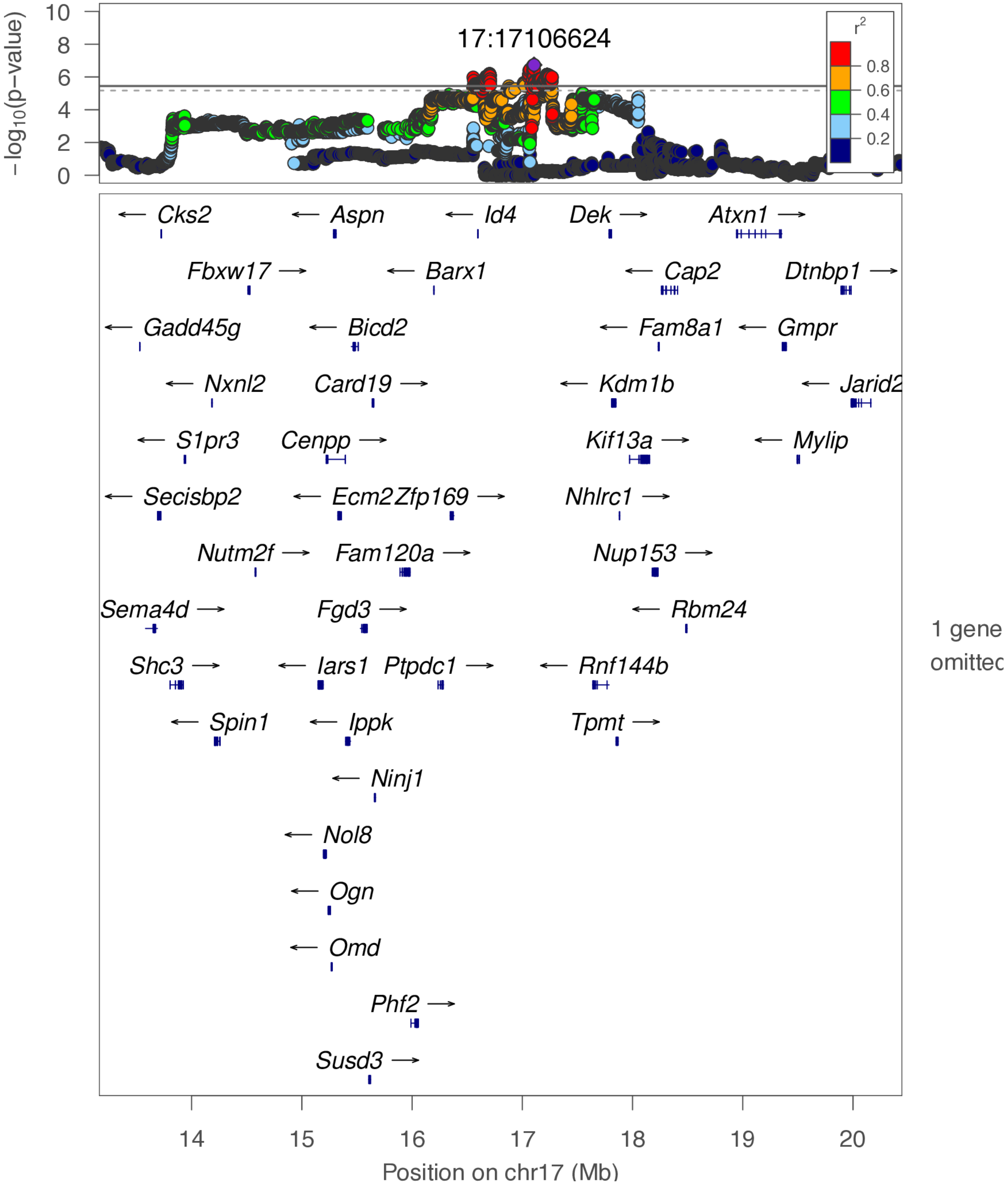

**chr18:76.7 Mb - Genomic SEM - narrow**

Lead: 18:76,732,183 | Genes in QTL interval: 2

QTL interval: chr18:76,403,146-77,077,249

Displayed region: chr18:76,180,337-77,457,164

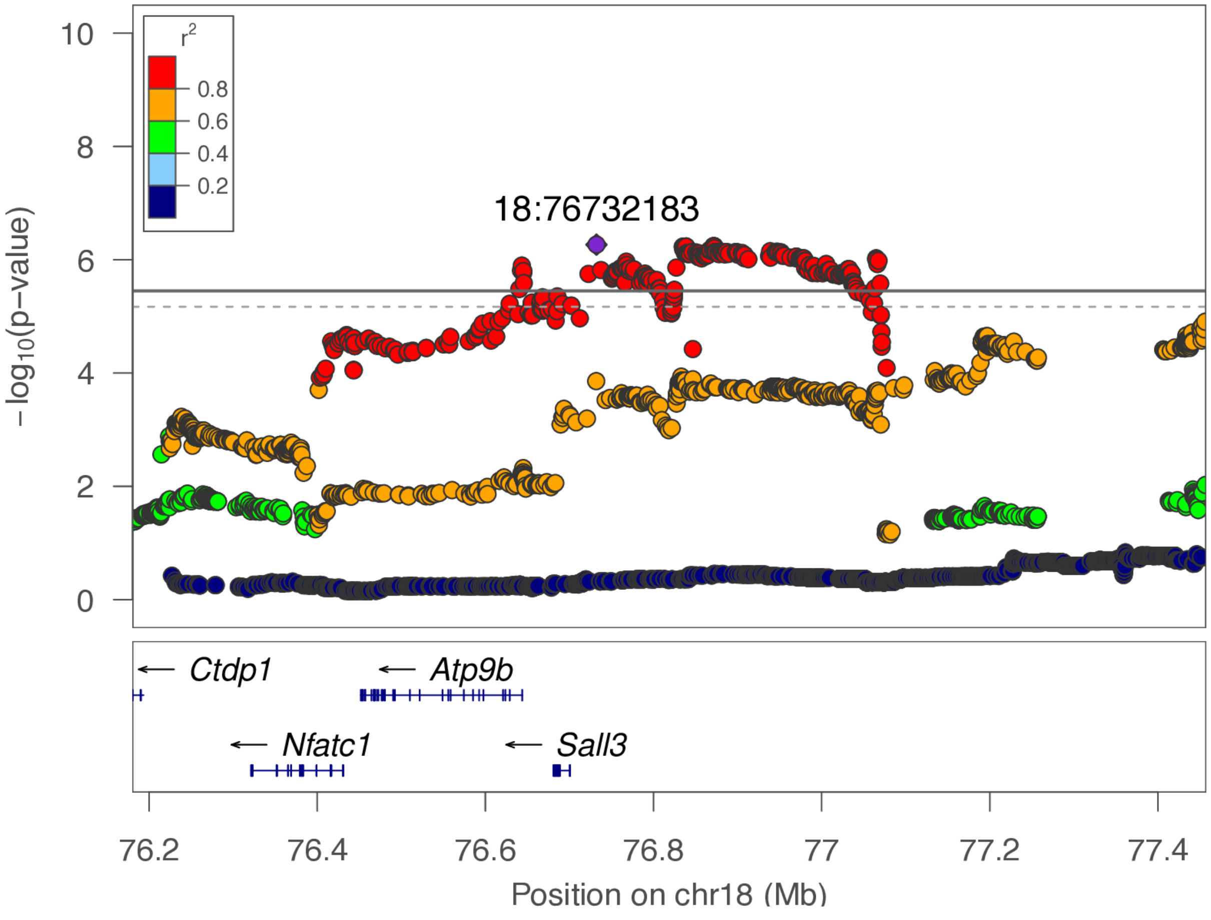

**chr18:76.7 Mb - Genomic SEM - broad**

Lead: 18:76,732,183 | Genes in QTL interval: 2

QTL interval: chr18:76,403,146-77,077,249

Displayed region: chr18:73,380,337-80,257,164

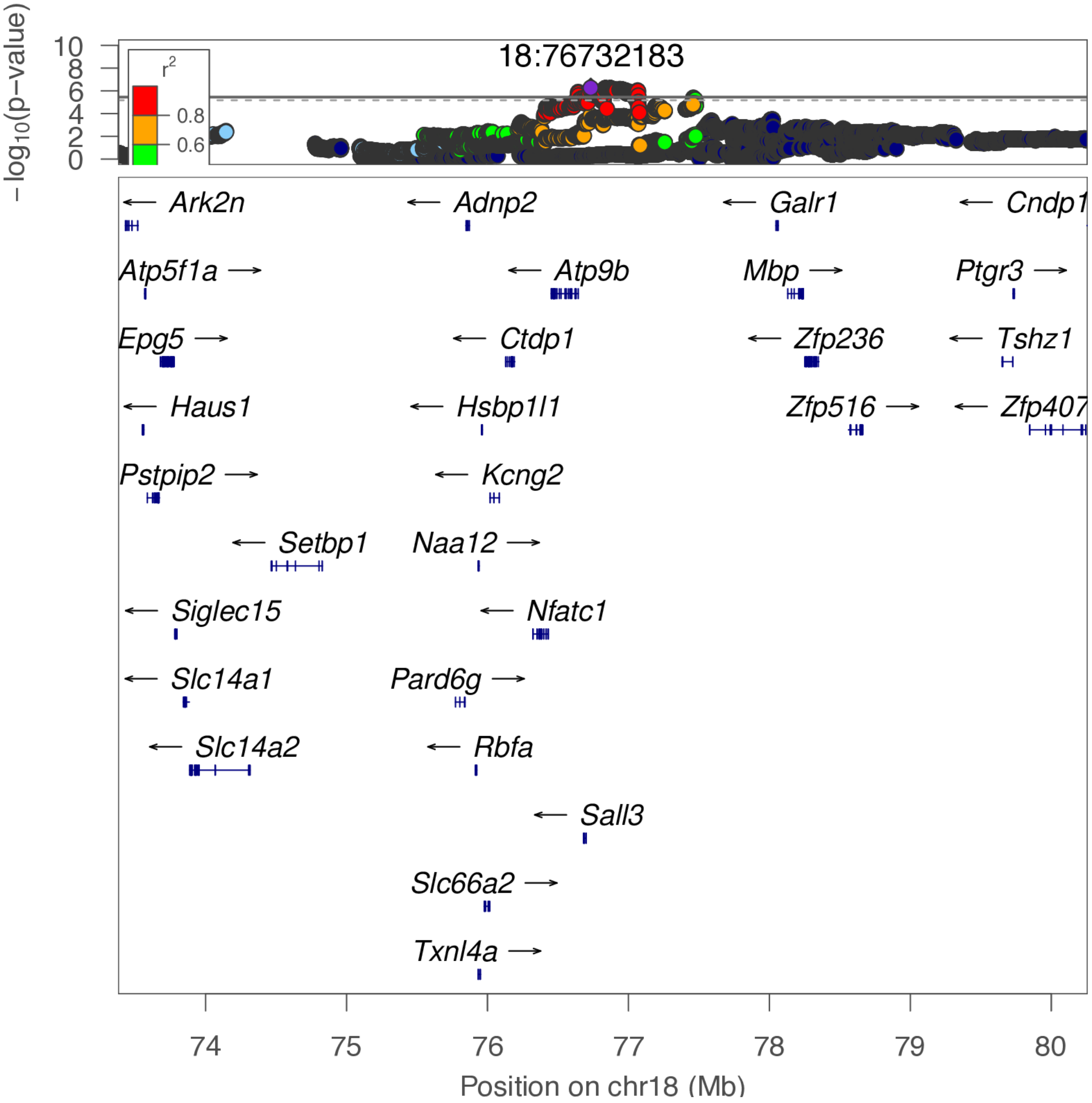

**chr19:54.5 Mb - Genomic SEM - narrow**

Lead: 19:54,543,879 | Genes in QTL interval: 9

QTL interval: chr19:54,045,028-54,654,195

Displayed region: chr19:53,845,028-54,854,195

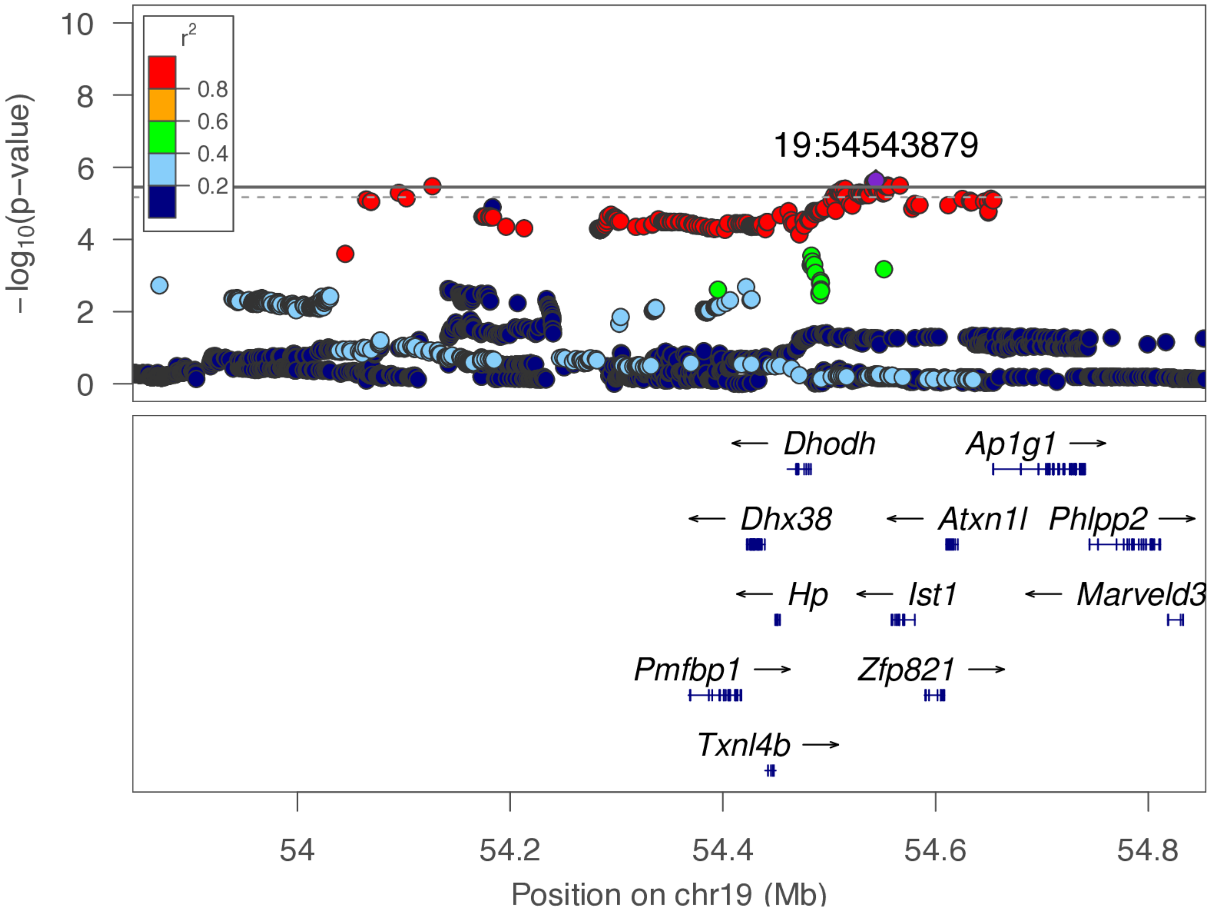

**chr19:54.5 Mb - Genomic SEM - broad**

Lead: 19:54,543,879 | Genes in QTL interval: 9

QTL interval: chr19:54,045,028-54,654,195

Displayed region: chr19:51,045,028-57,654,195

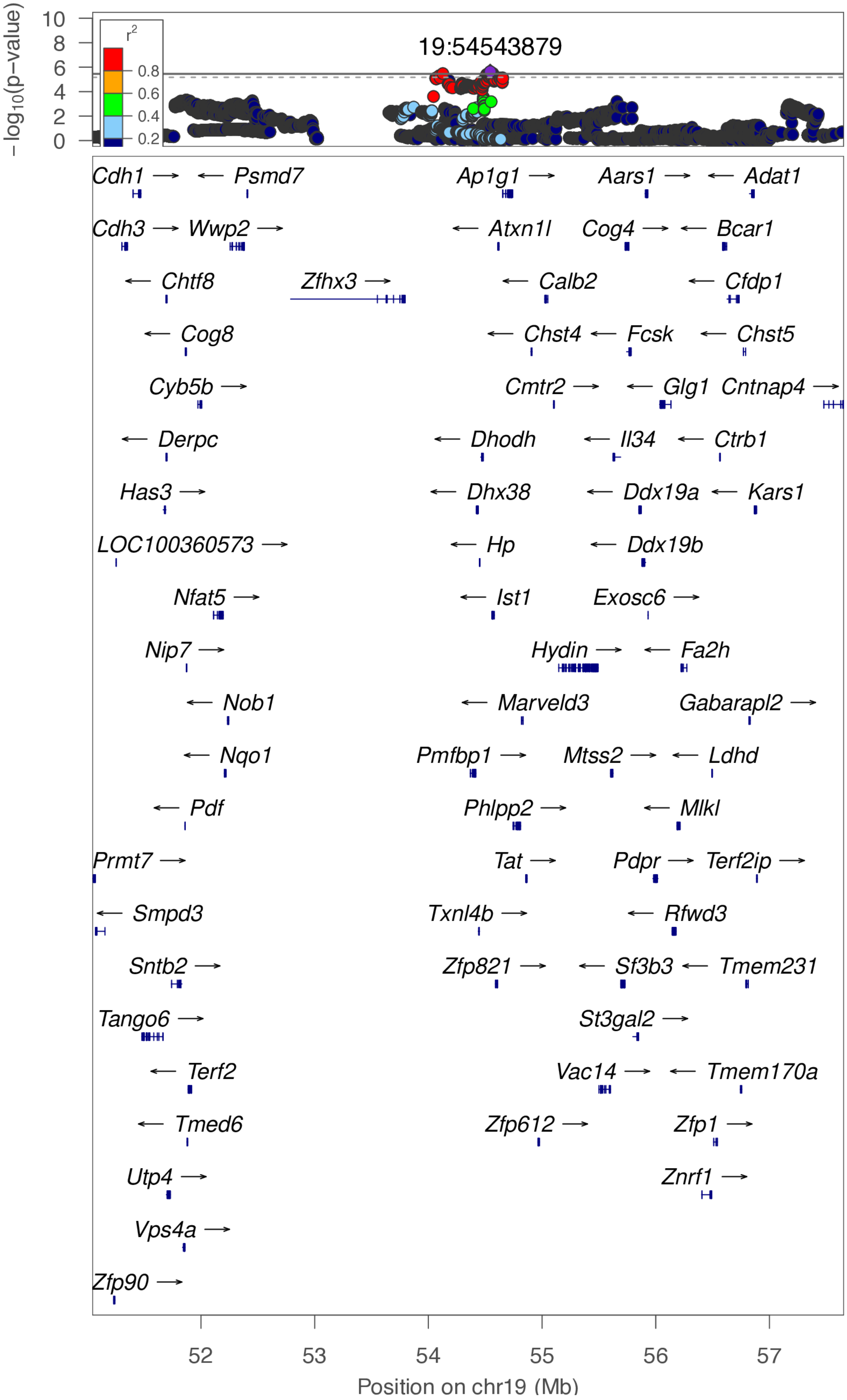

**chr20:16.6 Mb - Genomic SEM - narrow**

Lead: 20:16,638,901 | Genes in QTL interval: 1

QTL interval: chr20:16,331,360-16,641,441

Displayed region: chr20:16,131,360-16,841,441

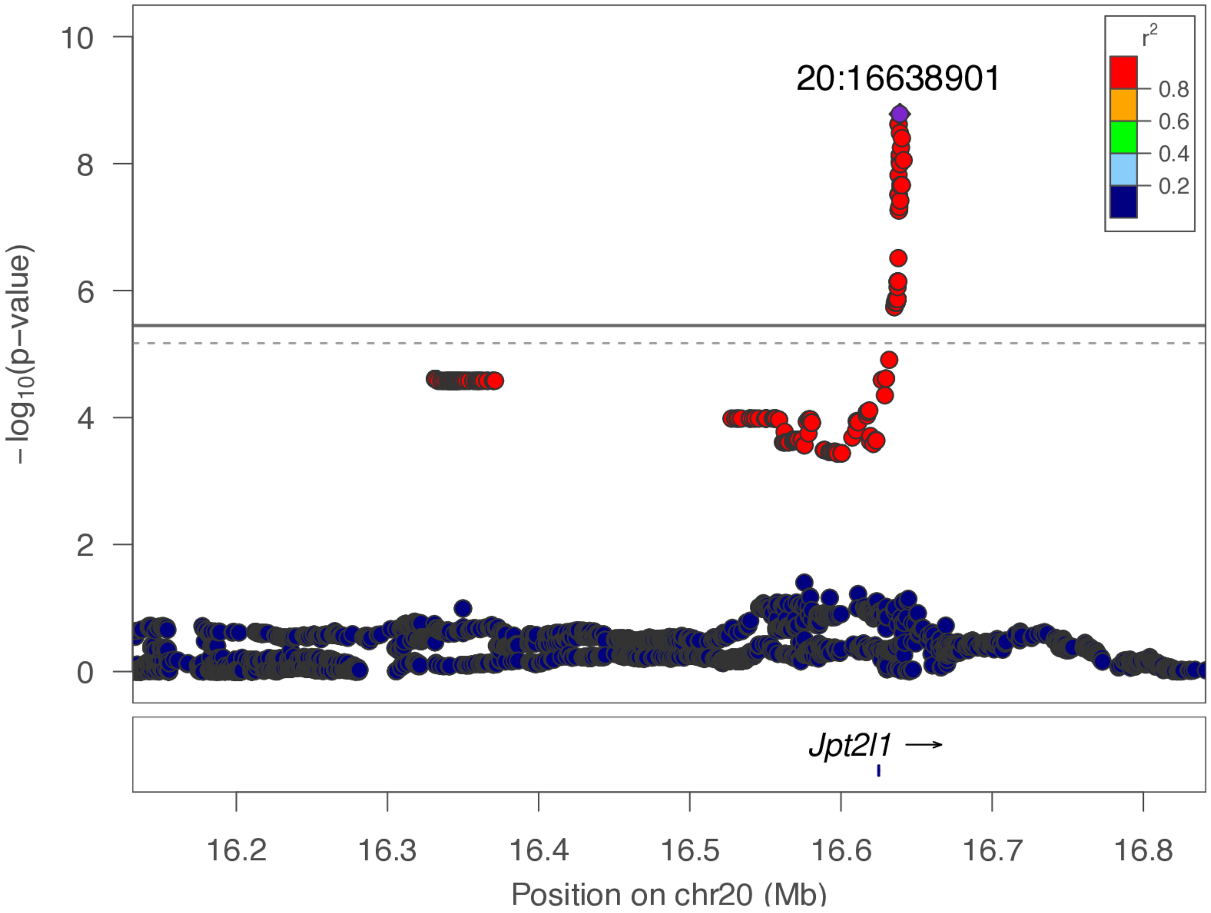

**chr20:16.6 Mb - Genomic SEM - broad**

Lead: 20:16,638,901 | Genes in QTL interval: 1

QTL interval: chr20:16,331,360-16,641,441

Displayed region: chr20:13,331,360-19,641,441

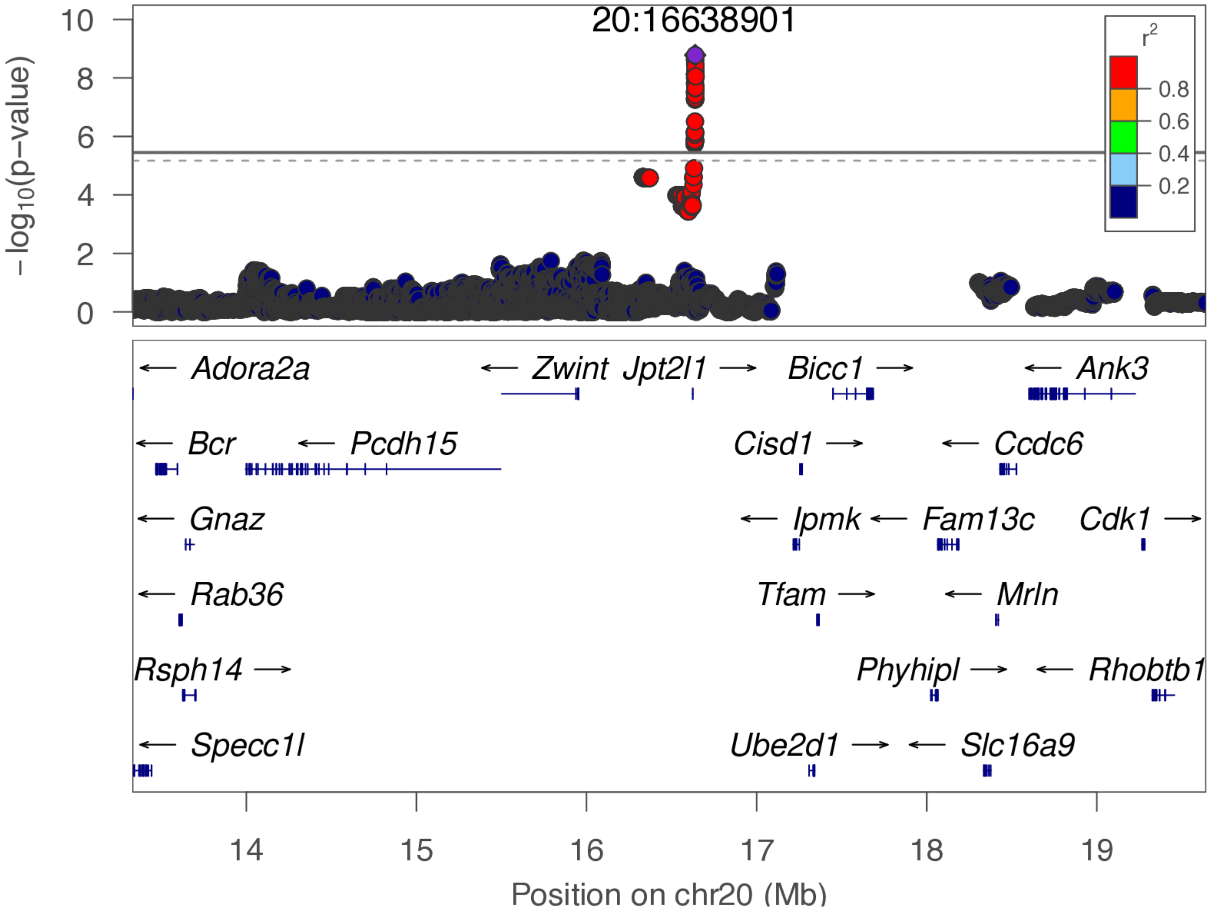

**chr20:54.7 Mb - Genomic SEM - narrow**

Lead: 20:54,711,825 | Genes in QTL interval: 4

QTL interval: chr20:54,546,255-56,013,408

Displayed region: chr20:54,346,255-56,213,408

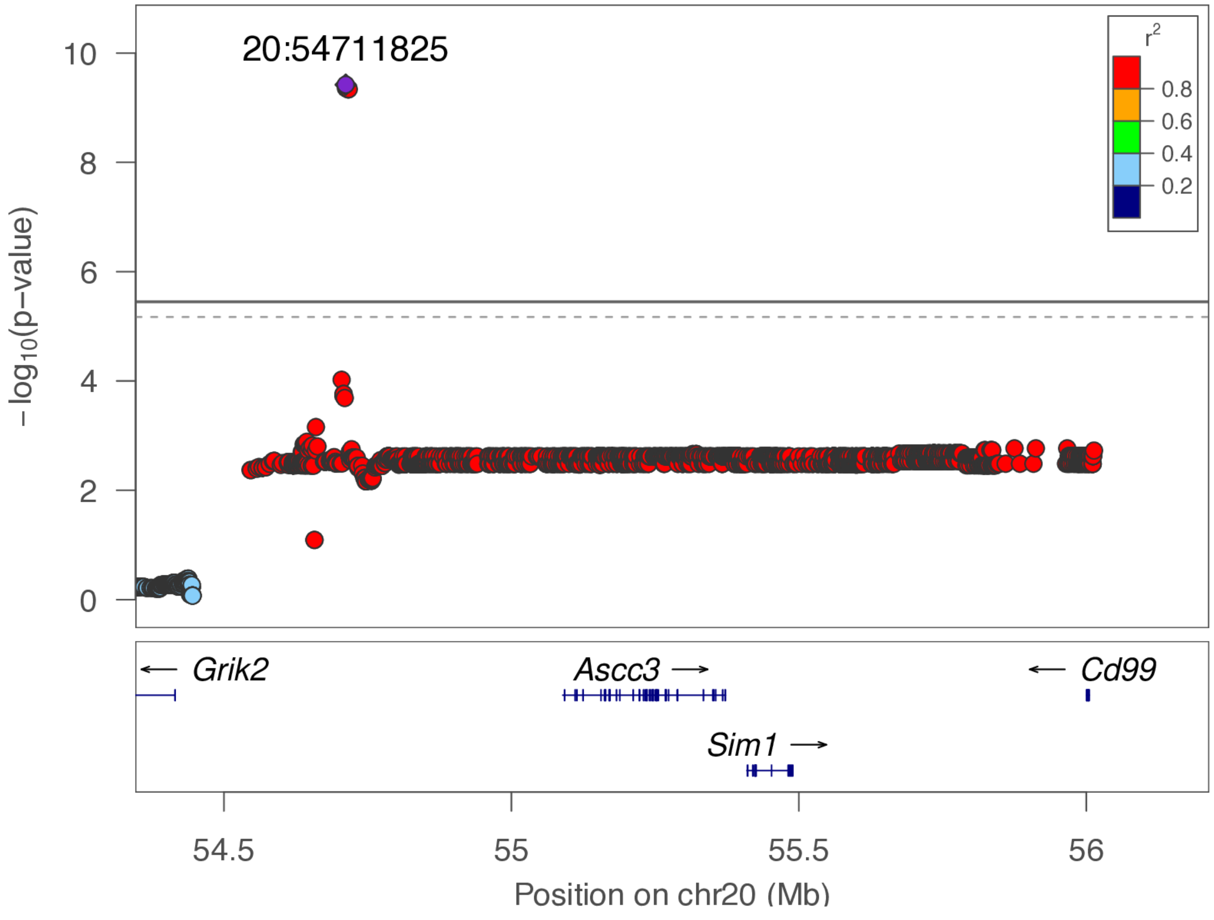

**chr20:54.7 Mb - Genomic SEM - broad**

Lead: 20:54,711,825 | Genes in QTL interval: 4

QTL interval: chr20:54,546,255-56,013,408

Displayed region: chr20:51,546,255-59,013,408

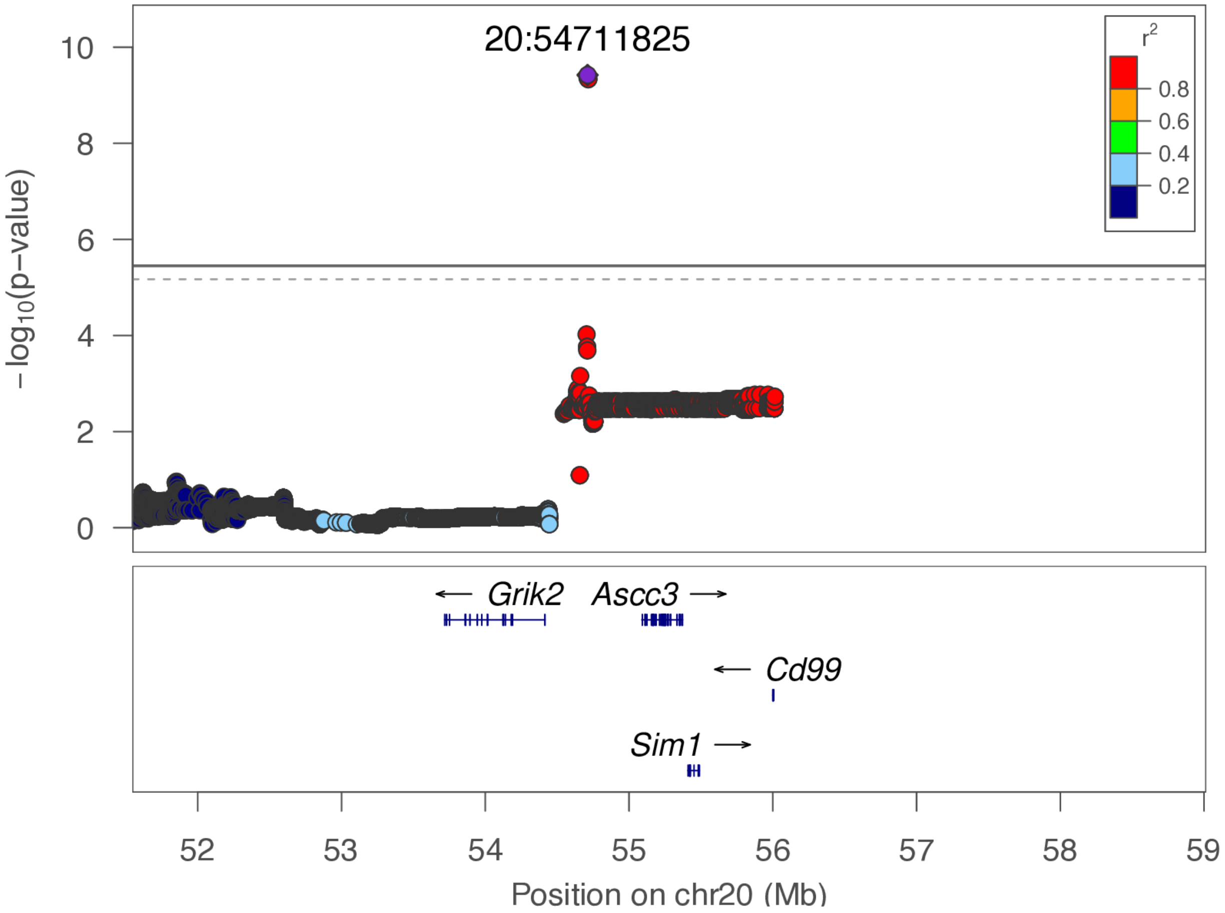

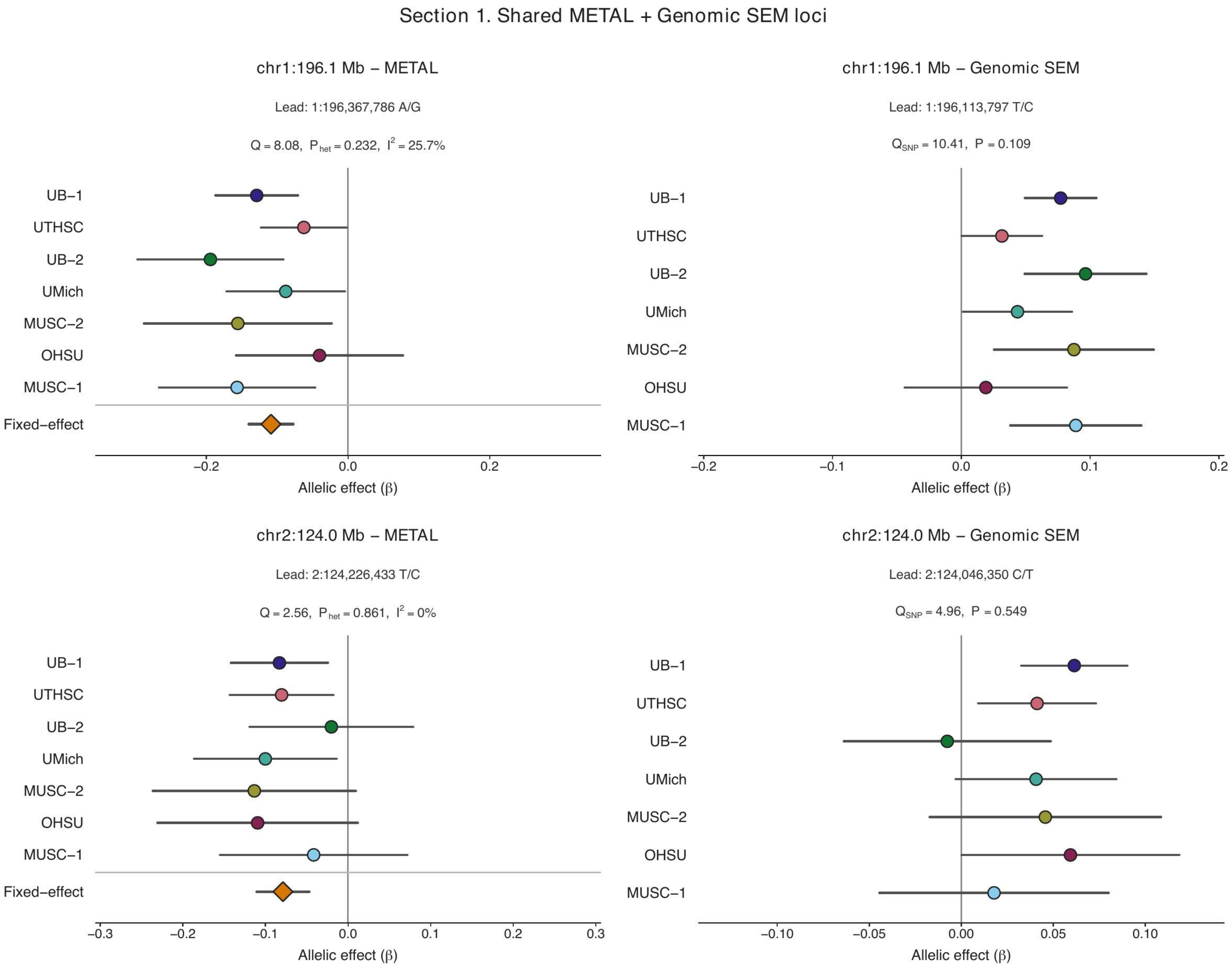


**Supplementary Figure 6. Cross-cohort effect structures at METAL- and Genomic SEM-supported loci.**

Forest plots show cohort-specific allelic-effect estimates ($\beta$) and 95% confidence intervals at each exact method-specific lead SNP. Loci supported by both methods are shown as paired METAL and Genomic SEM panels, followed by the remaining method-specific loci. For METAL, orange diamonds show the fixed-effect summary estimate; Cochran's $Q$, $P_{het}$, and $I^2$ summarize between-cohort heterogeneity. For Genomic SEM, $Q_{SNP}$ tests whether the cohort-specific SNP-effect pattern departs from that predicted by the selected factor model; no pooled summary estimate is shown. Where METAL and Genomic SEM select different lead SNPs within the same QTL, signed effects should not be compared directly without accounting for linkage-disequilibrium phase. Exact lead-SNP and allele information is provided above each panel.

### Section 1. Shared METAL + Genomic SEM loci

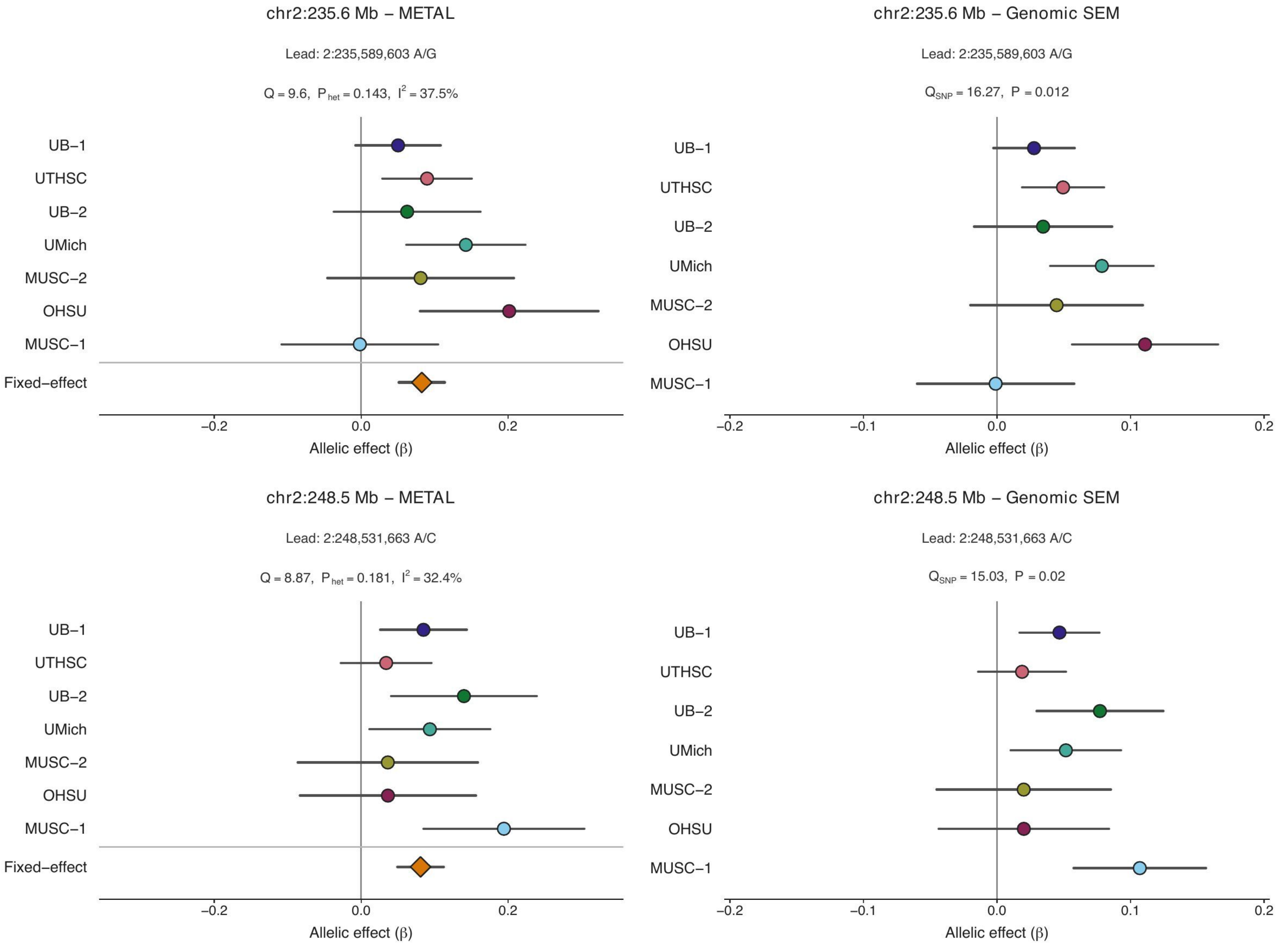

### Section 1. Shared METAL + Genomic SEM loci

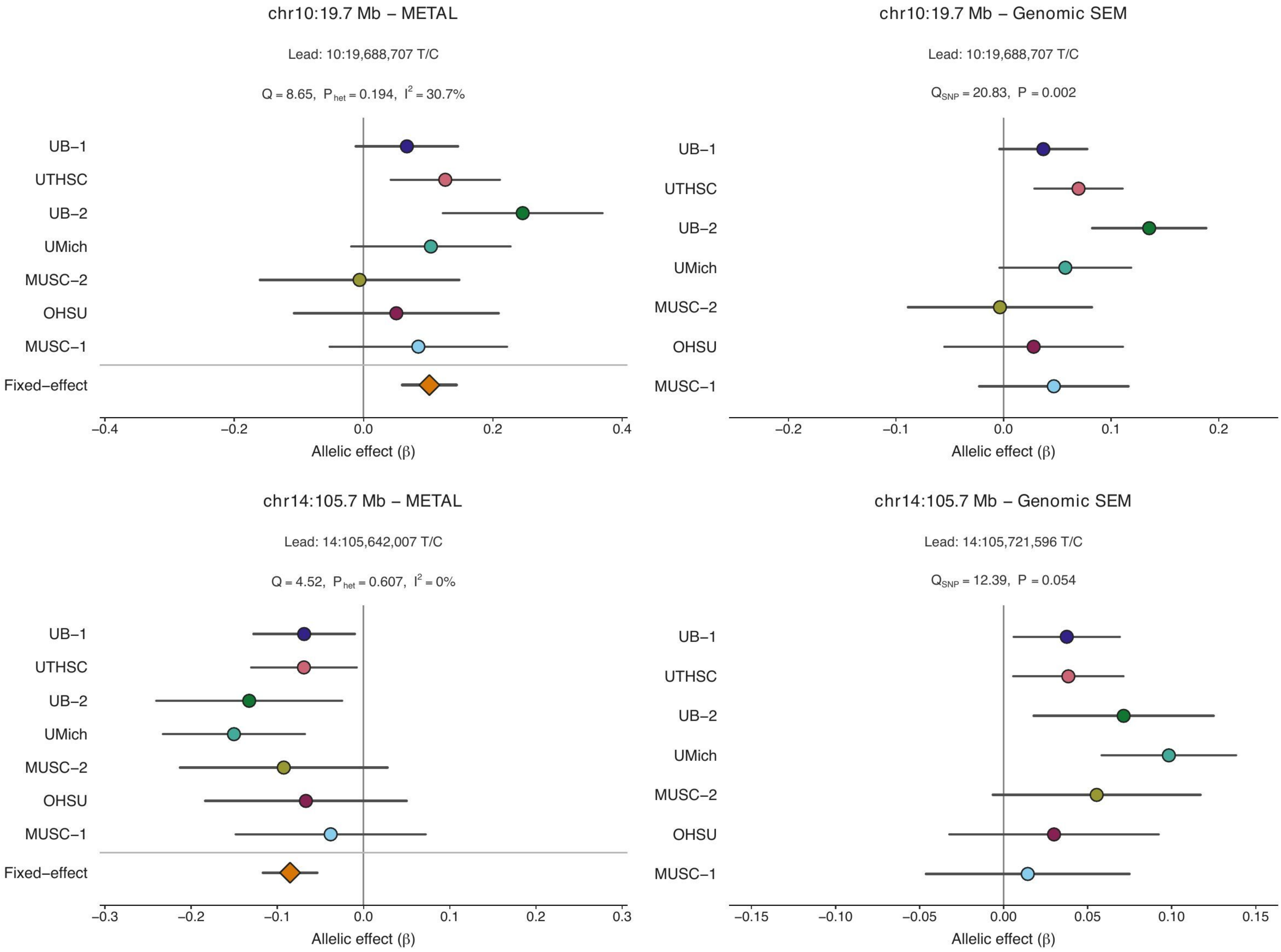

### Section 2. Remaining METAL–supported loci

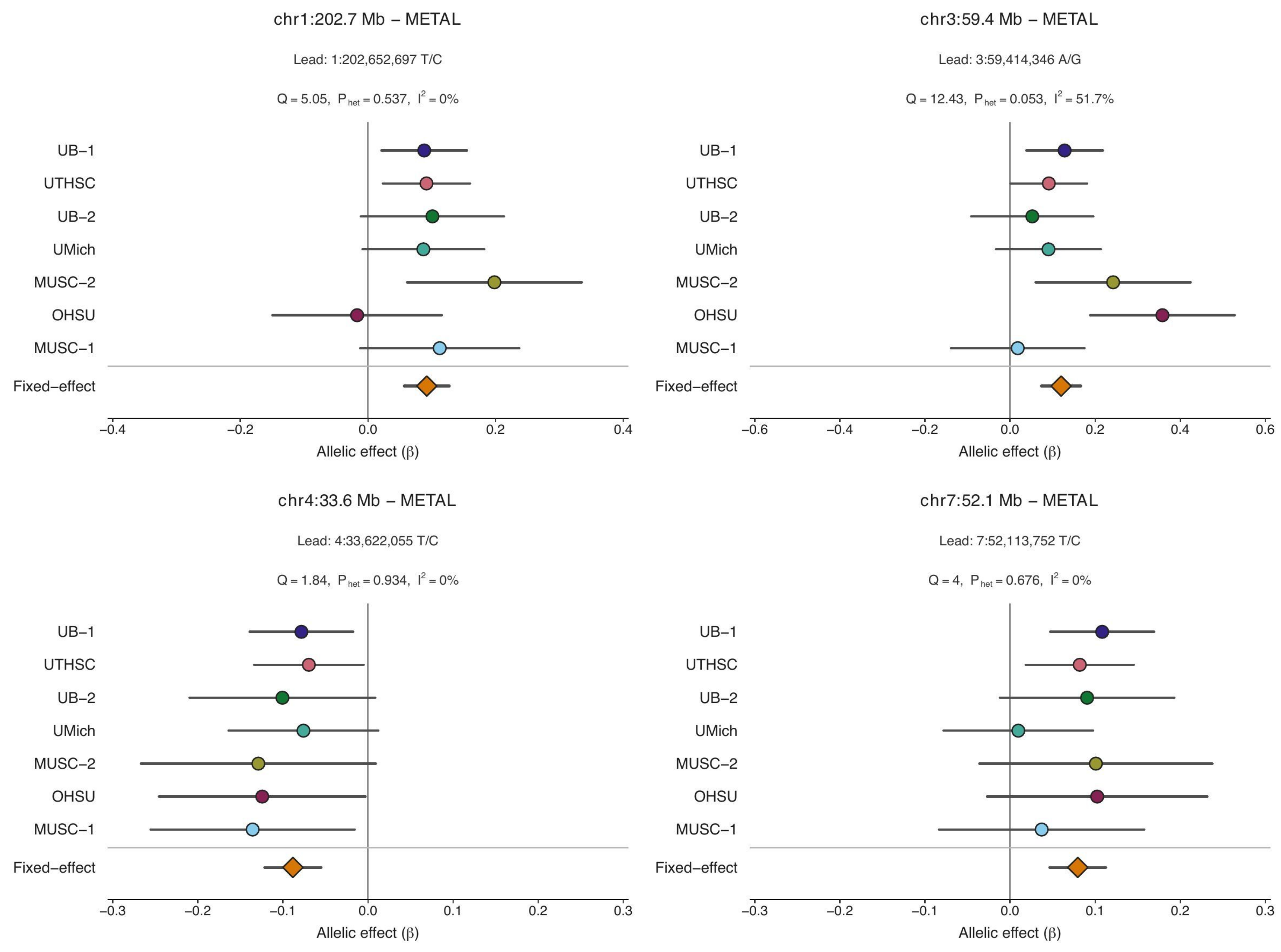

### Section 2. Remaining METAL–supported loci

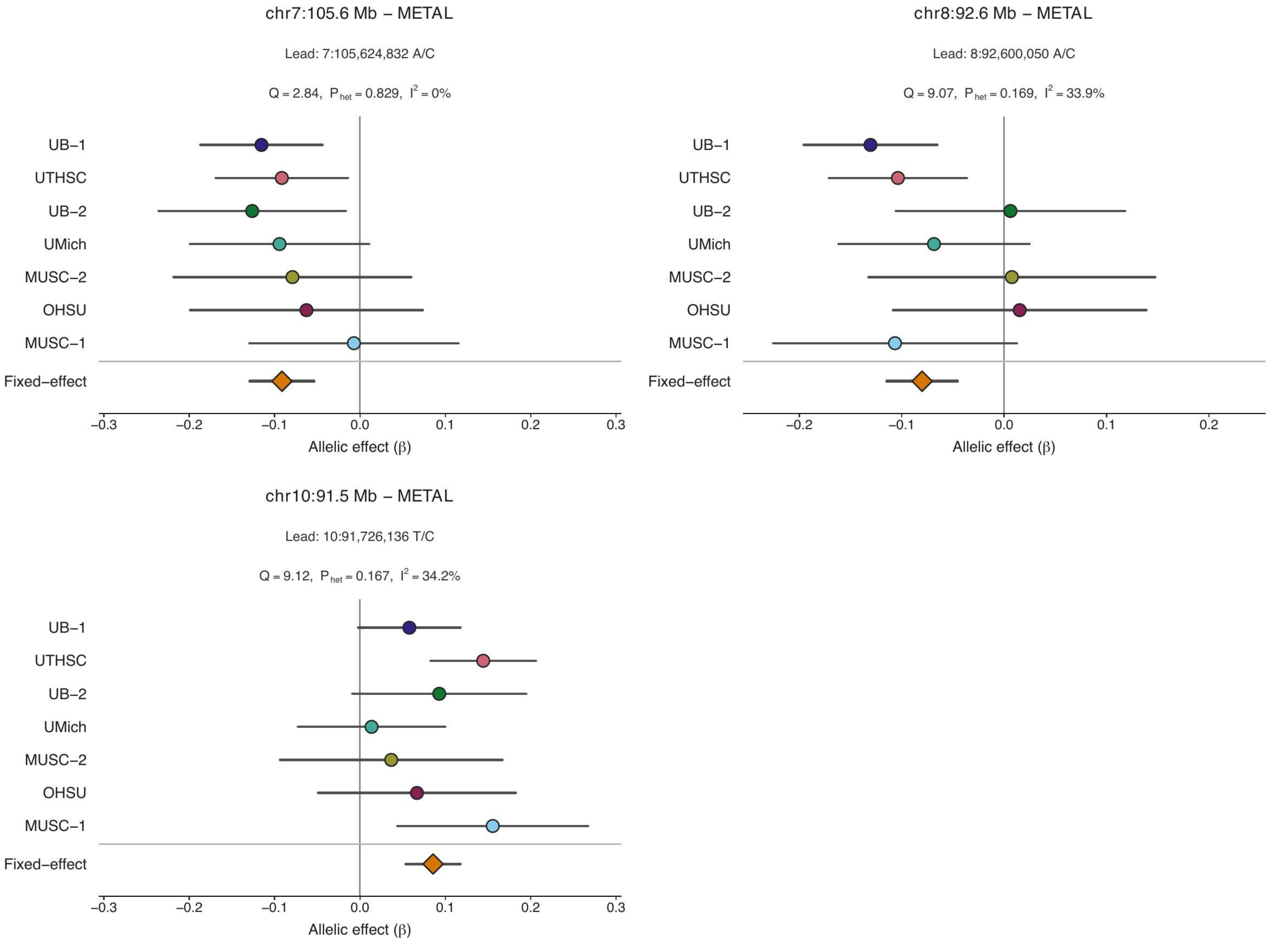

### Section 3. Remaining Genomic SEM–supported loci

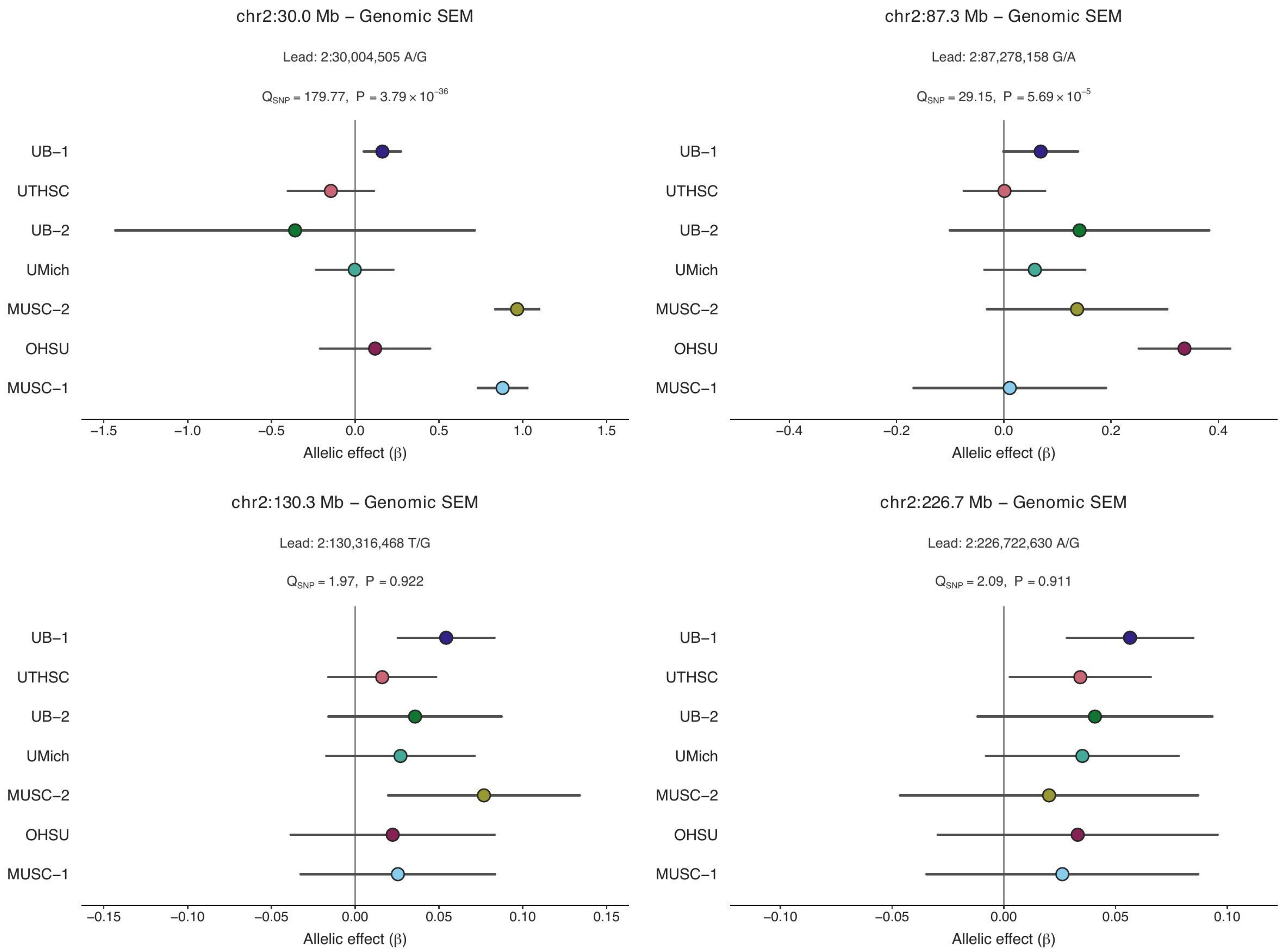

### Section 3. Remaining Genomic SEM–supported loci

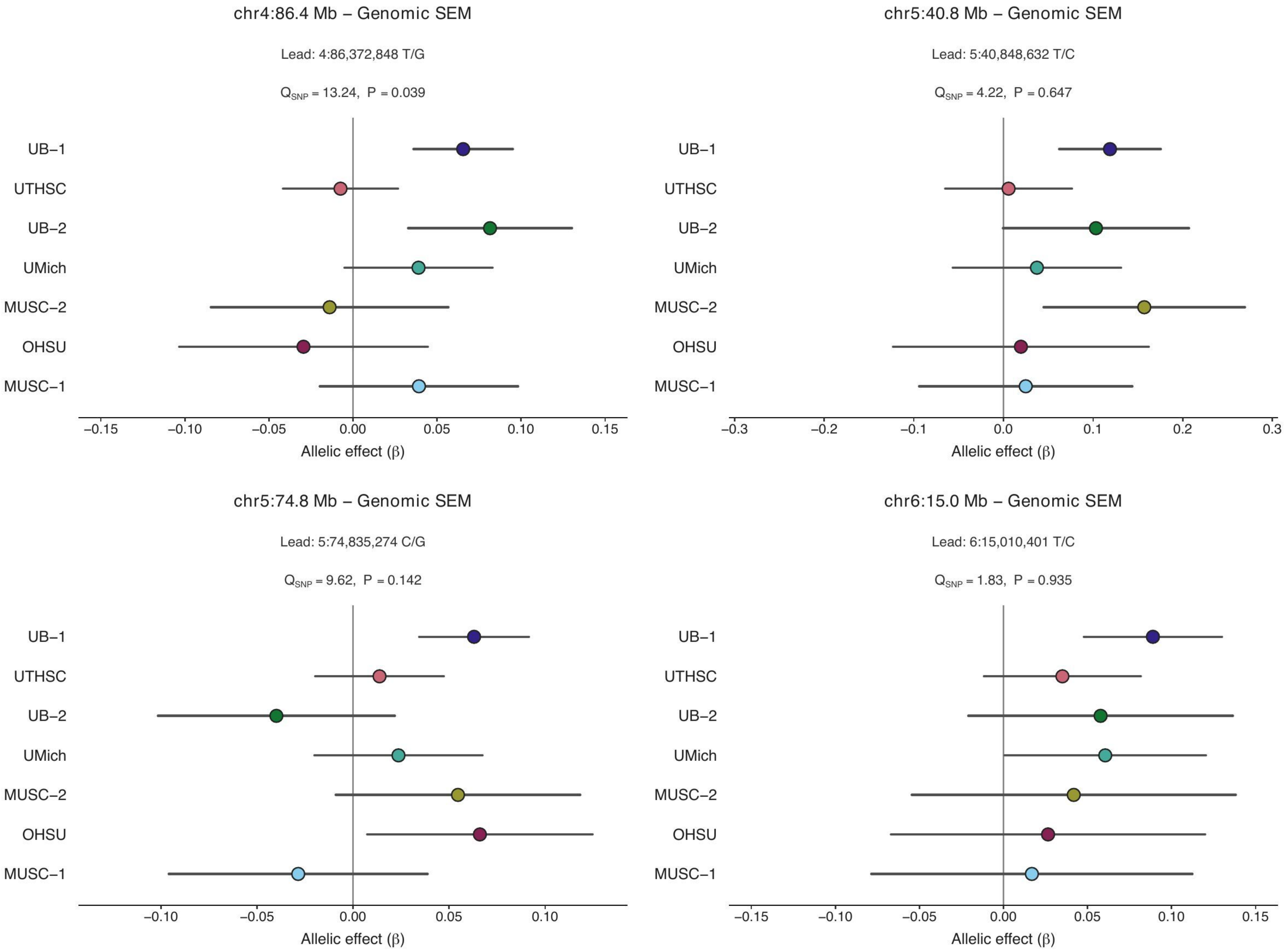

### Section 3. Remaining Genomic SEM–supported loci

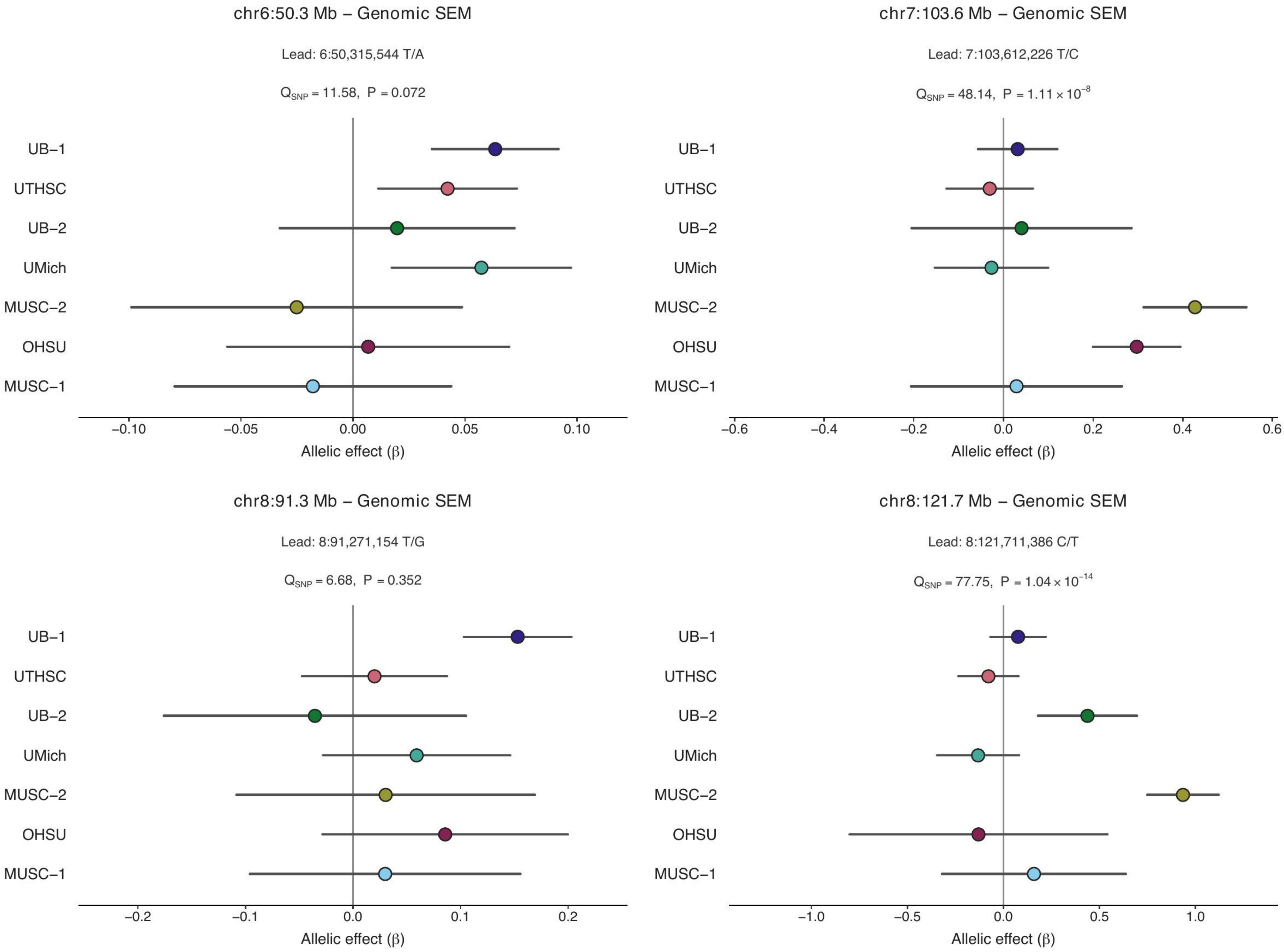

### Section 3. Remaining Genomic SEM–supported loci

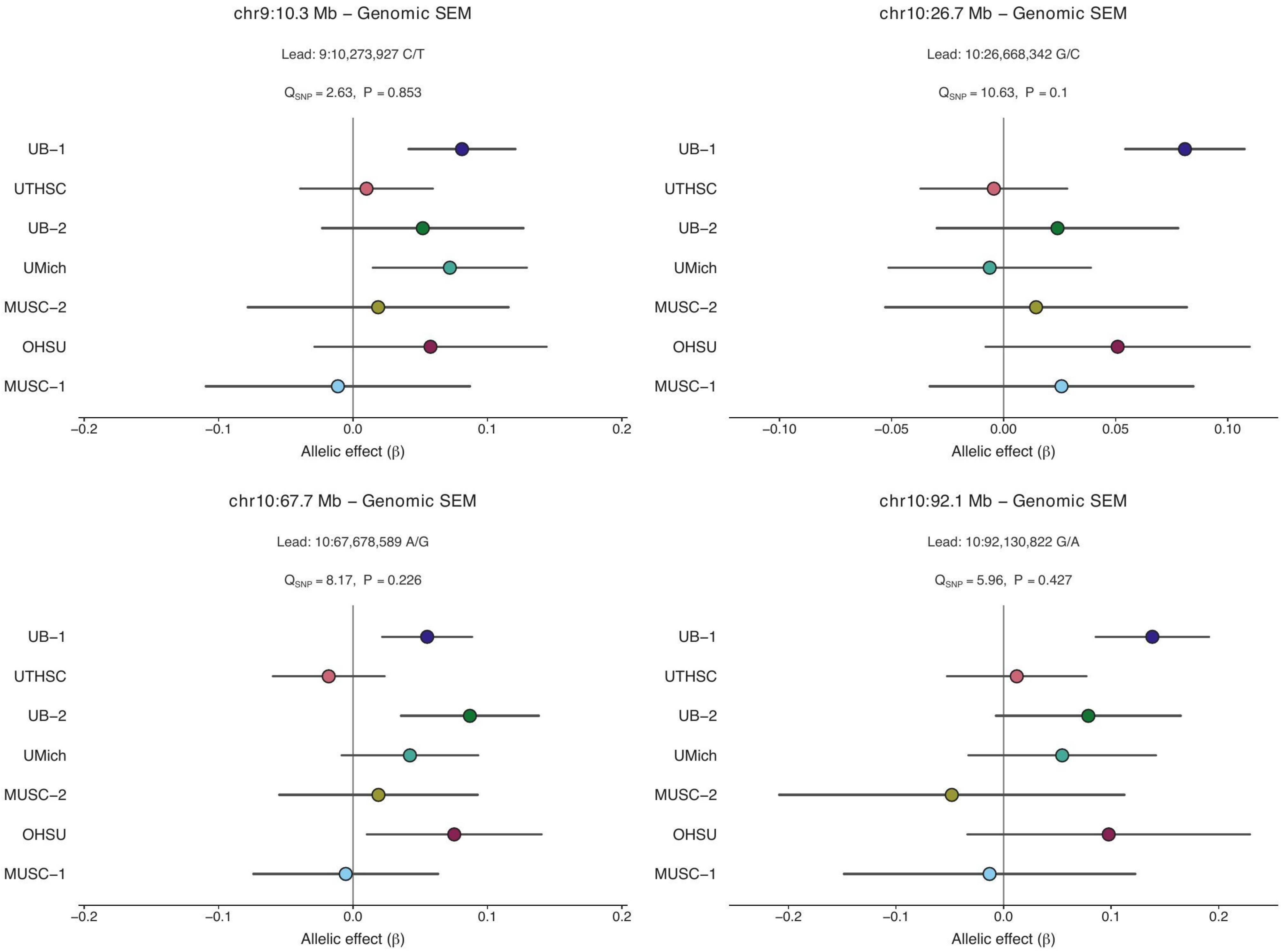

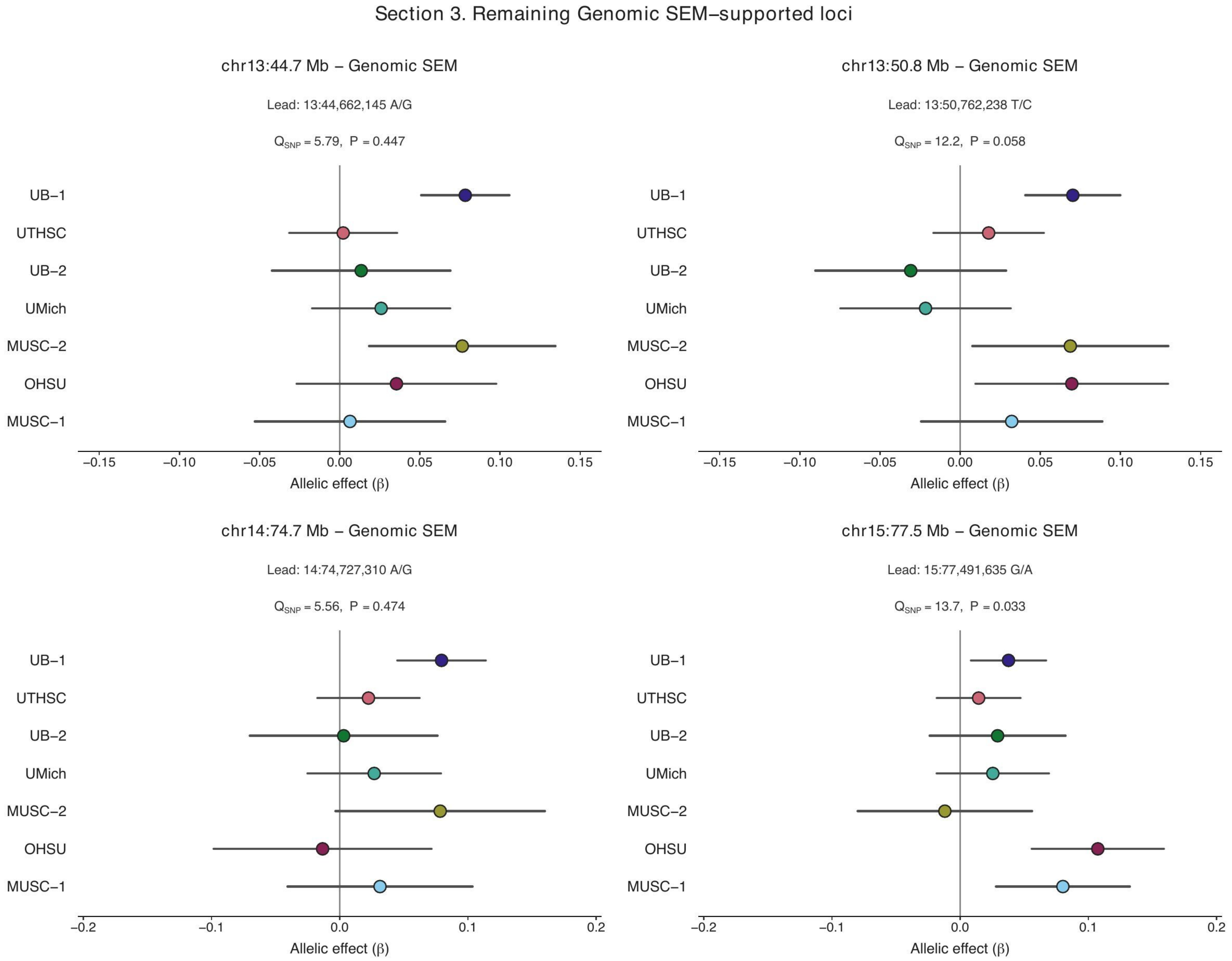
Section 3. Remaining Genomic SEM–supported loci
chr13:44.7 Mb – Genomic SEM
Lead: 13:44,662,145 A/G
$Q_{SNP}$ = 5.79, P = 0.447
chr13:50.8 Mb – Genomic SEM
Lead: 13:50,762,238 T/C
$Q_{SNP}$ = 12.2, P = 0.058
chr14:74.7 Mb – Genomic SEM
Lead: 14:74,727,310 A/G
$Q_{SNP}$ = 5.56, P = 0.474
chr15:77.5 Mb – Genomic SEM
Lead: 15:77,491,635 G/A
$Q_{SNP}$ = 13.7, P = 0.033
UB−1
UTHSC
UB−2
UMich
MUSC−2
OHSU
MUSC−1
Allelic effect (β)
−0.15
−0.10
−0.05
0.00
0.05
0.10
0.15
−0.2
−0.1
0.0
0.1
0.2

### Section 3. Remaining Genomic SEM–supported loci

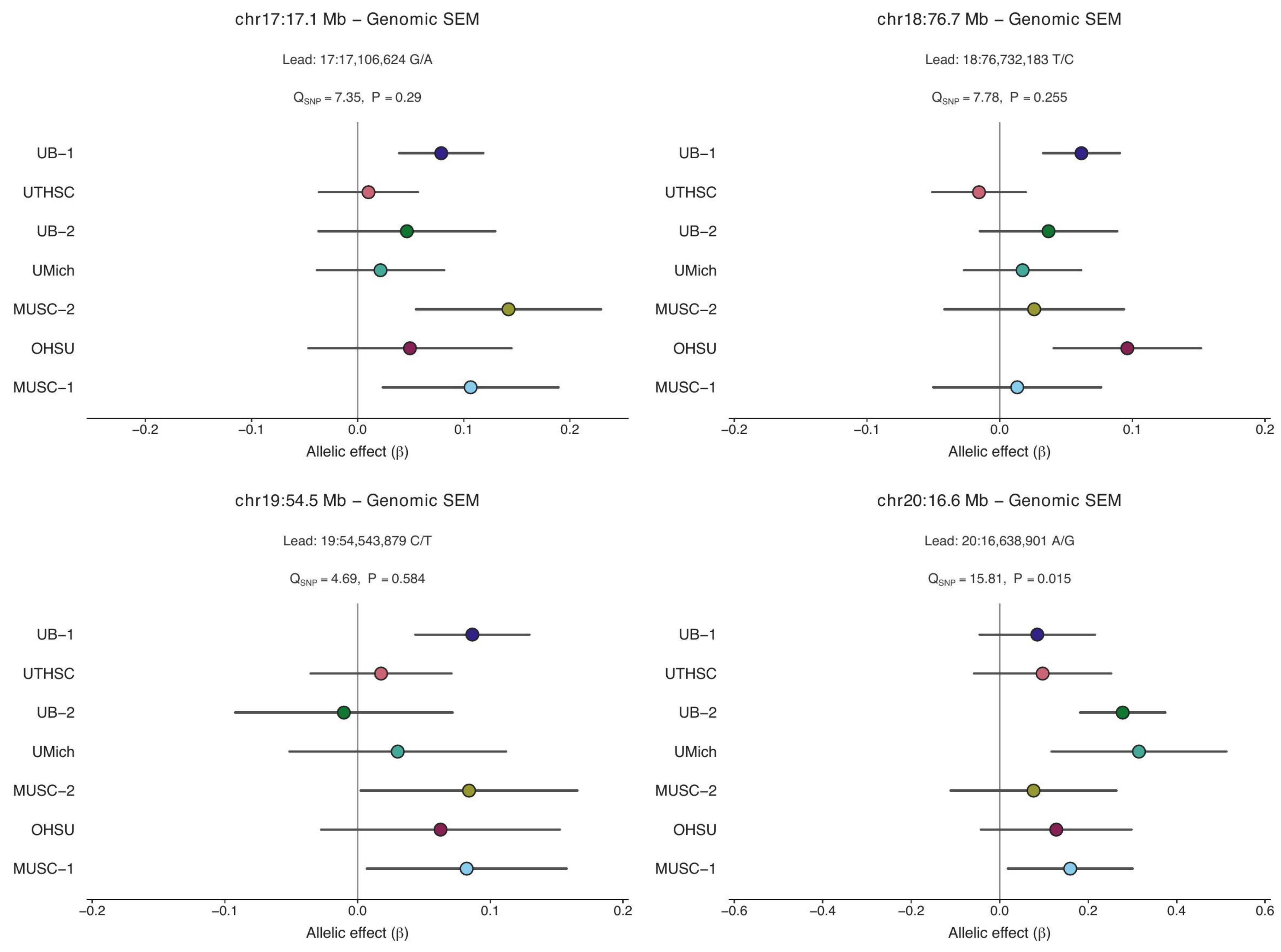

### Section 3. Remaining Genomic SEM–supported loci

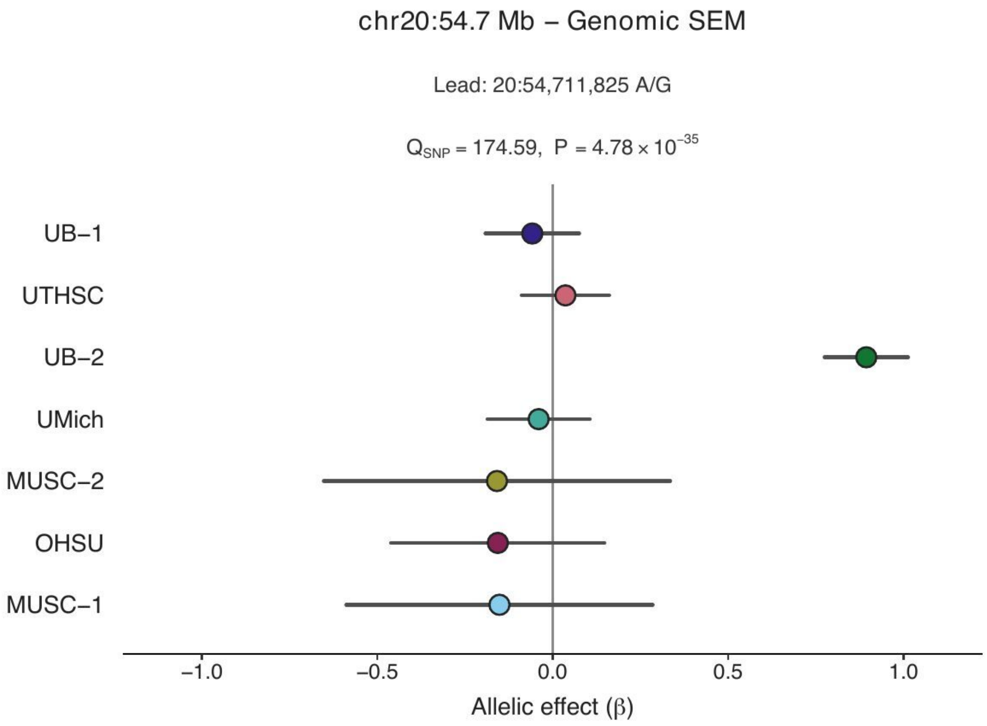
